# Evaluating molecular modeling tools for thermal stability using an independently generated dataset

**DOI:** 10.1101/856732

**Authors:** Peishan Huang, Simon K. S. Chu, Henrique N. Frizzo, Morgan P. Connolly, Ryan W. Caster, Justin B. Siegel

**Affiliations:** Biophysics Graduate Group, University of California, Davis, California, United States of America; Genome Center, University of California, Davis, California, United States of America; Microbiology Graduate Group, University of California, Davis, California, United States of America; Department of Biochemistry & Molecular Medicine, University of California, Davis, California, United States of America; Department of Chemistry, University of California, Davis, California, United States of America

**Keywords:** Thermal stability, Rosetta Molecular Modeling Suite, molecular modeling, Protein Engineering

## Abstract

Engineering proteins to enhance thermal stability is a widely utilized approach for creating industrially relevant biocatalysts. Computational tools that guide these engineering efforts remain an active area of research with new data sets and develop algorithms. To aid in these efforts, we are reporting an expansion of our previously published data set of mutants for a β-glucosidase to include both measures of T_M_ and ΔΔG, to complement the previously reported measures of T_50_ and kinetic constants (*k*_cat_ and *K*_M_). For a set of 51 mutants, we found that T_50_ and T_M_ are moderately correlated with a Pearson correlation coefficient (PCC) of 0.58, indicated the two methods capture different physical features. The performance of predicted stability using five computational tools are also evaluated on the 51 mutants dataset, none of which are found to be strong predictors of the observed changes in T_50_, T_M_, or ΔΔG. Furthermore, the ability of the five algorithms to predict the production of isolatable soluble protein is examined, which revealed that Rosetta ΔΔG, ELASPIC, and DeepDDG are capable of predicting if a mutant could be produced and isolated as a soluble protein. These results further highlight the need for new algorithms for predicting modest, yet important, changes in thermal stability as well as a new utility for current algorithms for prescreening designs for the production of soluble mutants.

## INTRODUCTION

A common goal of enzyme engineering is the enhancement of thermal stability.^1^ For industrial applications improving a proteins’ robustness to thermal challenges or half-life at elevated temperature can often be the deciding factor for the commercialization of a biocatalyst.^2–5^ Currently, the most common approach for improving thermal stability is through directed evolution methodologies,^6,7^ which, can be time-consuming, costly, and limited in the ability to search sequence space. Computational design tools to predictably identify mutations, single and combinatorial that enhance thermal stability are rapidly developing and growing in popularity.^8–11^ However, accurate predictions using computational tools to guide protein stability design remains an active area of research and is not always successful.

The use of large data sets on the mutational effect on protein stability, such as ProTherm, is often used to train computational methods for predicting thermal stability. The data sets utilized generally composed of the equilibrium constant of unfolding (*K*_U_) or the melting temperature of an enzyme (T_M_).^12^ In our previous study, we determined the thermal stability of 79 β -glucosidase B (BglB) variants by finding T_50_ – a type of kinetic stability that is determined by the temperature at which a mutant’s residual activity is reduced by 50% after a heat-challenge over a defined time.^4,12,13^ When analyzing this set of mutants using two established computational programs (Rosetta ΔΔG and FoldX PSSM) for predicting thermal stability, we found that there was no significant correlation between the predictions and the observed T_50_.^14^

One hypothesis explaining the poor predictive performance of the algorithms with the BglB dataset is that the algorithms are trained on T_M_, a direct measure of structural thermal stability, but being used to predict T_50_, an indirect measure of the protein’s thermal stability.^12^ Alternatively, the poor performance could have come from the narrow T_50_ range (extreme variants are +6.06 °C and -5.02 °C from wild type), as the algorithms are generally trained on larger changes in thermal stability and ± 5 °C may be within the error of the currently developed algorithms. In this study, we evaluated both hypotheses. To assess if there is a significant difference in T_M_ and T_50_ we developed a data set of 51 BglB mutants (Figure 1) in which both thermal stability measurements, T_50_ and T_M_, are measured.

**Figure 1.**
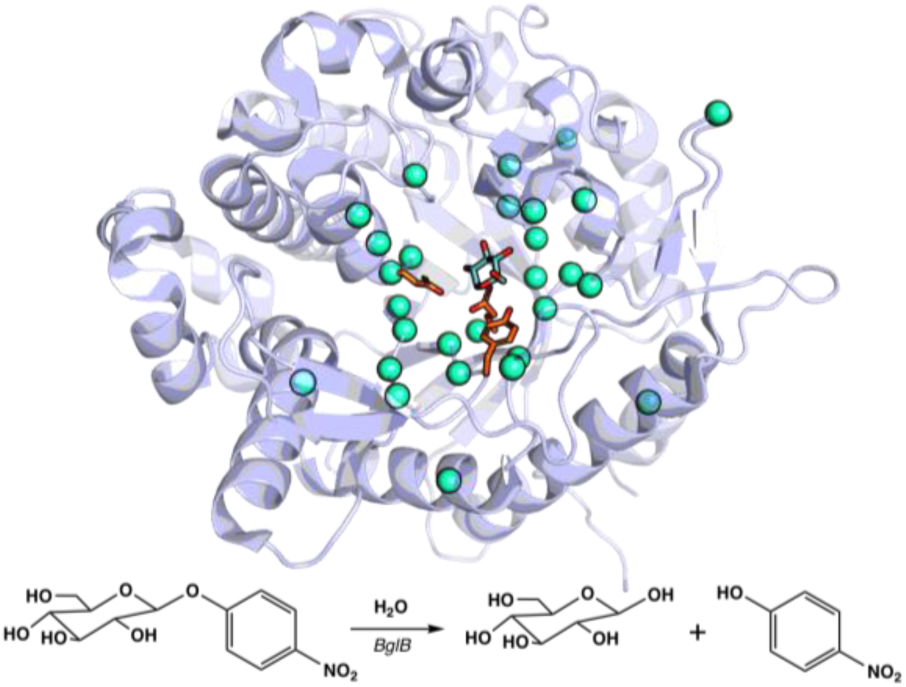
Structure of BglB (PDB ID: 2JIE) from bacterium *Paenibacillus polymyxa.* PyMOL rendering^15^ of BglB showing the 28 sequence-positions (teal spheres) of the 51 mutants randomly selected for T_M_ analysis. The reaction scheme of the hydrolysis of 4-nitrophenyl β-D-glucopyranoside by BglB used in the T_50_ study.^14^

Interestingly, for the set of 51 measurements, there is only a modest correlation between T_50_ and T_M_, with a PCC of 0.58. This highlights the difference in the physical properties being measured using these two techniques, T_M_ being the thermal stability of the protein’s structural elements and T_50_ reporting on the thermal stability to irreversible denaturation. However, similar to the previous study,^14^ the relationship between the predicted stability with the experimental T_M_ only results in a weak correlation not only with the previous algorithms evaluated (Rosetta ΔΔG and FoldX PSSM) but also three other commonly used methods: ELASPIC, DeepDDG, and PoPMuSiC. This result suggests that while the two measurements are reporting on different physical properties, this is not the key factor that led to the low predictive accuracy of established algorithms on this data set.

To evaluate the second hypothesis, that the changes in thermal stability of the BglB data set are too small for current algorithms, we investigated the ability of the algorithms to predict soluble protein production. Mutations that reduce thermal stability (T_M_) by > 18 °C also fail to yield soluble, isolatable protein. Analysis of computational algorithms enriched for insoluble mutants based on the predicted energetics showed a significant enrichment was observed for Rosetta ΔΔG. This supports the hypothesis that the lack of performance on the BglB data set is due to the narrow range in thermal stability changes observed. This highlights the need for new algorithms for predicting modest, yet important, changes in thermal stability as well as a new utility for current algorithms for prescreening designs for the production of soluble protein.

## METHODS

### Mutant Selection, protein expression, and purification

Out of 79 mutants of Beta-glucosidase B (BglB) that were previously characterized with T_50_ data,^14^ 51 variants with plasmid readily available were transformed into chemically competent *E. coli* BLR(DE3) cells. The variants were produced and purified as previously described.^14^ Expression was carried out by growing a 5mL overnight culture in a 50 mL falcon tube with breathable seal in Terrific broth (TB) medium with kanamycin while shaking at 250 rpm at 37 °C. After the initial overnight culture cells were spun down and resuspended in fresh TB with kanamycin with 1mM Isopropyl β-D-1-thiogalactopyranoside (IPTG) in a 50mL falcon tube with breathable seal and incubated while shaking at 250 rpm at 18 C for 24 hours. Then the cells were then spun down, lysed, and purified samples using immobilized metal ion affinity chromatography as previously described.^14^ The purity of the protein samples was analyzed using 12-14% SDS-PAGE (SI 1-1), the yield was assessed based on the A280 for proteins that appeared >75% pure in the SDS-PAGE analysis.

### Melting Temperature assay

The melting temperature (T_M_) of BglB was determined using the Protein Thermal Shift (PTS) ™ kit (Applied BioSystem ®, from Thermo Fisher). Standard protocols provided by the manufacturer were used. Protein concentrations ranged from 0.1-0.5 mg/mL and fluorescence reading was monitored with a QuantaStudio™ 3 System from 20 °C to 90 °C. The temperature melting curve was first smoothened with a 20-step sliding window average (SI 4). T_M_ was determined from the average of 4 replicates at which the derivative was largest, and all melting curves can be found in SI 2.

### ΔΔG Calculations from T_M_

Calculations were carried out in a similar manner as previously described.^15^ Briefly, to derive ΔG°_unfolding_ the fluorescence intensity was first translated into fraction of folded (P_f_) and unfolded (P_u_) proteins at different temperatures starting from the minimum fluorescence (*F*_*min*_) to the maximum fluorescence (*F*_*max*_) shown in eq 1.

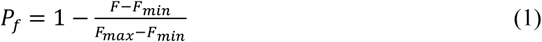

By taking a two-state folding-unfolding model, the equilibrium constant of unfolding (*K*_u_) at different temperatures is then given by:

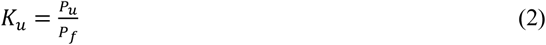

By plotting ln *K*_u_ against 1 / *T* using Van’t Hoff method shown in eq 3 (SI 2), the − ^Δ*H*^/_*R*_ is defined by the slope, ^Δ*S*^/_*R*_ is the y-intercept, T is temperature, and *R* is the ideal gas constant.

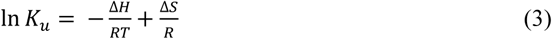

The Gibbs free energy of protein-unfolding (ΔG°_unfolding_) can then be determined using eq 4, where ΔG°_unfolding_ is the unfolding energy at 298K as T. All calculations can be found under SI 4.

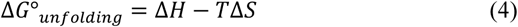

### Molecular modeling

Five popular, readily accessible, and recently developed force-field and machine-learning-based molecular modeling methods were evaluated for their ability to recapitulate the experimental data: Rosetta ΔΔG^16^, FoldX^17^, ELASPIC^18^, DeepDDG^19^, and PoPMuSiC.^20^ The crystal structure of BglB (PDB ID: 2JIE) was used across five different algorithms. First, using a previously described method^14^, the 2JIE structure was used as input to the Rosetta ΔΔG application and run as previously described (SI 5). Briefly, 50 poses of the wild type and the mutant were generated for which 15 energy terms are reported from the score function used.^16^ The three lowest system energy scores out of 50 from WT and mutant were averaged to give the final Rosetta ΔΔG score.

Second, for the FoldX position-specific scoring metric (PSSM) protocol, the 2JIE structure was first minimized for any potential inaccurate rotamer assignment using the RepairPDB application.^17^ The repaired PDB structure was mutated with single-point mutants and then modeled using FoldX PSSM. The model was scored based on 17 terms within FoldX force-field.^17^ Third, the ELASPIC protocol first constructs a homology model of the wild type using the crystal structure, sequence, molecular, and energetics information. Using the standard procedure described, FoldX algorithm was used to construct the mutant model. Combining the energetic information from FoldX along with 74 other features, Stochastic Gradient Boosting of Decision Trees are used to make the final prediction of the mutational change.^18,21^ Fourth, using a curated dataset derived from Protherm database, DeepDDG used their previously described Shared Residue Pair (SRP) neural network structure to make stability prediction.^19^ Lastly, the fifth method evaluated is PoPMuSiC where the stability of the wild type structure and mutant were estimated using 13 statistical potential terms and additional two terms that count for the volume differences of the residues between WT and the mutant.^20^

A Pearson correlation (PCC) analysis was performed between the change in total system energy (ΔTSE) of the five computational methods and the two thermal stability methods (T_M_ and T_50_). Additionally, the available individual features within the Rosetta ΔΔG and FoldX PSSM force field were further evaluated against the T_M_ dataset for correlation.

Finally, the ΔTSE was also evaluated against isolated soluble protein and un-isolated insoluble protein for its statistical significance. Using the student T-test, the p-values for the five computational methods were obtained by treating the two categories between isolated soluble and insoluble protein as samples that are independent of each other with the variance being treated as unequal.

## RESULTS

### Evaluating the Relationship between T_M_ and T_50_

To the best of our knowledge, there has not been a large data set (>50 data points) directly comparing the T_M_ and T_50_ relationship for a single set of protein mutants produced and characterized uniformly. It is important to distinguish both T_M_ and T_50_ methods since the measurements are quantifying and reporting different structural and functional properties. T_M_ is defined by the temperature at which half the enzyme is found in the unfolded state over folded state^12,22^ and is often evaluated through denaturation assays, from which the thermodynamic measurements (ΔG_unfolding_) can be obtained.^22^ This method is generally a lower throughput method as purified protein is required to get an accurate measurement for the structural properties for the mutant being evaluated. T_50_ measures the temperature of half-inactivation that leads to irreversible unfolding^11,23^, and it is determined by the reduction of half of the enzymatic activity due heat-challenges.^12^ This is a very common assay for protein engineering due to its compatibility with high throughput assays and the ability to use cell lysates to evaluate function.

To complement our previously measured dataset of T_50_, 51 proteins were selected randomly and evaluated for T_M_ using the Protein Thermal Shift™ assay in order to compare T_50_ and T_M_. The wild type BglB T_M_ is 45.97 ± 1.03 °C, meanwhile the previously determined T_50_ is 39.9 ± 0.1 °C.^14^ When evaluating the entire data set, the T_M_ range is between 37.1 and 54.3 °C, slightly larger than what was observed for T_50_ which is between 34.9 and 46.0 °C (SI 1-2). The highest T_M_ in this data set is E167A, 54.3 °C (+8.33 °C), which is also observed to have a similar increase in T_50_ from wild type (+6.06 °C).^14^ The variant that has the lowest T_M_ in this dataset is found to be E225A, with a ΔT_M_ of -8.9 °C, which has corresponding T_50_ of -3.1 °C.

The relationship between T_50_ and T_M_ is plotted in Figure 2A. The Pearson coefficient correlation (PCC) of 0.58 shows the two methods are moderately positively correlated. Correlation between methods increased in cases where mutations resulted in extremely stable and unstable products, for example E167A and E225A, respectively. This is an expected result as for small changes (<3 °C) in thermal stability; the differences in Measurement methods would be expected to play a more significant role than for larger changes (>5 °C).

**Figure 2.**
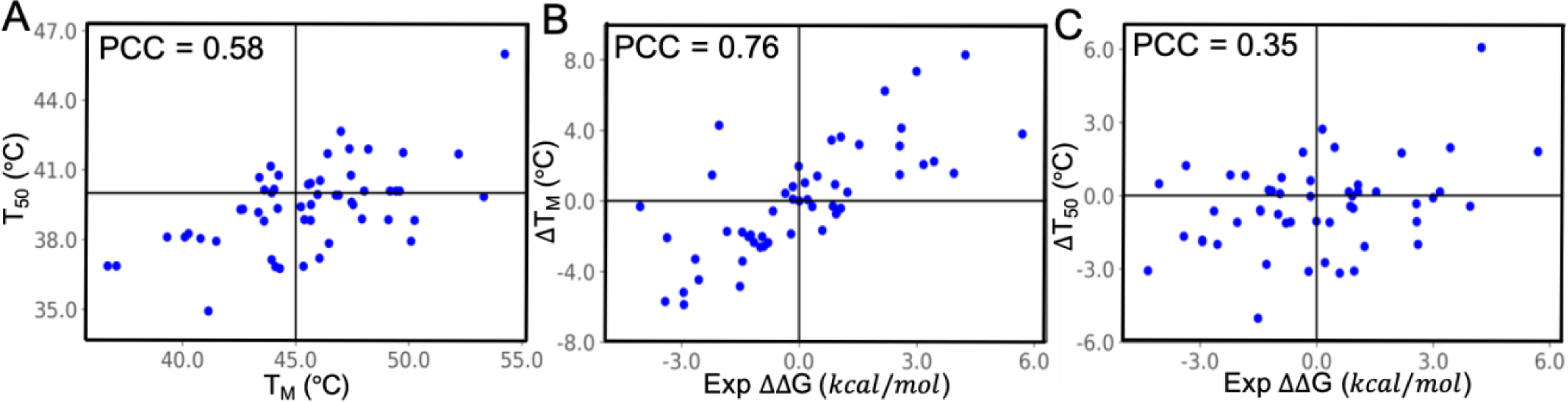
Comparison of two different experimental thermal stability dataset and experimentally-derived ΔΔG. (A) The relationship for each mutant between T_50_ and T_M_. The PCC of 0.58 illustrates the two methods are modestly positively correlated with mutations that are in the extreme ends of the temperature range (± 5 °C) are generally in agreement. (B) The evaluation of ΔT_M_ with experimentally-derived ΔΔG shows the two qualities are highly correlated (PCC = 0.76) unlike (C) the relationship between ΔT_50_ and experimentally-derived ΔΔG with a PCC of 0.35.

### Evaluating computational stability tools using the BglB T_M_ dataset

The computational evaluation of protein stability of the current experimental T_M_ dataset was analyzed in the same manner as our previous study did for T_50_.^14^ An energetically evaluated model for each mutant was generated using established computational methods, and subsequently plotted as a function of T_M_ to evaluate the calculated energies related to the observed T_M_. The PCC for the most commonly evaluated term, the difference in total system energy of the mutants versus the wild type protein (ΔTSE), was found to be highest for FoldX PSSM (PCC = -0.34) with ΔT_M_ (Fig 3). Similarly, the FoldX PSSM correlations with experimentally-derived ΔT_50_ data is found to be -0.21. The overall relationship between the ΔTSE and the current thermal stability dataset slightly improved for FoldX, DeepDDG, and PoPMuSiC, while Rosetta ΔΔG and ELASPIC remained relatively unchanged with no significant correlation.

**Figure 3.**
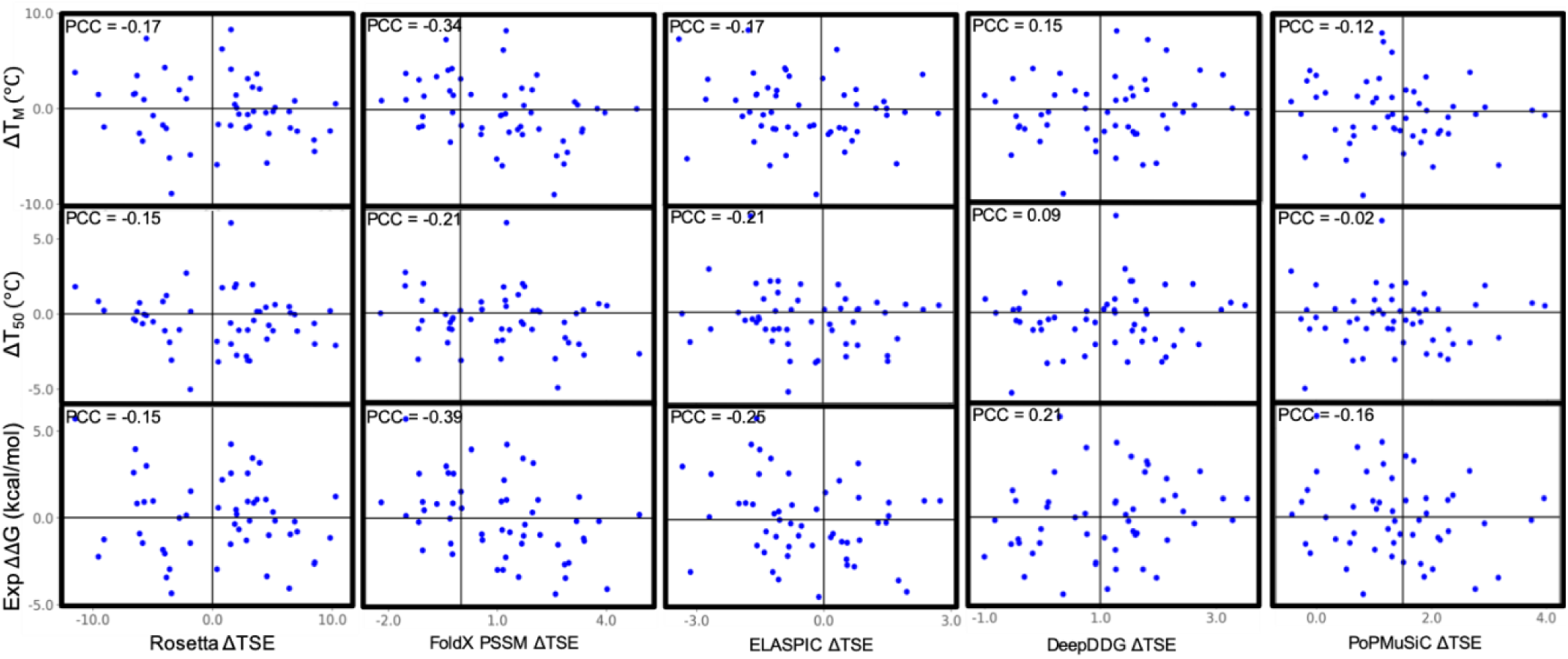
Evaluation of the five algorithms ΔTSE versus the experimentally-derived ΔΔG, and T_M_ and T_50_ dataset. The Pearson correlation for each performance against three types of experimental data was analyzed. No algorithm resulted in a significant correlation between the calculated energies and the observed T_M_, T_50_, or ΔΔG for this dataset.

An analysis of each energetic term from Rosetta ΔΔG and FoldX PSSM did not uncover any specific parameter in either method’s energetic evaluation that was strongly correlated with the T_M_ data set, as was previously observed for the T_50_ data set^14^ (SI 3). The strongest PCC for T_M_ against any of the available energetic terms is 0.39 for the Δbackbone clash term from FoldX PSSM and -0.31 for the Omega energy term from Rosetta ΔΔG. Moreover, since the previous performance of the algorithms were evaluated using experimentally-derived ΔΔG in kcal/mol, the same evaluation was done by converting the T_M_ dataset into ΔΔG (Fig 3). The PCC of -0.39 between experimental ΔΔG and ΔTSE for FoldX PSSM outperformed four other algorithms that were compared. The correlation between experimental ΔΔG with ΔTSE was not unexpected as T_M_ is relatively correlated to ΔΔG with a PCC of 0.76 (SI 1-3).

Based on this analysis, it is apparent that the general performance of all given methods only weakly correlates with the experimentally determined effects of the mutations. This data fails to support the hypothesis that the lack of a previously observed correlation of these established computational tools with observed changes in thermal stability in the BglB data set is due to the difference in the physical property being measured.

### Protein expression prediction

The current data set consists primarily of modest thermal stability changes of <5 °C, calculated to be ± 4 kcal/mol of the native, and therefore may be challenging for current computational methods to predict. However, this change is only analyzing a fraction of the 129 mutants tested in the overall BglB data set. Of the 129 mutants, only 92 were found to be produced and isolated in a soluble form. As all purification procedures are conducted at room temperature, with the wild type T_50_ of 39.9 °C, any reduction of >18 °C would result in denatured, insoluble protein. Therefore, it seemed pertinent to evaluate if any of the five algorithms could differentiate variants in this data set that could be isolated as soluble protein versus those that were not able to be separated as soluble protein.

For this evaluation, all the previous 129 mutants reported were assessed using Rosetta ΔΔG, FoldX PSSM, ElASPIC, DeepDDG, and PoPMuSic in the same manner as carried out for T_50_ and T_M_. A mutant was generally considered soluble if it was observed on an SDS-PAGE analysis and had an A280 of >0.1 mg/ml. The wild type protein produced using the methods described generally resulted in an average A280 of 1.5 mg/ml, which would provide a >10-fold change in yield for mutants having an A280 less than 0.1 mg/ml. While factors other than thermal stability can affect production and isolation of soluble protein, it is assumed in this case that the primary factor that decreases soluble protein yield is from denaturation of the mutant protein either during expression or purification. The results of this analysis are presented in Figure 4.

**Figure 4.**
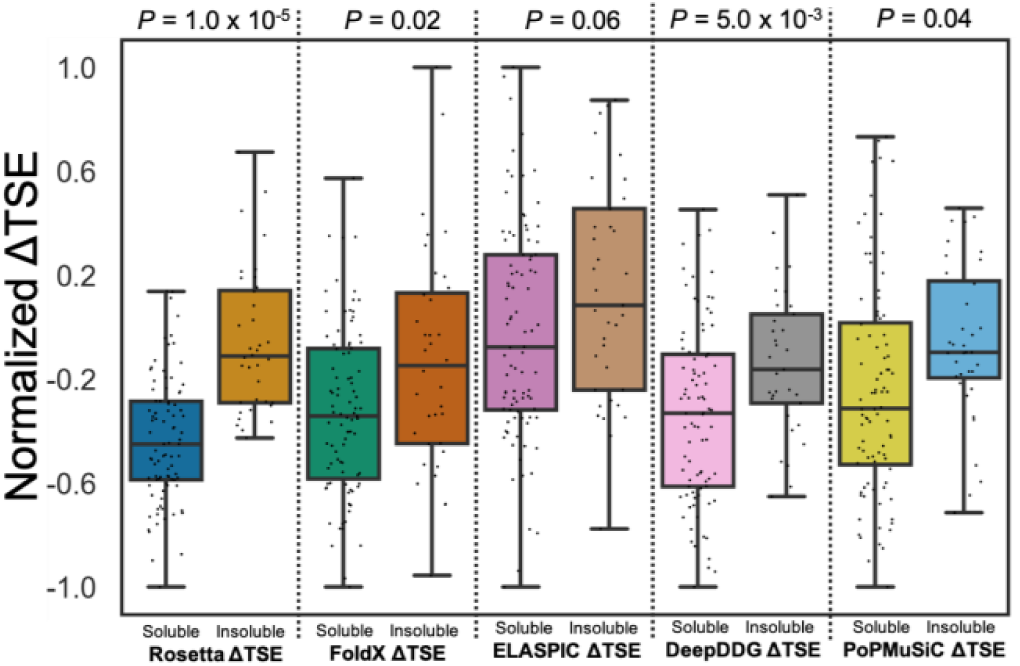
Computational prediction for the effect on mutant soluble protein production using five different algorithms. From left to right: Rosetta ΔΔG, FoldX PSSM, ELASPIC, DeepDDG, and PoPMuSiC of soluble and insoluble protein. In this case mutants that resulted in a significant (>10-fold) decrease in yield of purified soluble protein are considered insoluble. Significance in population differences is done using the Student’s t-test.

Of the five algorithms evaluated, Rosetta ΔΔG, ELASPIC, and DeepDDG can capture the enrichment of mutants isolated as a soluble protein. The differences were evaluated for statistical significance using the student T-test, and the highest among the top three methods was shown in Rosetta ΔΔG with a *p*-value of 1.0 × 10^−5^. In contrast, enrichment was lower for FoldX PSSM and PoPMuSiC with a *p*-value of 0.02 and 0.04, respectively.

A few outliers were observed in all methods except for ELASPIC (SI 1-4). For example, the mutant G15N for both Rosetta ΔΔG and FoldX PSSM was identified as severely energetically unfavorable, which is consistent with the observation that this variant was not able to be isolated as a soluble protein.

## DISCUSSION

Both T_M_ and T_50_ are methods commonly used to quantify different physical aspects of protein thermal stability, however there is relatively little information reported on the relationship of these two measurements. Using a data set of 51 protein mutants we observed that there is a moderate positive correlation (PCC of 0.58) between these two properties. Mutants with extreme stability changes, such as E164A (>6 °C), usually exhibit similar magnitude in T_M_ and T_50_ results. However, the majority of the mutants exhibit a change of 3 °C or less in this T_M_ and T_50_ data set being analyzed, in which range the effect of experimental variability may be significant, therefore analysis with larger data sets with more extreme stability changes may reveal an even stronger correlation of these two properties.

Consistent with our previous analysis, we found a lack of performance using established computational tools when predicting T_M_ and T_50_ from the wild type for this data set. According to Jia et al, stability prediction using the experimentally-derived free energy change of unfolding ΔΔG (Δ kcal/mol) outperforms the prediction using ΔT_M_ (Δ Celsius).^24^ However, in this case, we saw no significant change in predictive performance for all five computational tools when comparing to the experimentally-derived free energy change. In addition, we found that T_M_ and ΔΔG to be strongly correlated with this data set, which may suggest that the improved performance is only relevant for more diverse datasets composed of different proteins as opposed to mutants of a single protein.

While none of the computational methods demonstrated a strong predictive power for the mutants in this study, Rosetta ΔΔG, FoldX PSSM, ELASPIC, and PoPMuSiC all have previously been shown to have high correlations with experimental data (PCC between 0.69 to 0.83).^16,20,21,25^ A major factor that could differentiate this study from previous work is that this data set has an experimental ΔΔG range of ± ∼4 kcal/mol, which is significantly smaller than what the algorithms were usually trained on, which ranged from +8 to - 5 kcal/mol.^17^ This is further supported by the observation that Rosetta ΔΔG does have a strong predictive performance for our data set when including mutants presumed to have large effects on T_M_ and T_50_, because they failed to be isolated as soluble protein due to the substantial stability changes.

This study highlights the need for new computational tools that can more accurately predict modest changes in thermal stability, rather than major changes. This becomes particularly important because single-point mutants often increase thermal stability by a few degrees at a time, while major changes are more often produced from the synergistic effect of combining multiple mutations.^11,26–28^ Furthermore, as larger data sets of protein mutants with explicitly measured biophysical properties are generated, opportunities to explore combinations of molecular modeling and machine learning methods will become practical. These algorithms and data sets will enable the development of robust predictors of thermal stability.

## ASSOCIATED CONTENT

### Supporting Information (SI)

SI 1-1. SDS-PAGE images for 51 BglB mutants and WT. (PDF)

SI 1-2. A distribution analysis of temperatures observed for T_M_ and T_50_.

SI 1-3. PPC graph between ΔT_M_ and ΔΔG of BglB mutants. (PDF)

SI 1-4. Evaluation of five computational methods on protein expression. (PDF)

SI 2. Images of T_M_ fluorescence graphs, derivative graphs, and Van’t Hoff plot for 51 mutants and WT. (Zip)

SI 3. Rosetta ΔΔG and FoldX PSSM correlations graphs with ΔT_M_ and experimental ΔΔG. Excel files of all the parameters from data acquisition. Excel files of the total system energy for DeepDDG, ELASPIC, and PoPMuSiC (Zip)

SI 4. Jupyter notebook for all thermal stability data acquisitions with all T_M_ raw data files. (ipynb)

SI 5. Example files for Rosetta_ddg_monomer run. (Zip)

## AUTHOR INFORMATION

### Author Contributions

Data curation: PH, SKSC, HNF

Investigation: PH, SKSC

Methodology: PH, SKSC, MPC, RWC, JBS

Software: SKSC

Writing – original draft: PH

Writing – review & editing: PH, JBS, SKSC, MPC

## ACKNOWLEDGMENT

This work was supported by the University of California Davis, the National Institutes of Health [R01 GM 076324-11], the National Science Foundation [Award Numbers 1827246, 1805510, 1627539], and the National Institute of Environmental Health Sciences of the National Institutes of Health [Award Number P42ES004699]. The content is solely the responsibility of the authors and does not necessarily represent the official views of the National Institutes of Health, National Institute of Environmental Health Sciences, National Science Foundation, or UC Davis.

