## Supplementary material for "Evaluating molecular modeling tools for thermal stability using an independently generated dataset": SI_1

### Evaluating the relationship between $T_{50}$ and $T_M$ for an enzyme mutant library

Table of Contents:

1-1 SDS-PAGE Images for 51 mutants and WT Page 2

1-2 A distribution analysis of temperatures observed for $T_M$ and $T_{50}$ Page 2

1-3 Pearson correlation coefficient of $\Delta T_M$ and experimental $\Delta\Delta G$ Page 3

1-4 Evaluation of five computational methods on protein expression. Page 3

SI 2 file description Page 4

SI 3 file description Page 4

SI 4 file description Page 4

SI 5 file description Page 4

Supporting information 1-1. 12-14% SDS-PAGE for 51 BglB mutants and wild type (WT) using Protein Plus Kaleidoscope as ladder.

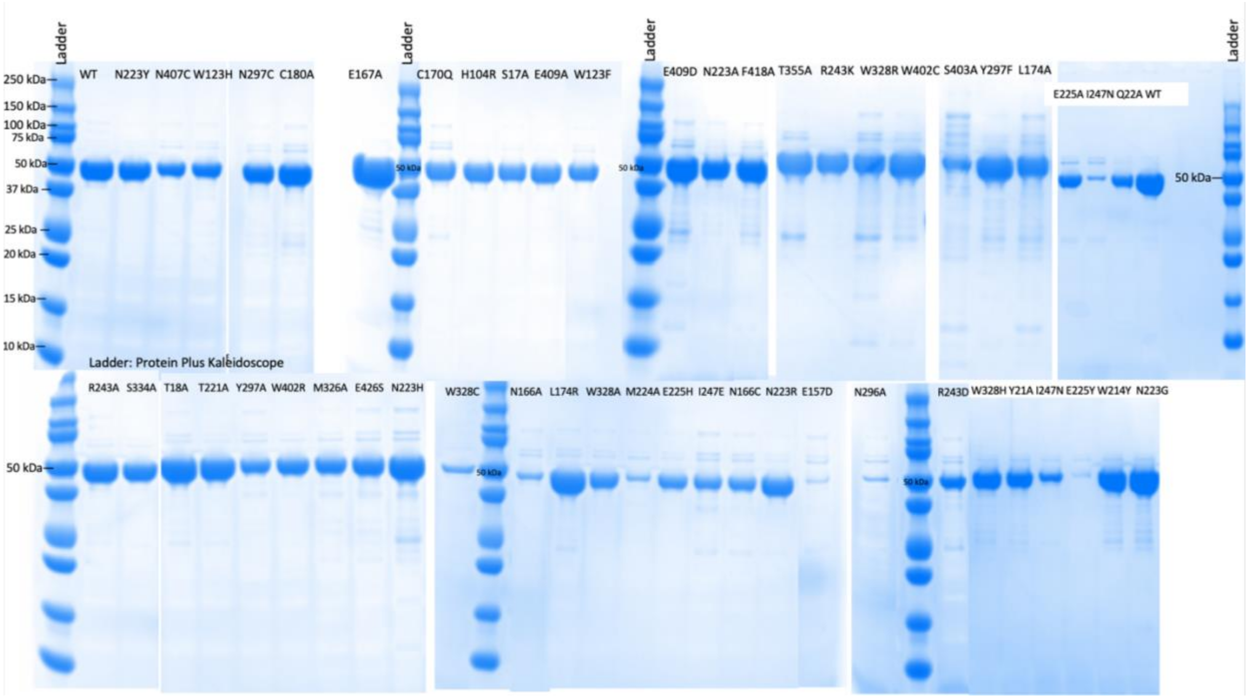


Supporting information 1-2. A distribution analysis of temperatures observed for each method. Black circles represent mutants and the red circle represents the native protein observed $T_M$ or $T_{50}$.

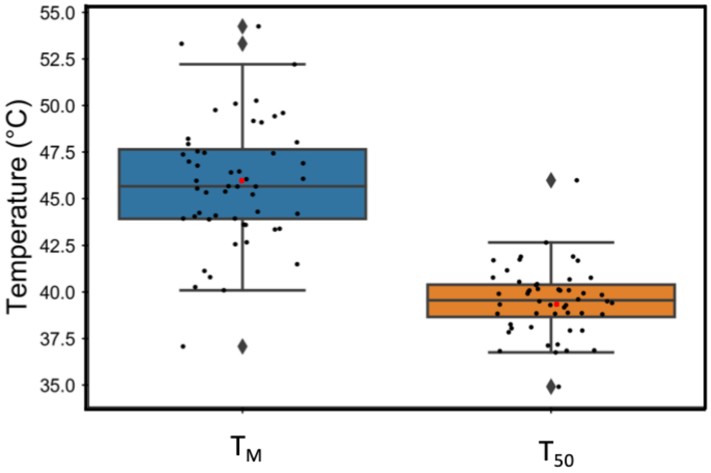

Supporting information 1-3. Pearson correlation coefficient (PCC) for $\Delta T_M$ and Experimental ΔΔG for 51 mutants of BglB were found to be 0.76.

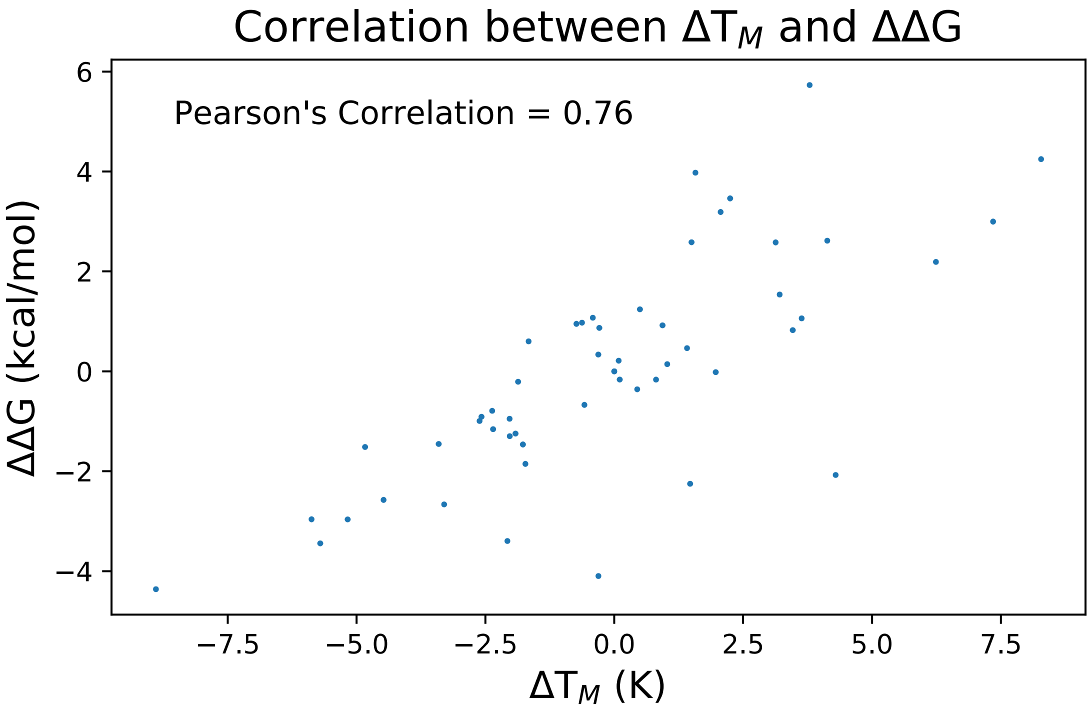


Supporting information 1-4. Evaluation of five computational methods on protein expression. Top panel is Rosetta ΔΔG, FoldX, and ELASPIC ΔTSE of soluble (Left) and insoluble (Right). Bottom panel is PoPMuSic and DeepDDG ΔTSE of soluble (Left) and insoluble (Right).

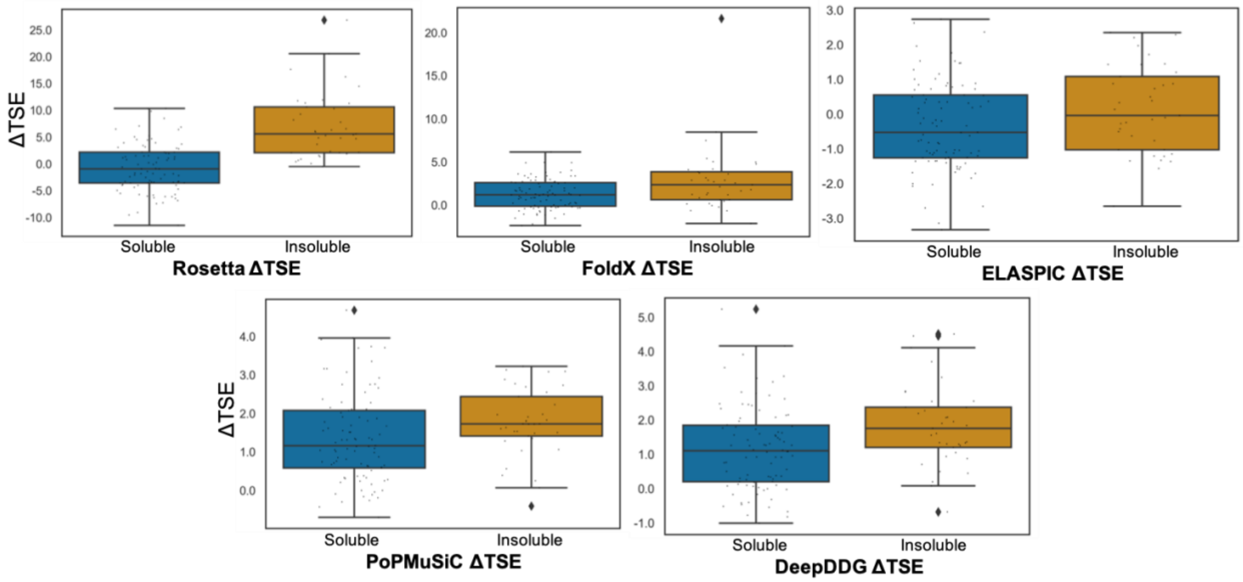

Supporting information 2.

The zip file contains images of fluorescence graphs, 1st derivative graphs, and Van't Hoff plot for 51 mutants in quadruplicates and 6 biological replicates for WT in .pdf format. All graphs were generated using matplotlib found in SI 4.

Supporting information 3.

The SI 3 file contains two folders (Rosetta ΔΔG and FoldX PSSM) consisted of PCC graphs of $\Delta T_M$ with each individual energy term described in the Rosetta ΔΔG and FoldX PSSM energy scoring protocols. Each folder also has a .csv file containing all the raw data from each algorithm for all previously described mutants. Lastly, the SI 3 file contains a Finalized_exp_dgg.csv file including all the thermodynamic parameters (ΔΔG, ΔΔH, ΔΔS, and $\Delta T_M$) derived from the fluorescence melting curve, as well as gel number that corresponds to each of the 51 mutants.

Supporting information 4.

The SI_script.ipynb script contains method used for data acquisitions to generate all the graphs from experimental $T_M$ data, as well as methods to obtain all the thermodynamics parameters (ΔΔG, ΔΔH, ΔΔS, and $\Delta T_M$). The folder also includes individual .csv files of all raw data used (fluorescence vs temperature) for data acquisitions.

Supporting information 5.

This folder contains bglb_apo.pdb, flags, mutant_file, and sub.sh file needed to execute Rosetta_ddg_monomer application that have been previously described in Kellogg 2011.
