## Supplementary figures and images for "Evaluating molecular modeling tools for thermal stability using an independently generated dataset"

### backbone clash.png

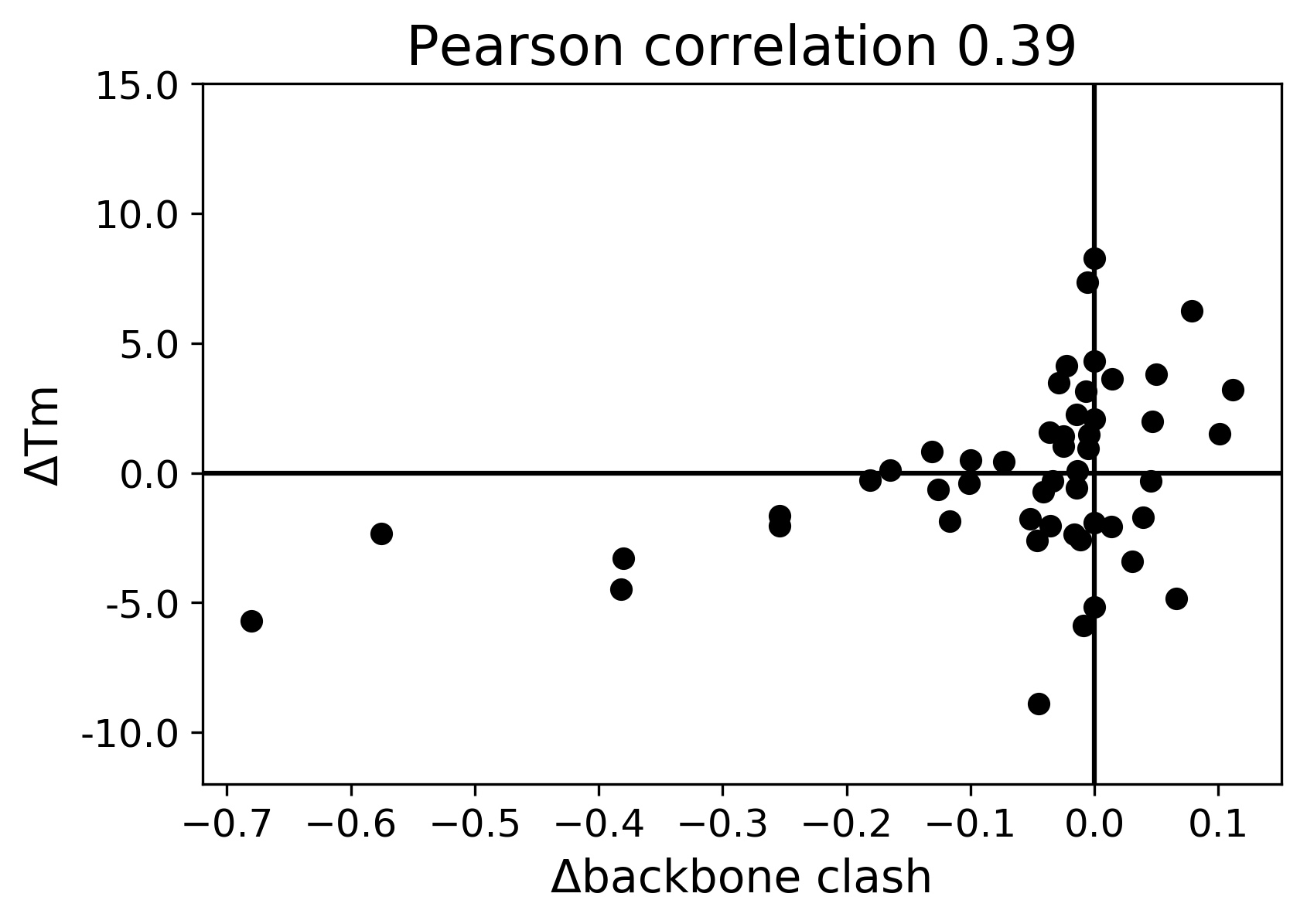

### Backbone Hbond.png

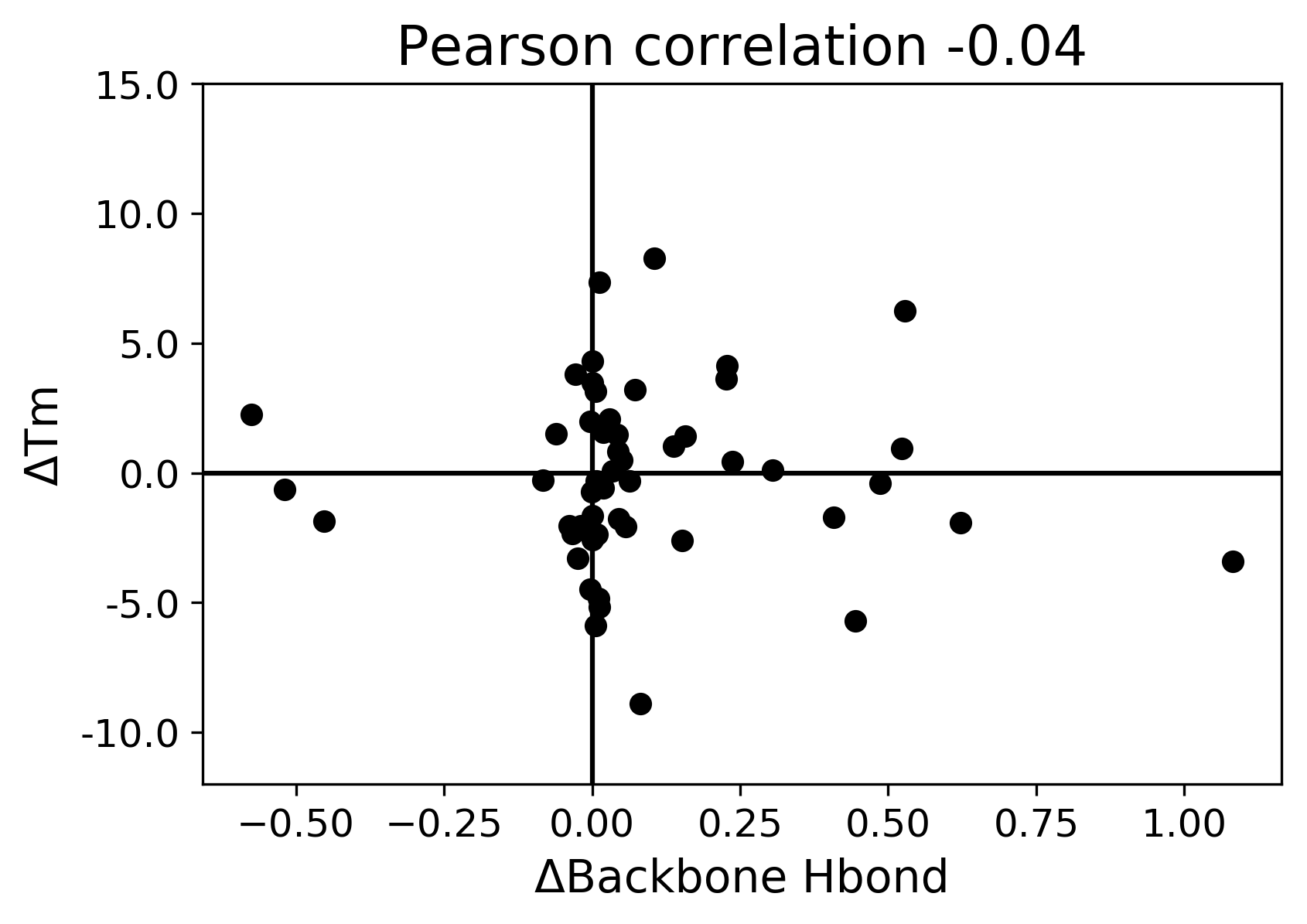

### C170Q_C10_fluorescence.pdf

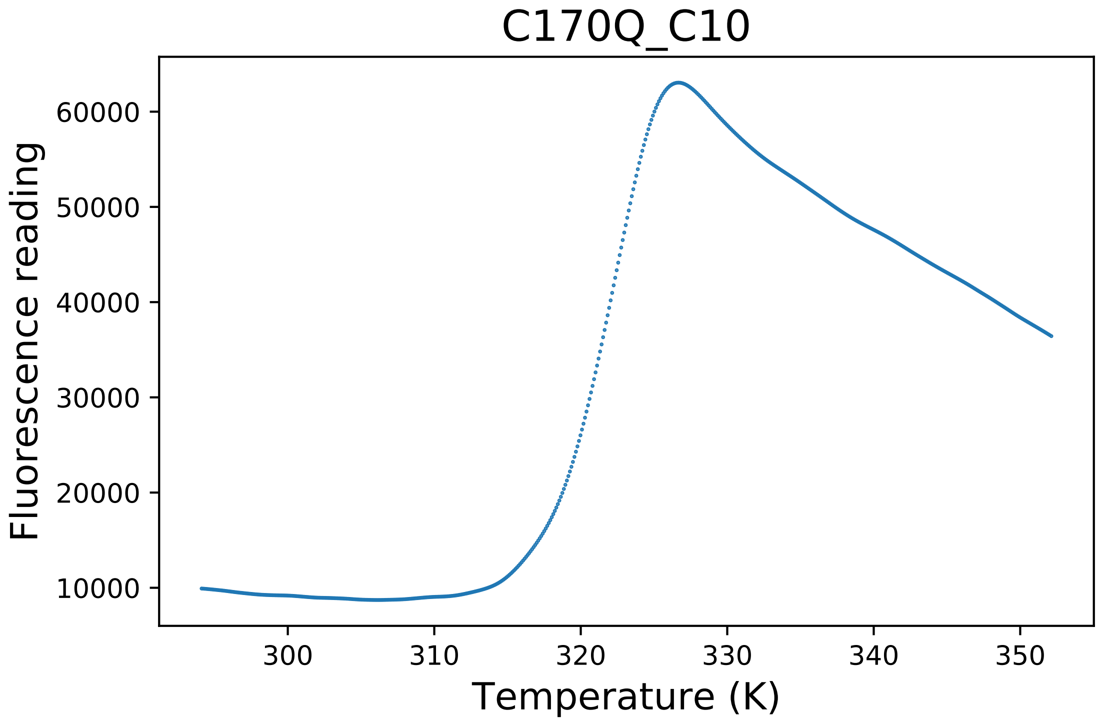

### dslf_fa13.png

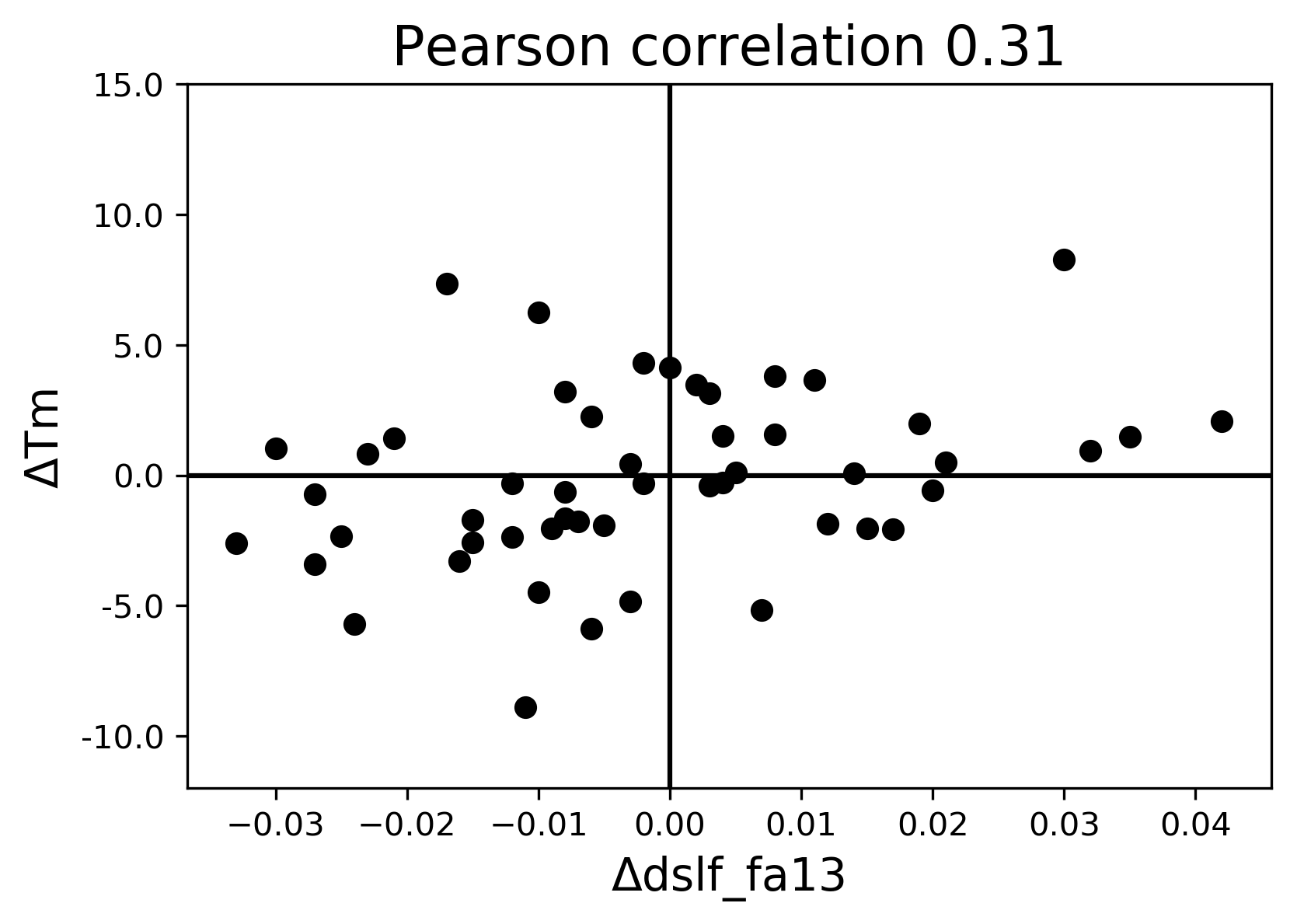

### E225H_C09_fluorescence.pdf

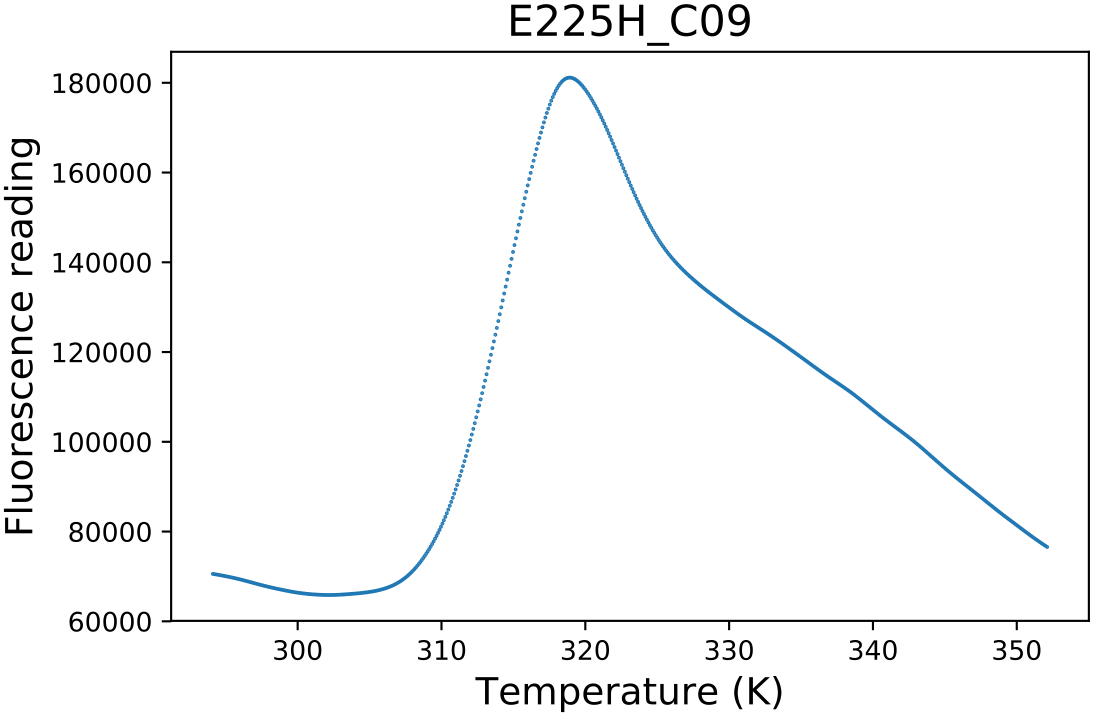

### E409A_C08_dg.pdf

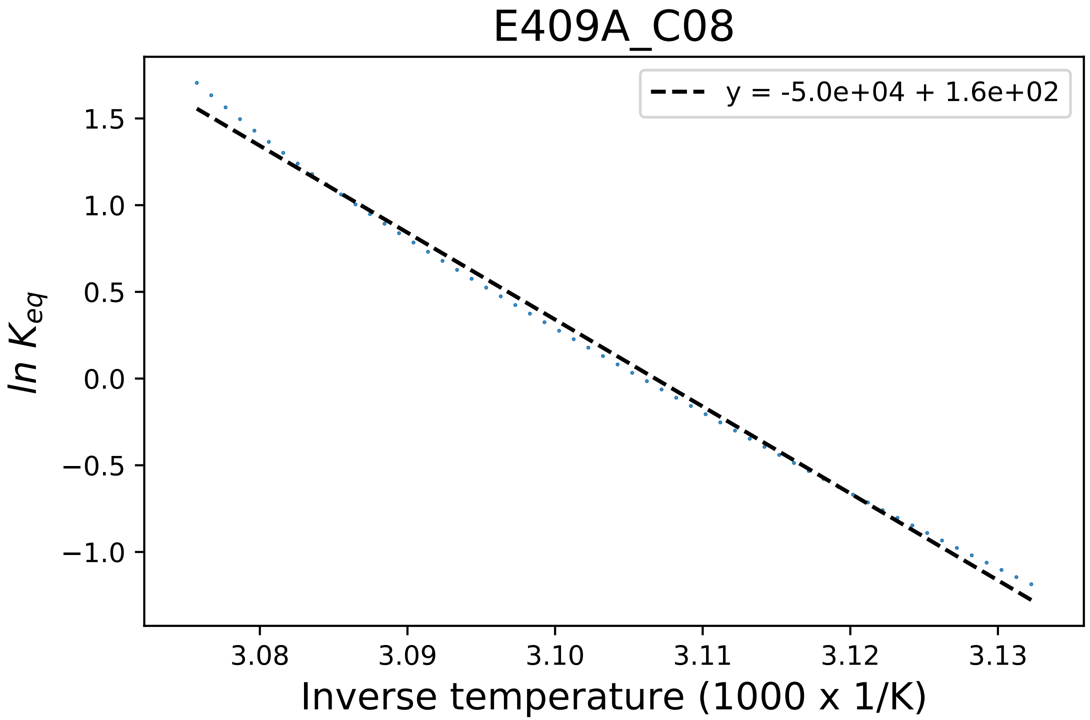

### E409D_F01_dg.pdf

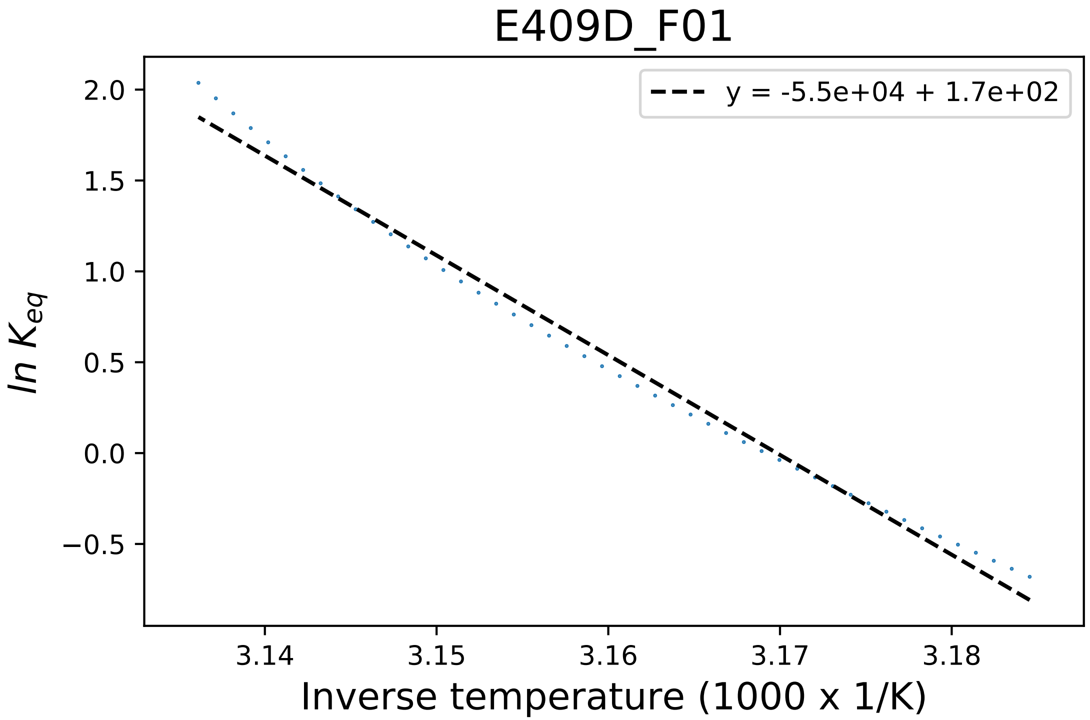

### Electrostatics.png

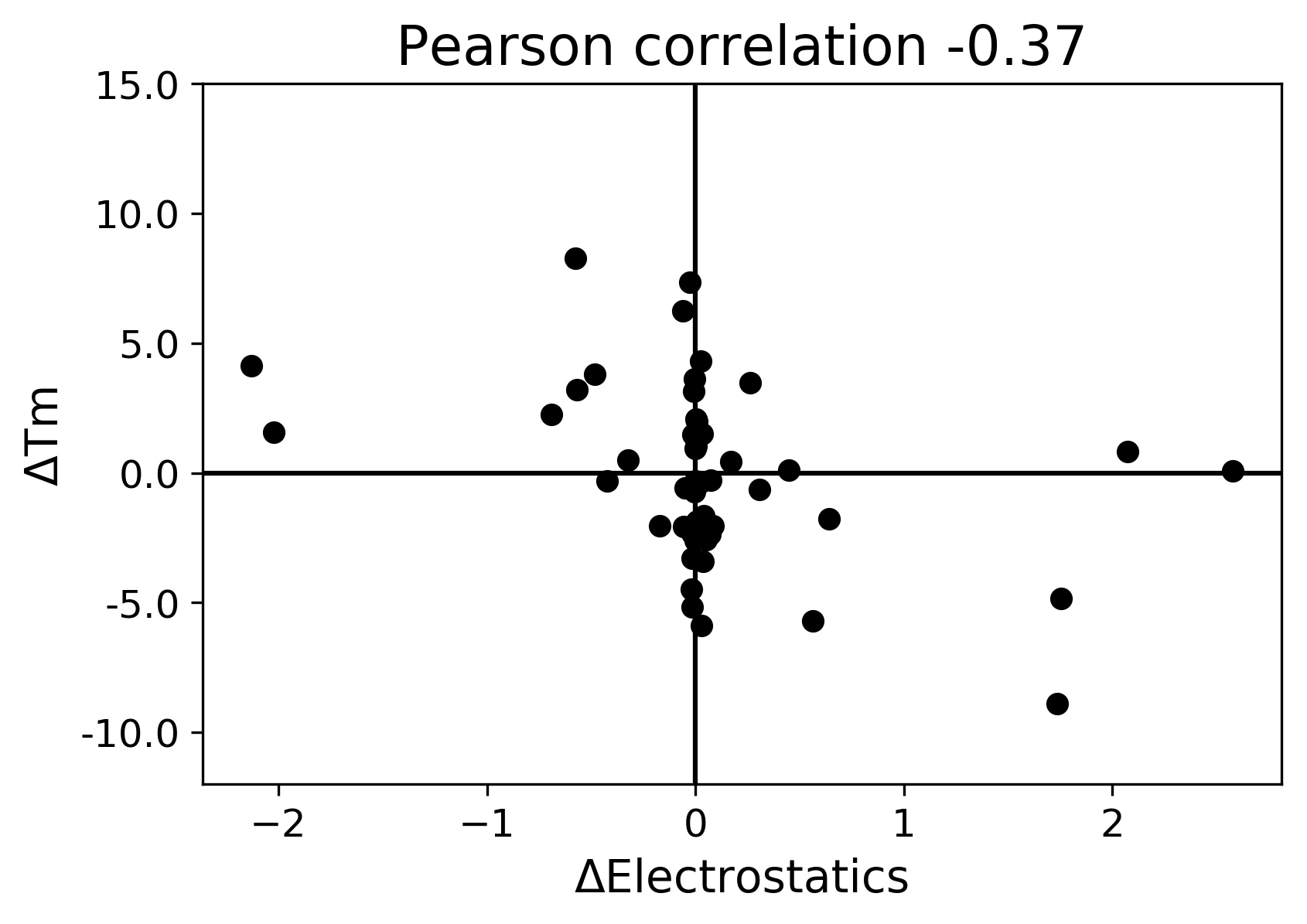

### energy ionisation.png

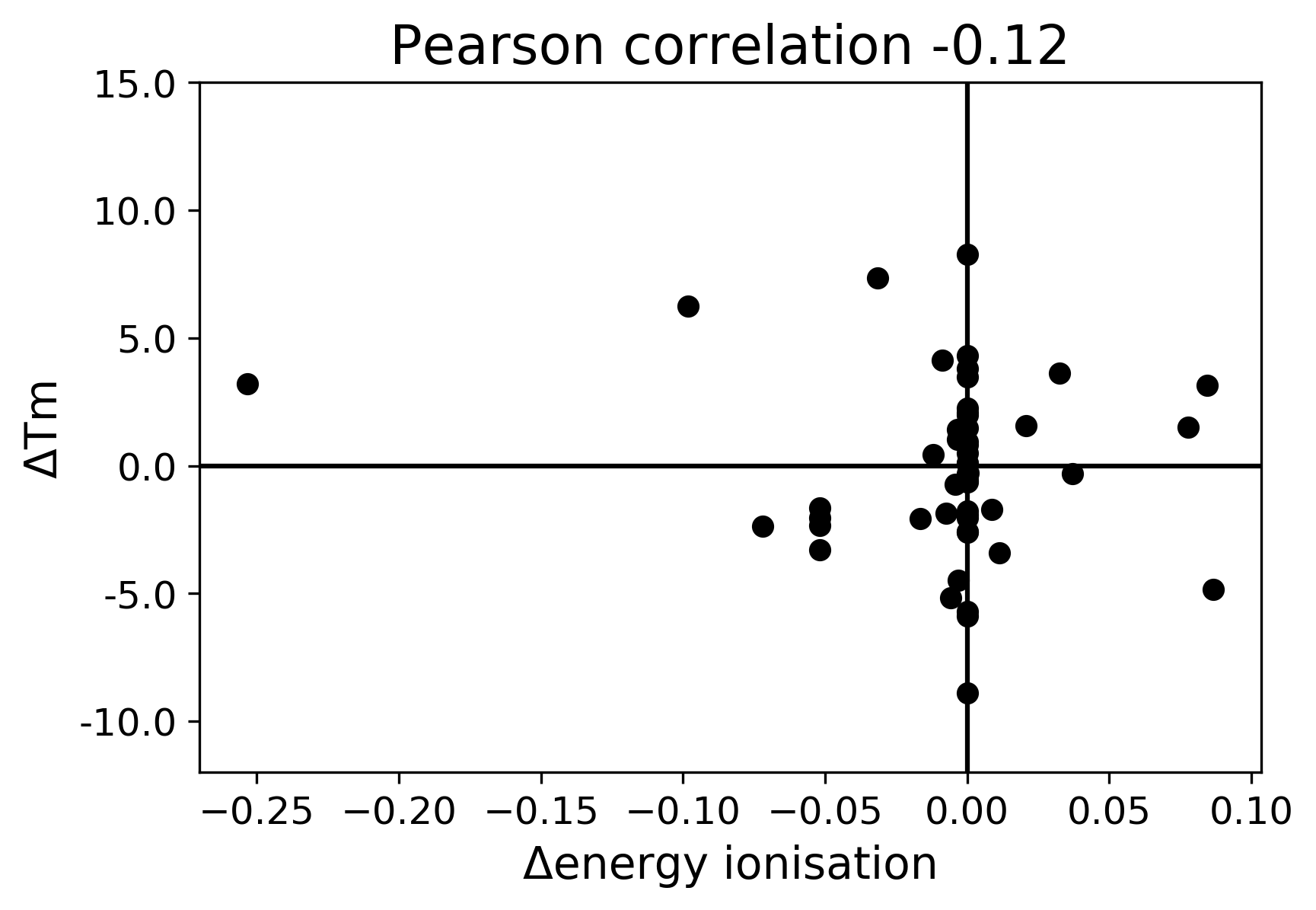

### entropy mainchain.png

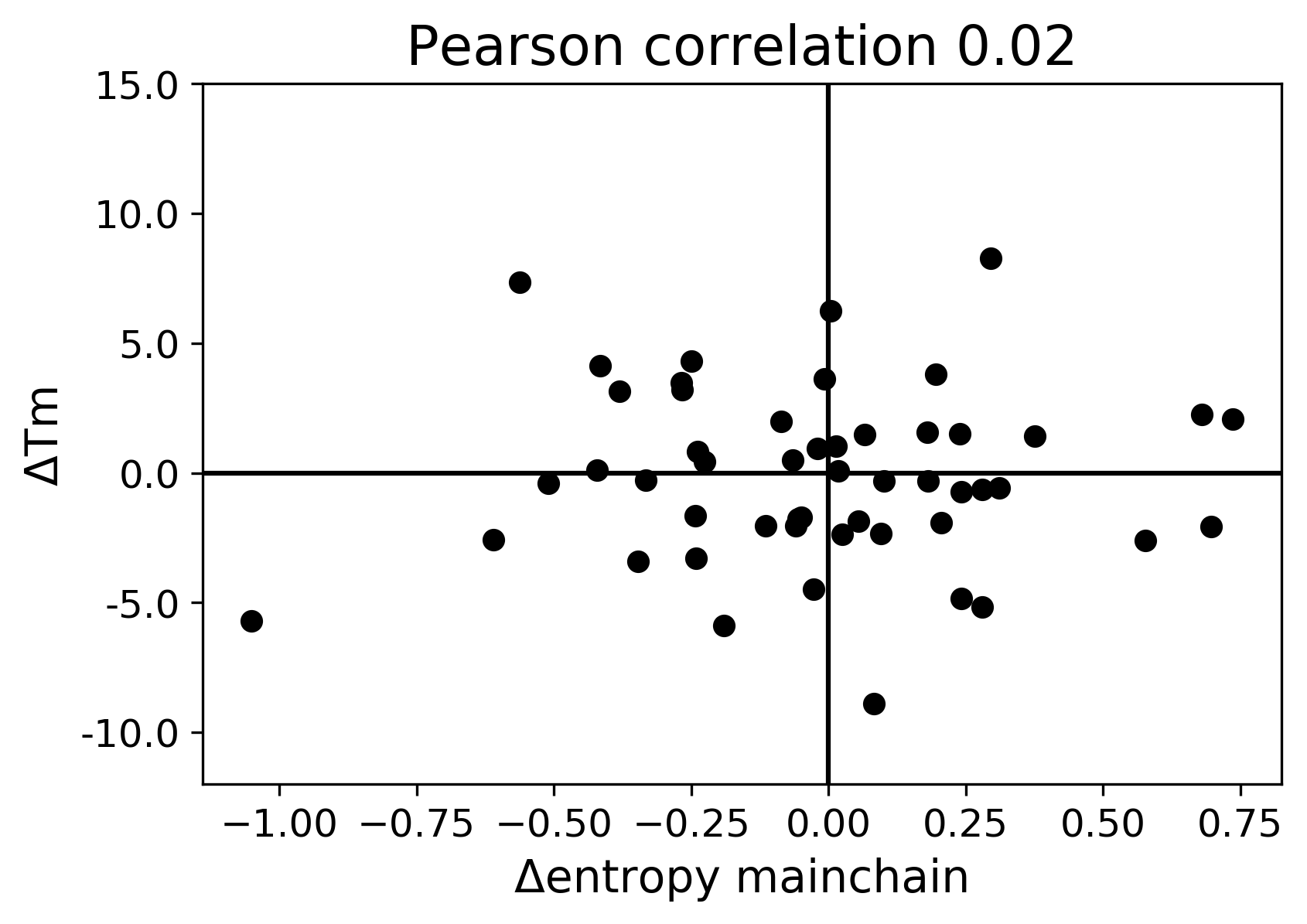

### Entropy sidechain.png

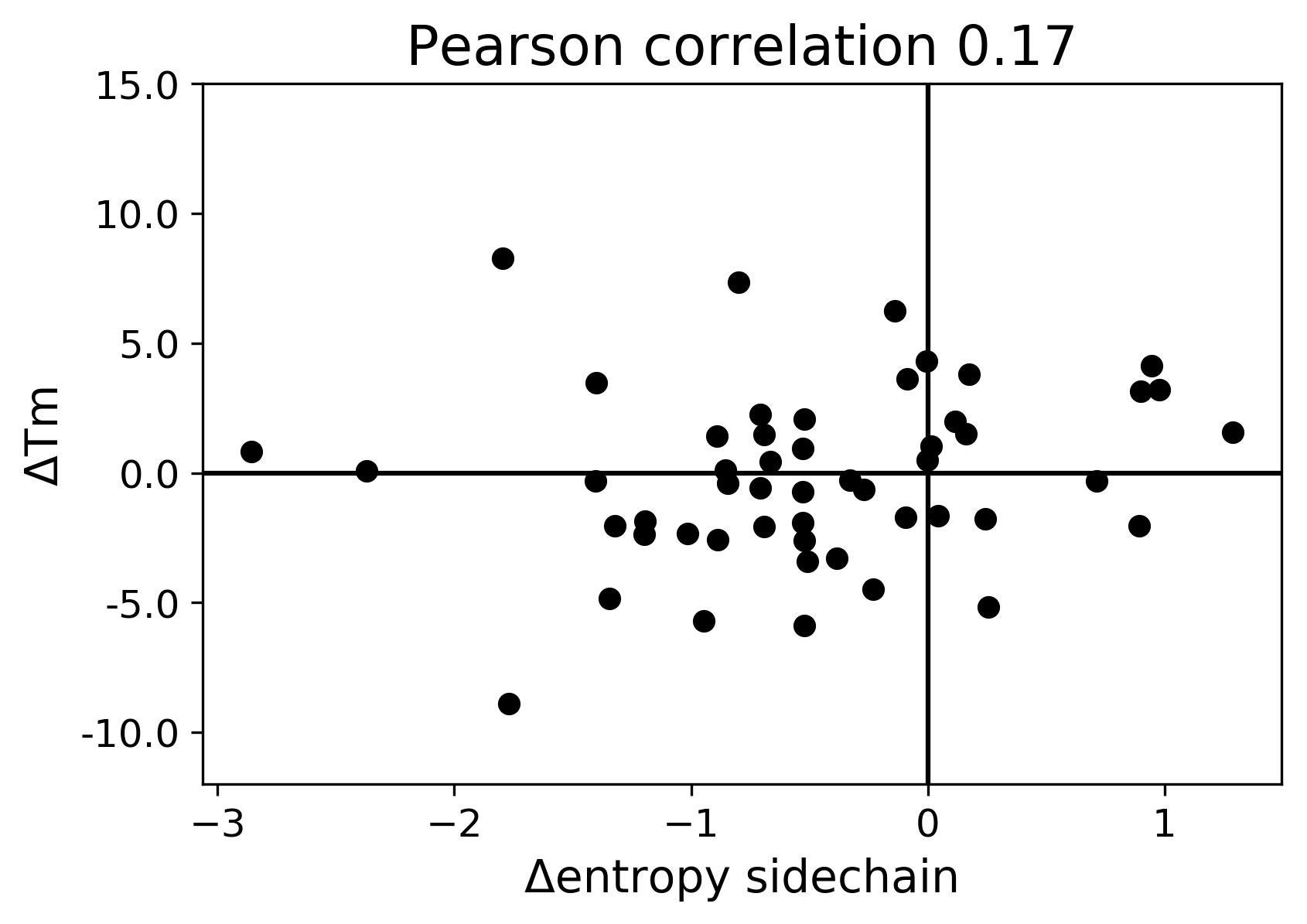

### fa_atr.png

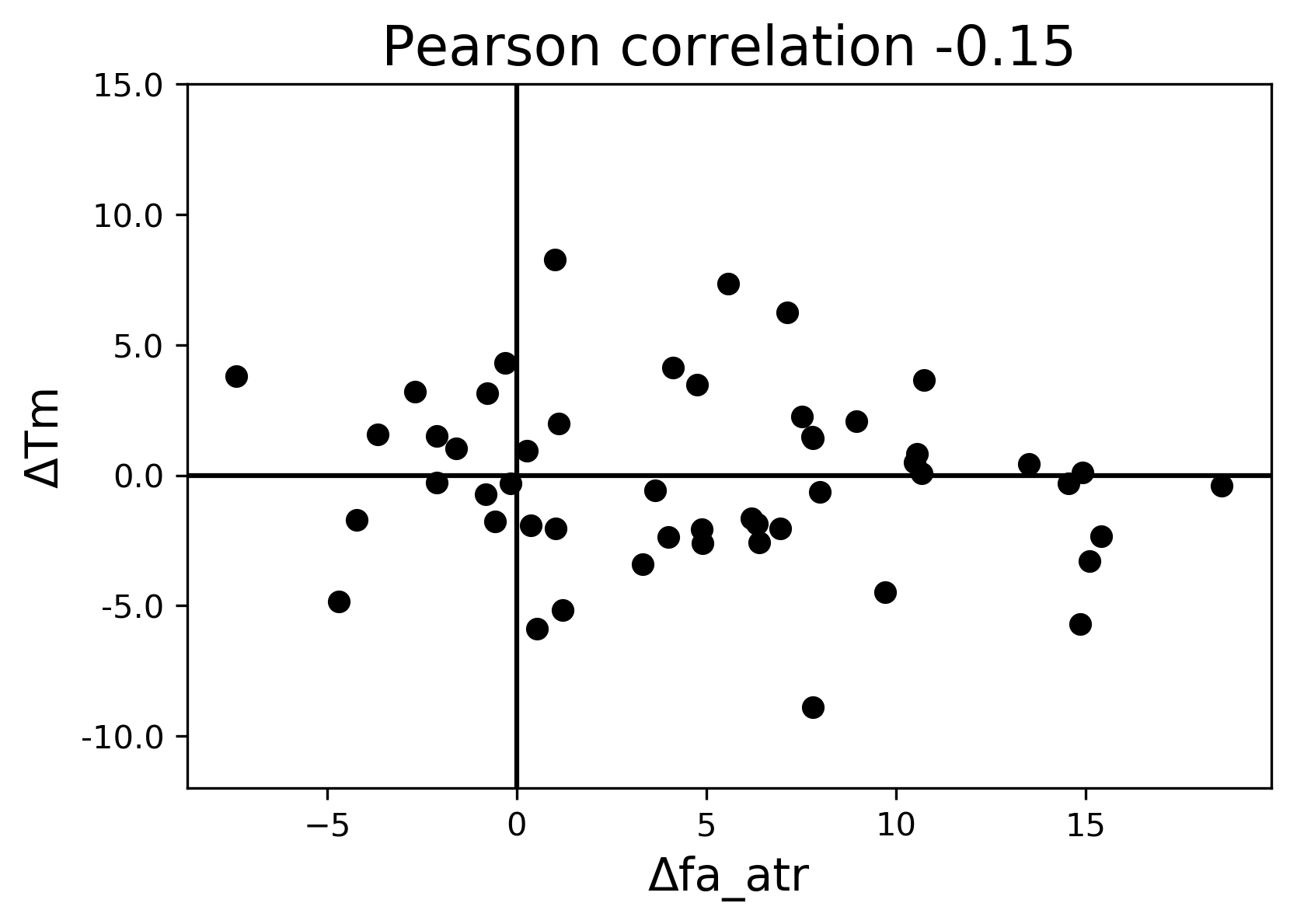

### fa_dun.png

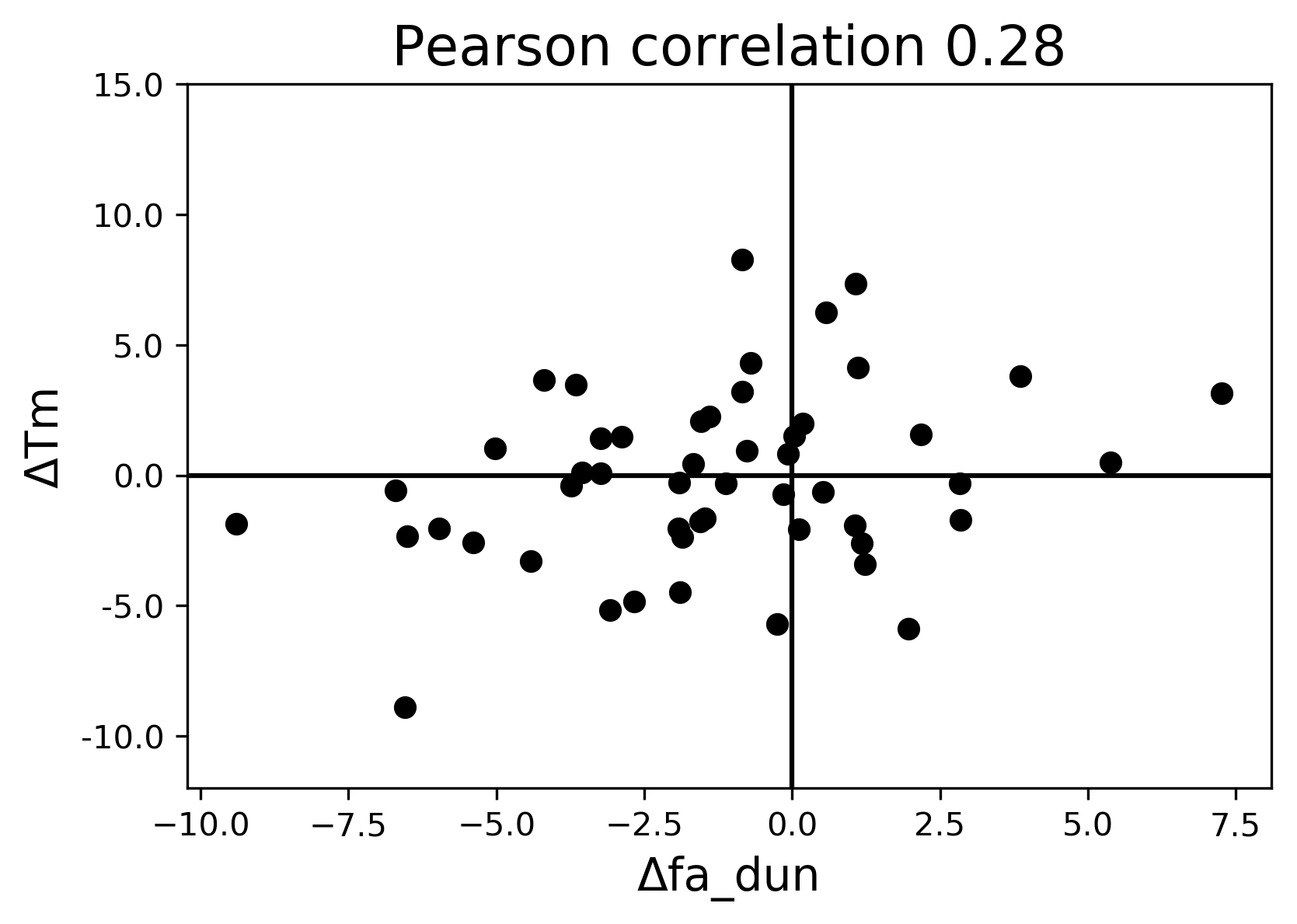

### fa_elec.png

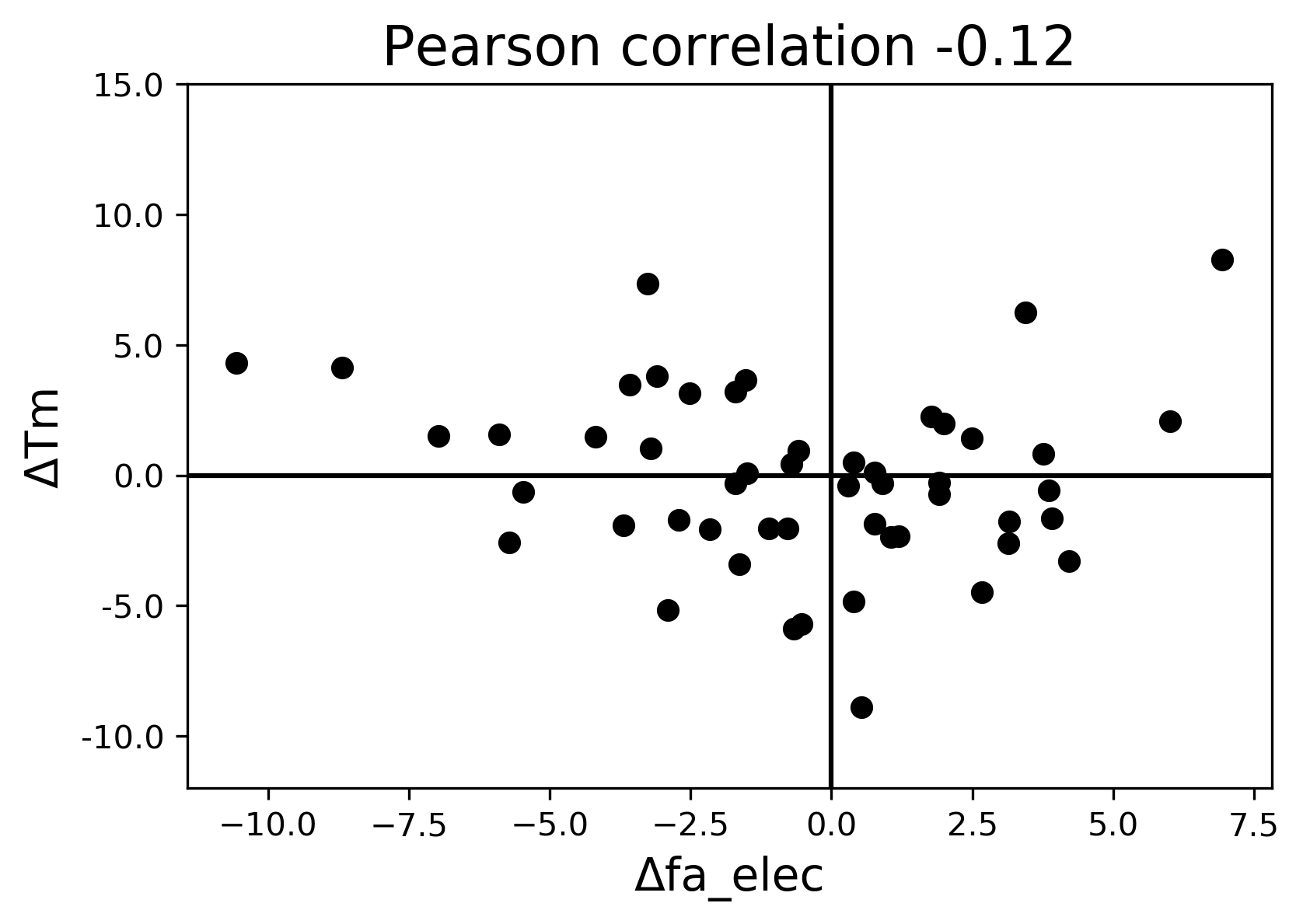

### fa_intra_rep.png

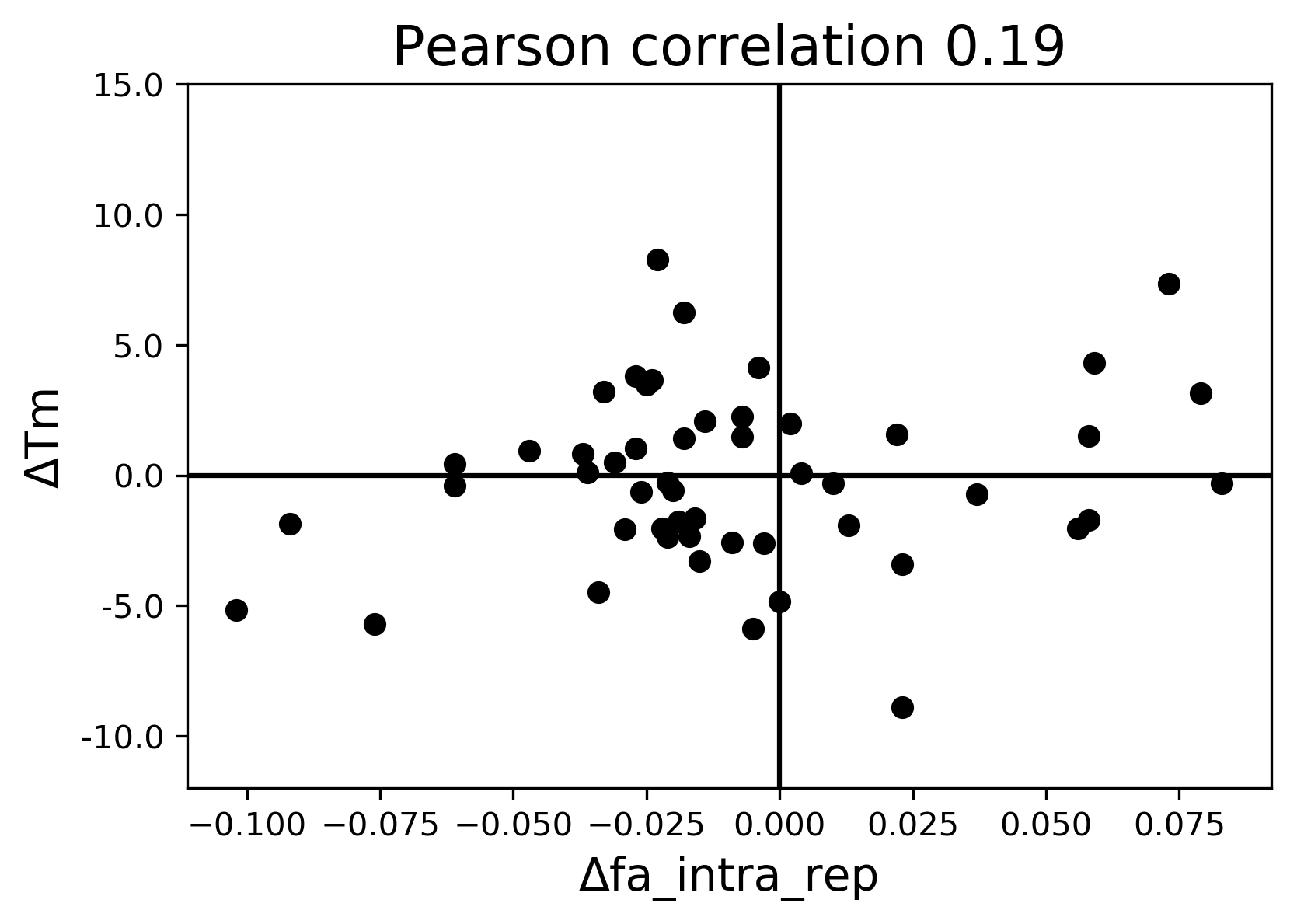

### fa_intra_sol_xover4.png

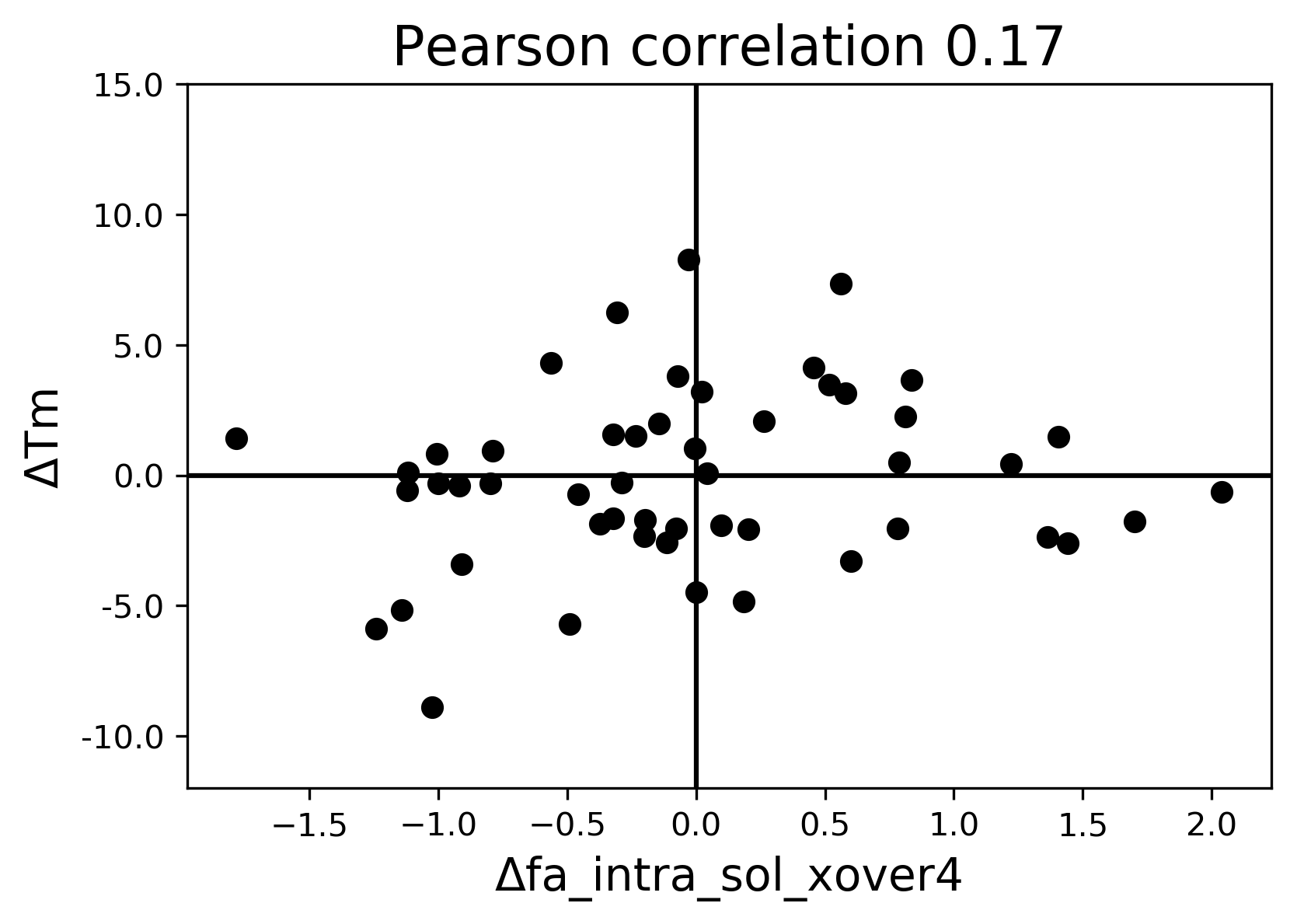

### fa_rep.png

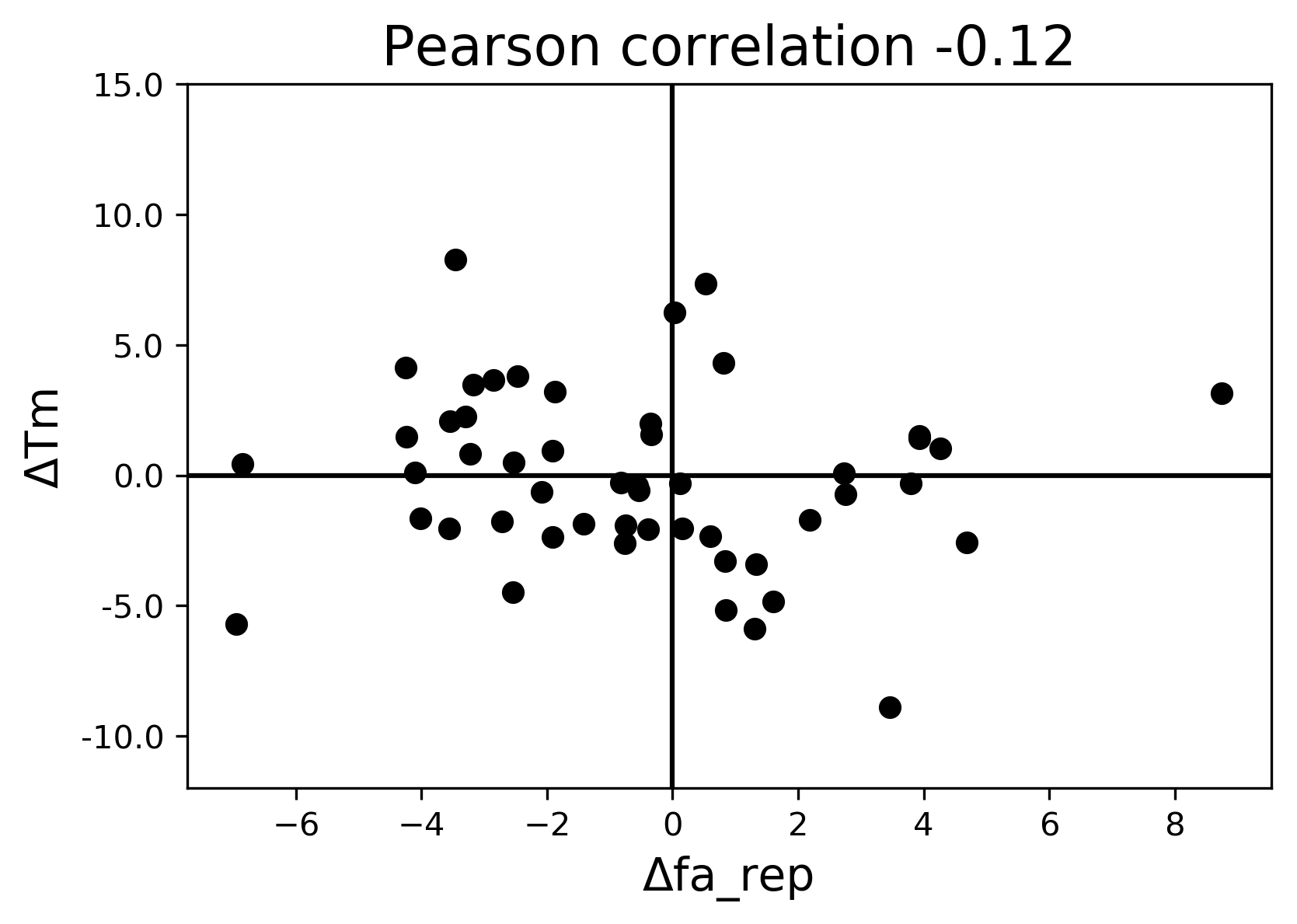

### fa_sol.png

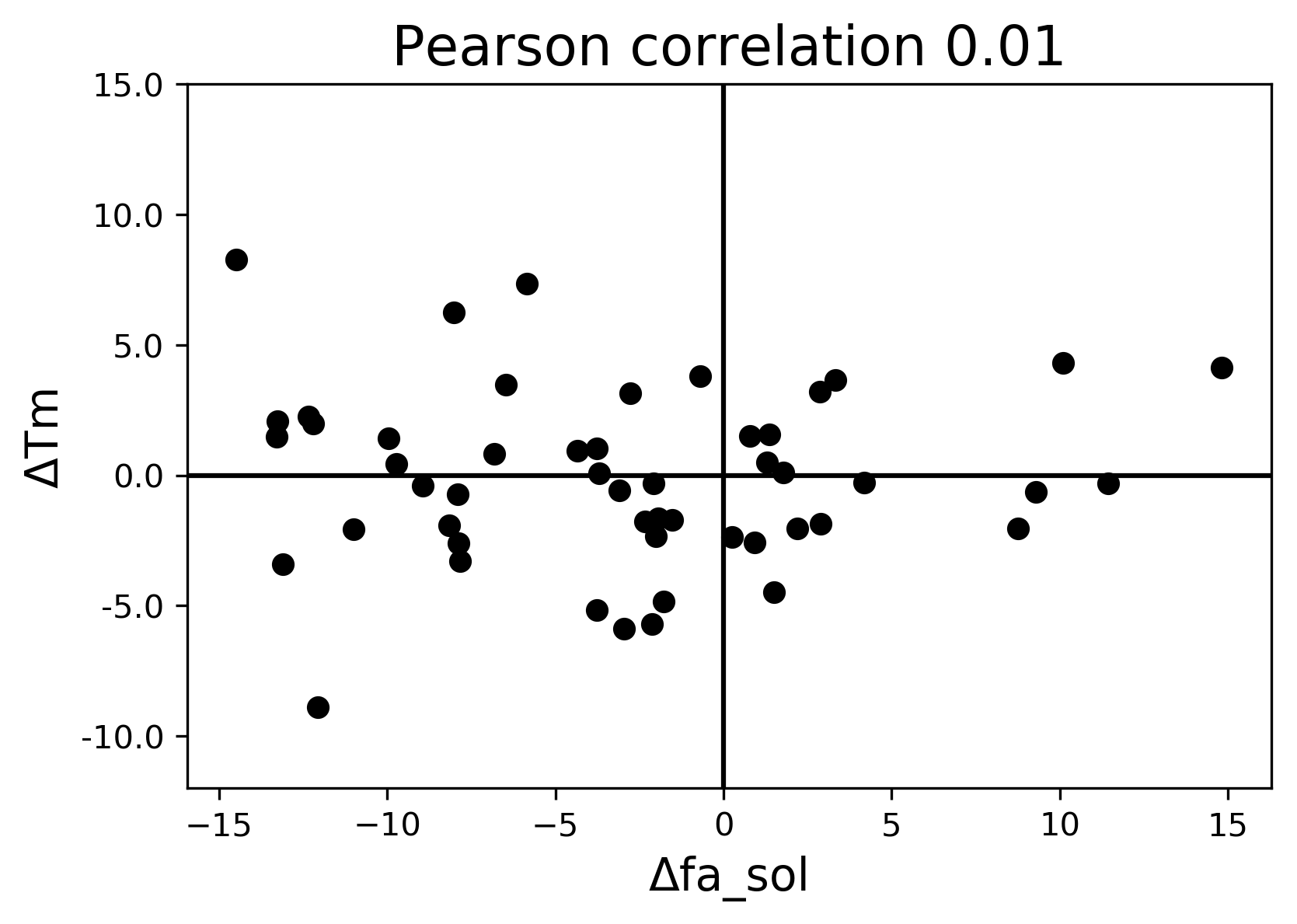

### H104R_F11_dg.pdf

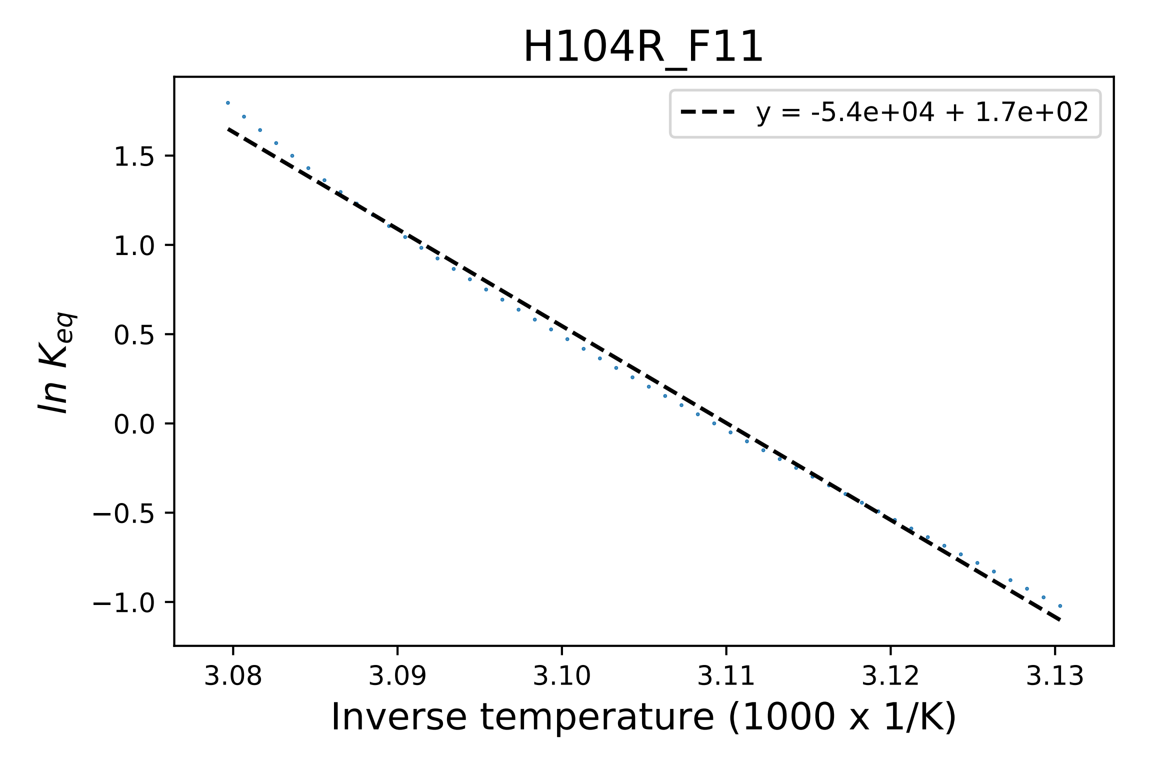

### H104R_F11_fluorescence.pdf

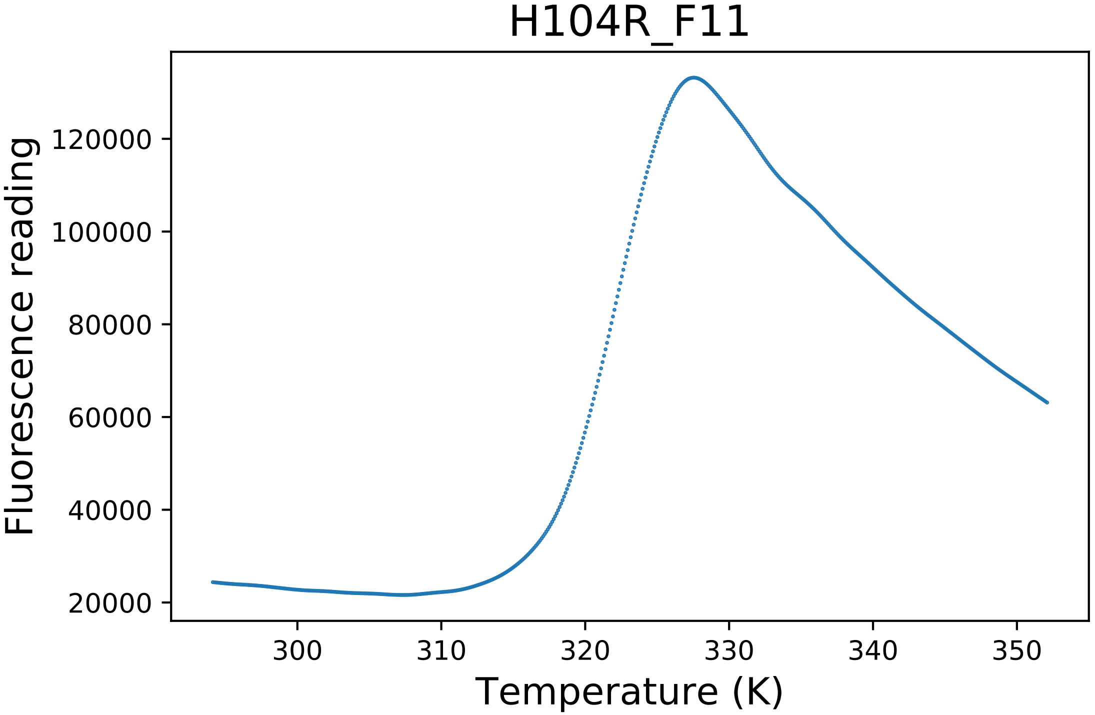

### hbond_bb_sc.png

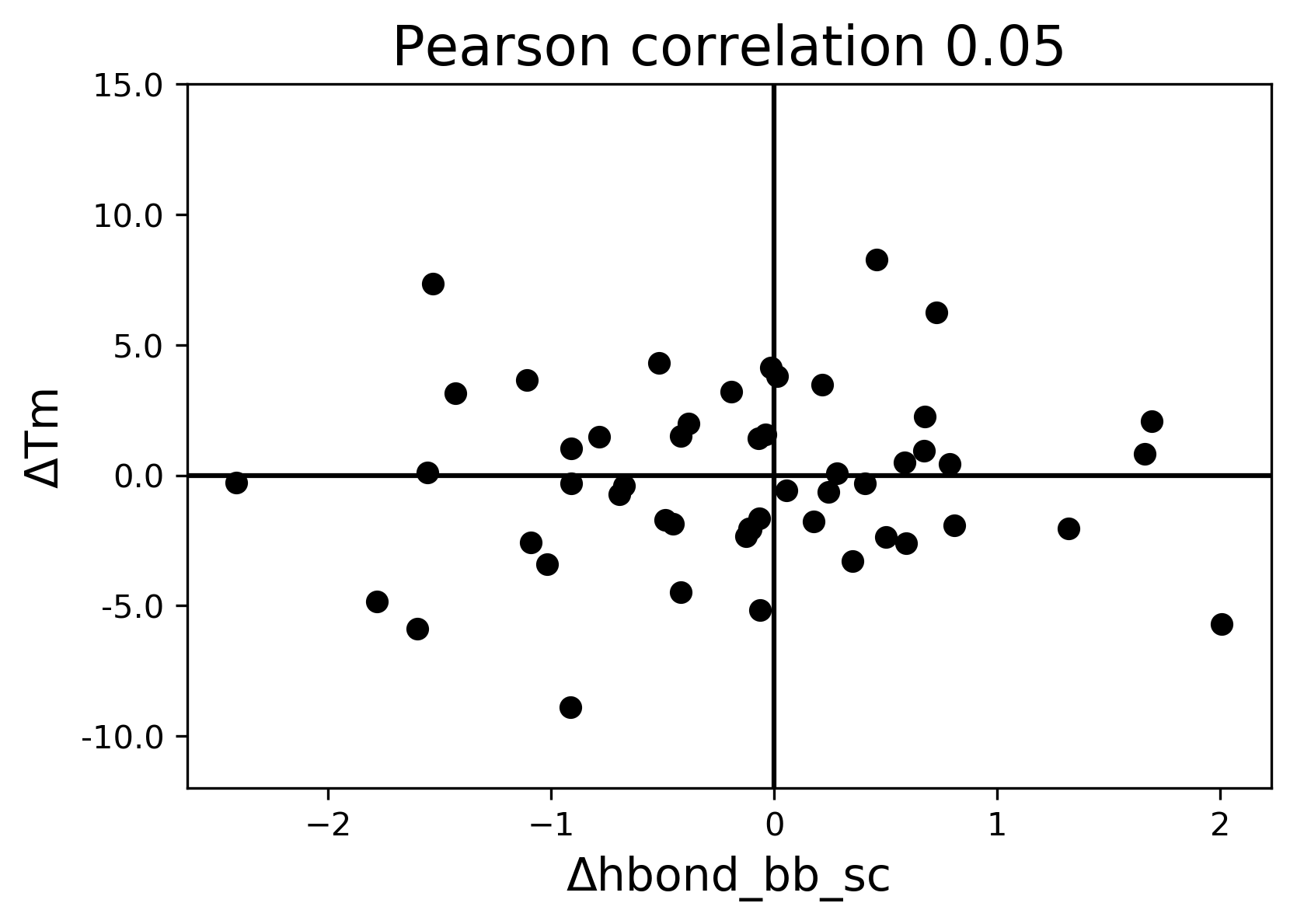

### hbond_lr_bb.png

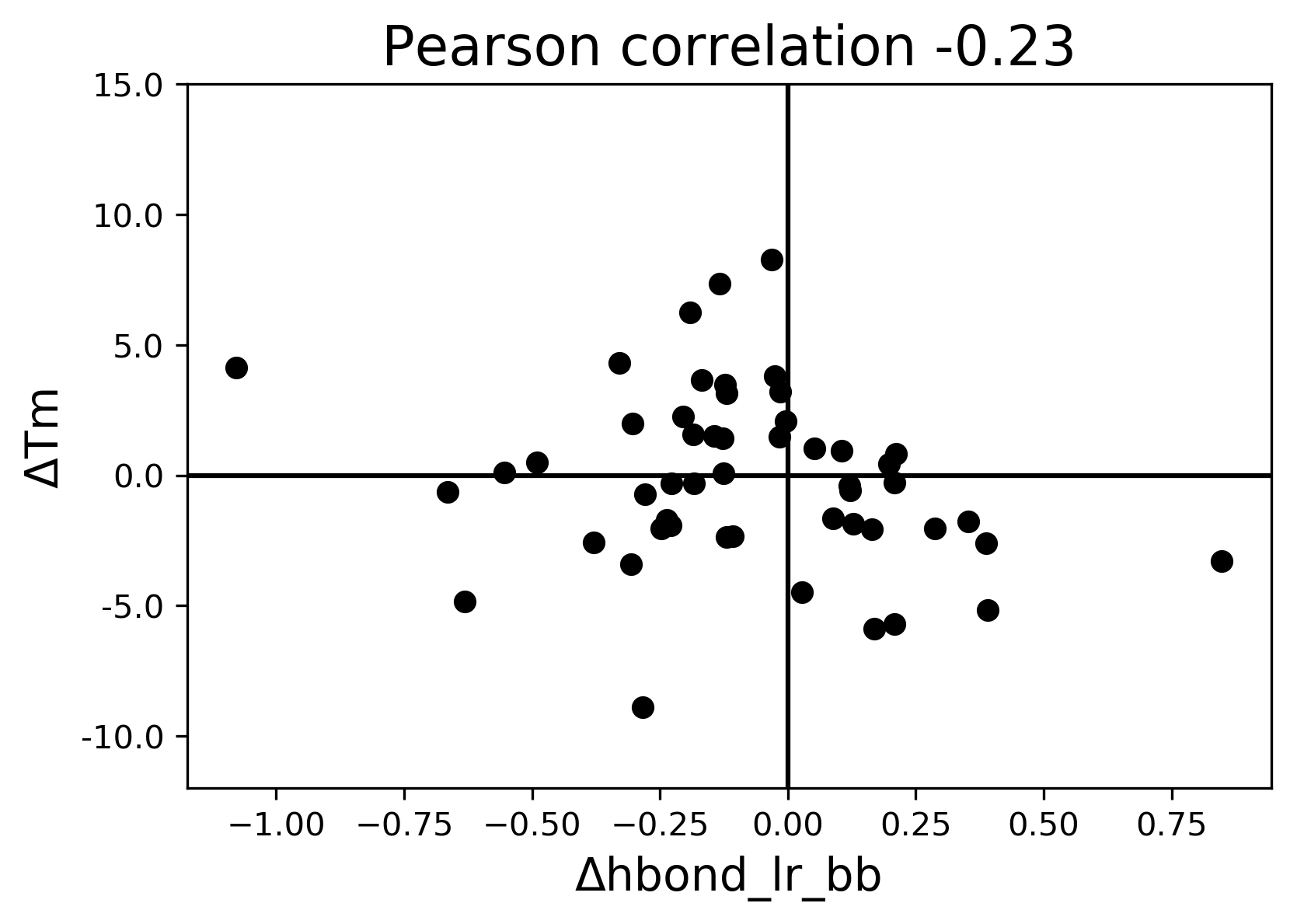

### hbond_sc.png

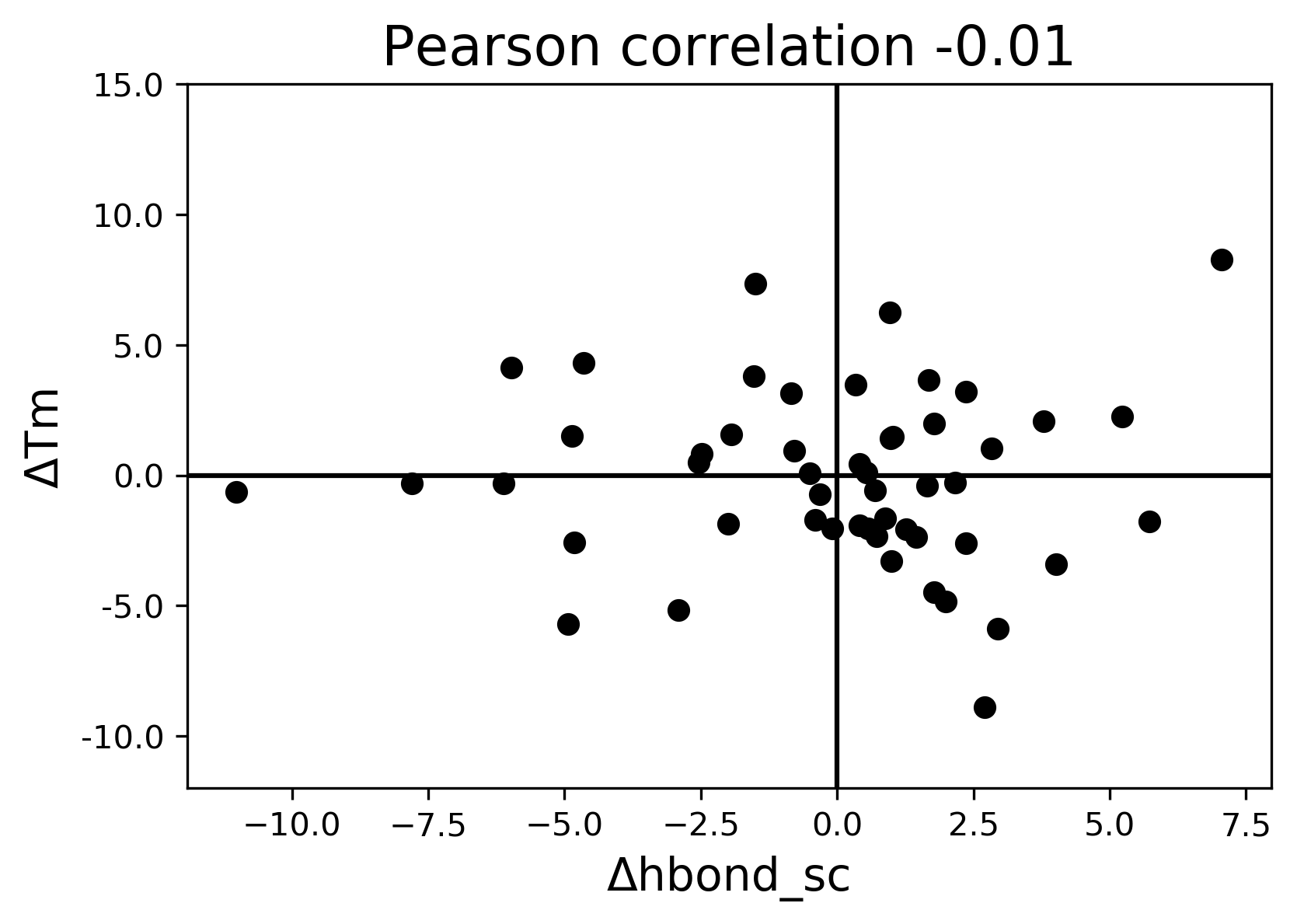

### hbond_sr_bb.png

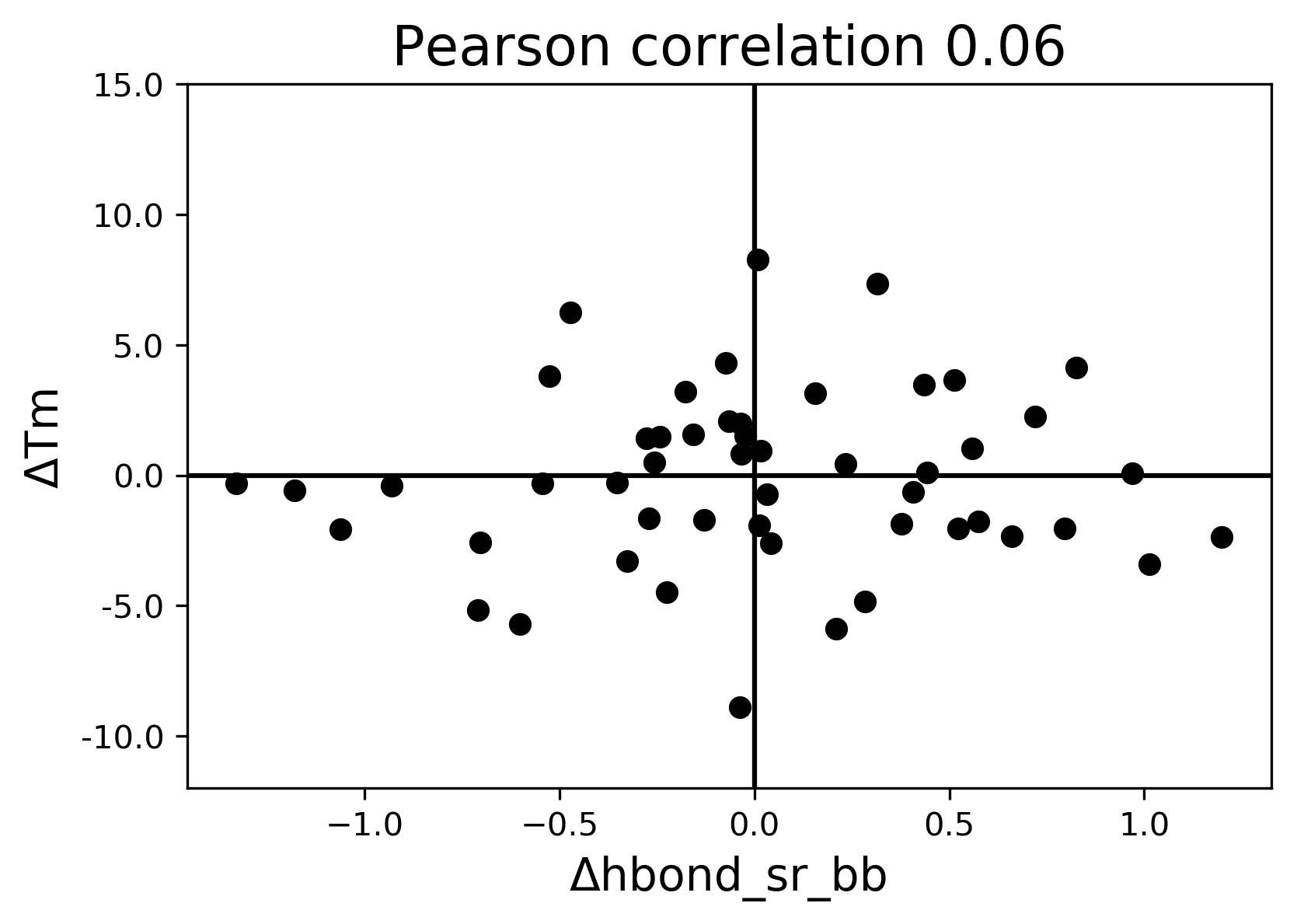

### helix dipole.png

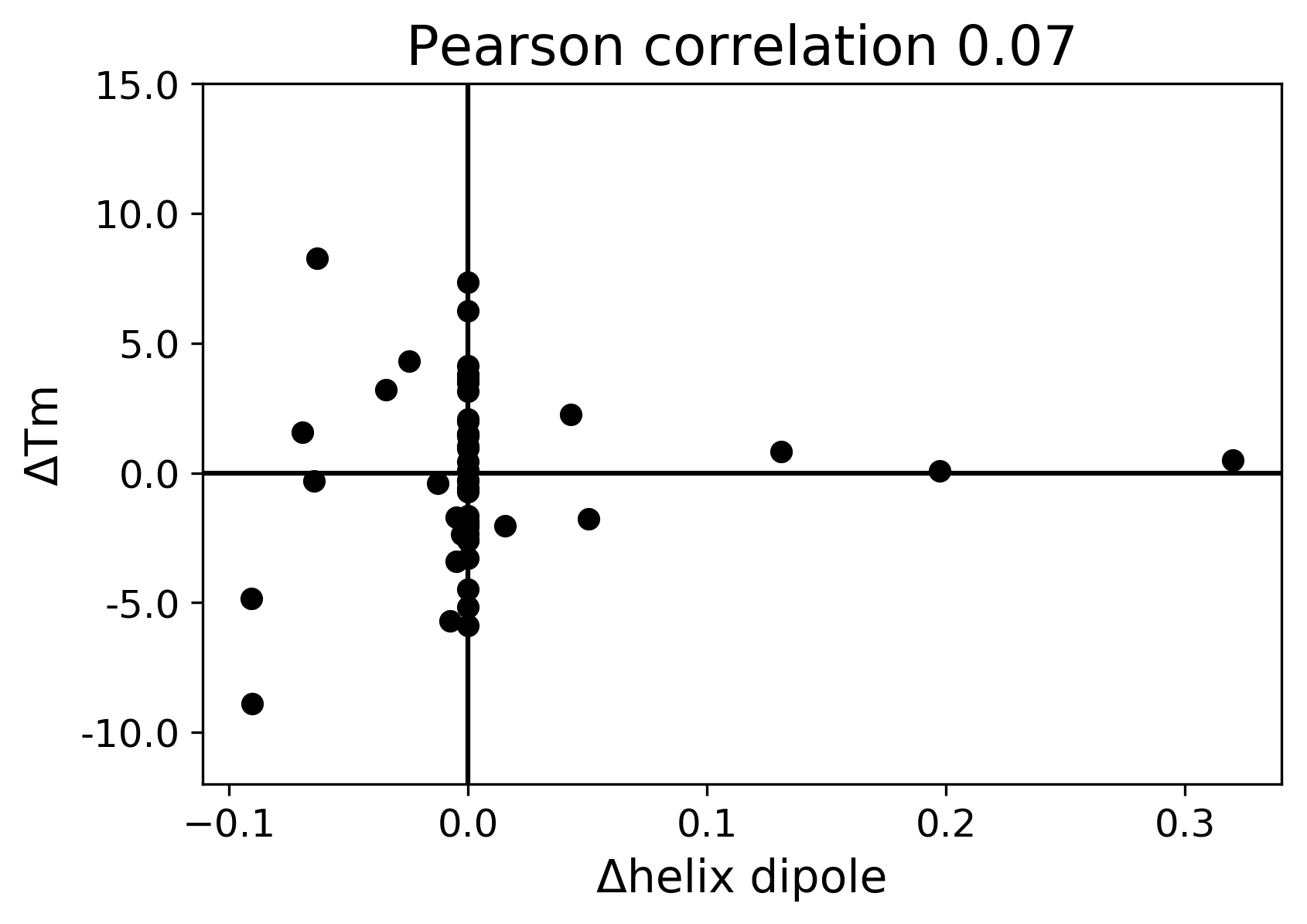

### I247N_B01_derivative.pdf

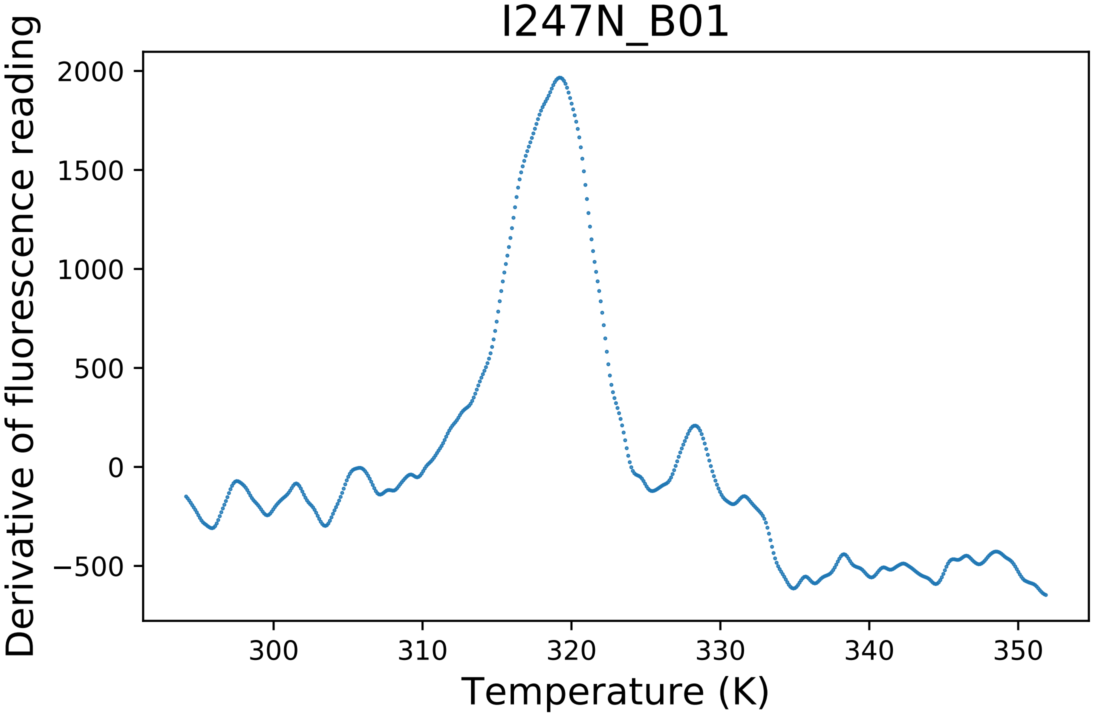

### I247N_B01_dg.pdf

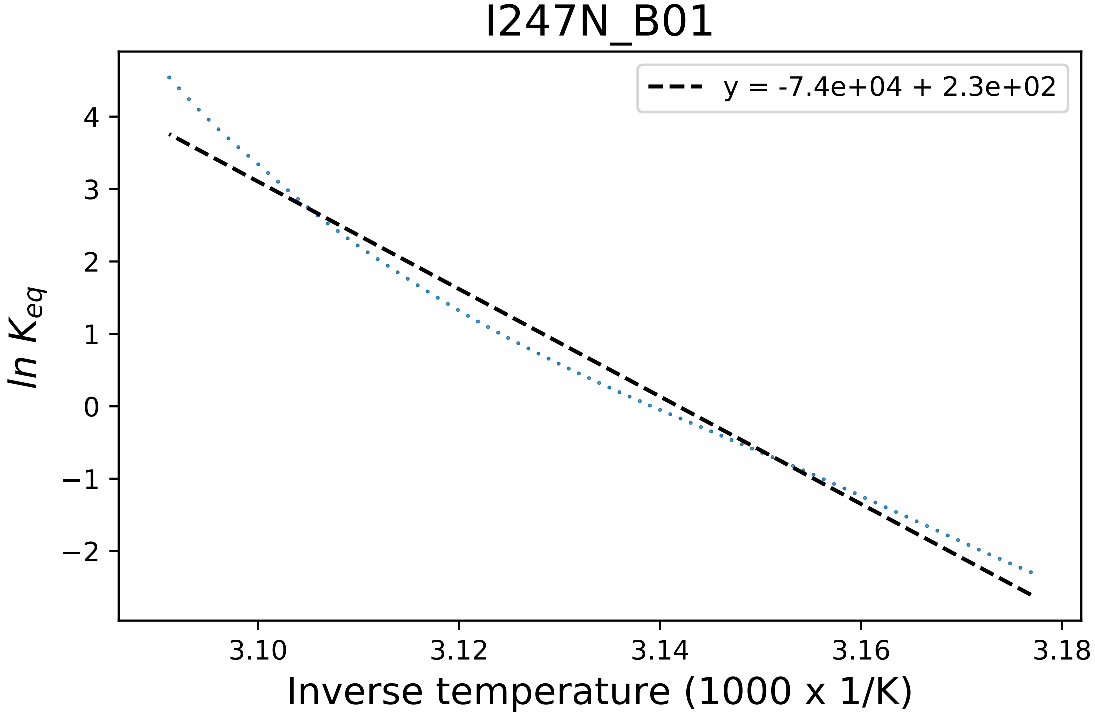

### L174A_E03_fluorescence.pdf

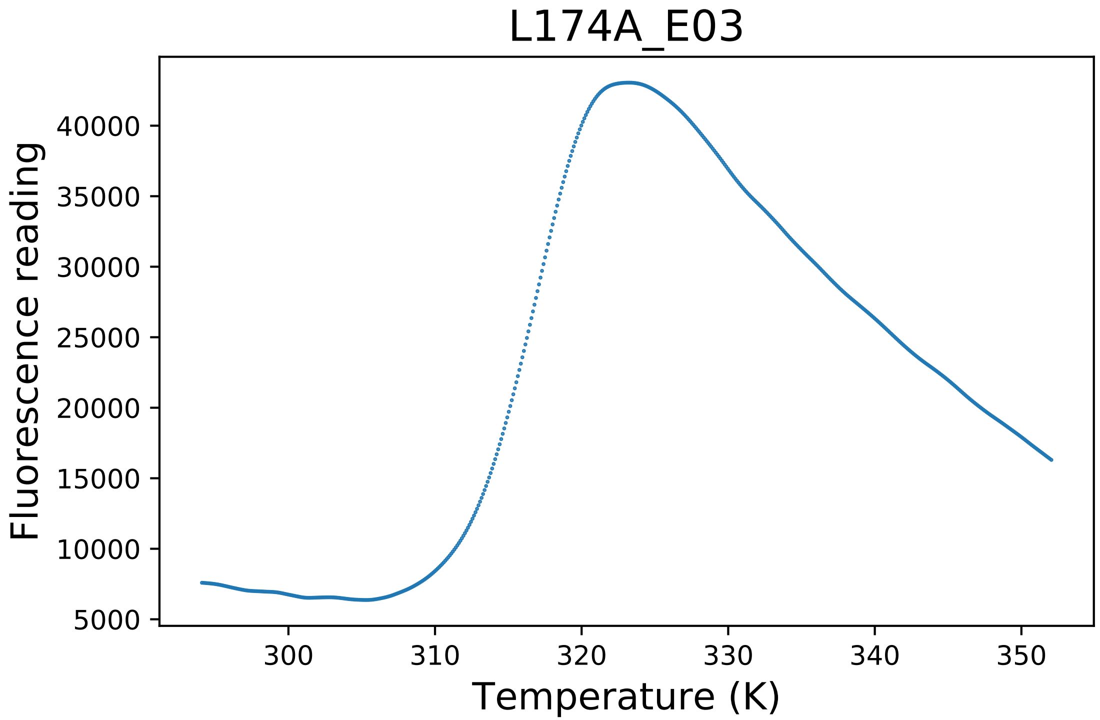

### L174R_G09_derivative.pdf

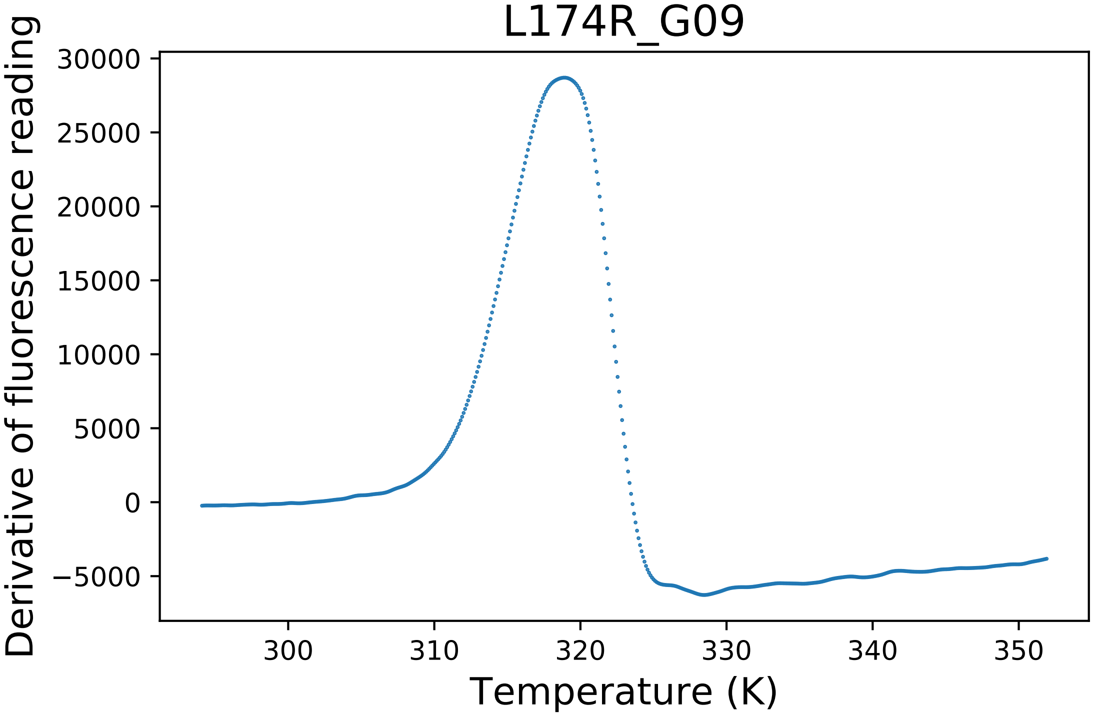

### L174R_G09_dg.pdf

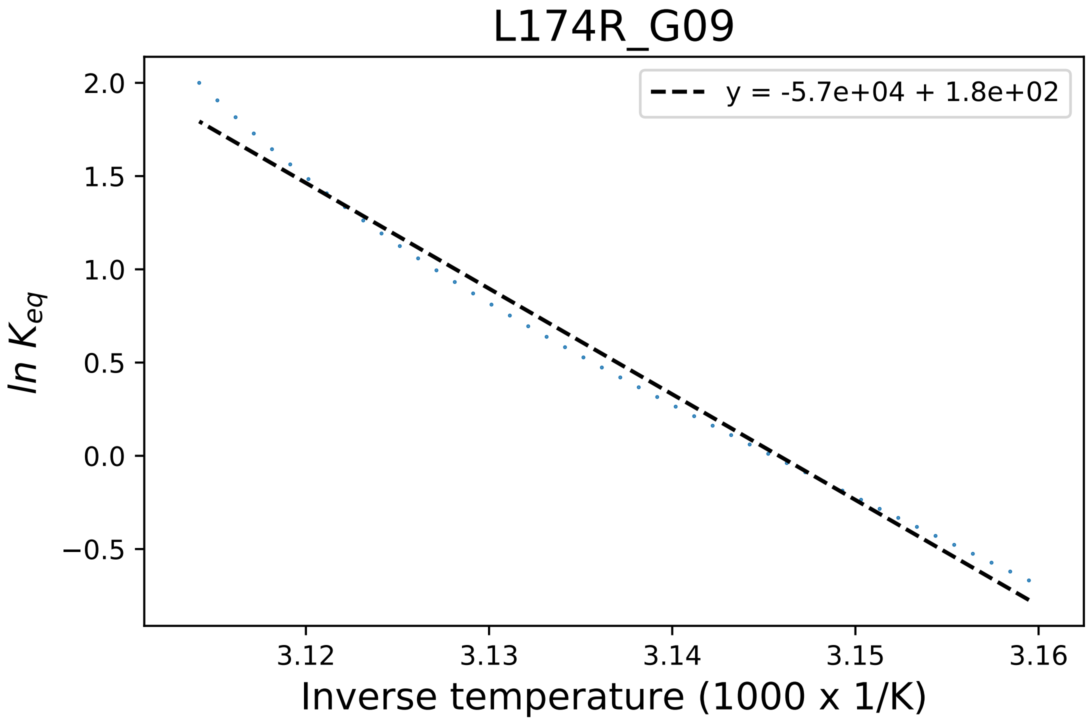

### L174R_G09_fluorescence.pdf

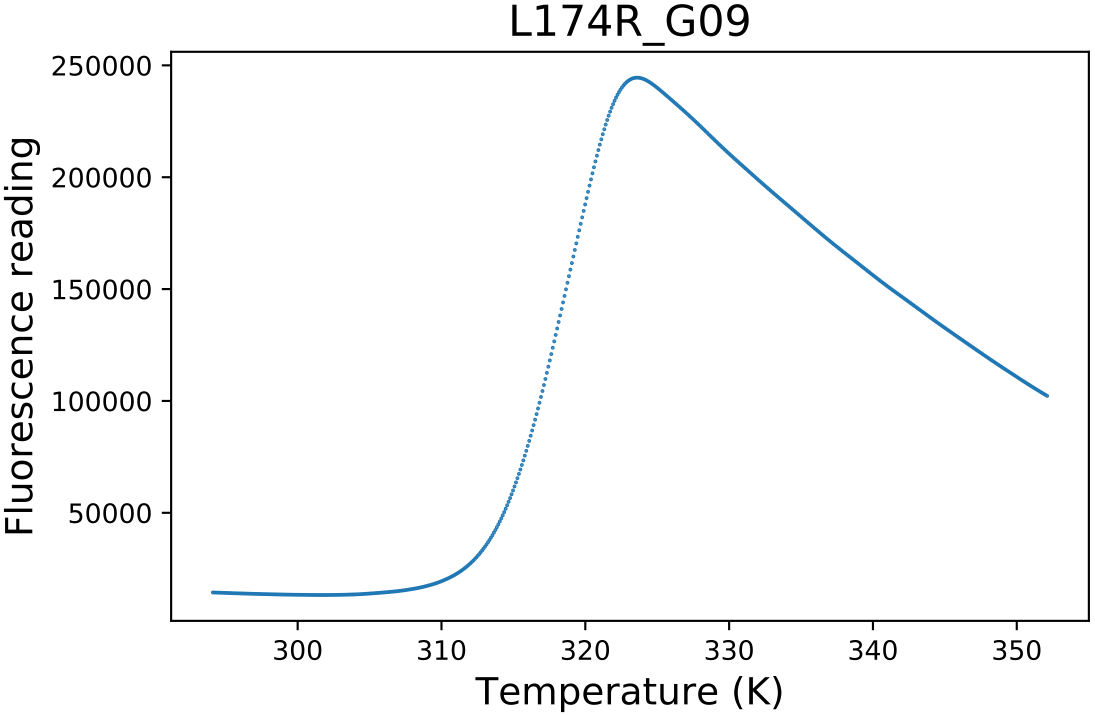

### lk_ball_wtd.png

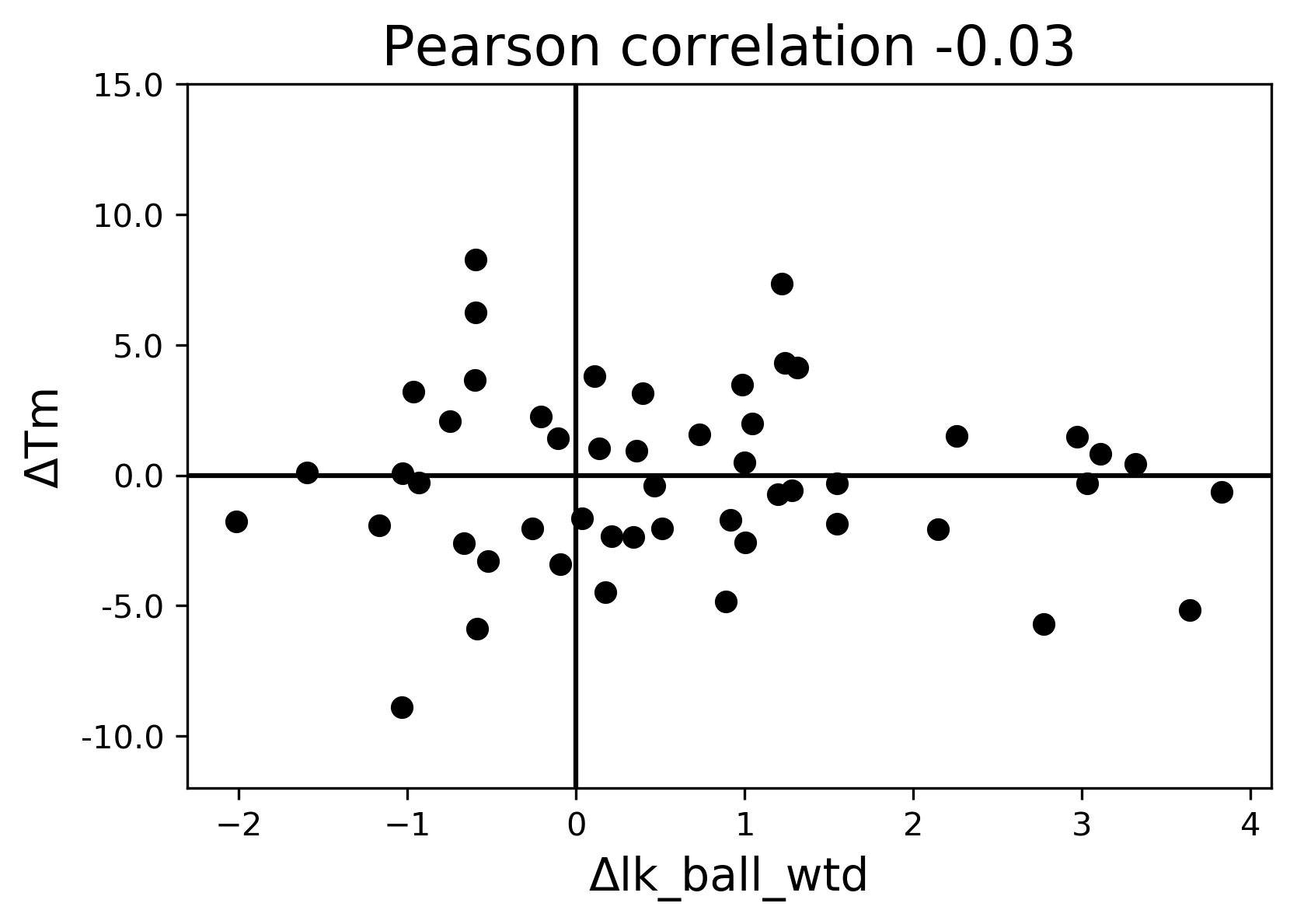

### M326A_D07_fluorescence.pdf

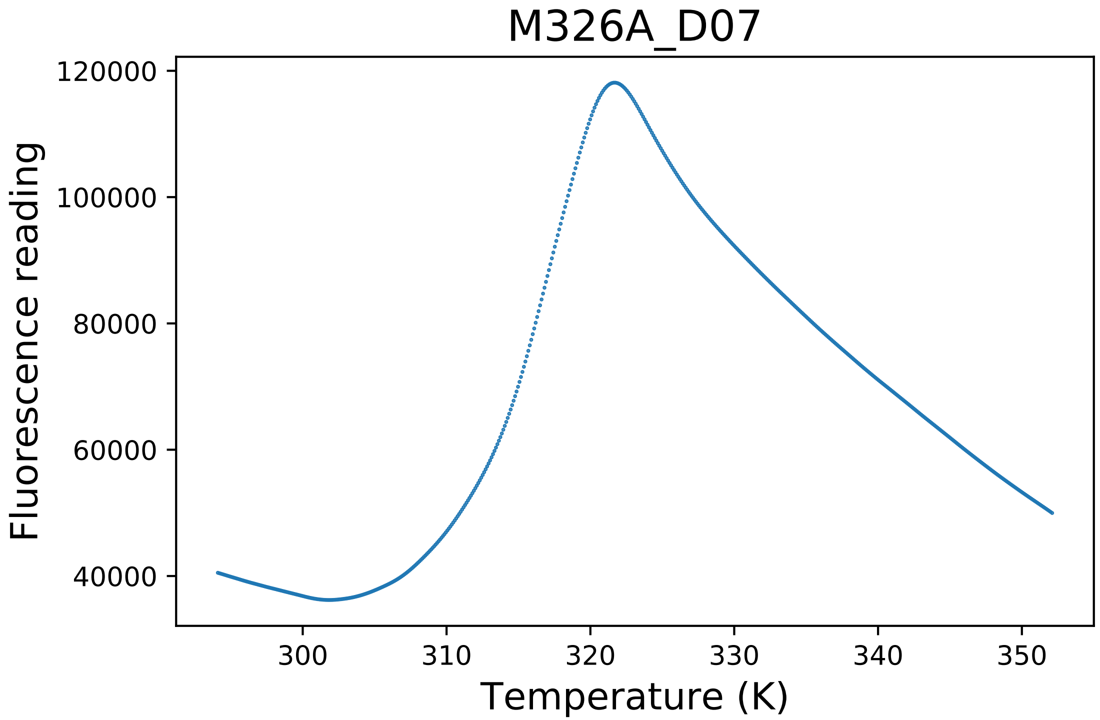

### M326A_D08_derivative.pdf

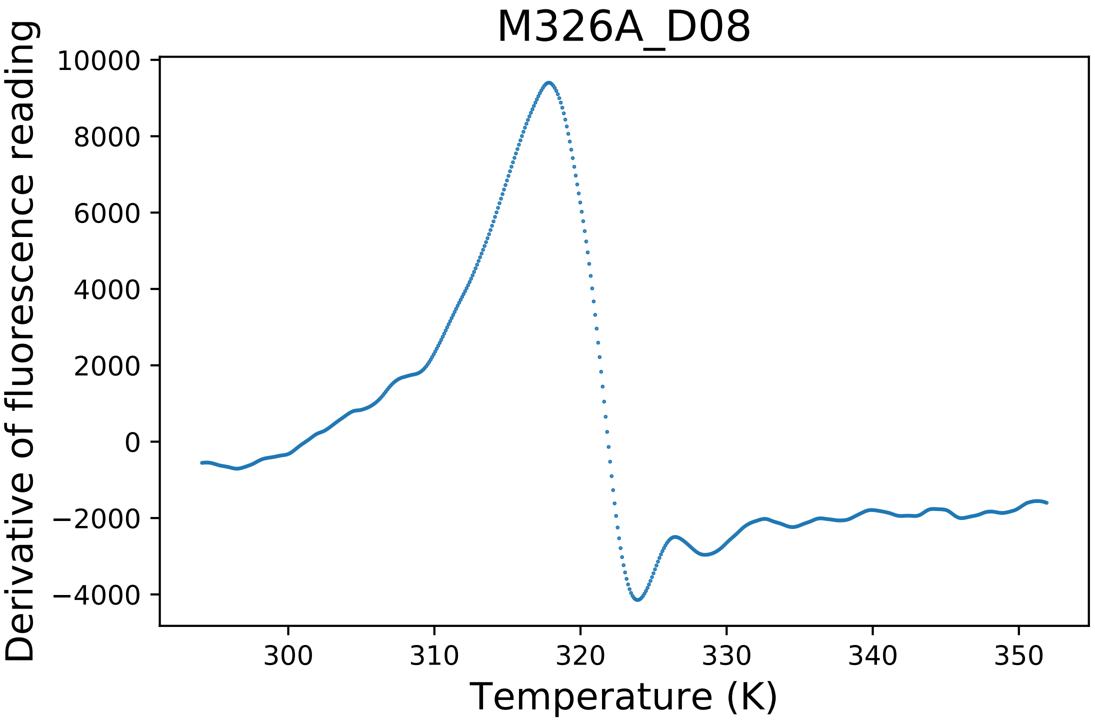

### N166A_B11_dg.pdf

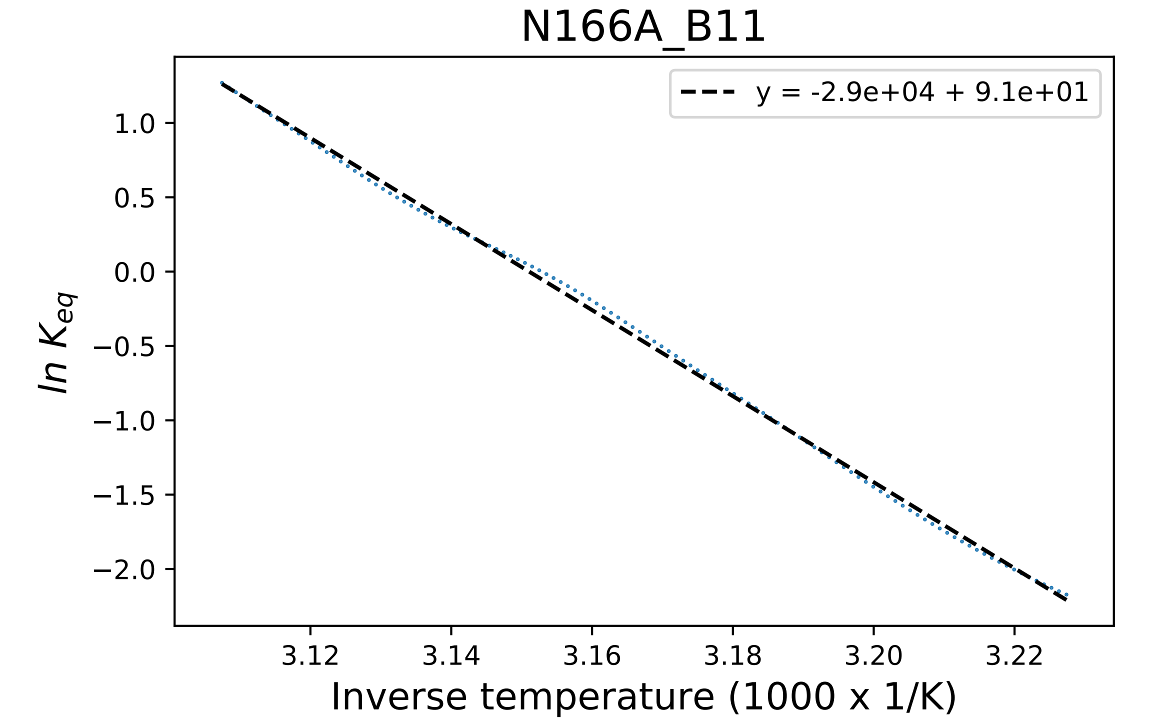

### N166C_F02_dg.pdf

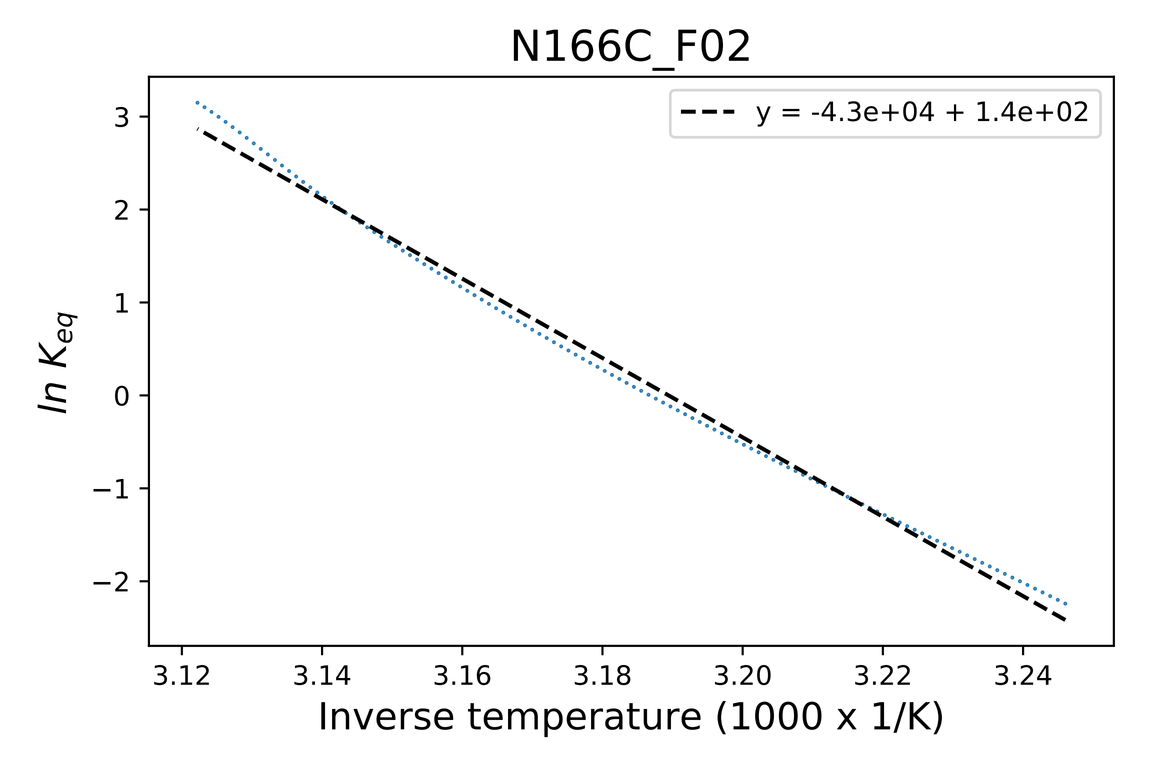

### N223A_F07_derivative.pdf

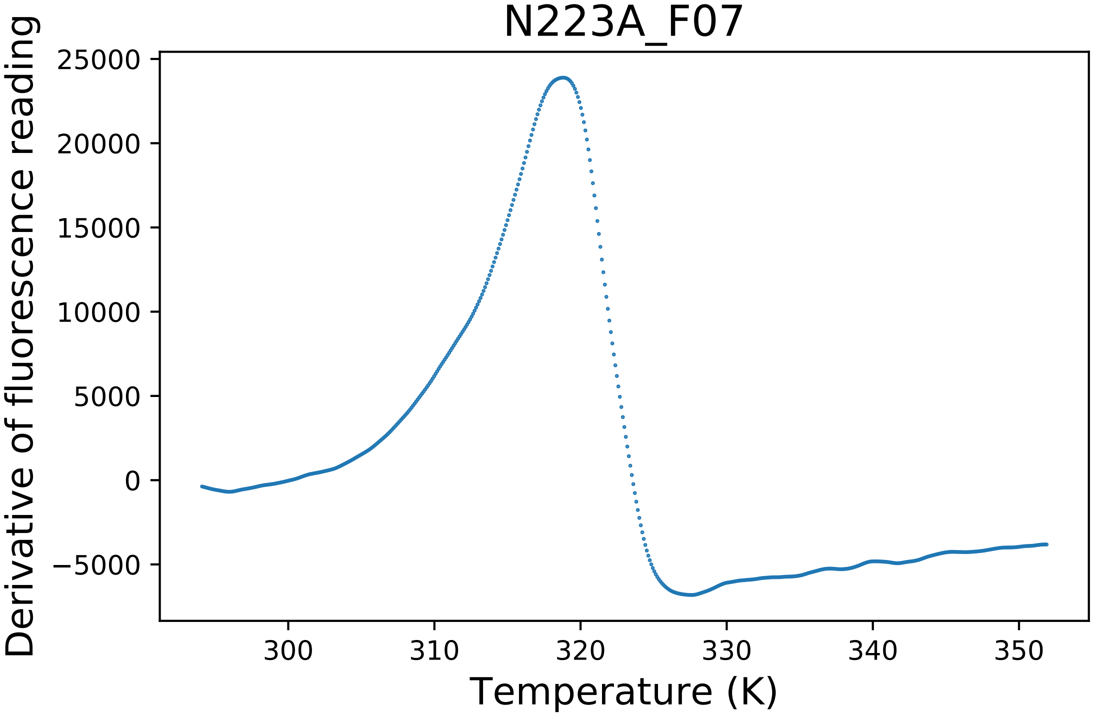

### N223G_C01_fluorescence.pdf

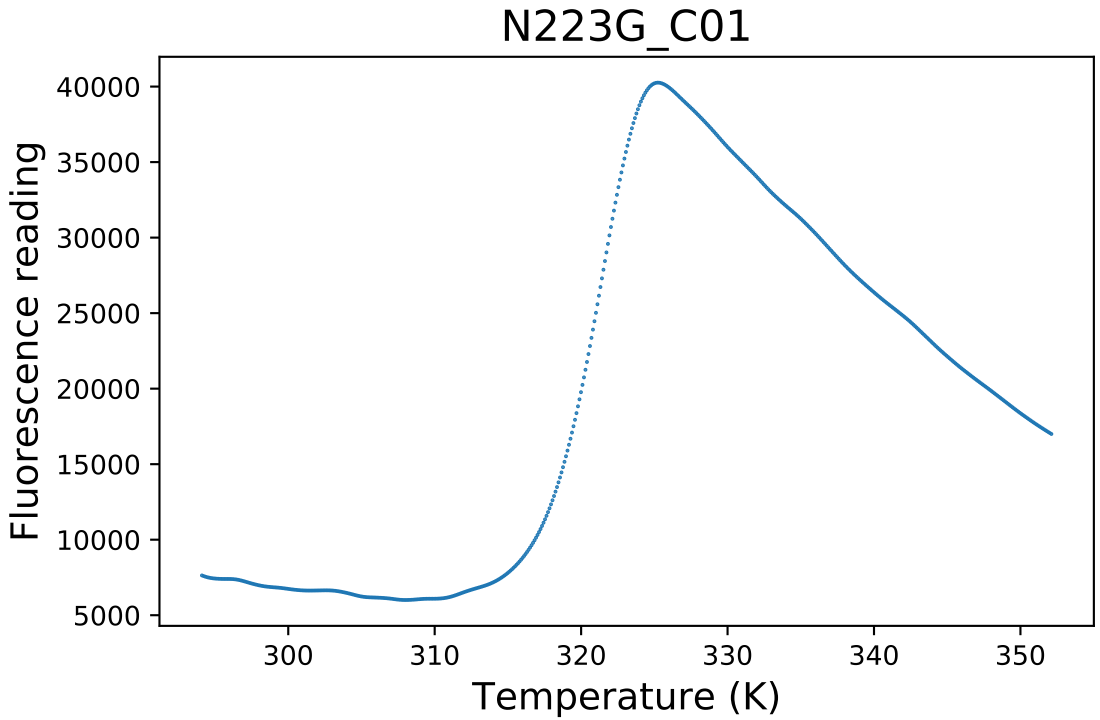

### N223G_C03_dg.pdf

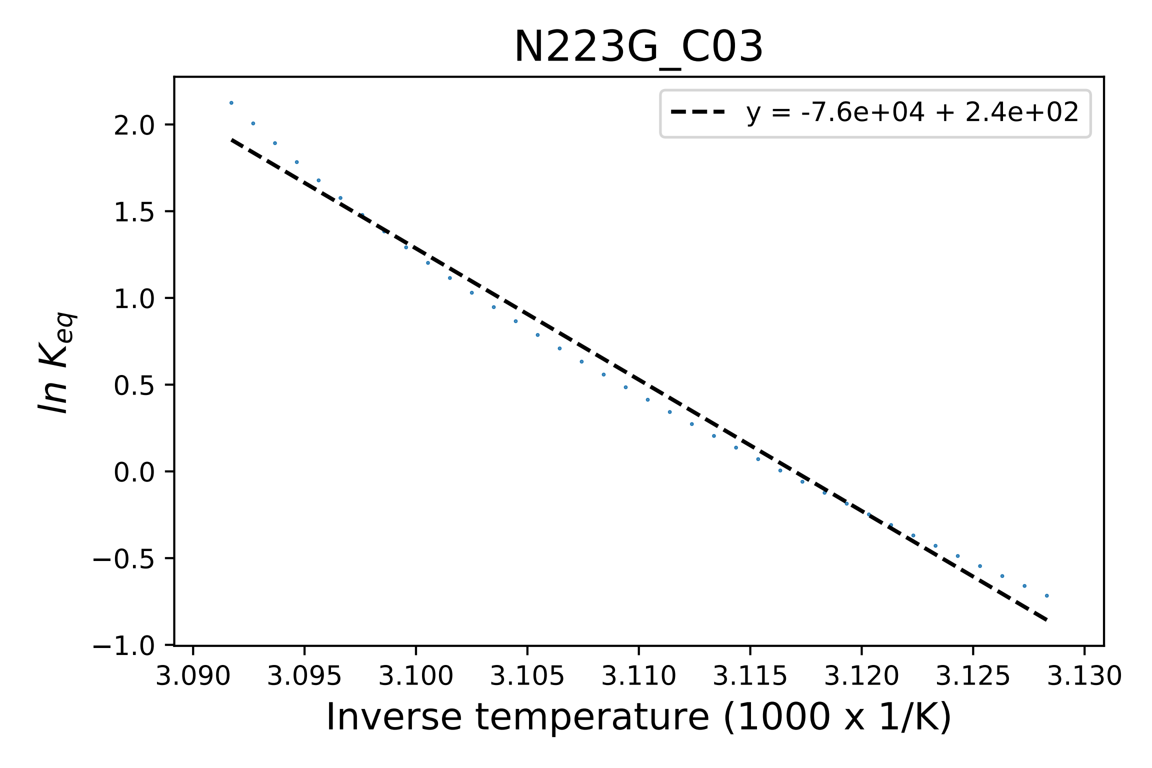

### N223R_G02_fluorescence.pdf

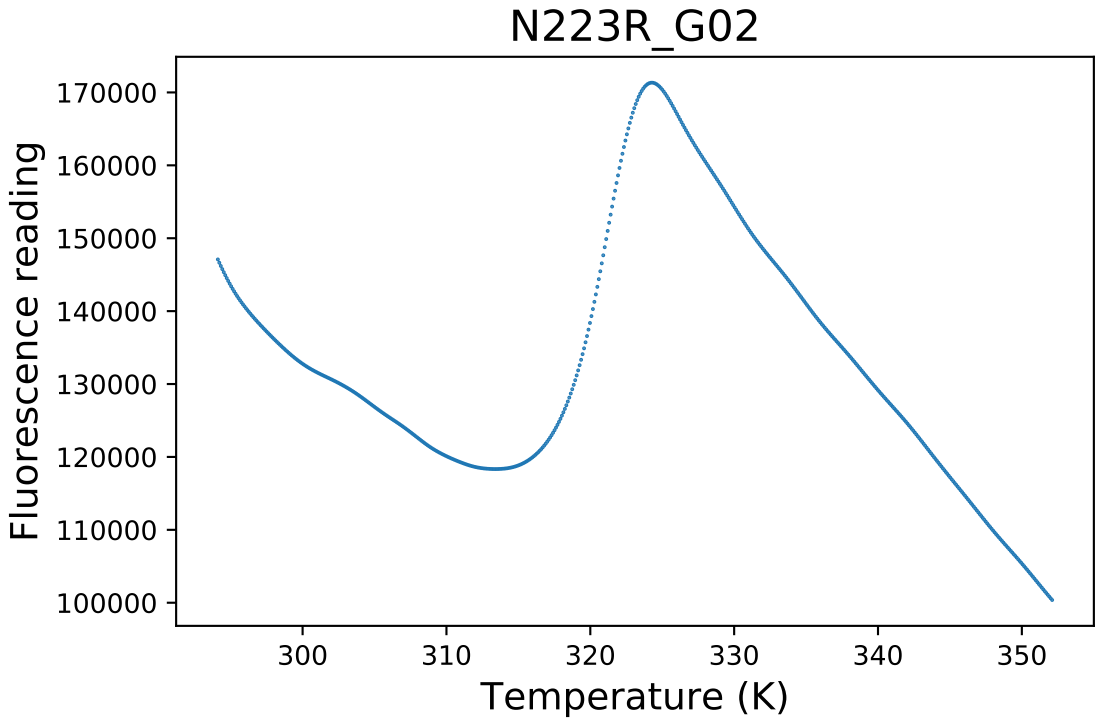

### N223R_G03_dg.pdf

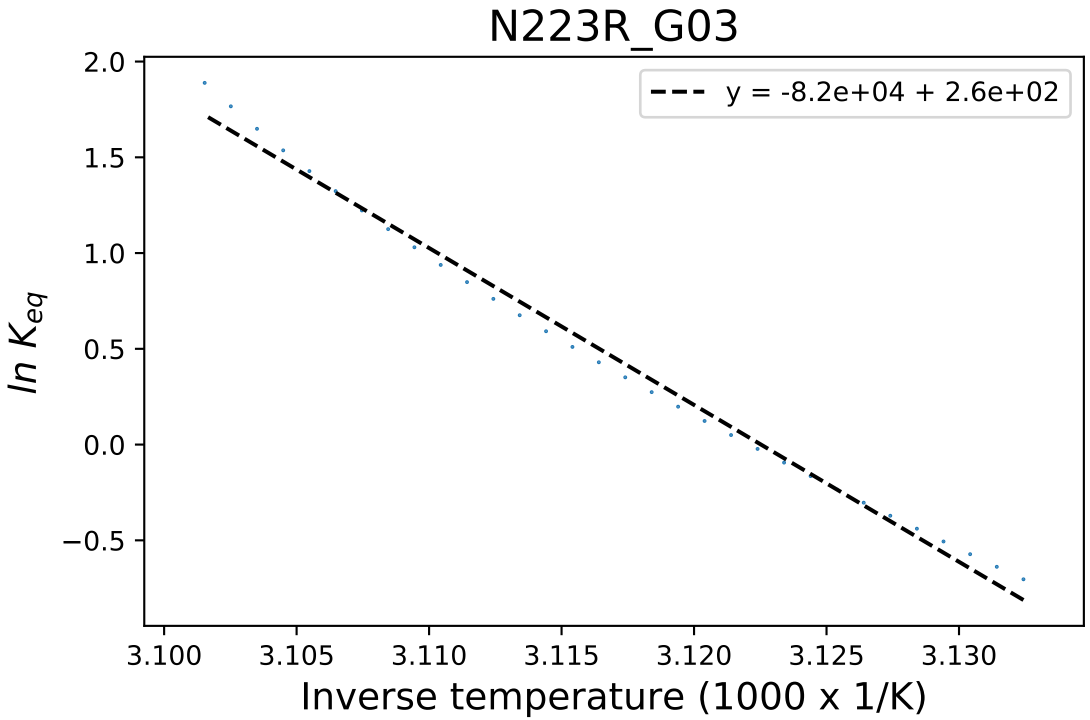

### N223Y_B02_fluorescence.pdf

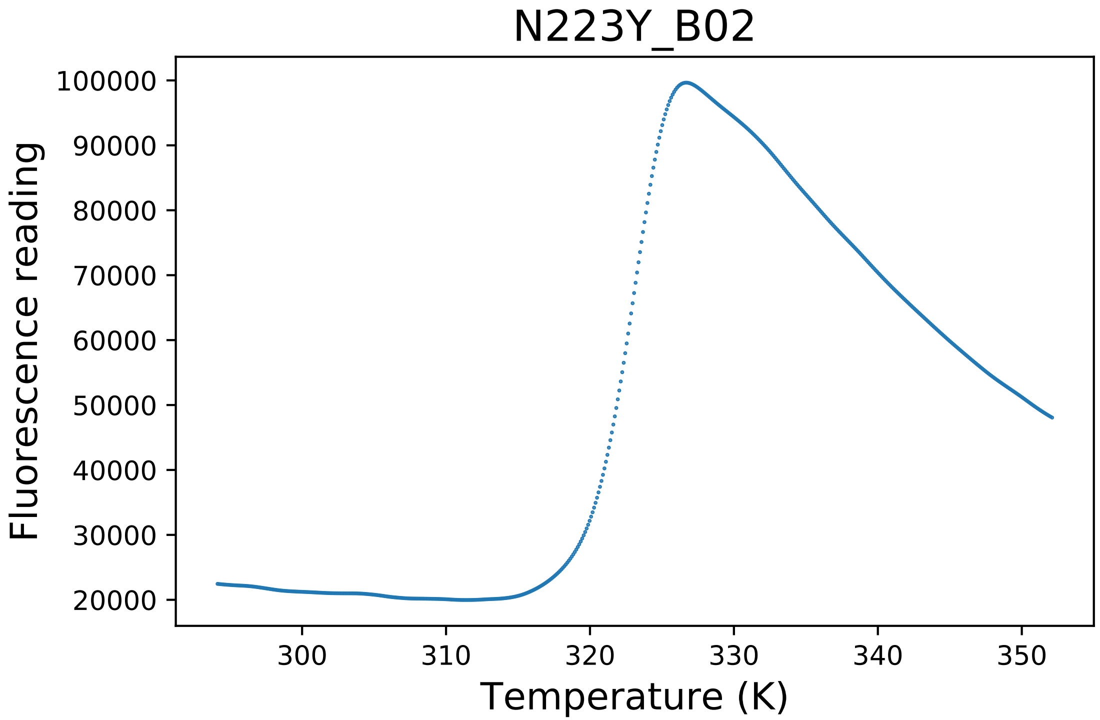

### N296A_B07_fluorescence.pdf

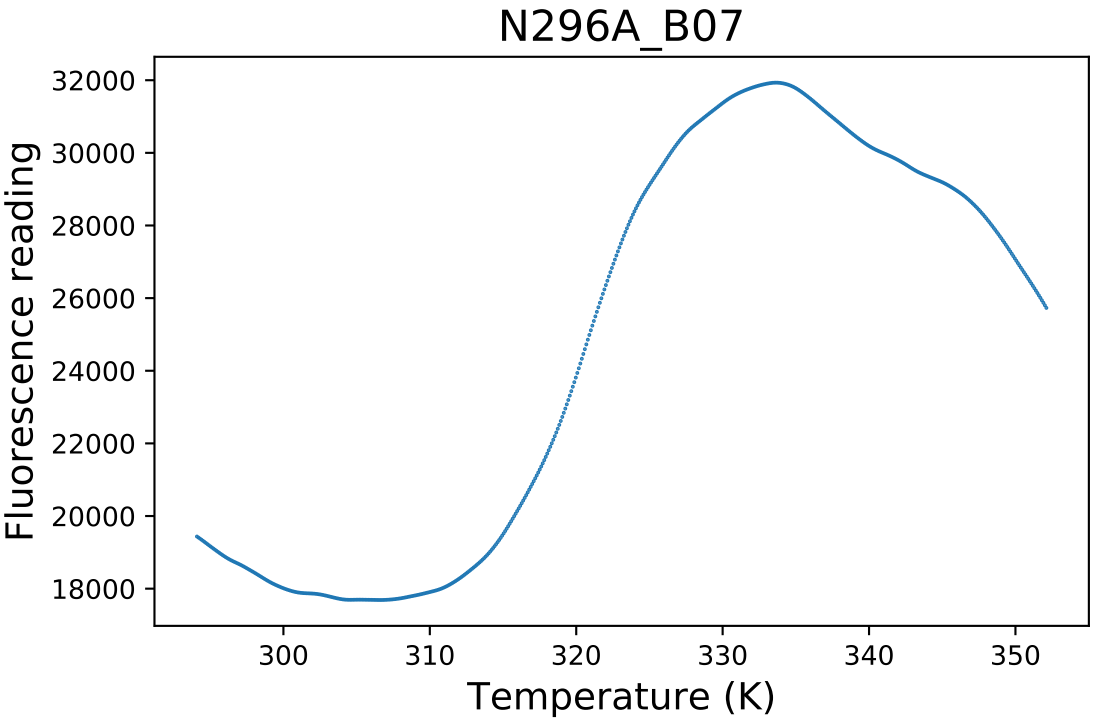

### N296C_B06_dg.pdf

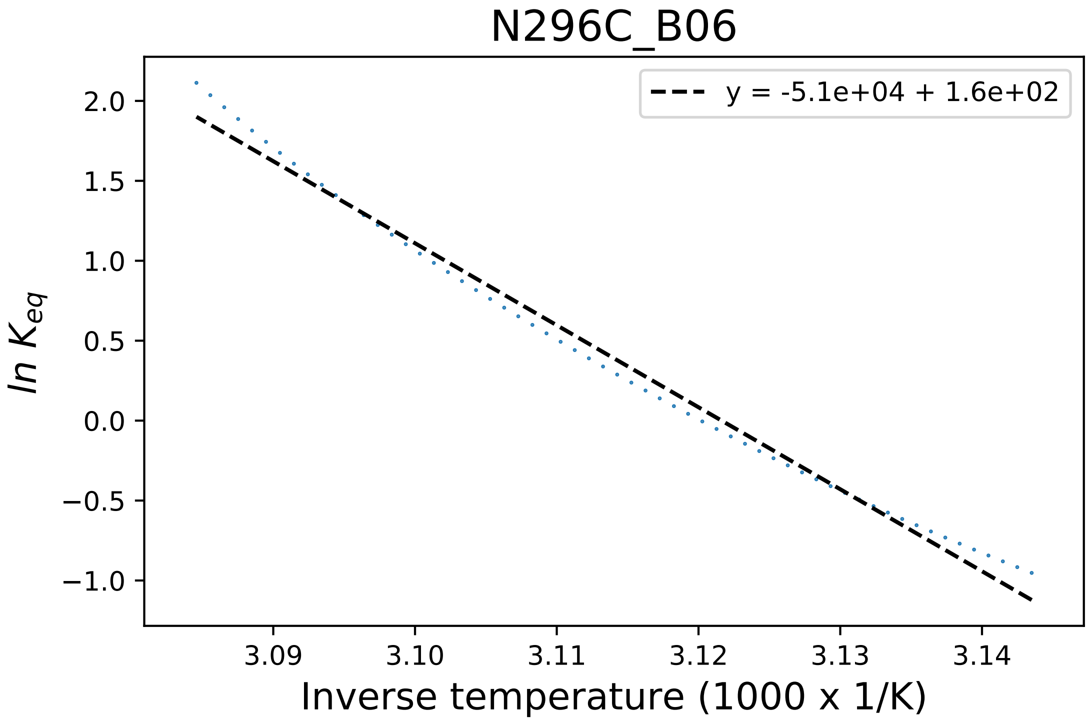

### N407A_A02_fluorescence.pdf

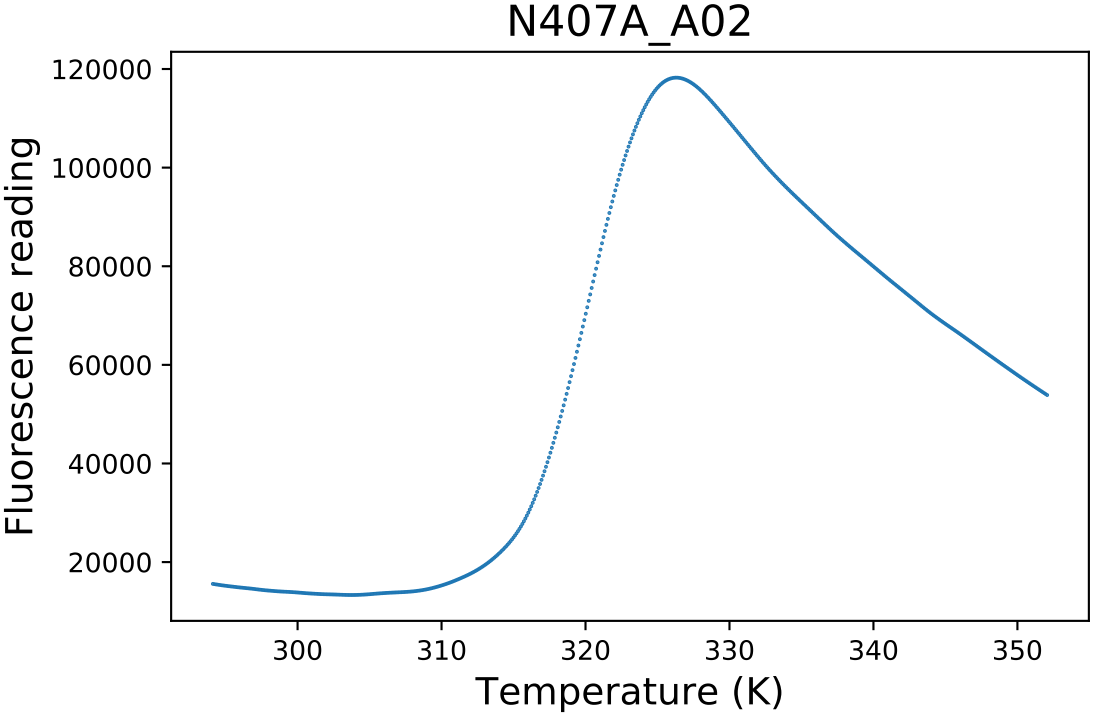

### N407A_A04_dg.pdf

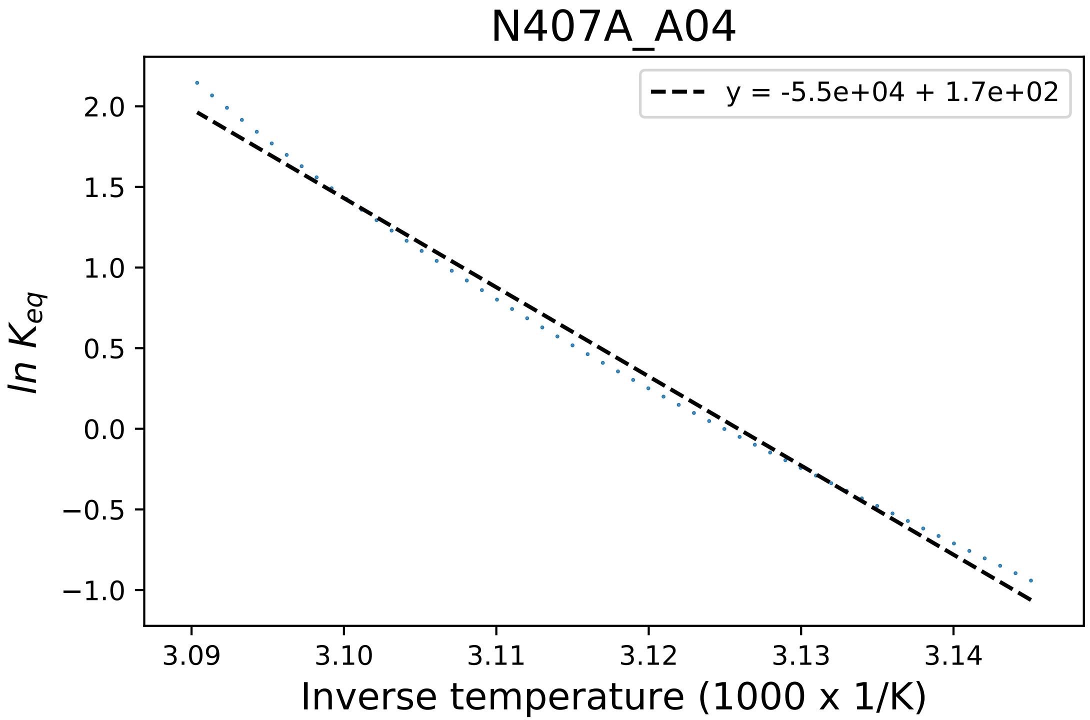

### N407C_C01_dg.pdf

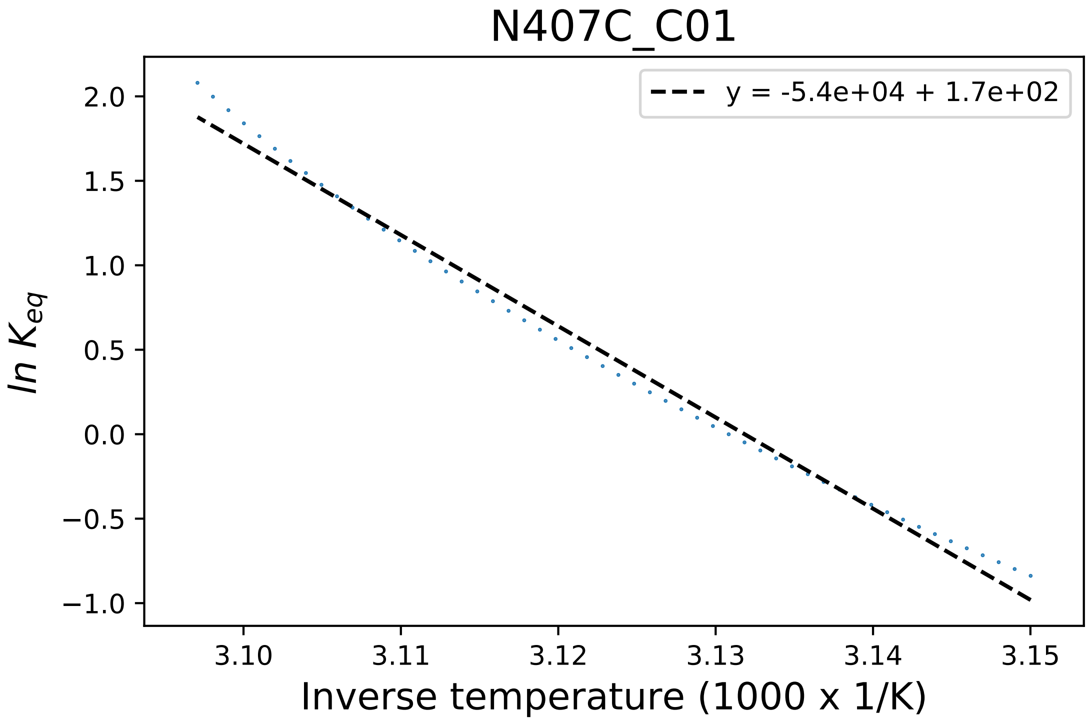

### N407C_C03_derivative.pdf

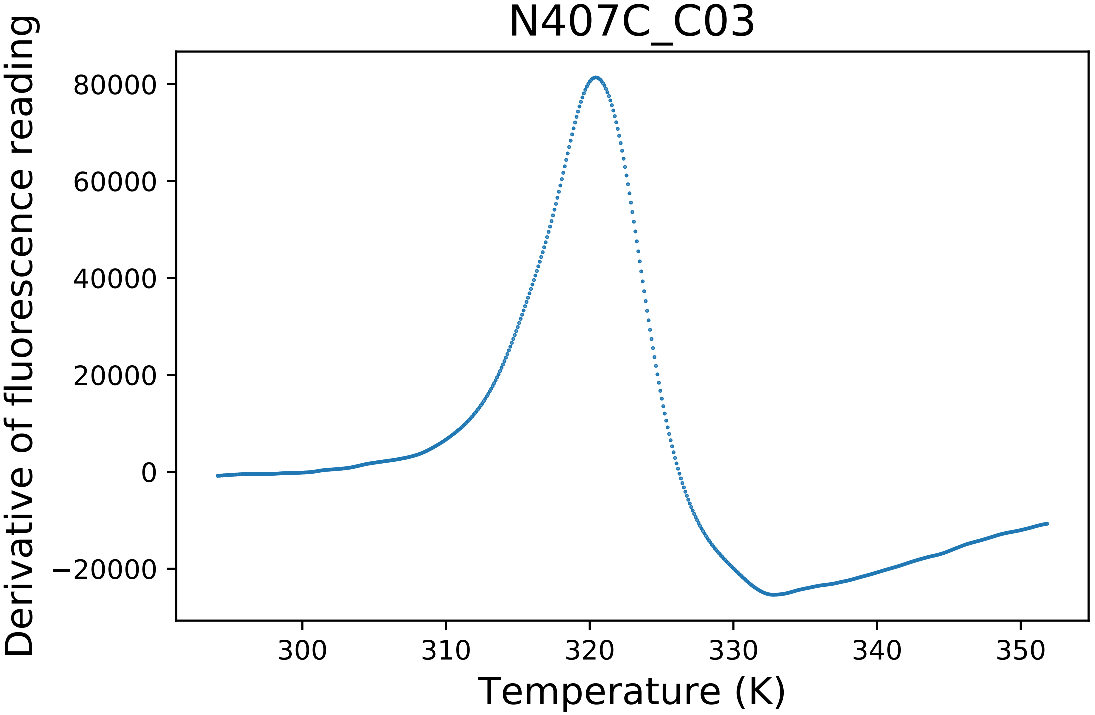

### omega.png

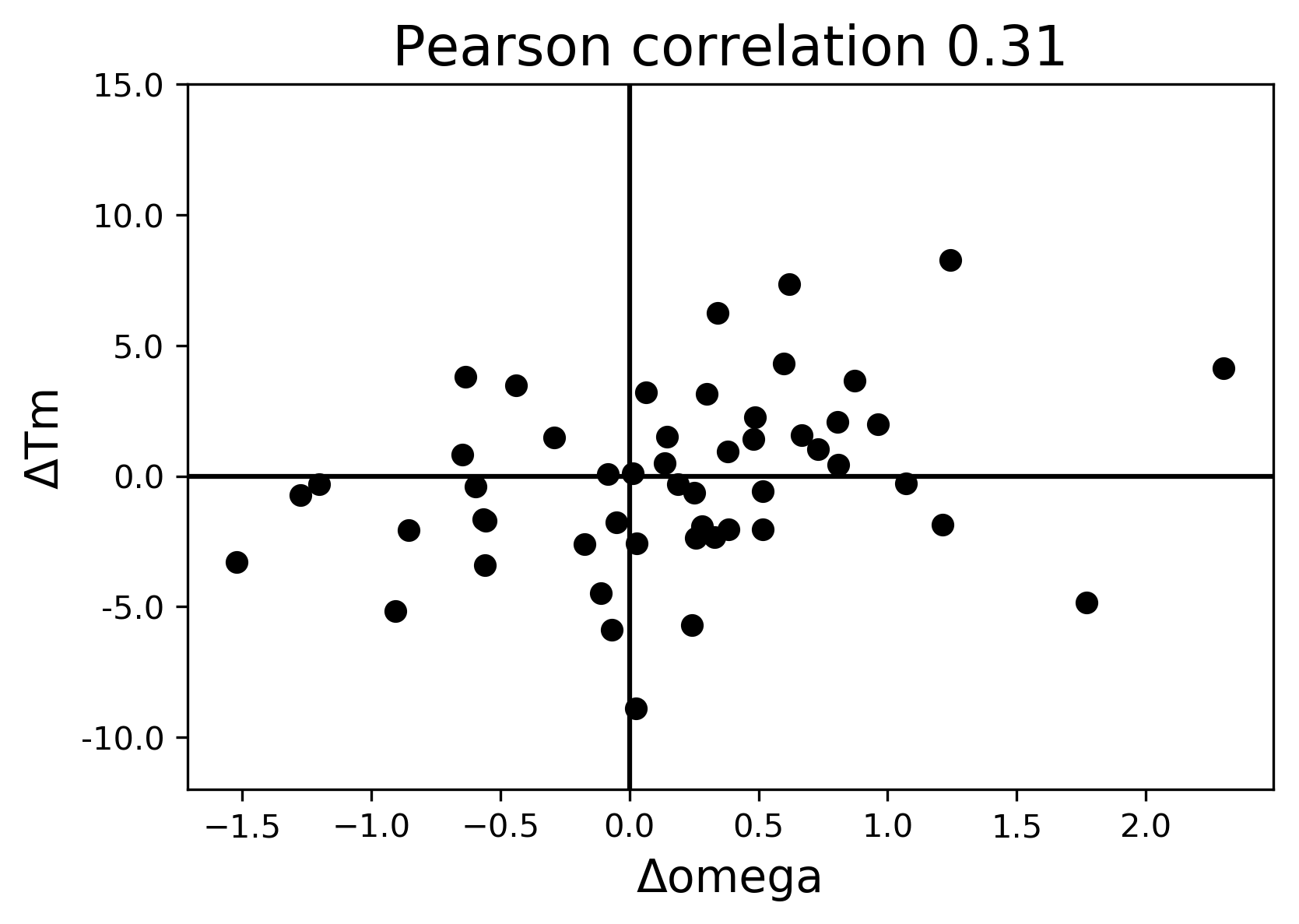

### p_aa_pp.png

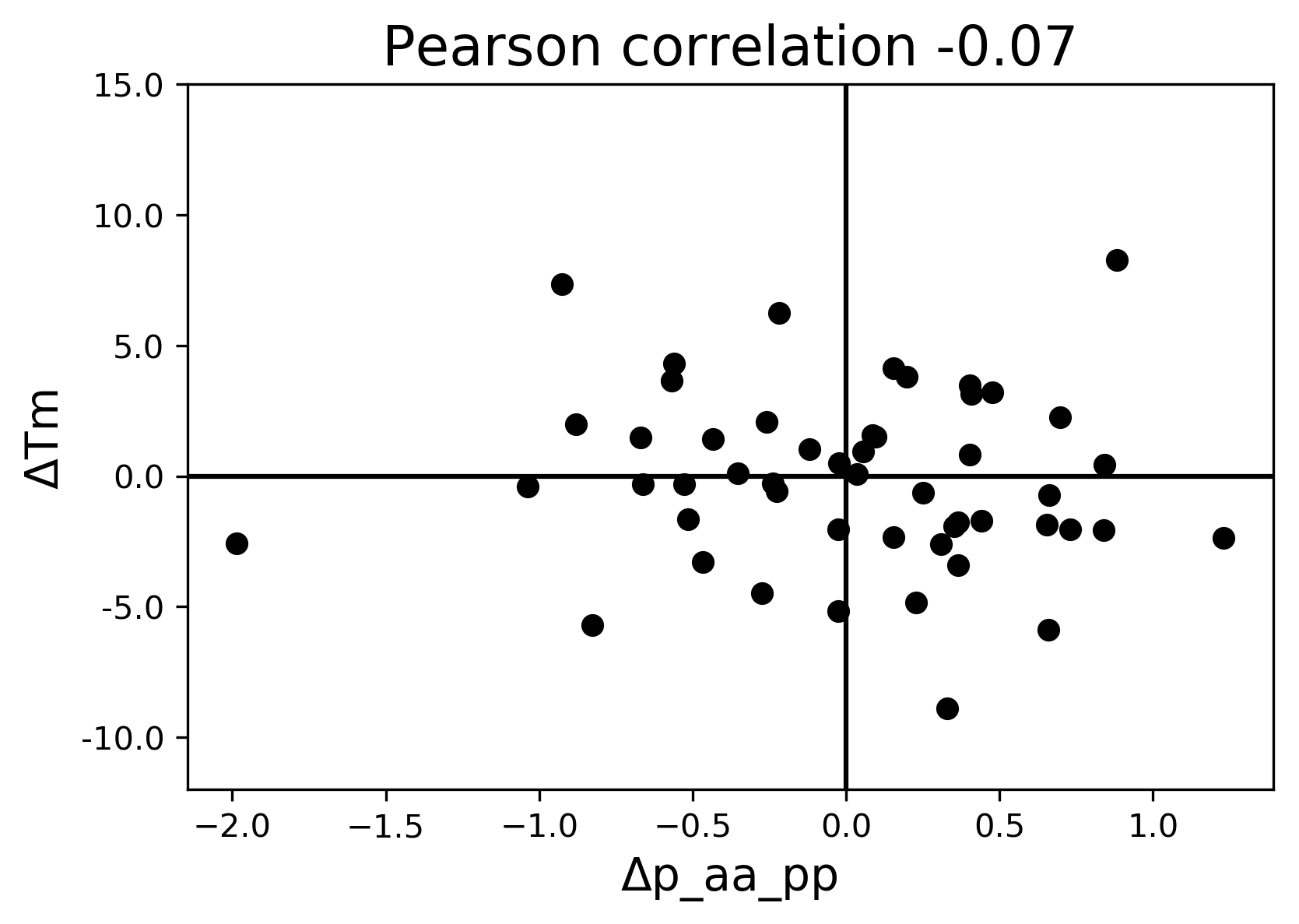

### pro_close.png

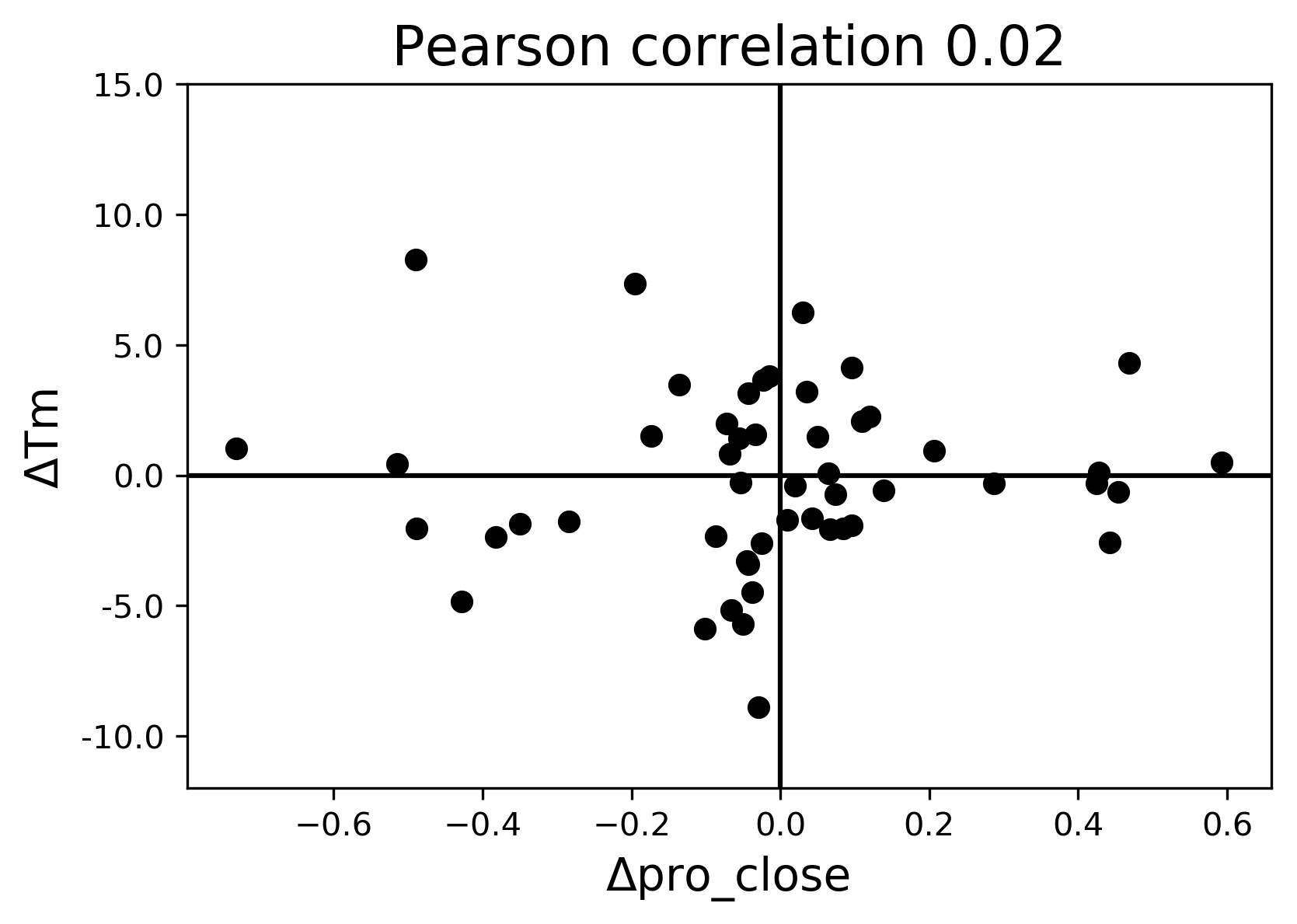

### Q22S_B07_derivative.pdf

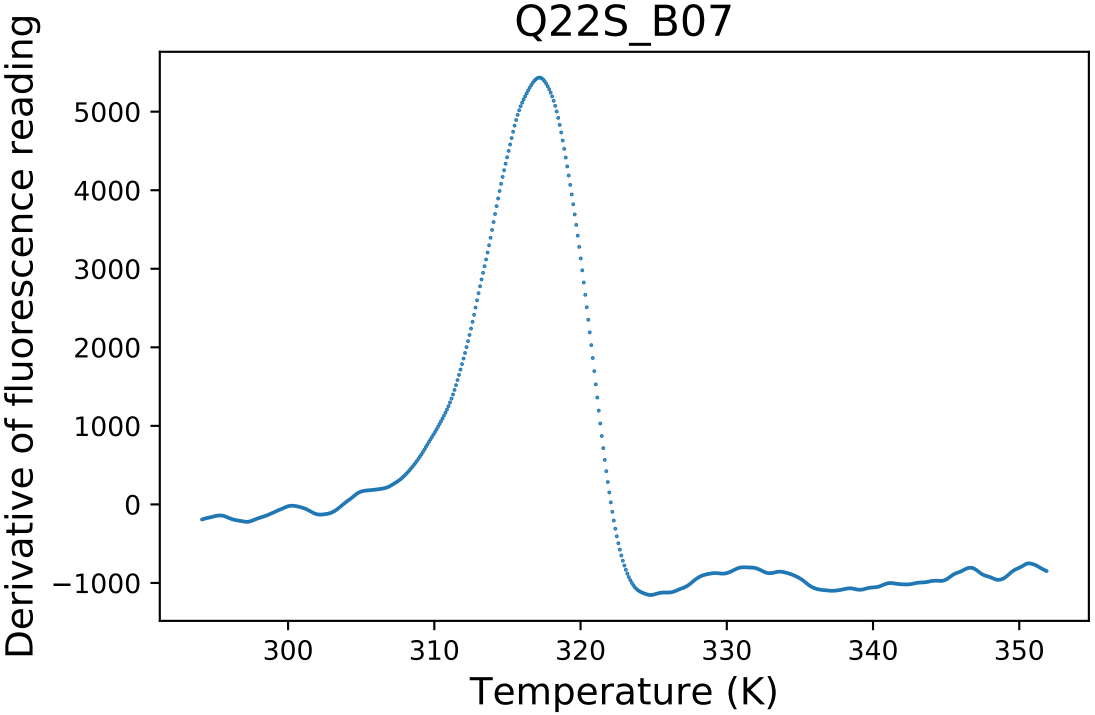

### R243A_A04_fluorescence.pdf

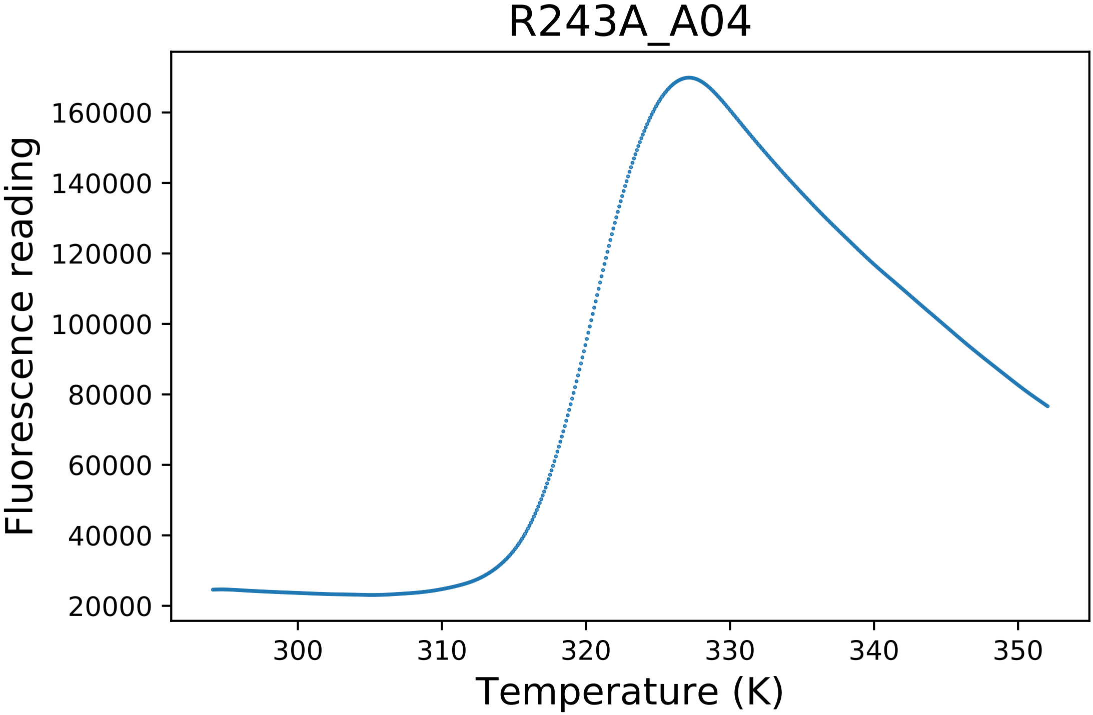

### R243D_E05_dg.pdf

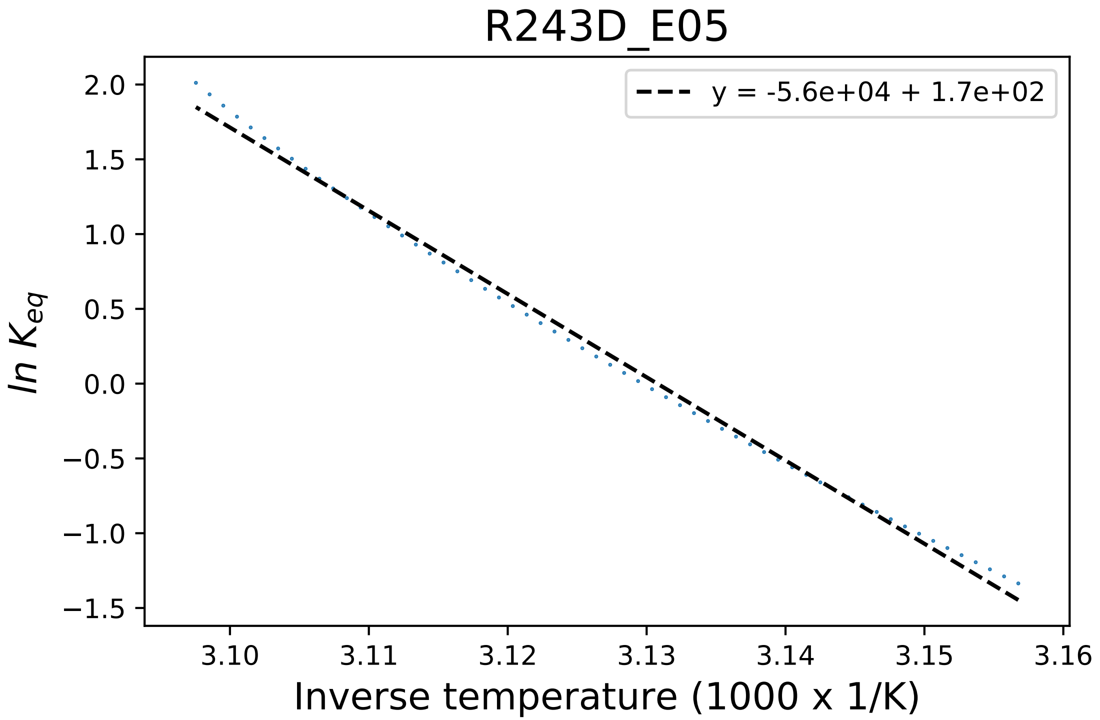

### R243K_C07_fluorescence.pdf

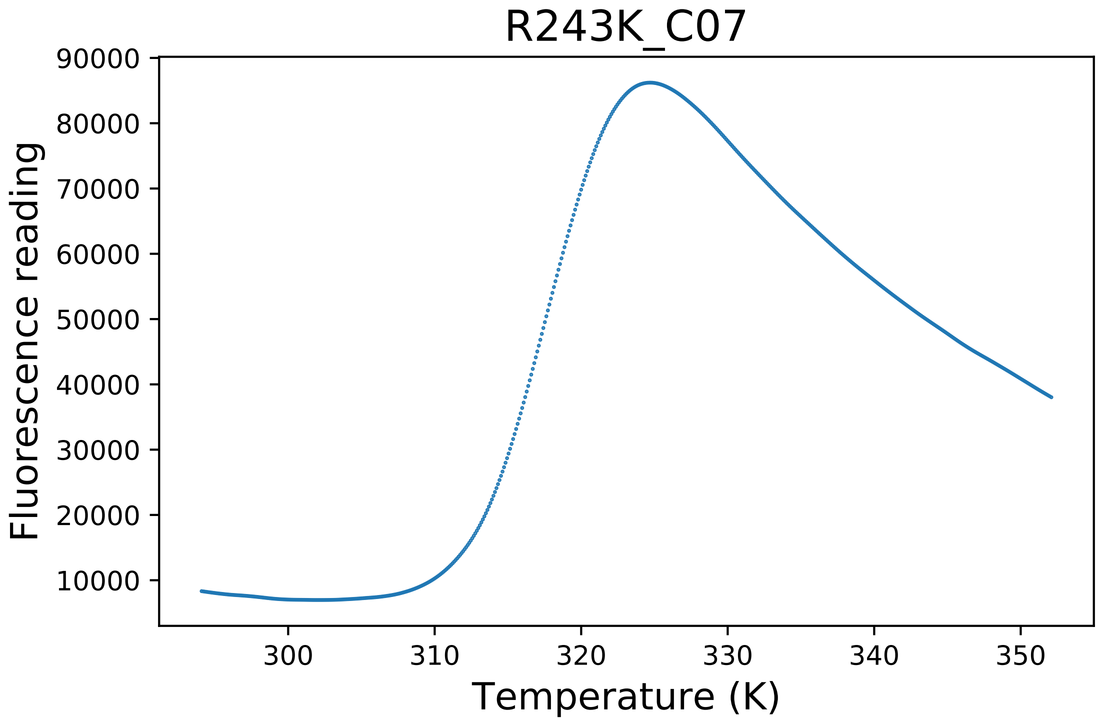

### rama_prepro.png

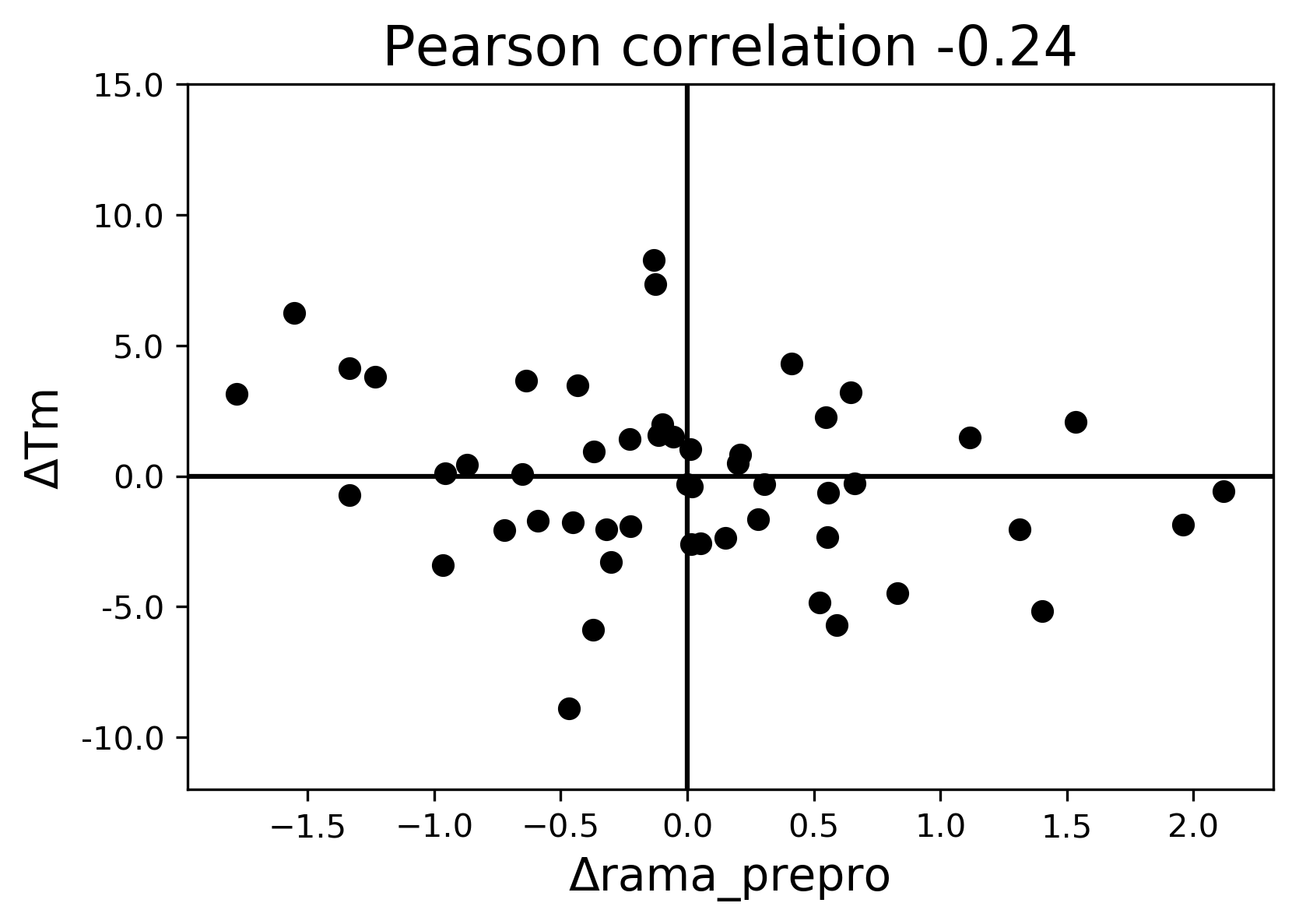

### ref.png

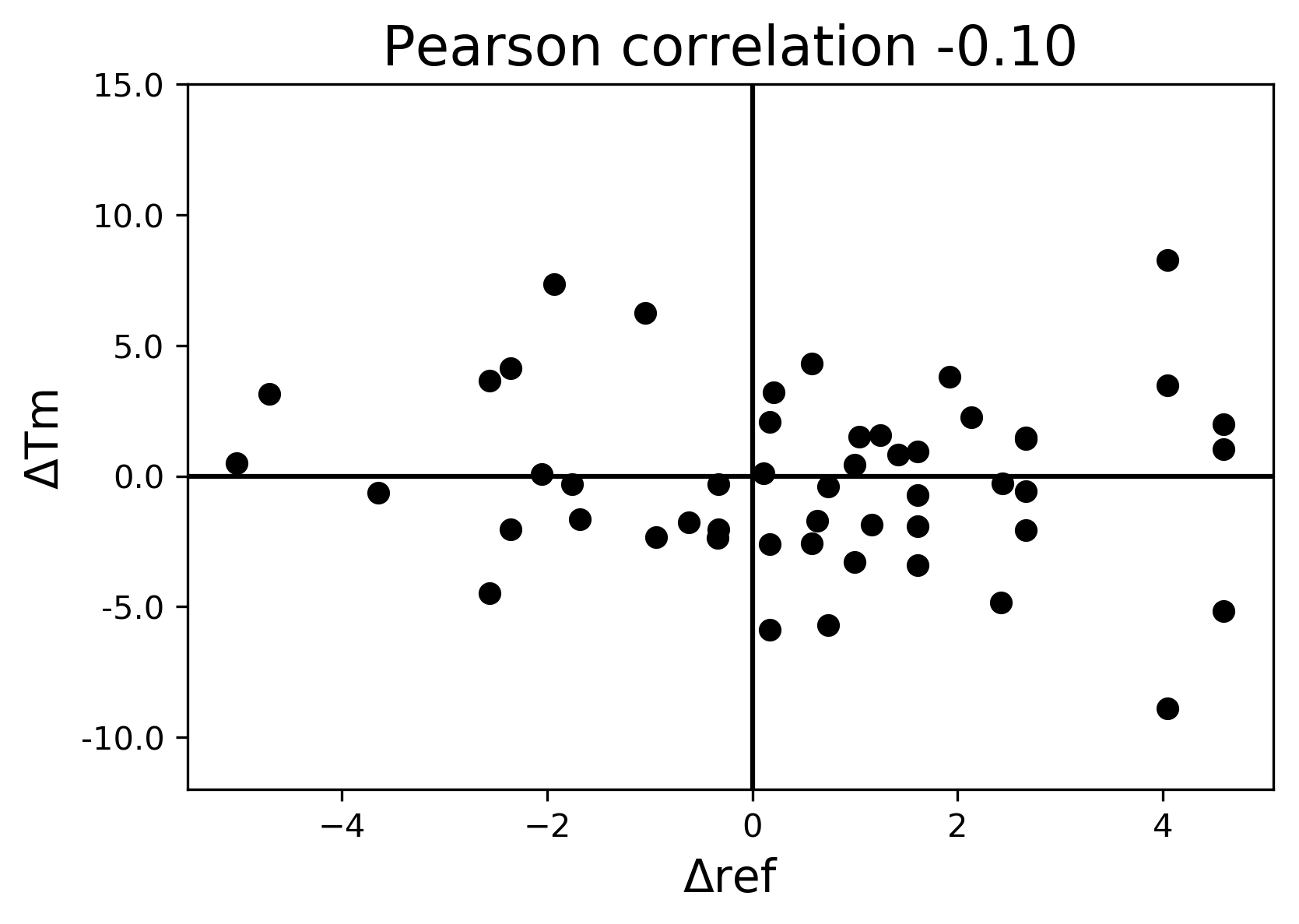

### S17A_G10_dg.pdf

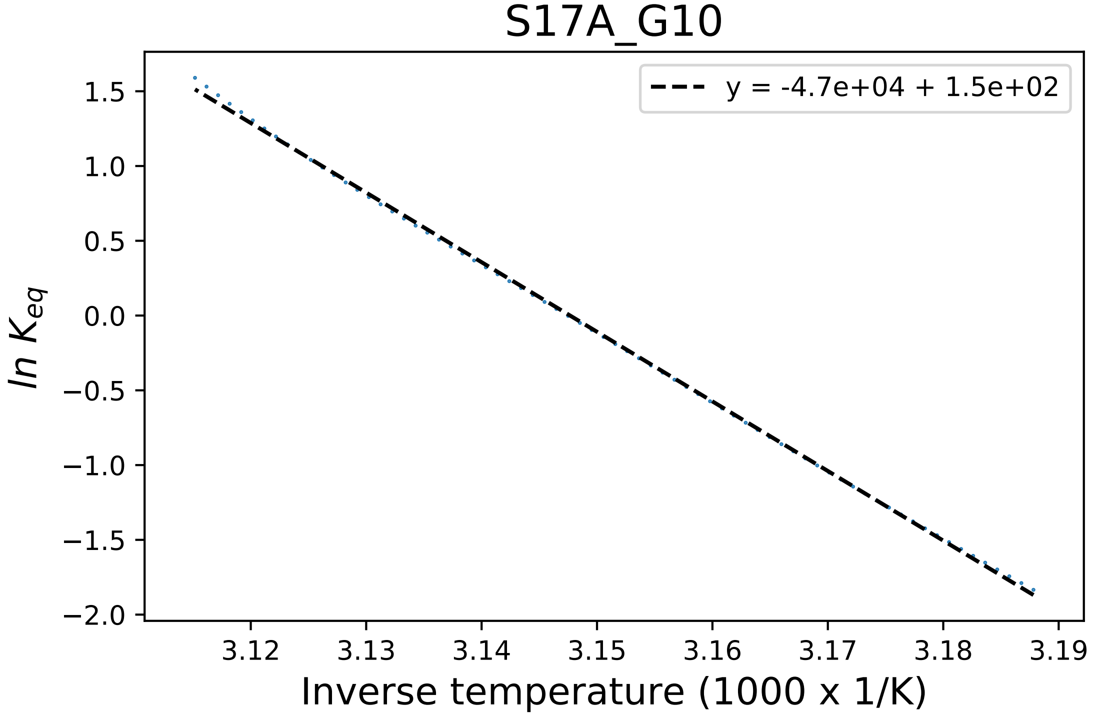

### S17A_G10_fluorescence.pdf

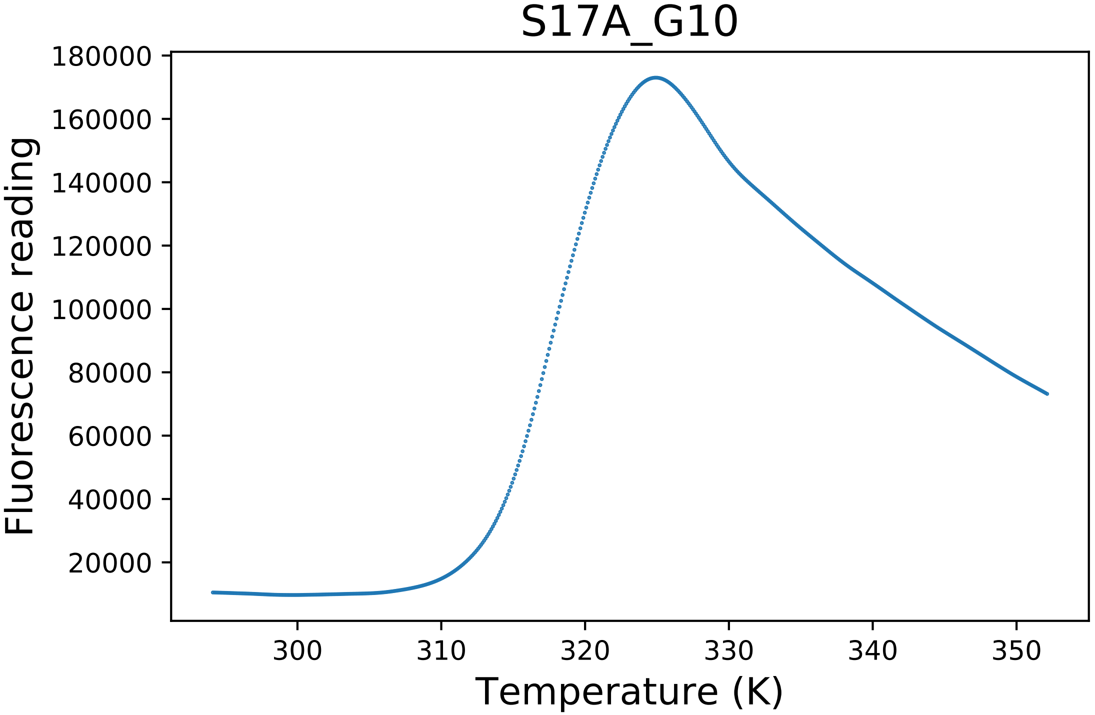

### S19A_A03_dg.pdf

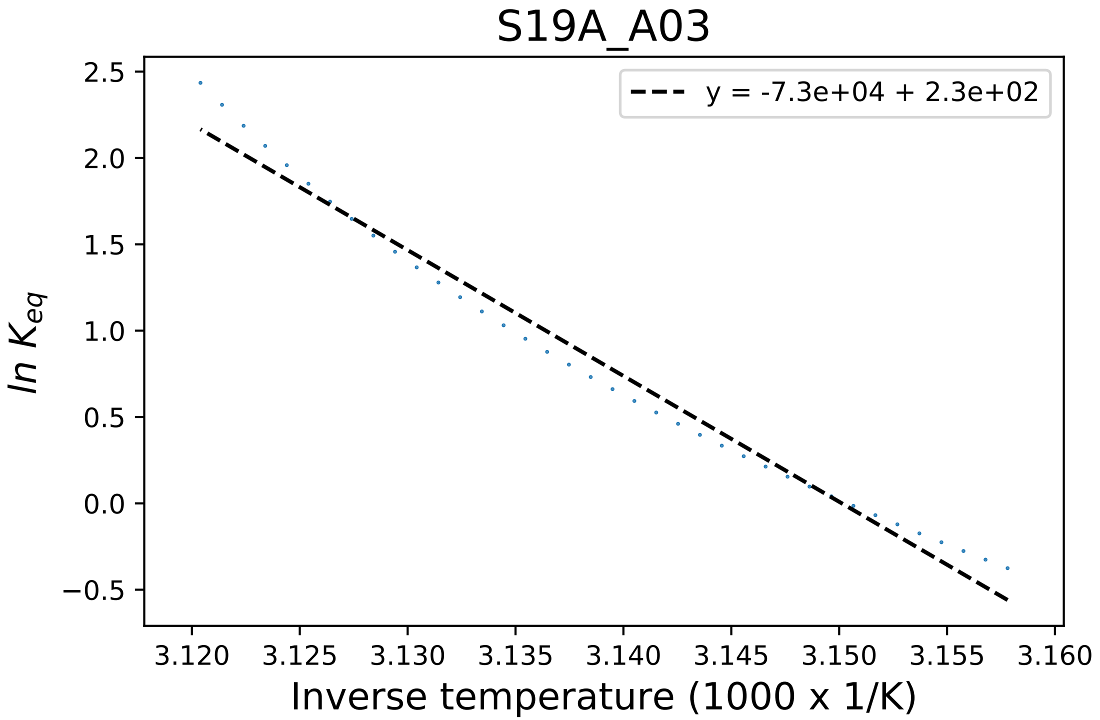

### S19A_A04_fluorescence.pdf

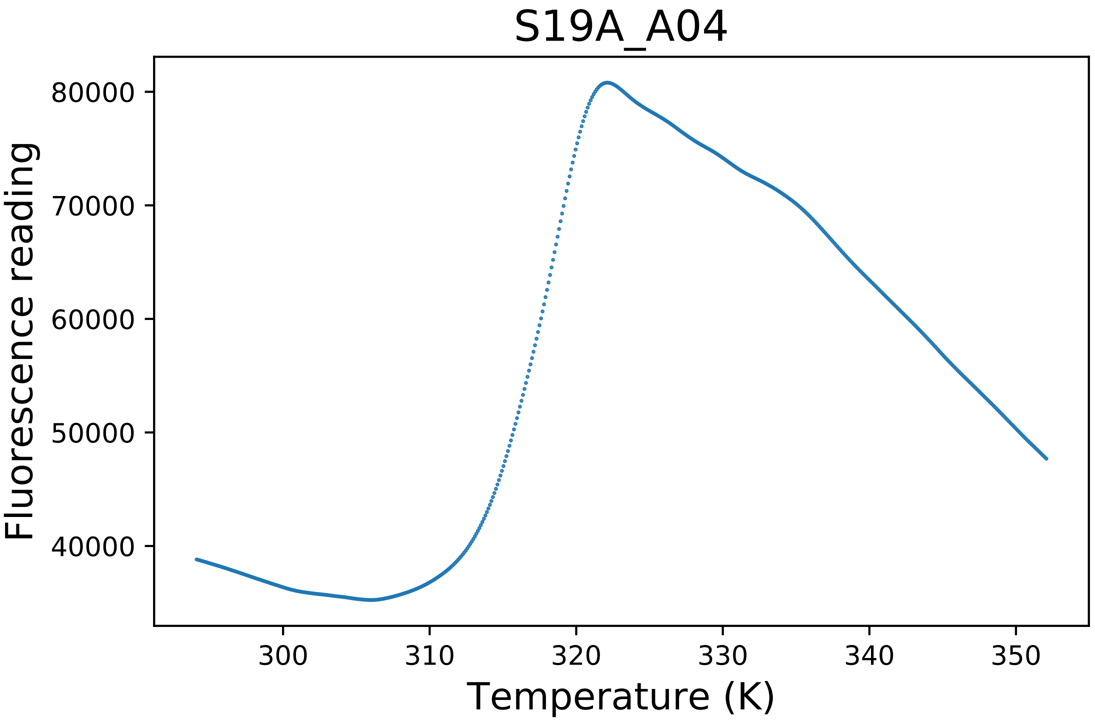

### S334A_B01_dg.pdf

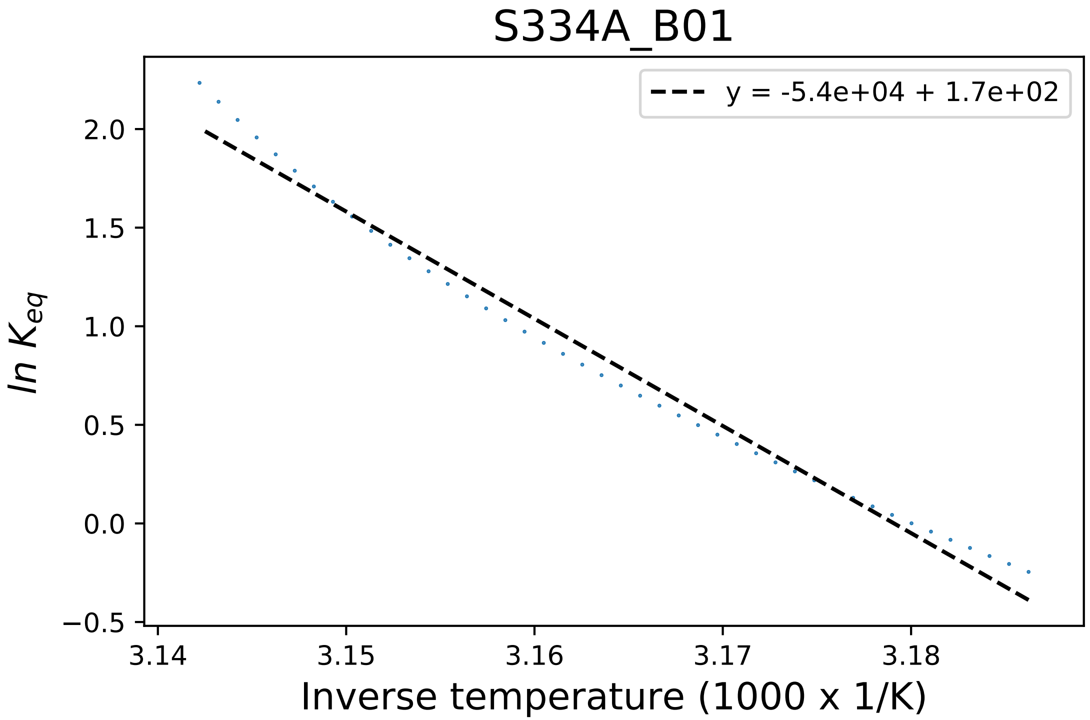

### S334A_B04_derivative.pdf

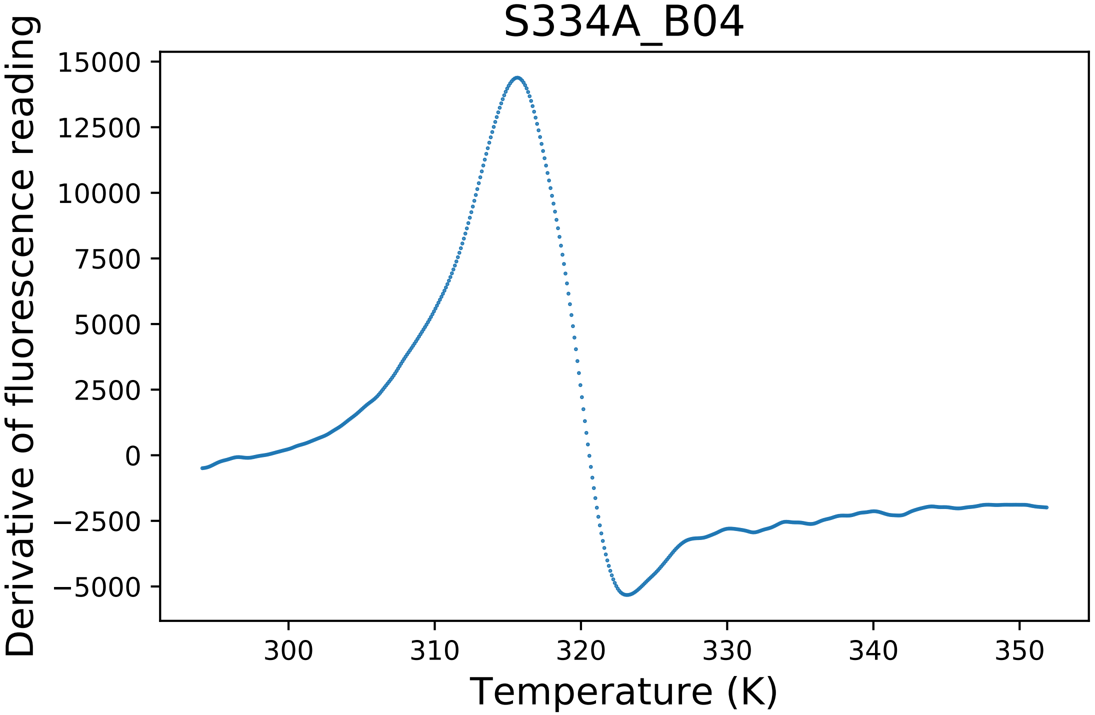

### S334A_B04_fluorescence.pdf

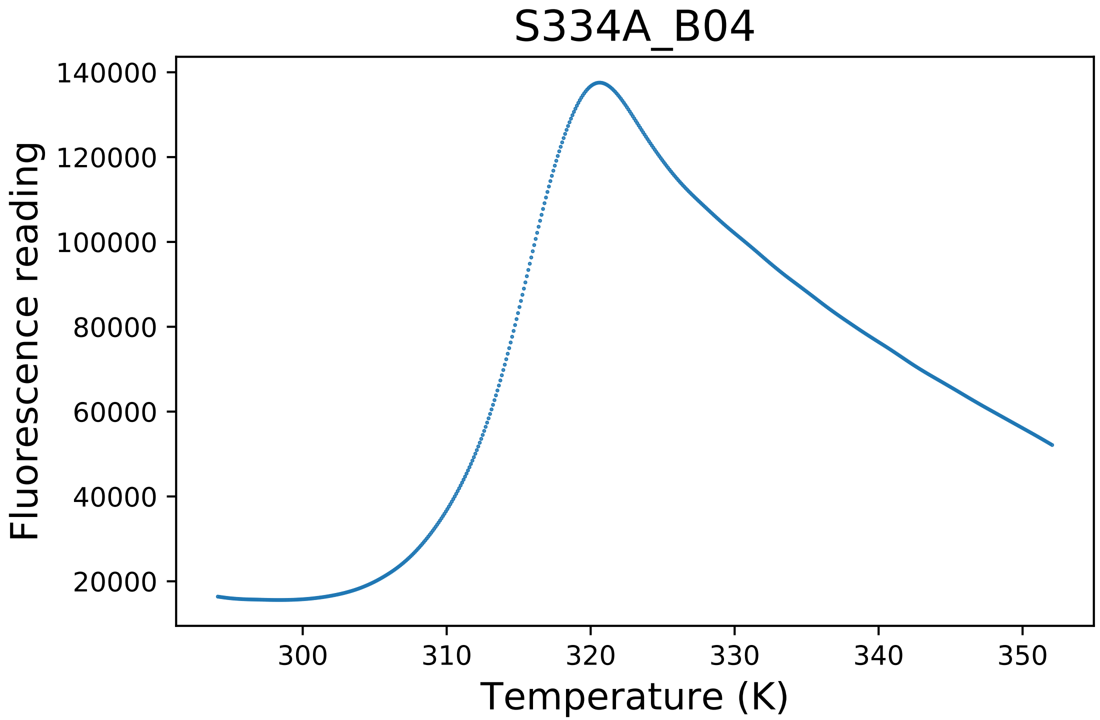

### S403A_E09_dg.pdf

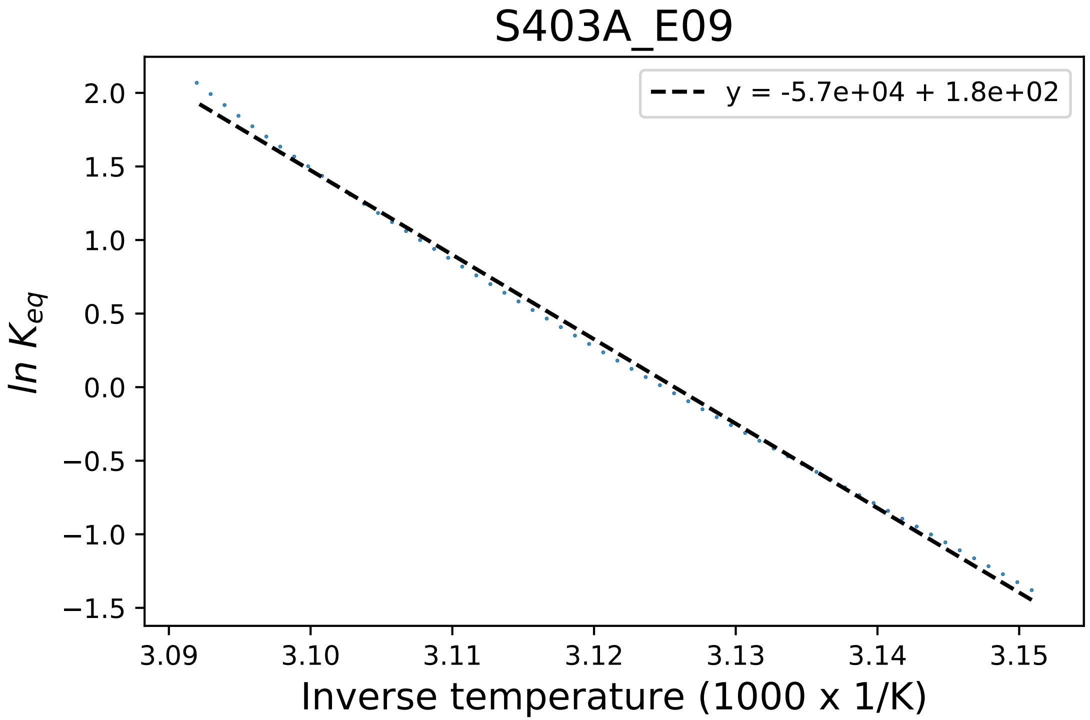

### S403A_E10_fluorescence.pdf

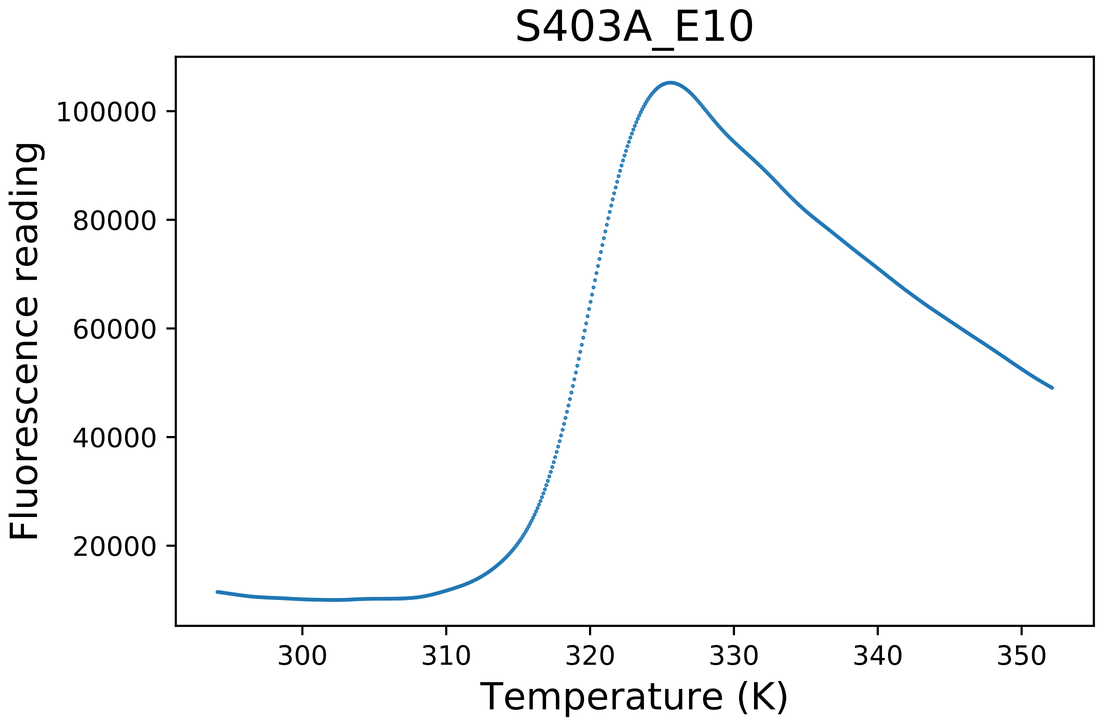

### Sidechain Hbond.png

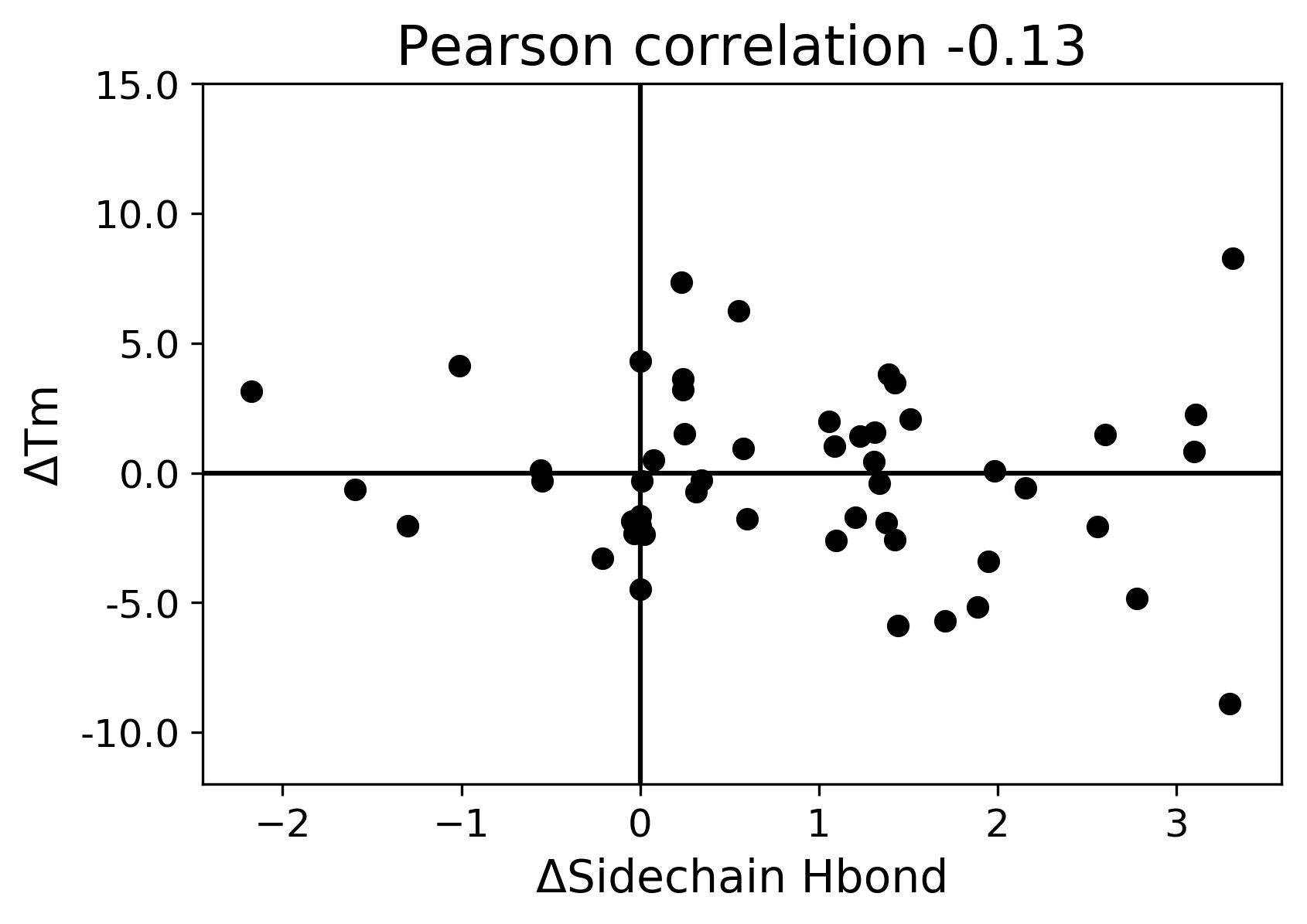

### Solvation Hydrophobic.png

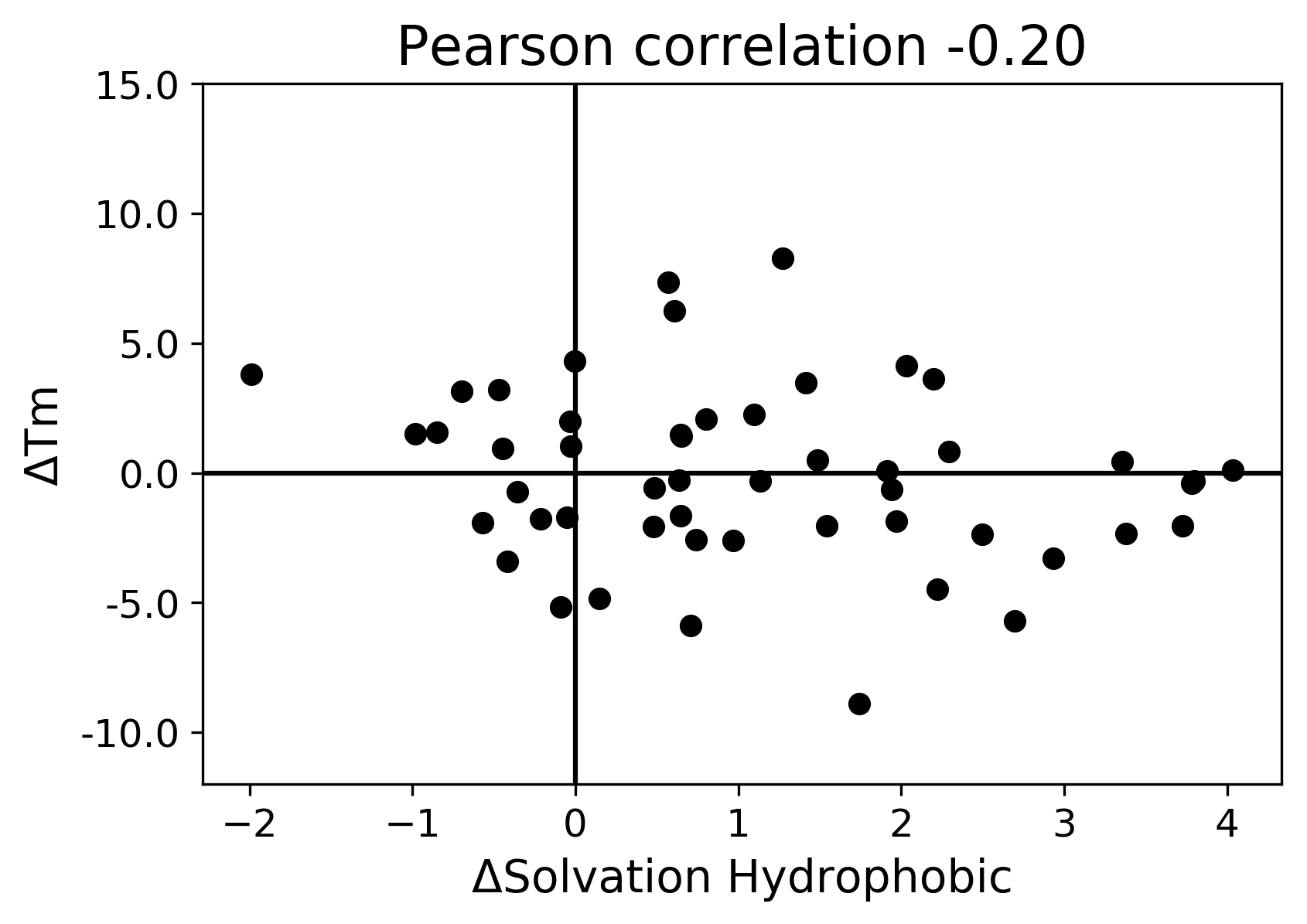

### Solvation Polar.png

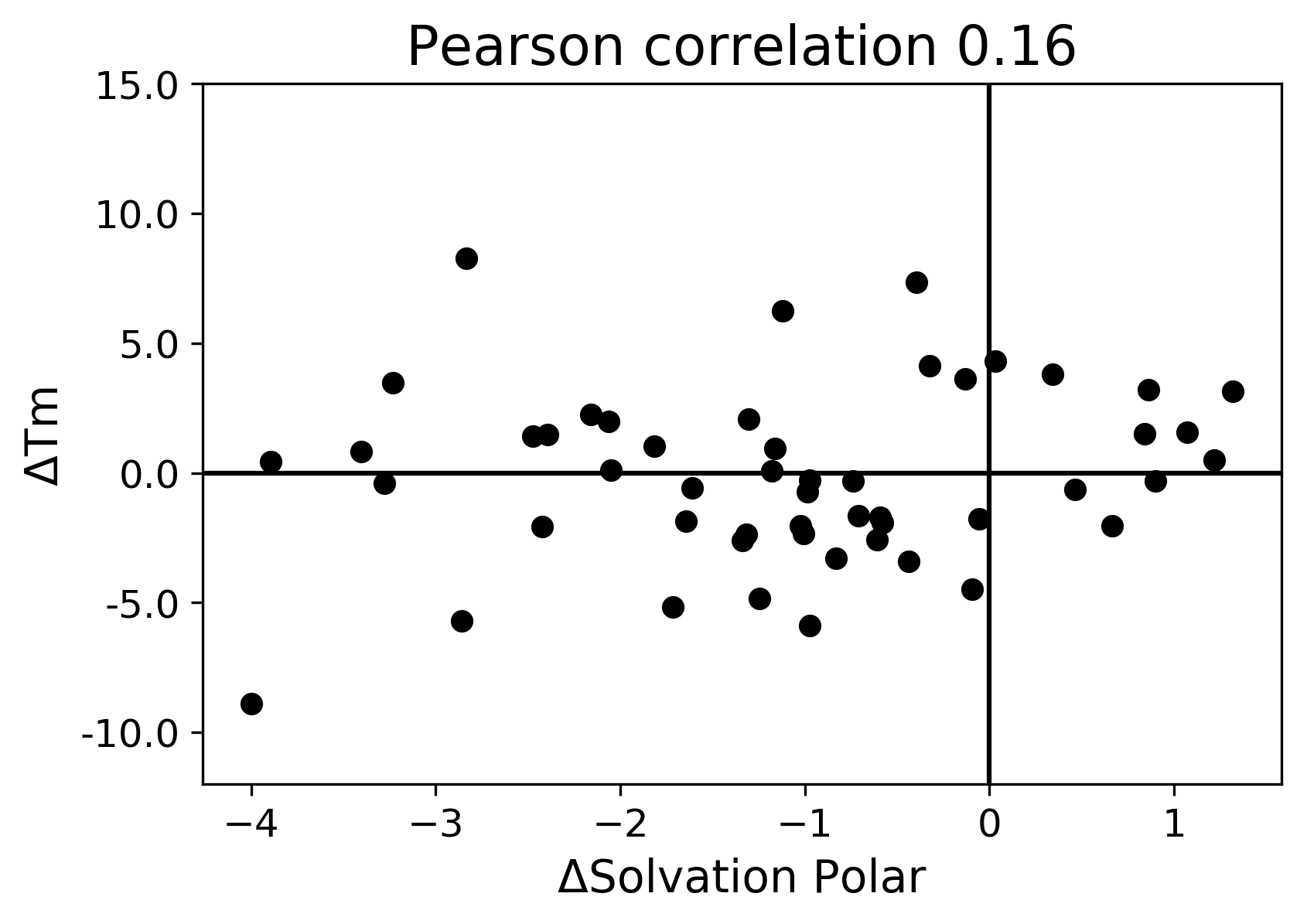

### T221A_F03_dg.pdf

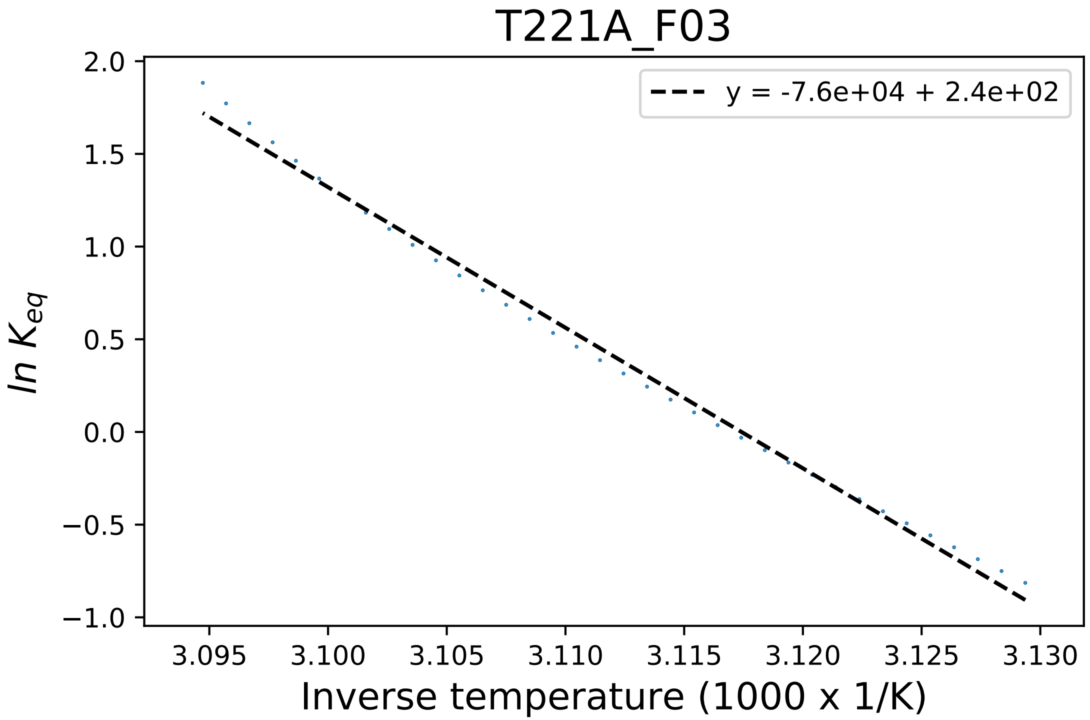

### T221A_F04_fluorescence.pdf

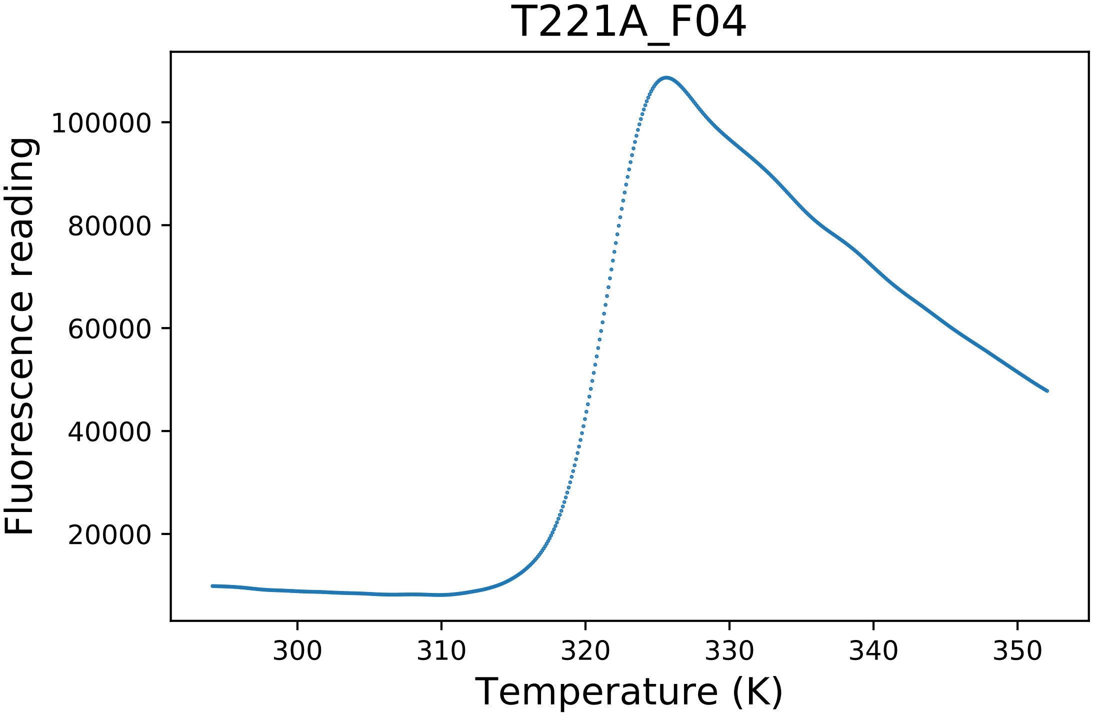

### torsional clash.png

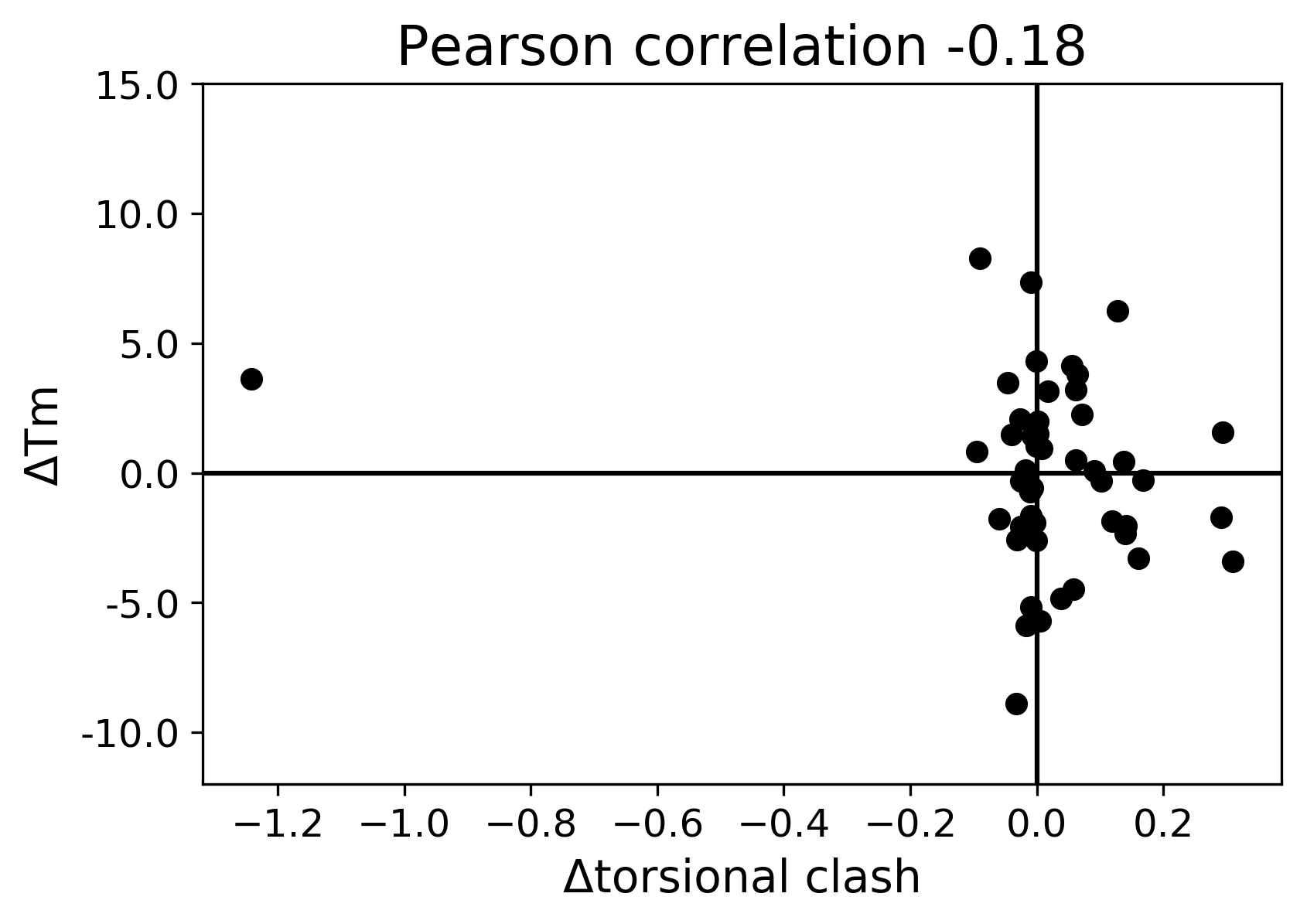

### total energy.png

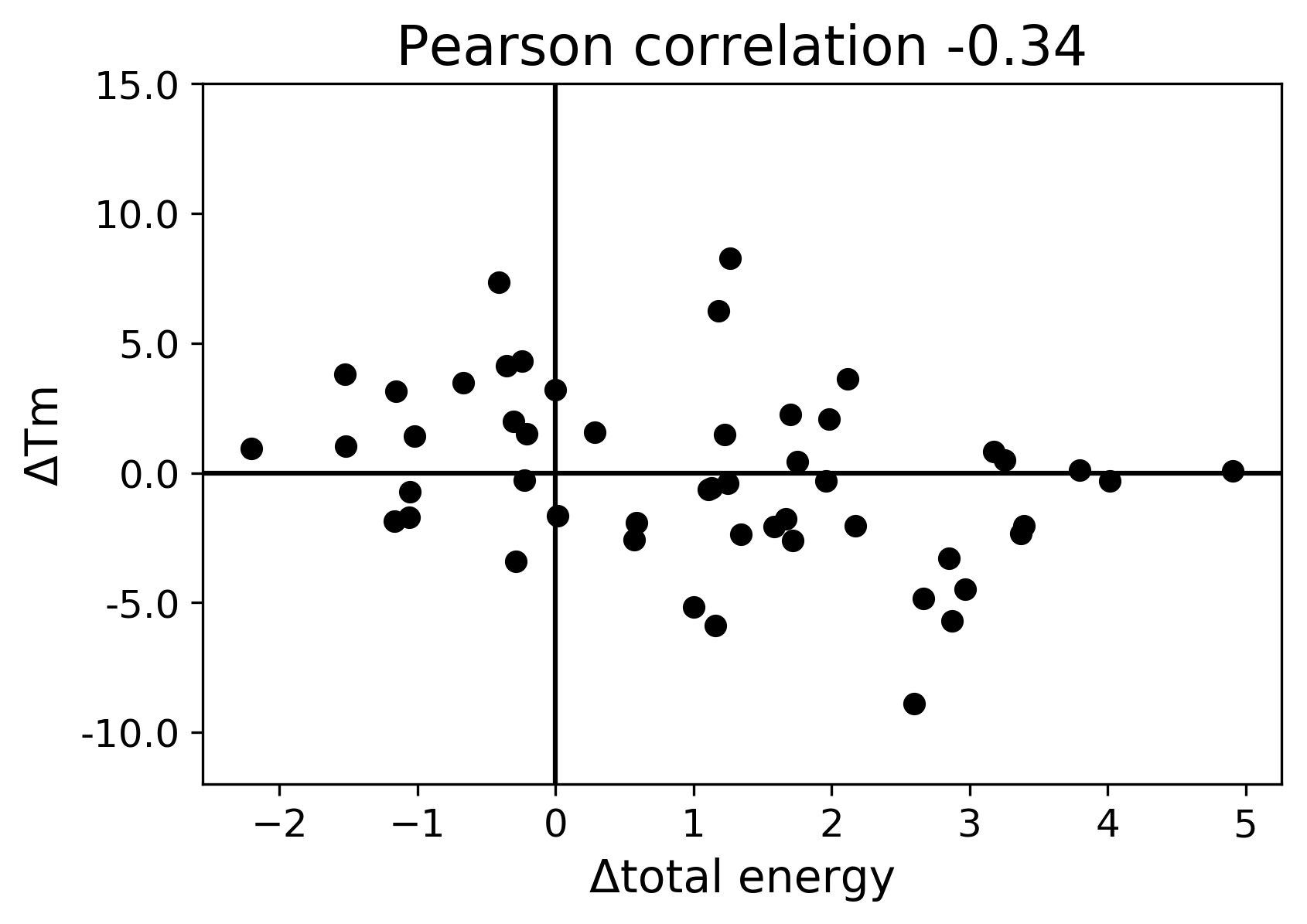

### total.png

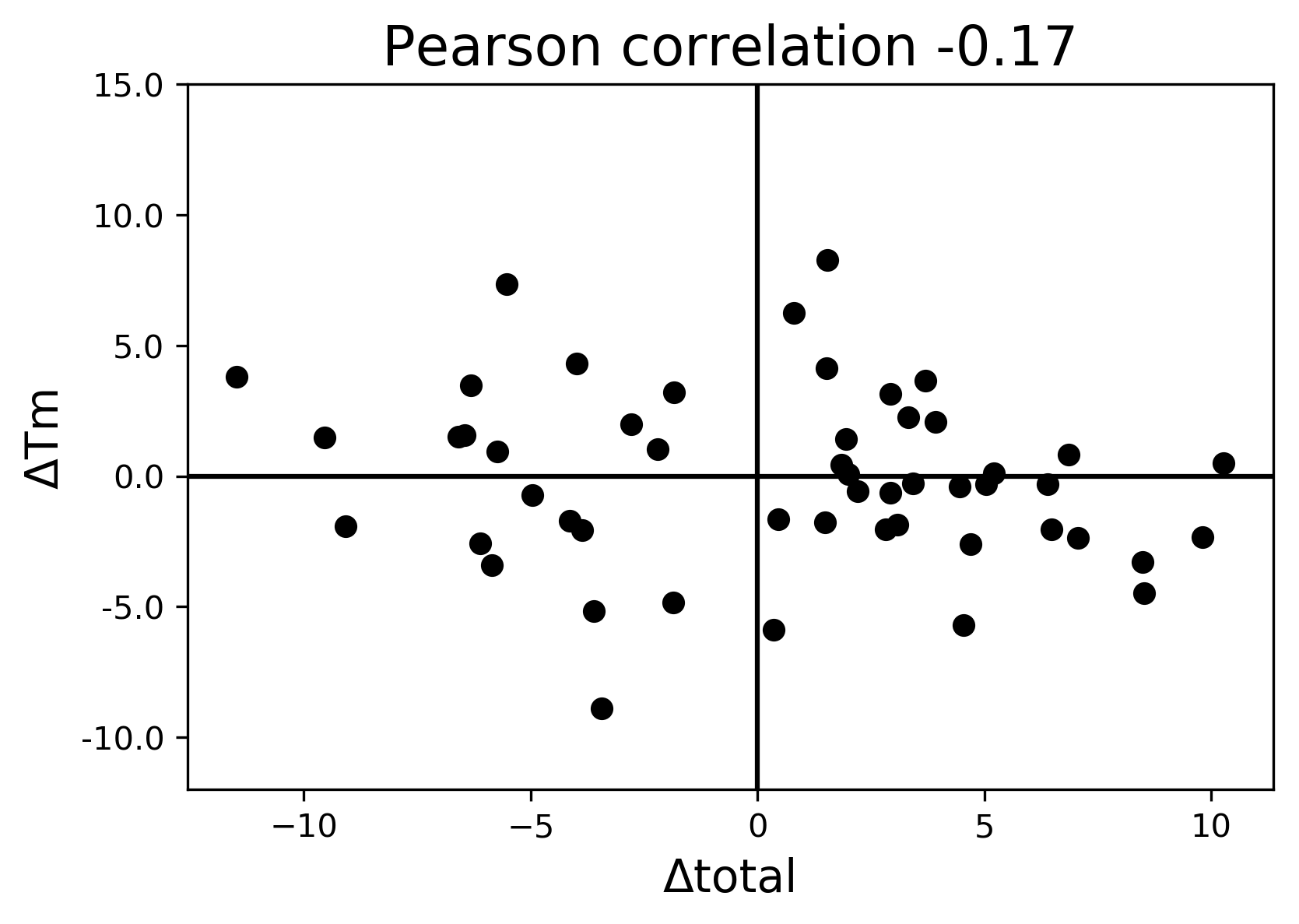

### Van der Waals clashes.png

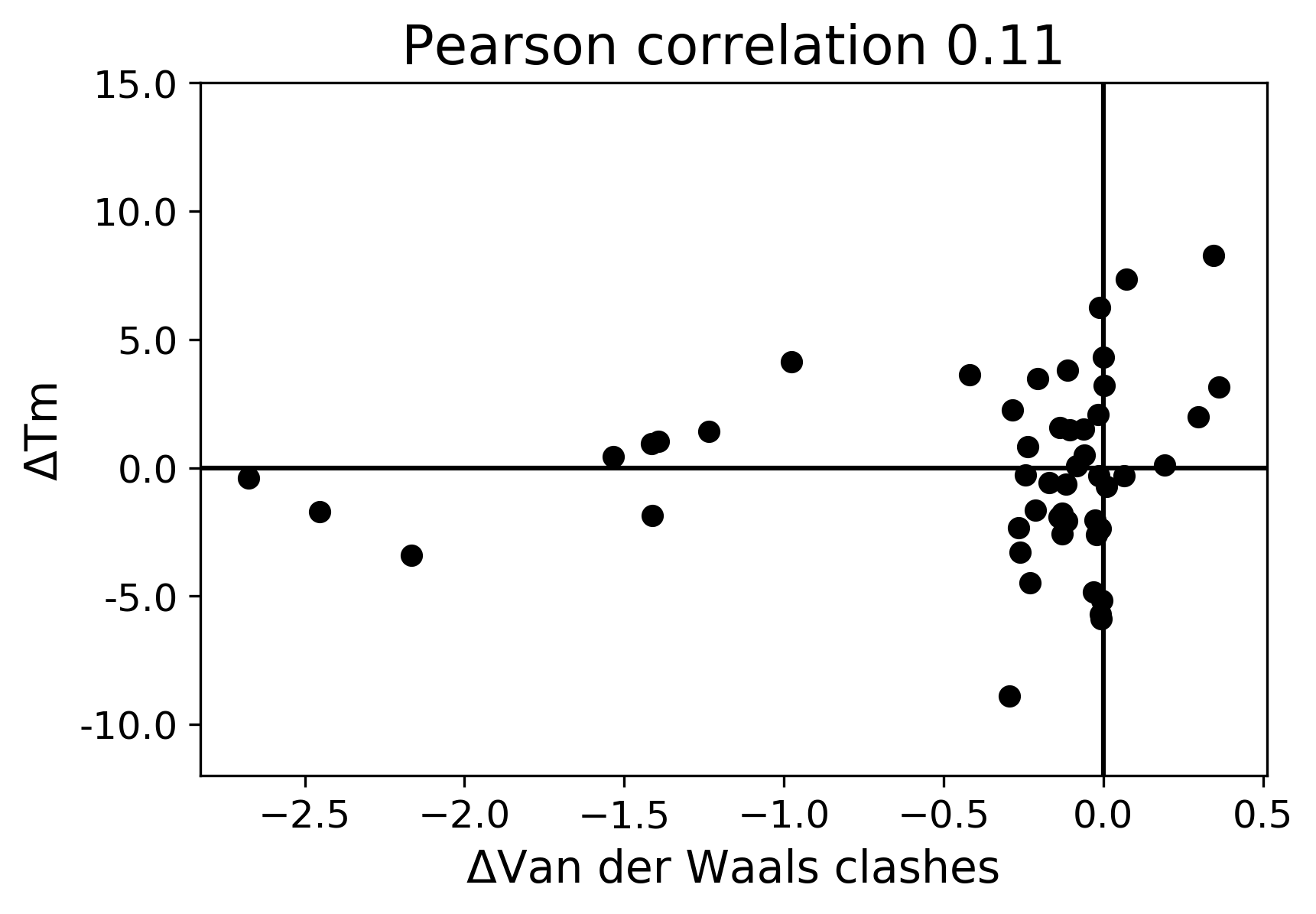

### Van der Waals.png

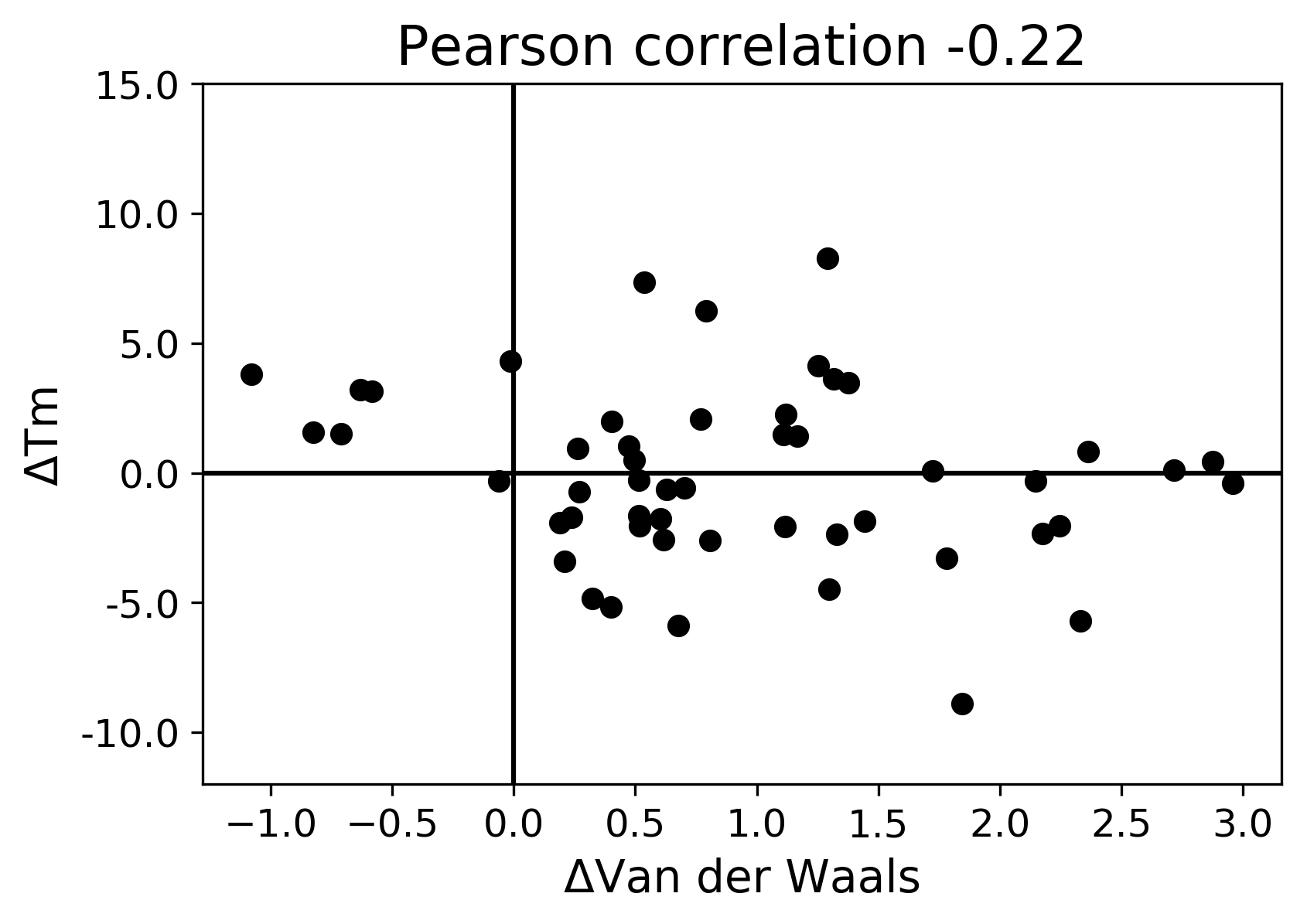

### W123F_G08_dg.pdf

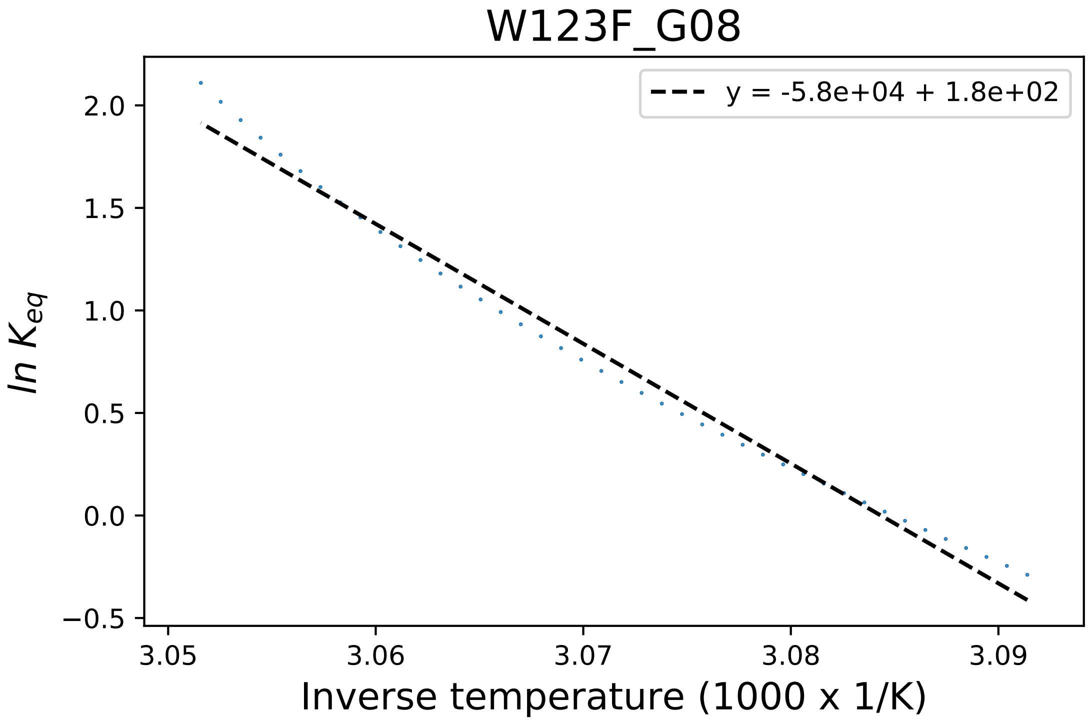

### W123H_F01_dg.pdf

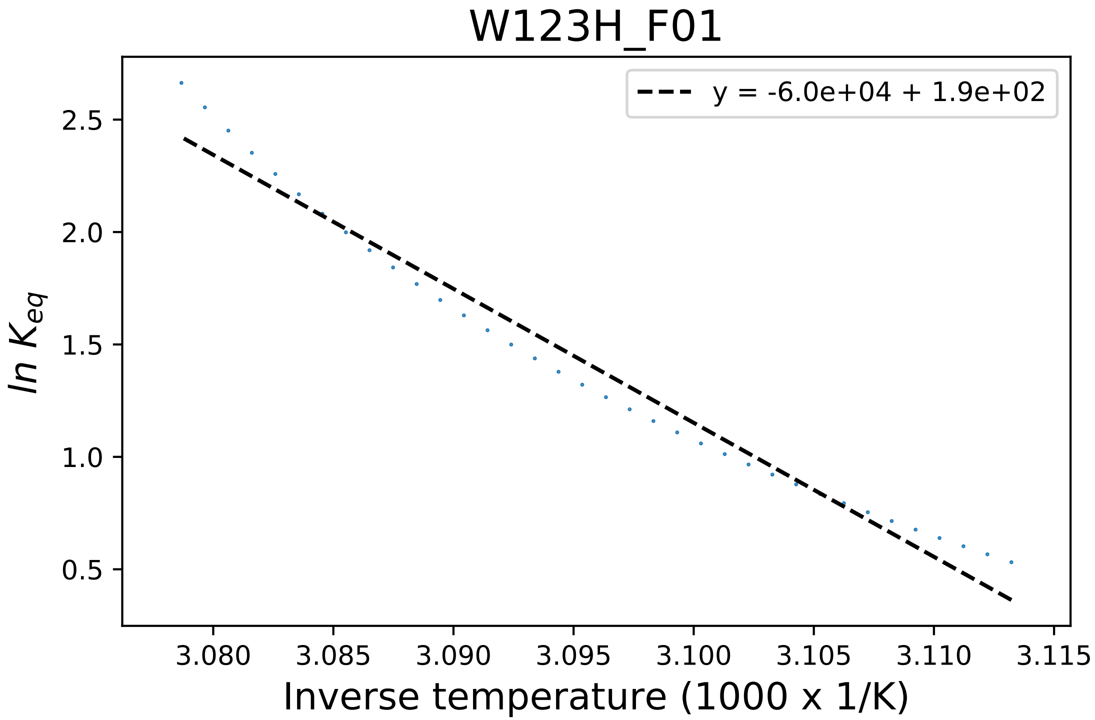

### W123H_F04_fluorescence.pdf

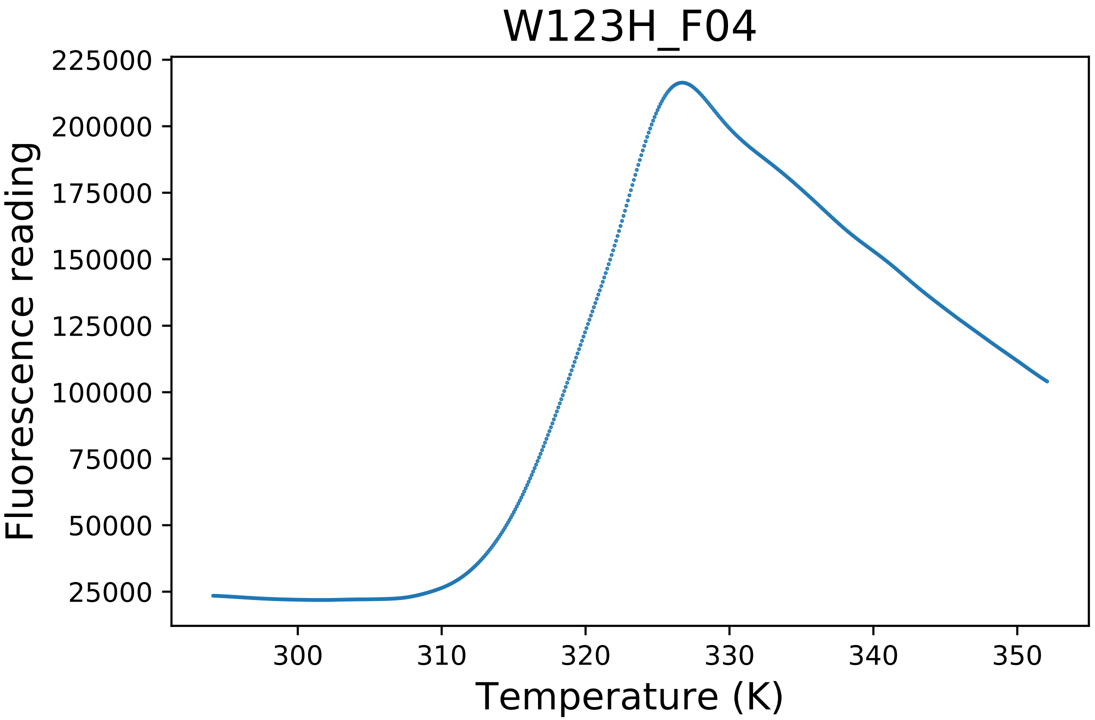

### W328C_A09_fluorescence.pdf

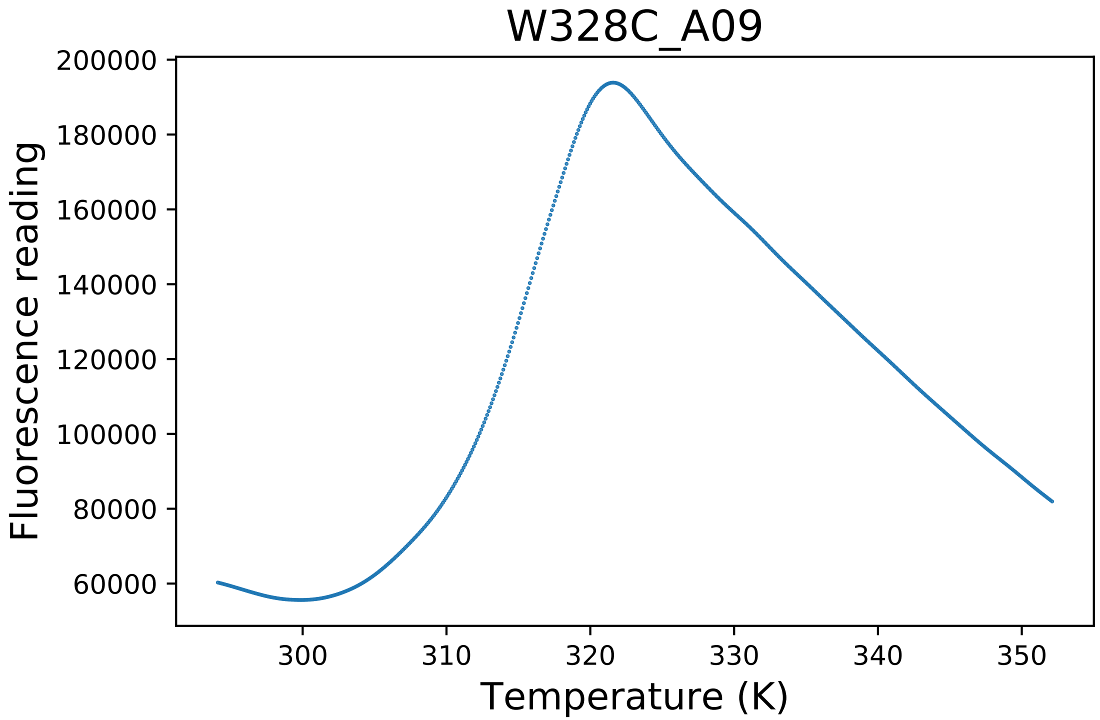

### W328C_A10_dg.pdf

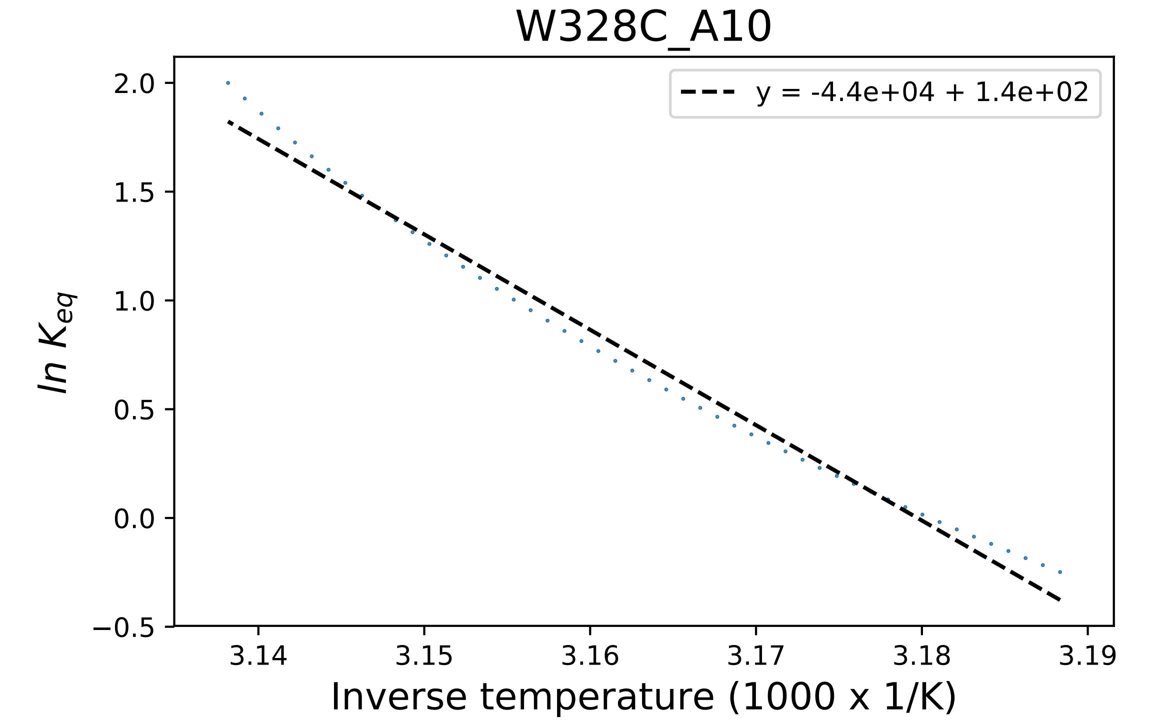

### W328C_A11_derivative.pdf

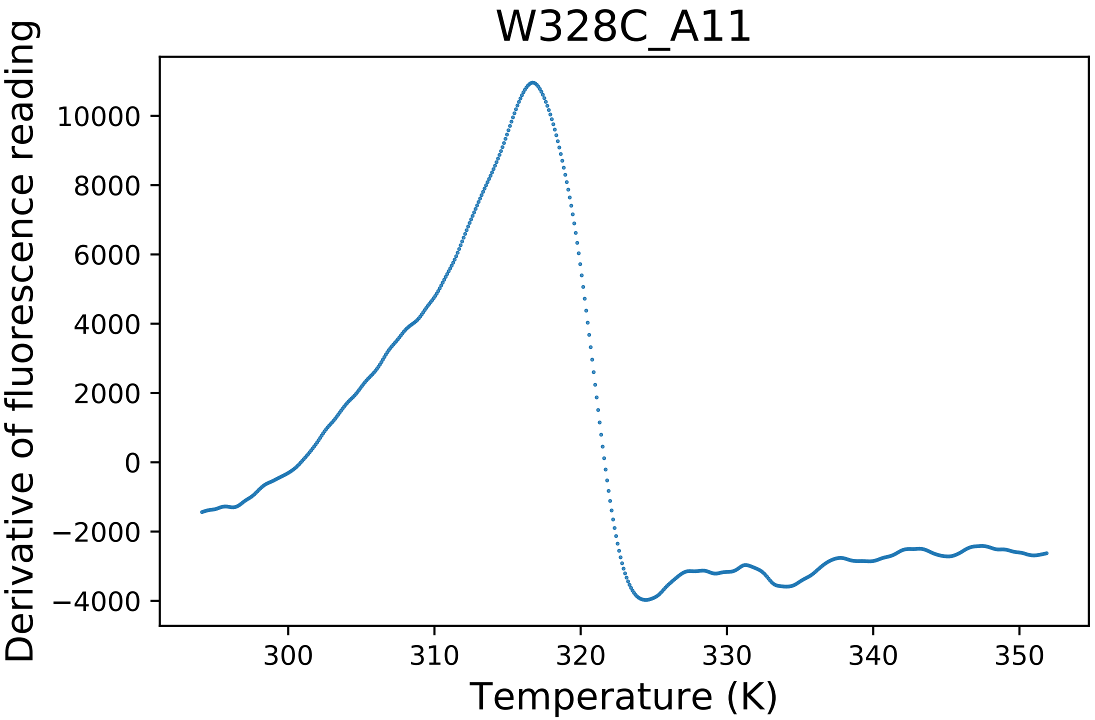

### W328H_A05_dg.pdf

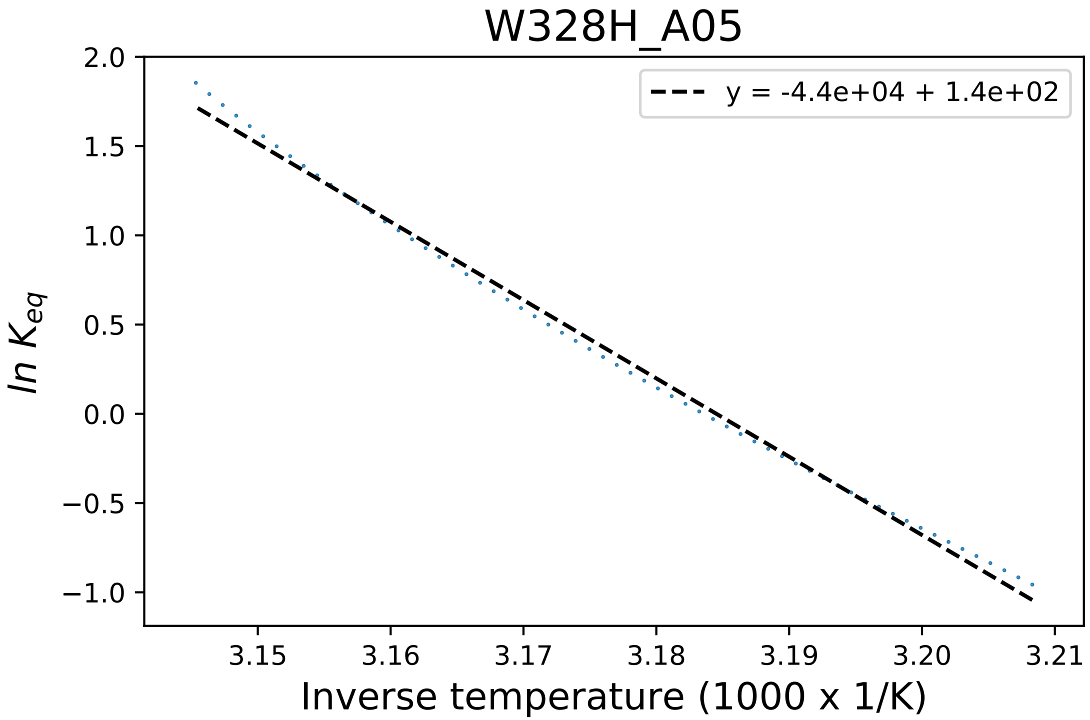

### W328H_A06_derivative.pdf

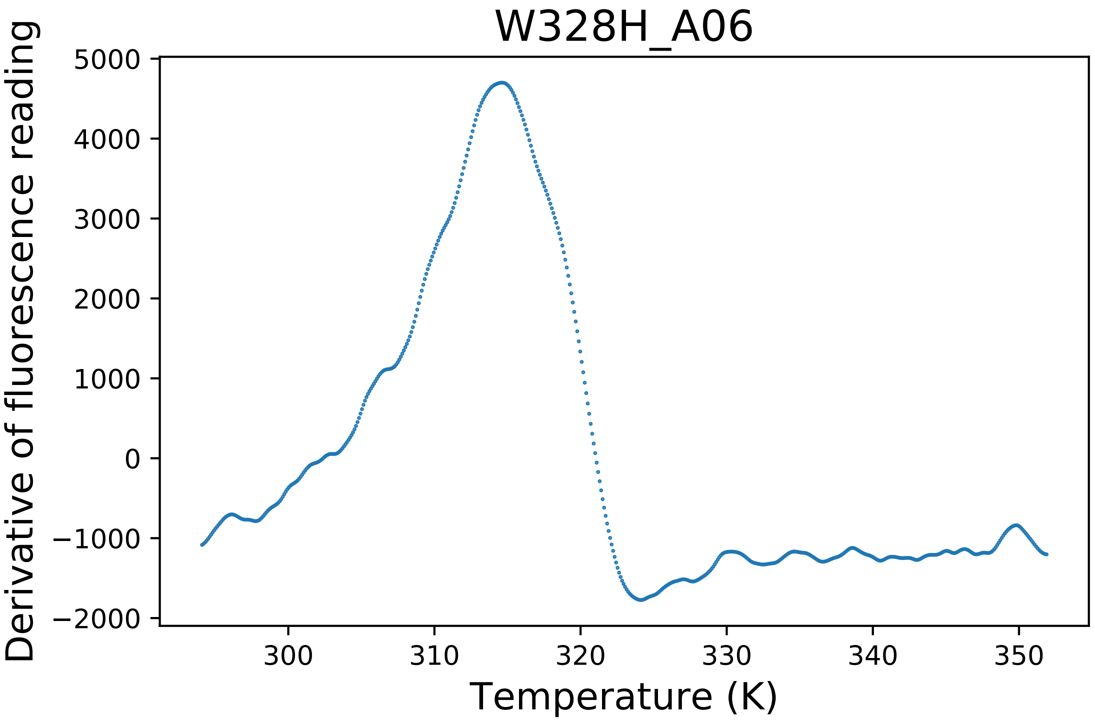

### W328H_A08_fluorescence.pdf

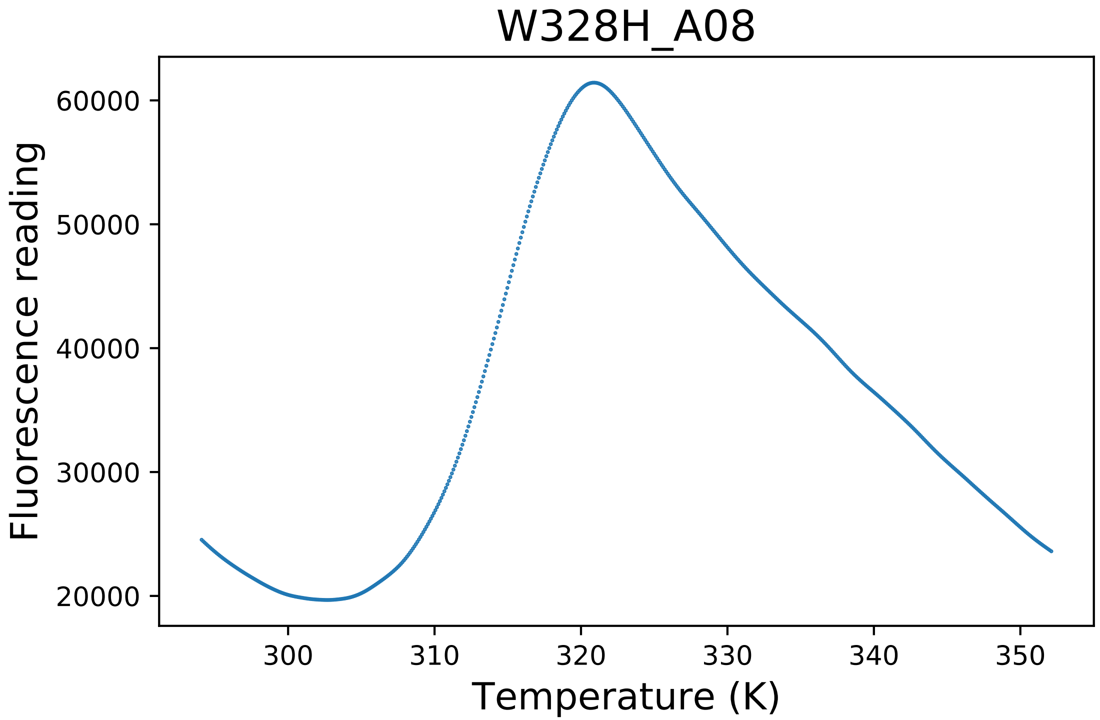

### W328R_D07_fluorescence.pdf

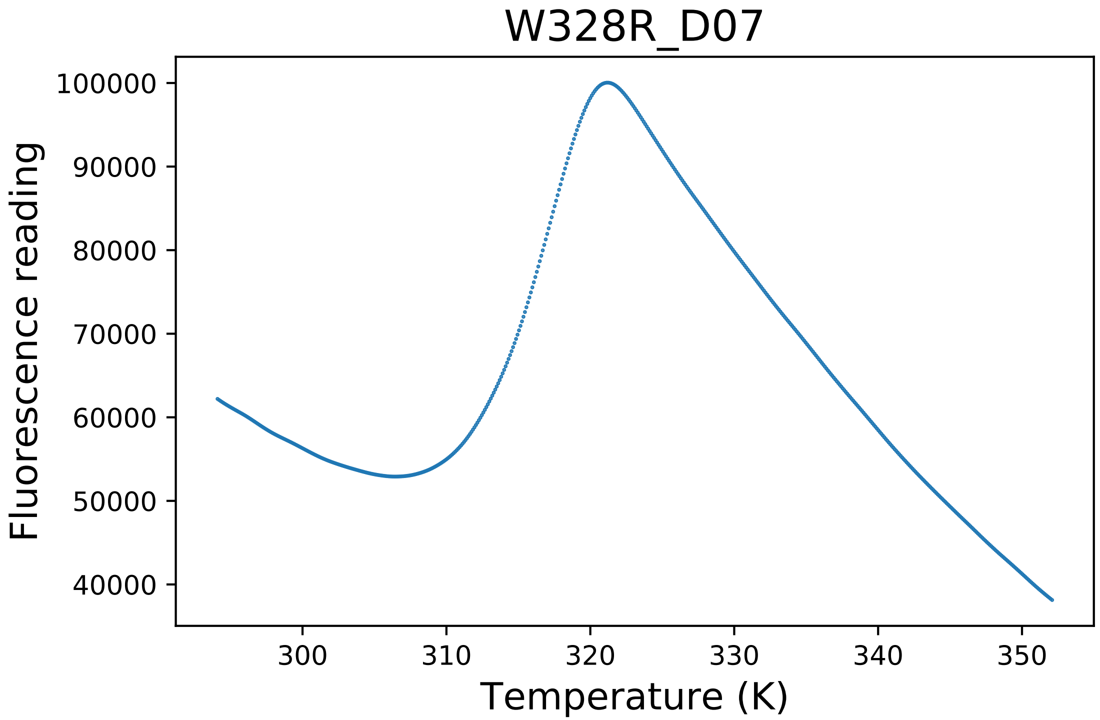

### W402C_E05_derivative.pdf

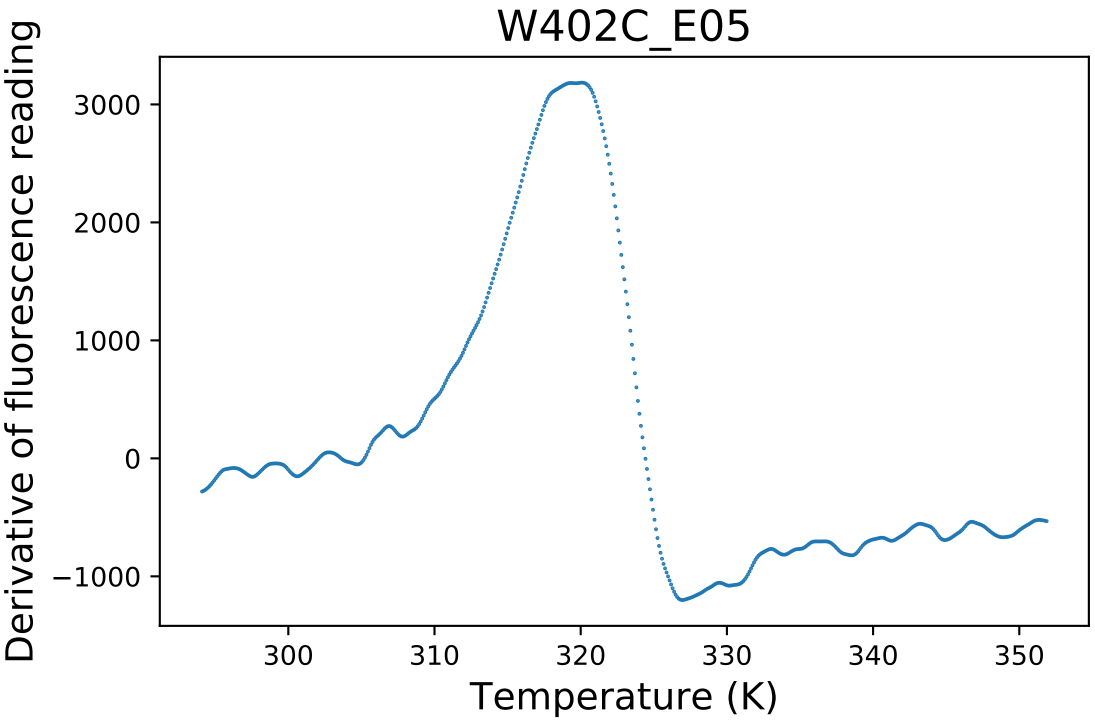

### W402C_E06_dg.pdf

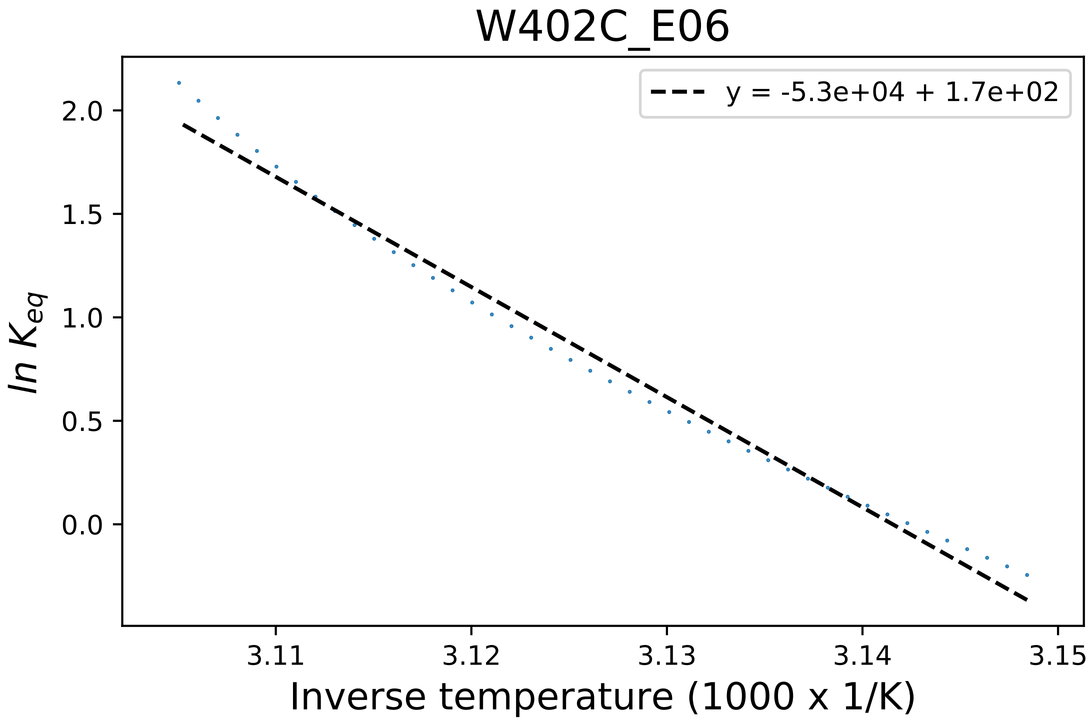

### W402R_A06_derivative.pdf

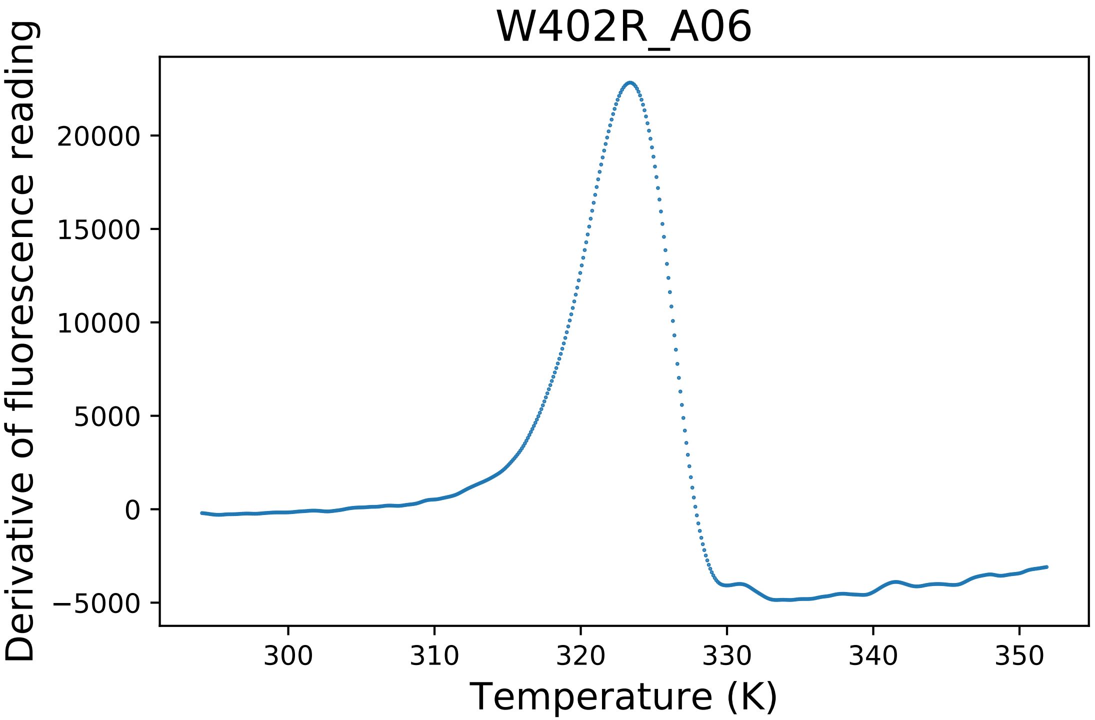

### WT1_A03_dg.pdf

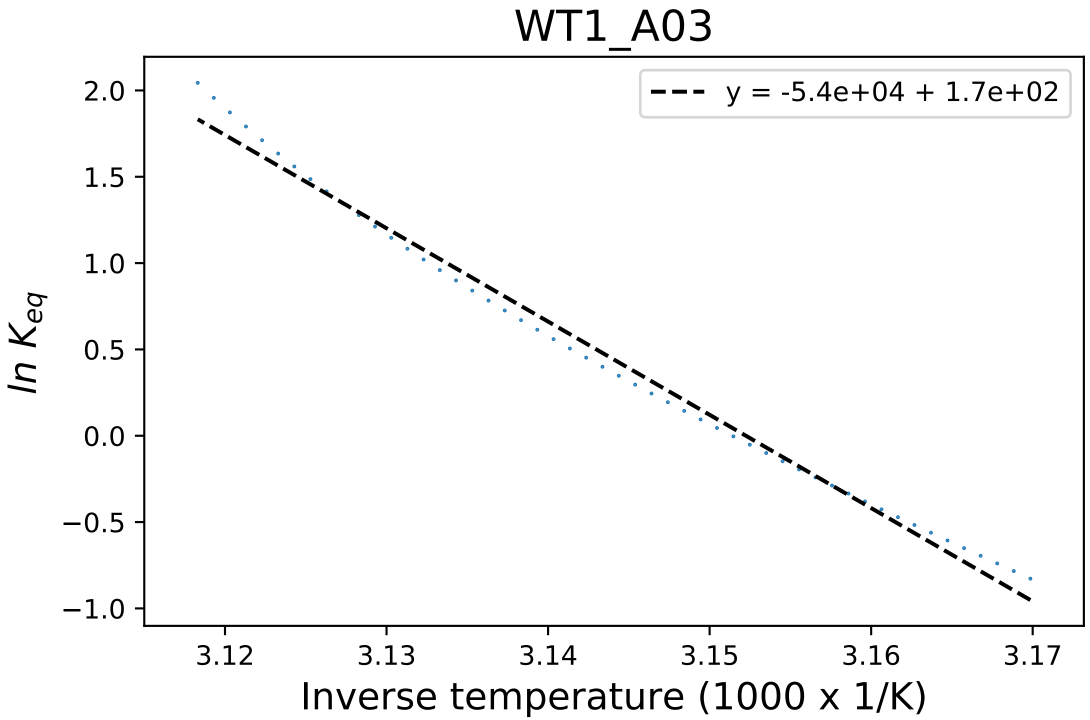

### WT2_H09_dg.pdf

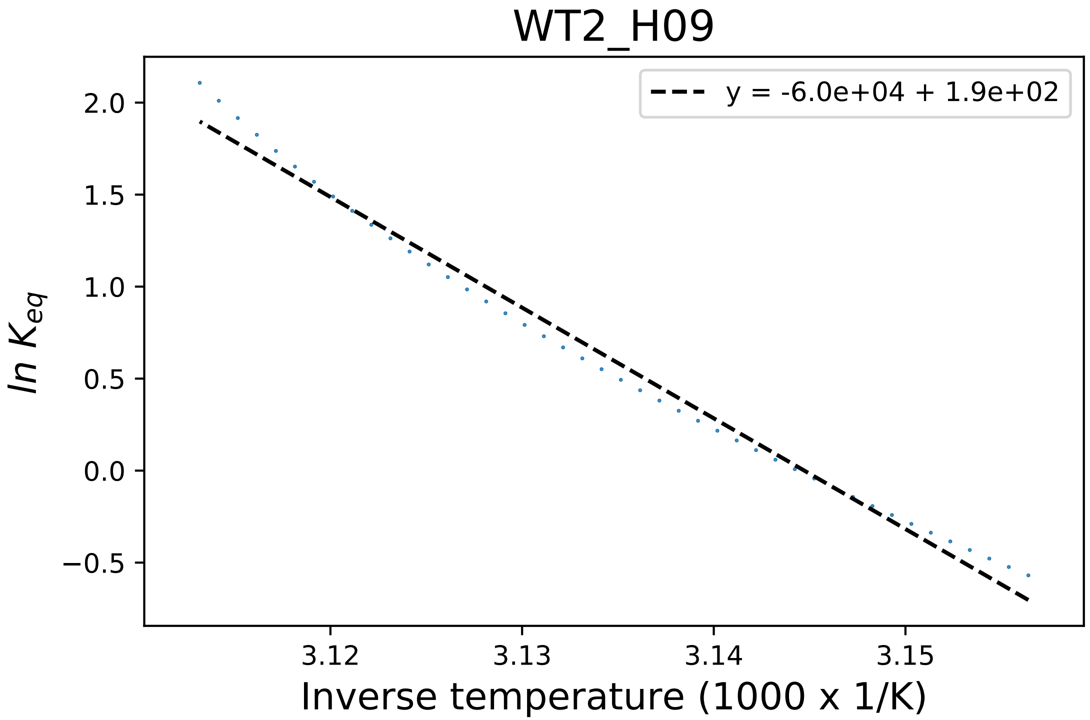

### WT4_A01_dg.pdf

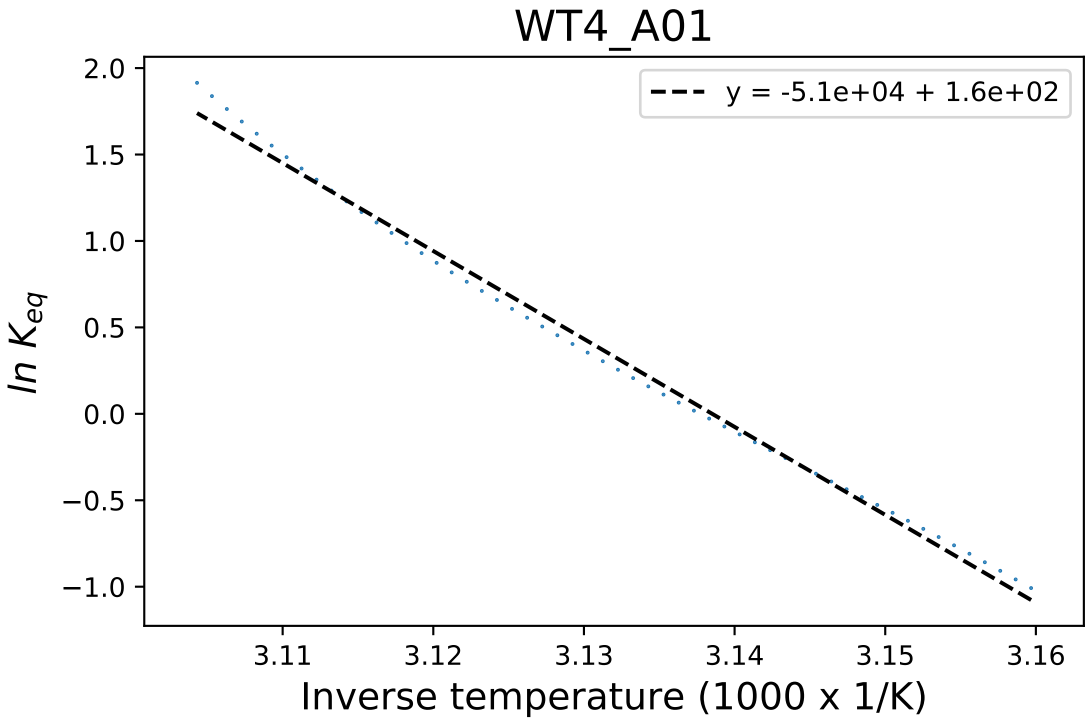

### WT4_A04_fluorescence.pdf

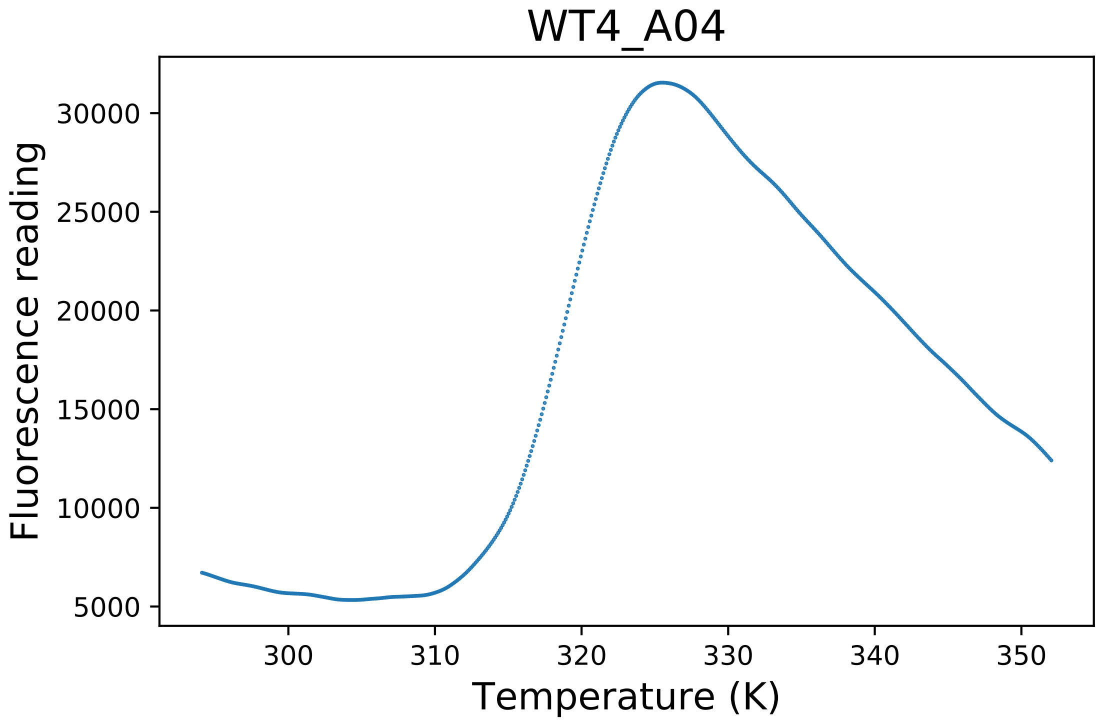

### WT6_D06_derivative.pdf

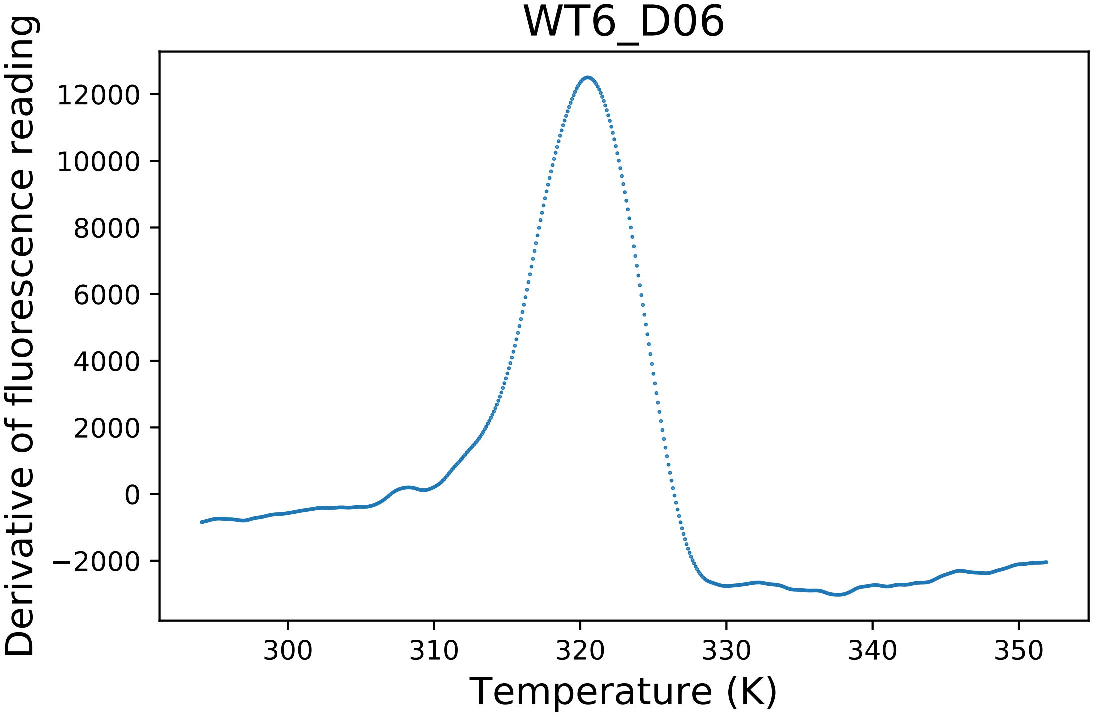

### WT6_D08_fluorescence.pdf

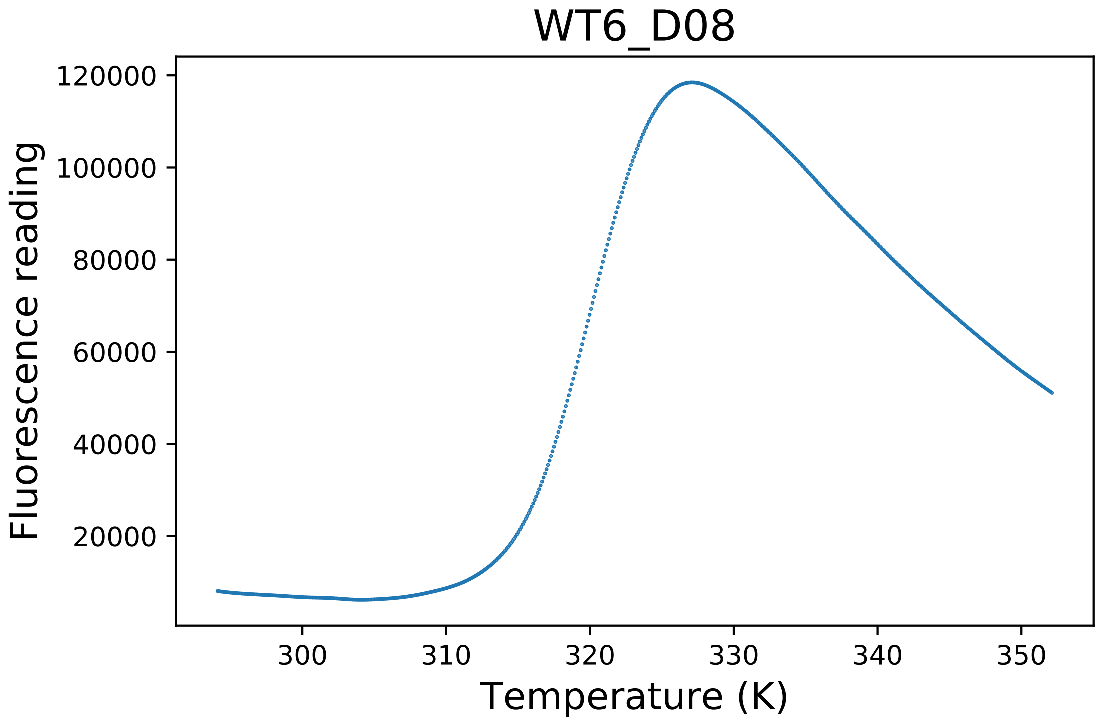

### Y21A_B04_derivative.pdf

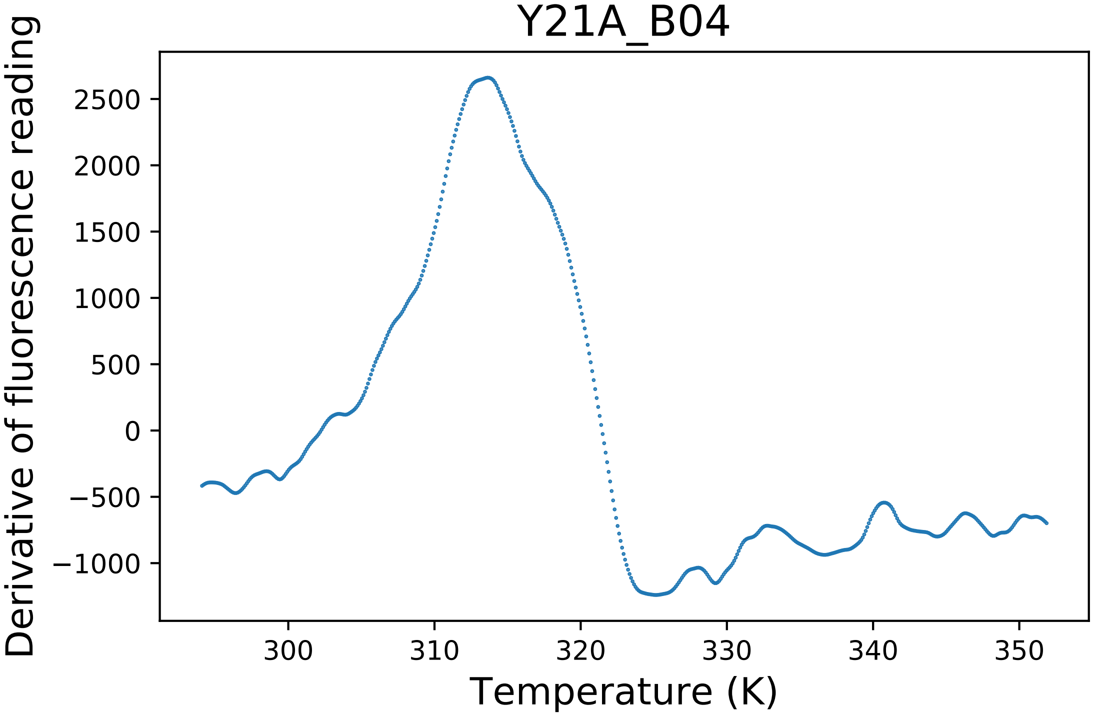

### Y297F_B01_fluorescence.pdf

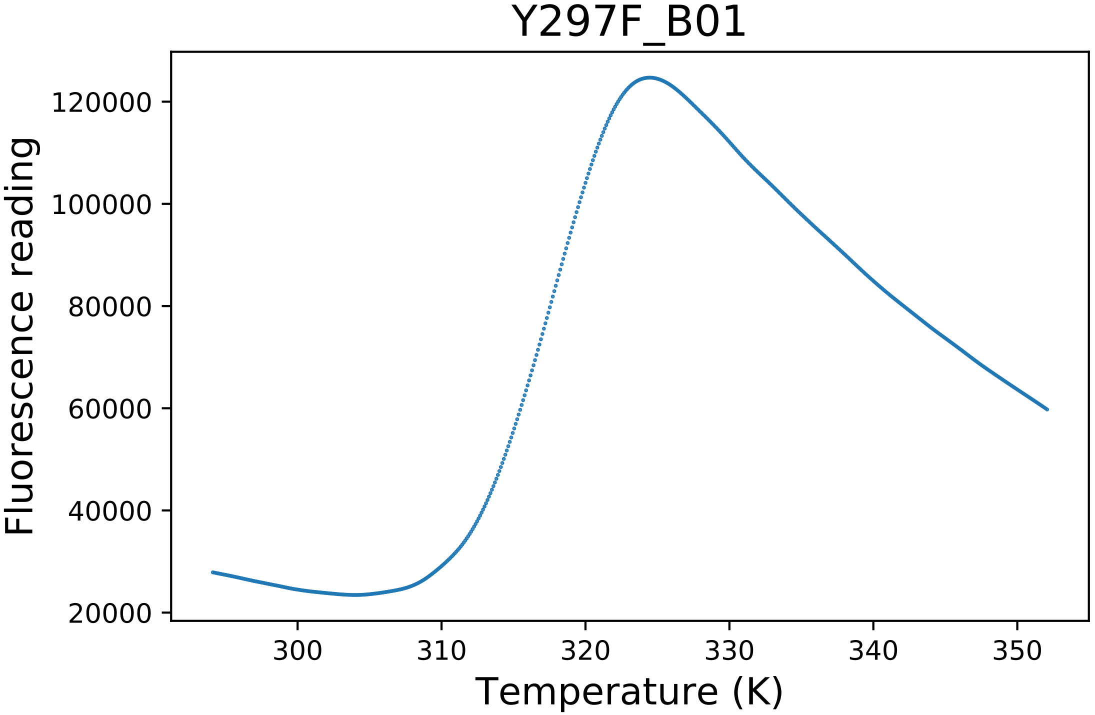

### Y297F_B04_derivative.pdf

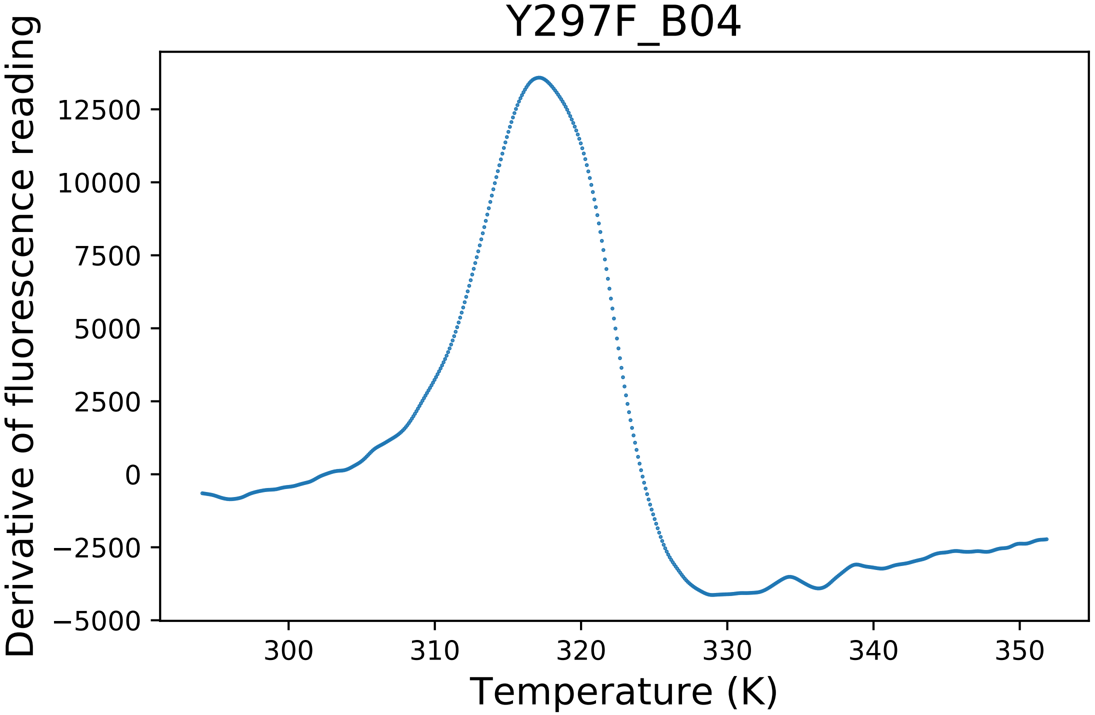

### Y297F_B04_dg.pdf

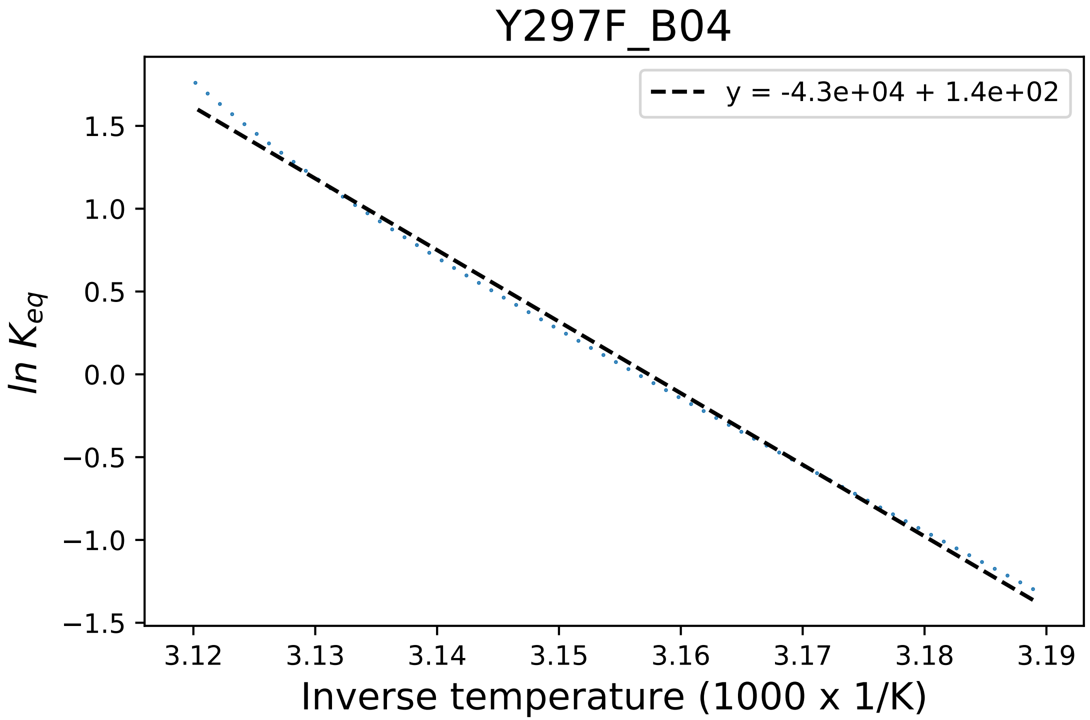

### yhh_planarity.png

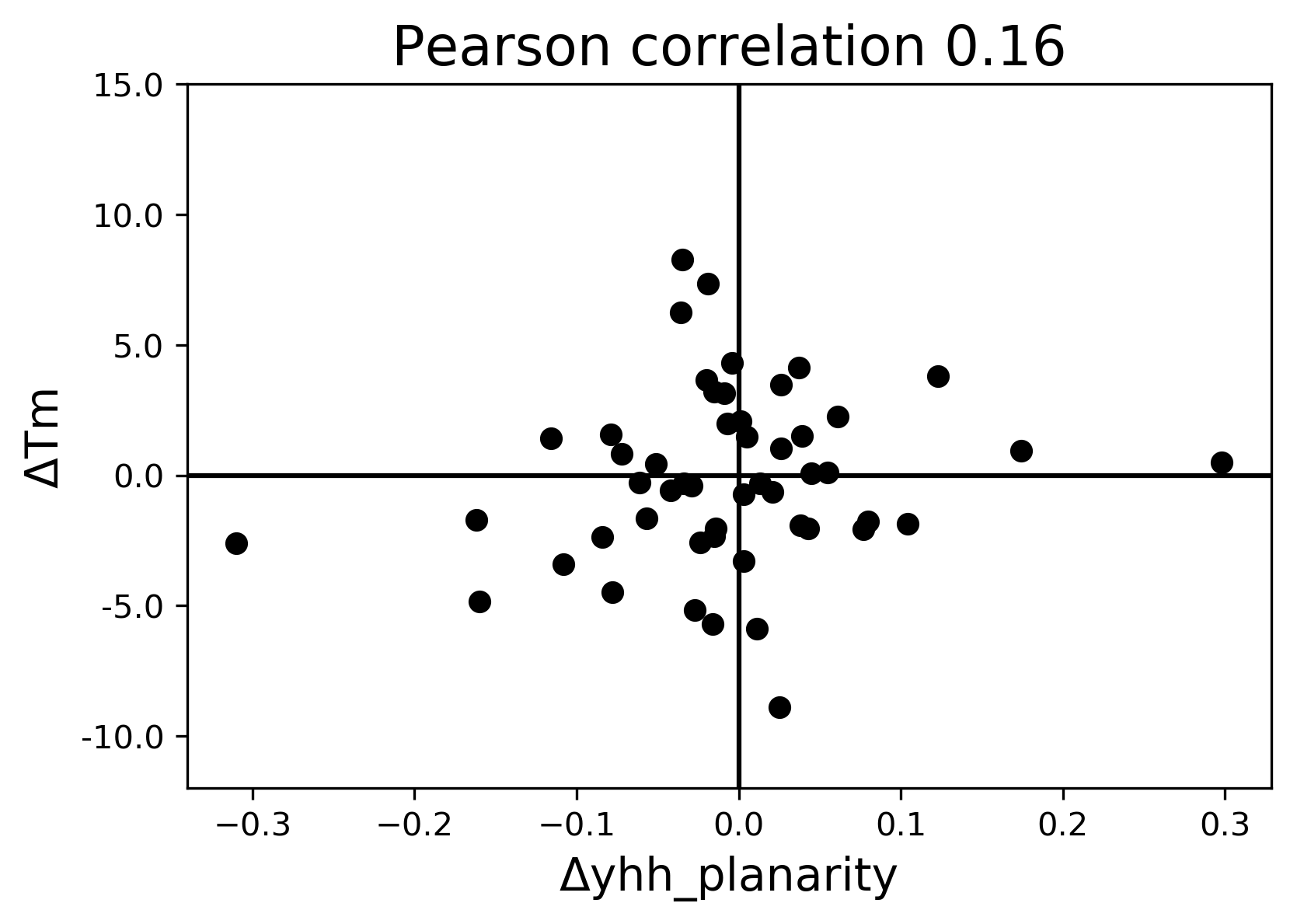
