## Supplementary figures and images for "Evaluating molecular modeling tools for thermal stability using an independently generated dataset"

### C170Q_C11_fluorescence.pdf

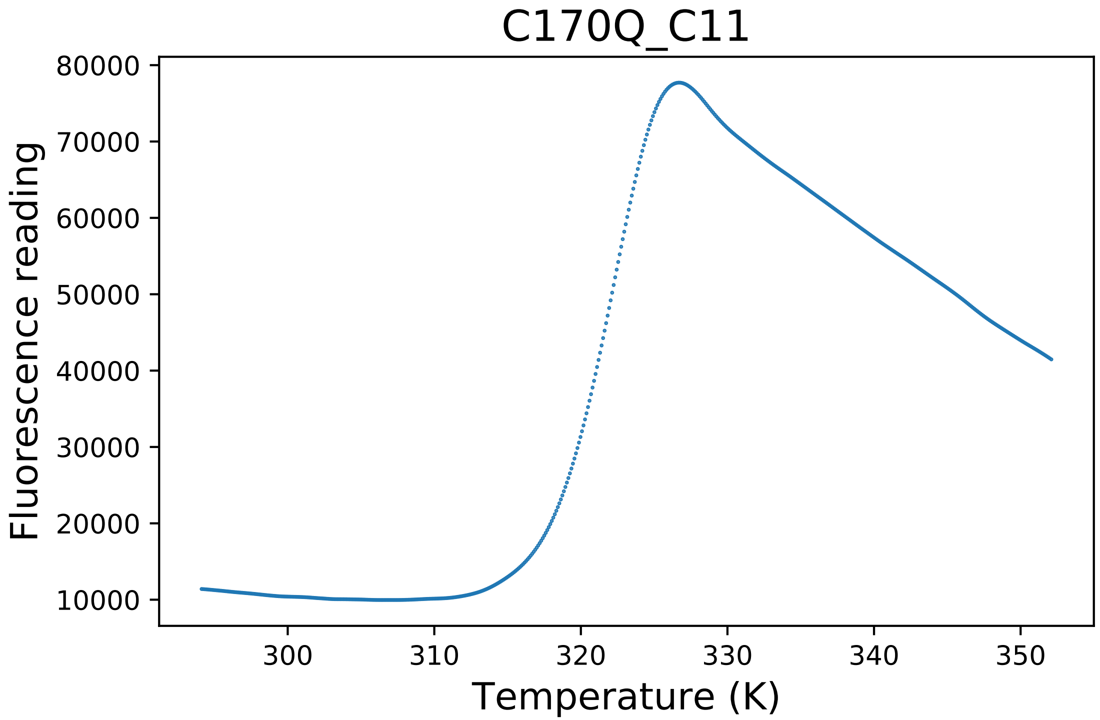

### E157D_A06_dg.pdf

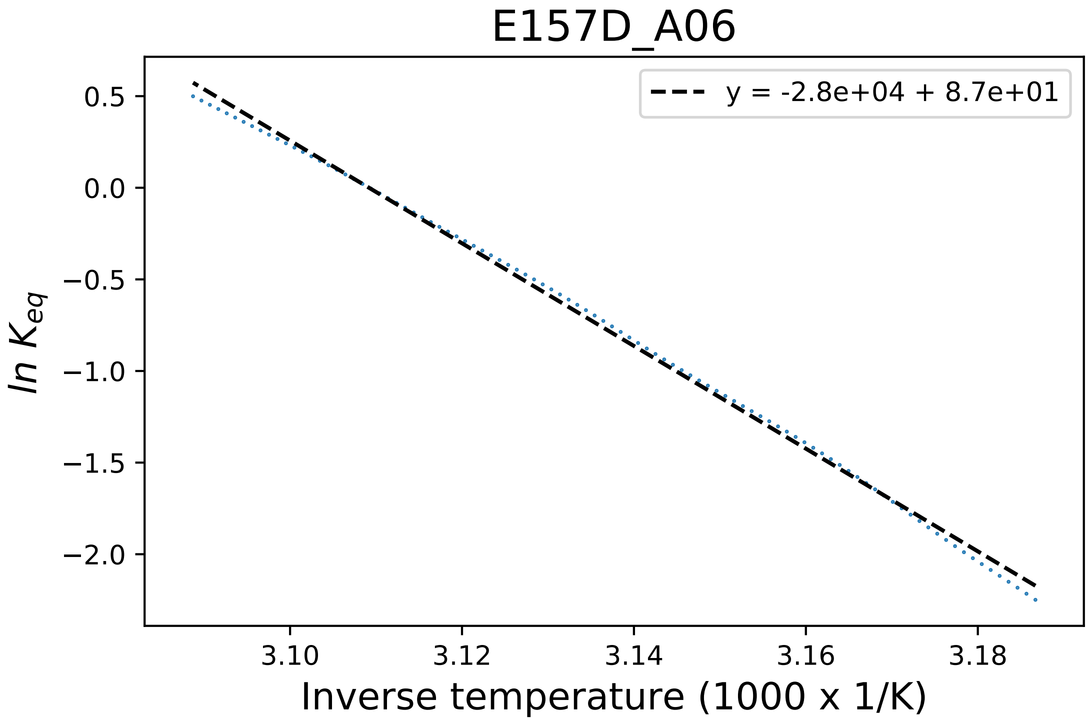

### E167A_A09_fluorescence.pdf

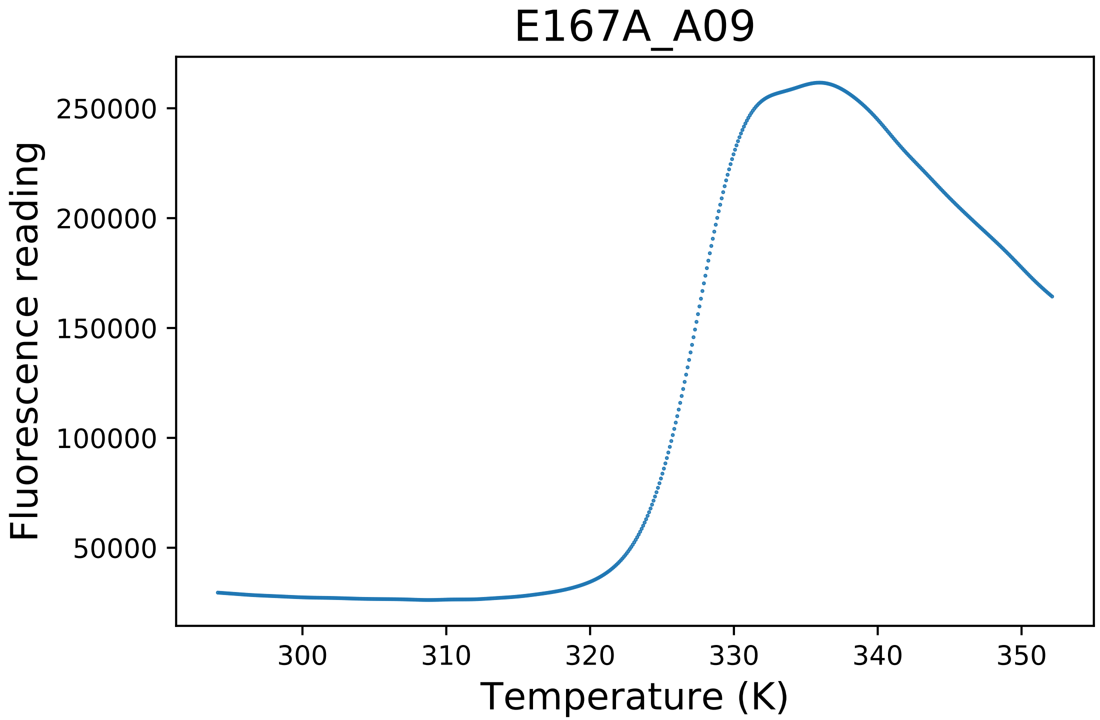

### E225A_A01_dg.pdf

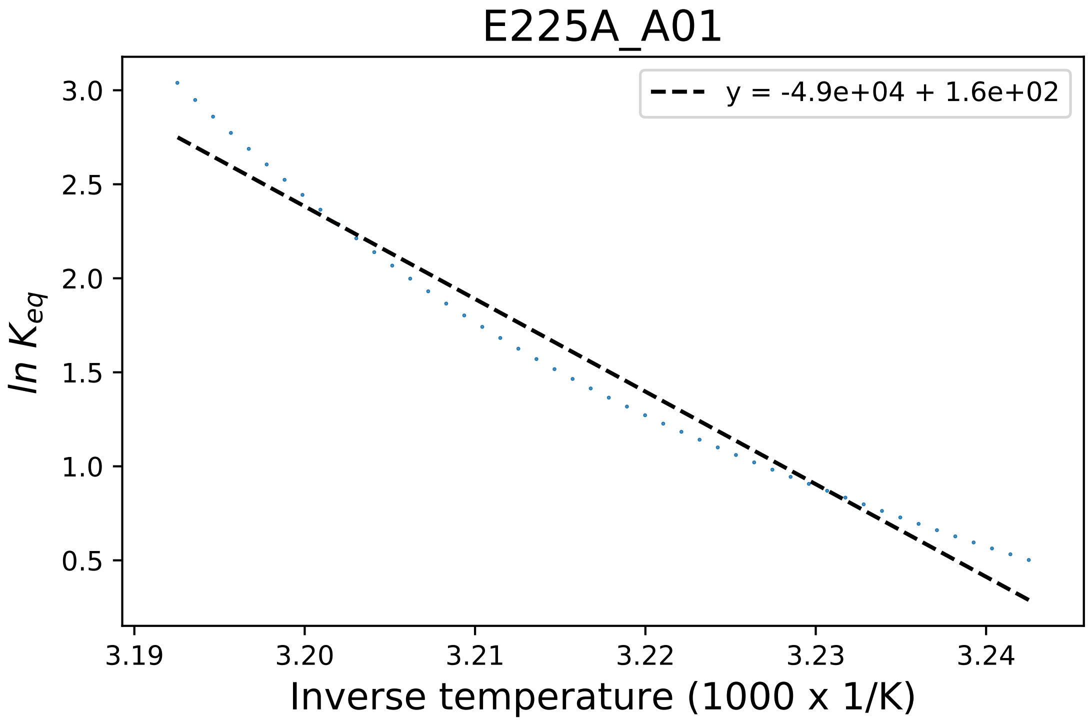

### E225H_C11_dg.pdf

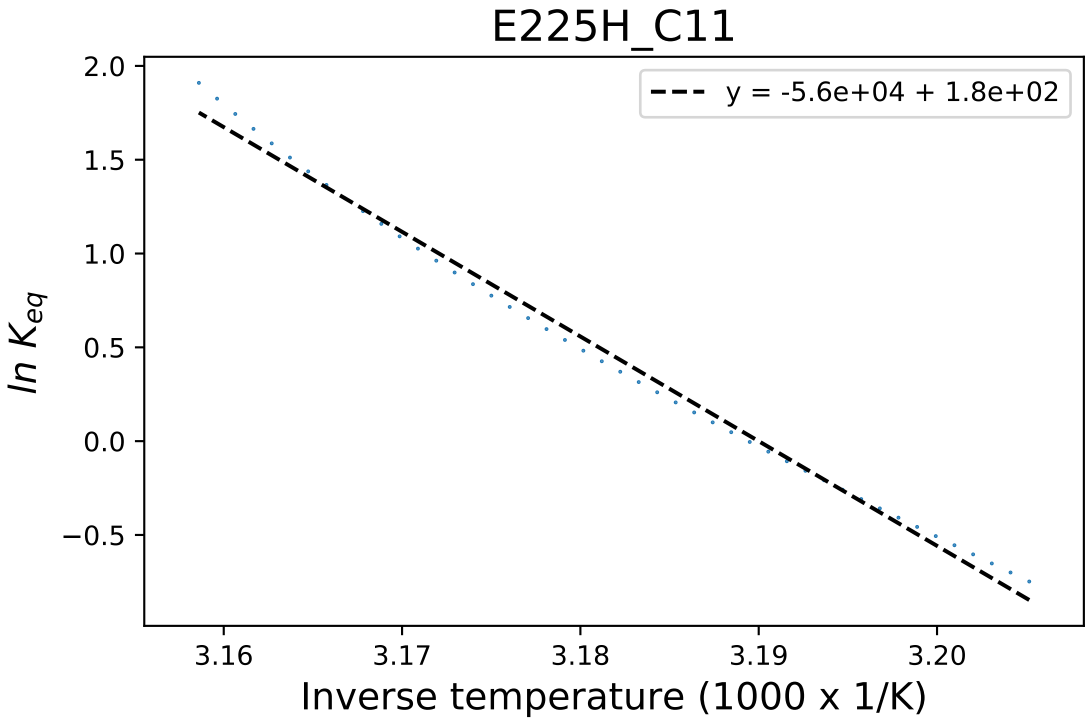

### E426S_E06_dg.pdf

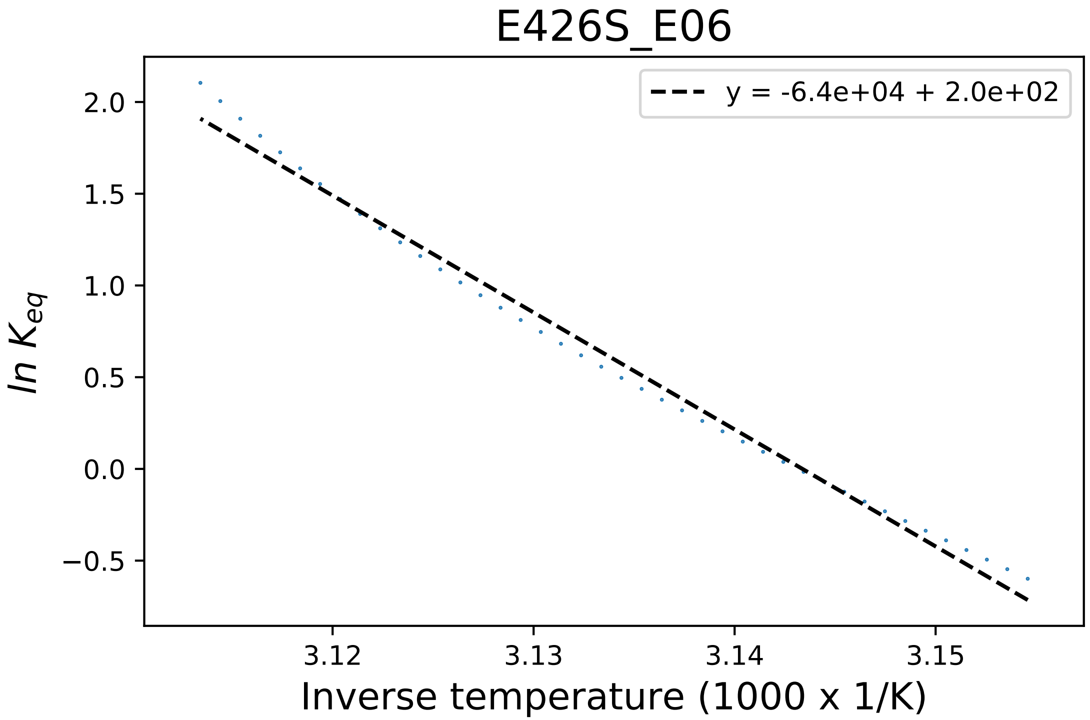

### F418A_H02_fluorescence.pdf

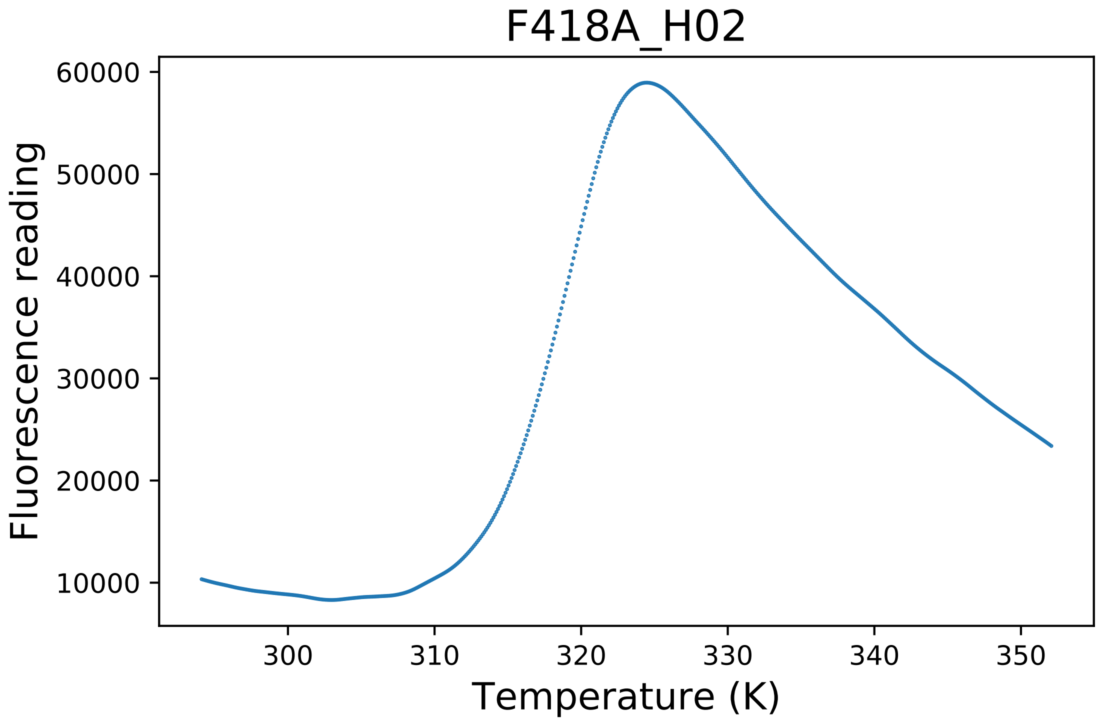

### I247N_B02_derivative.pdf

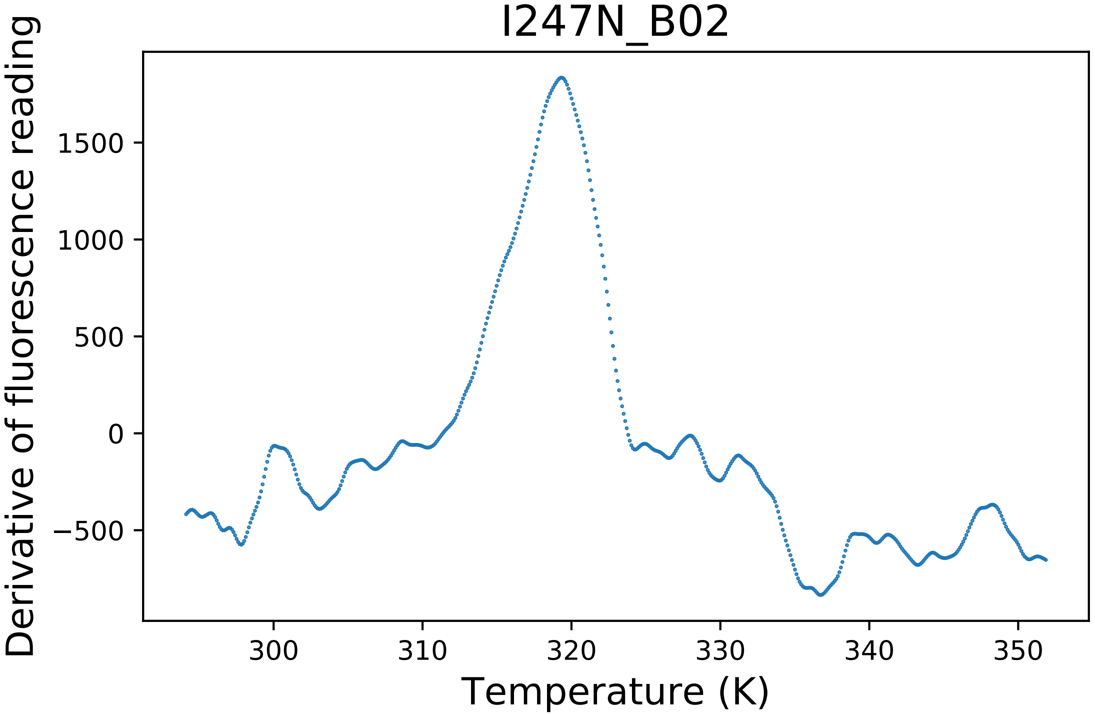

### L174A_E03_derivative.pdf

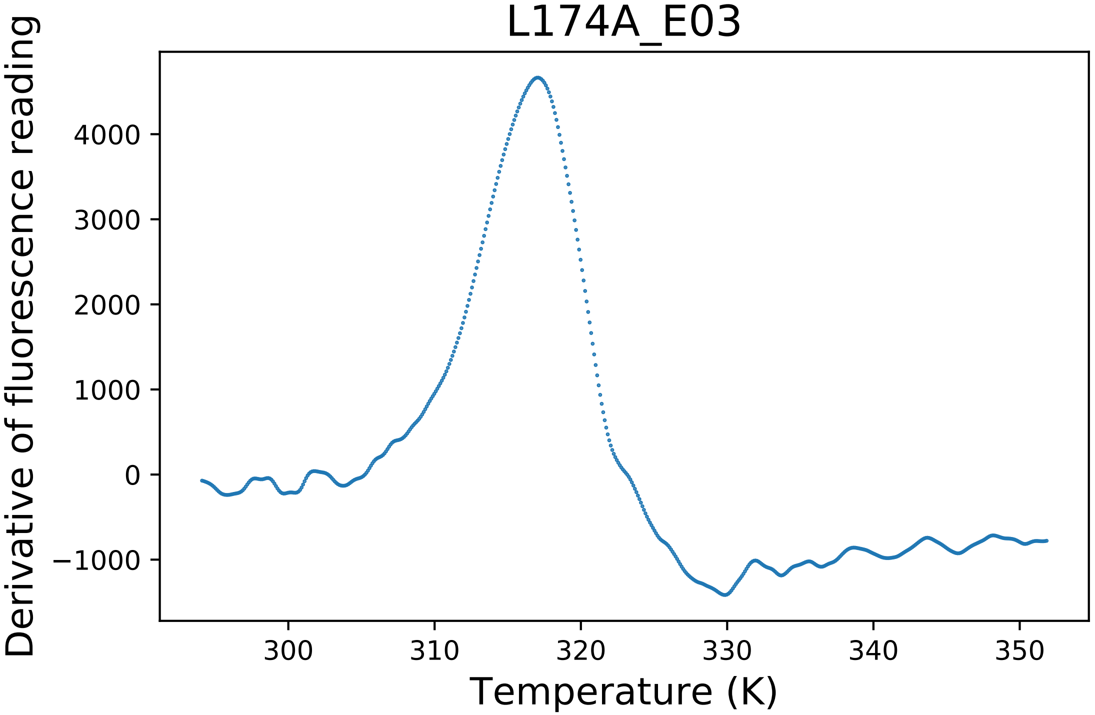

### L174A_E04_dg.pdf

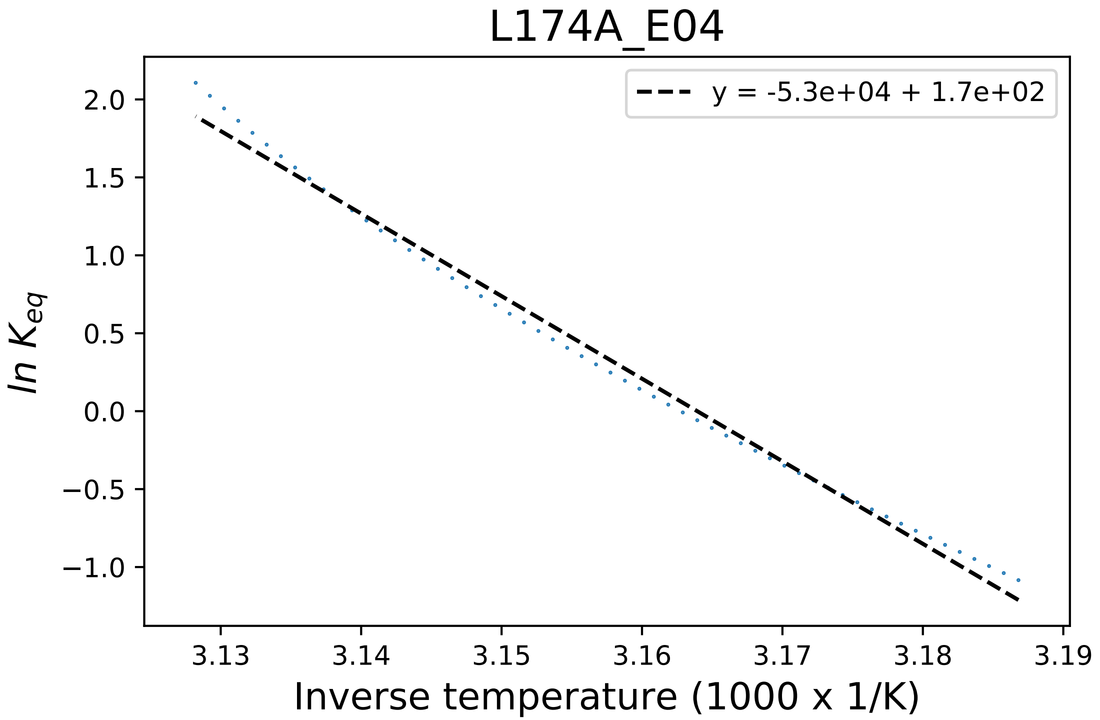

### N223A_F05_dg.pdf

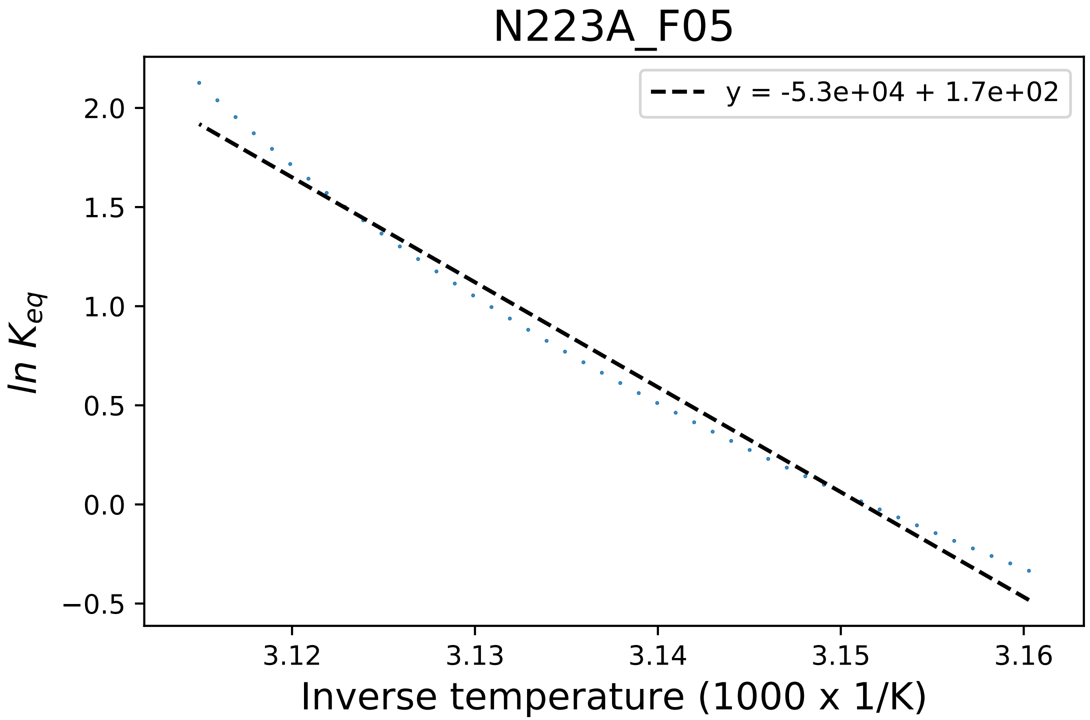

### N223A_F06_derivative.pdf

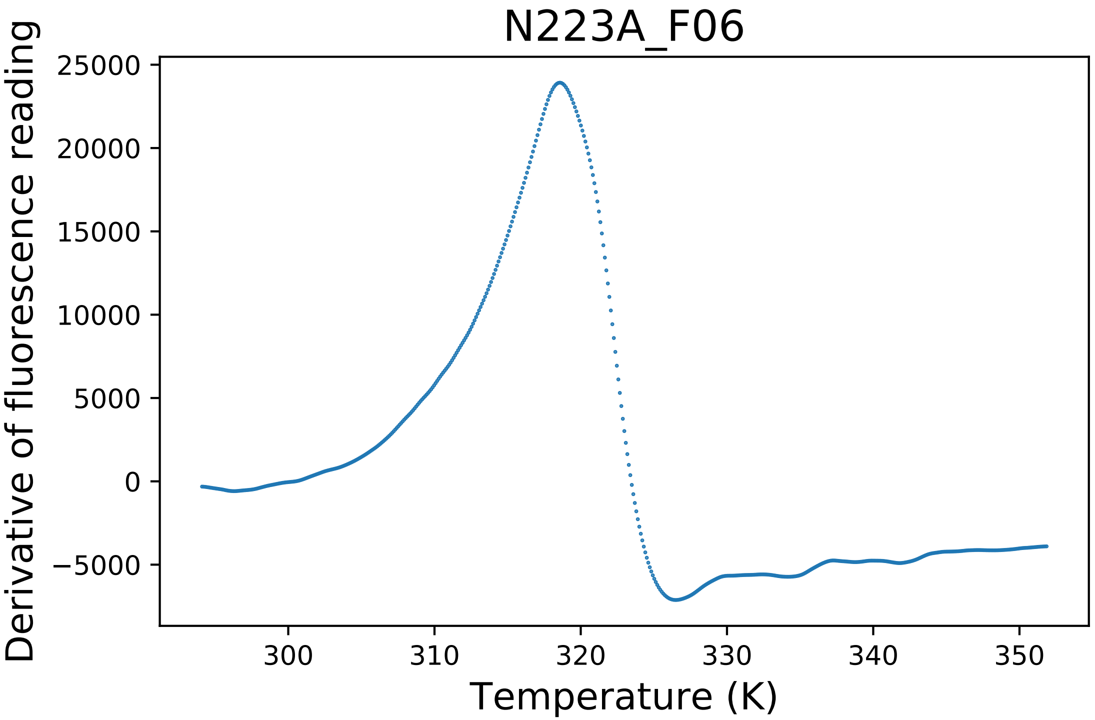

### N223R_G03_fluorescence.pdf

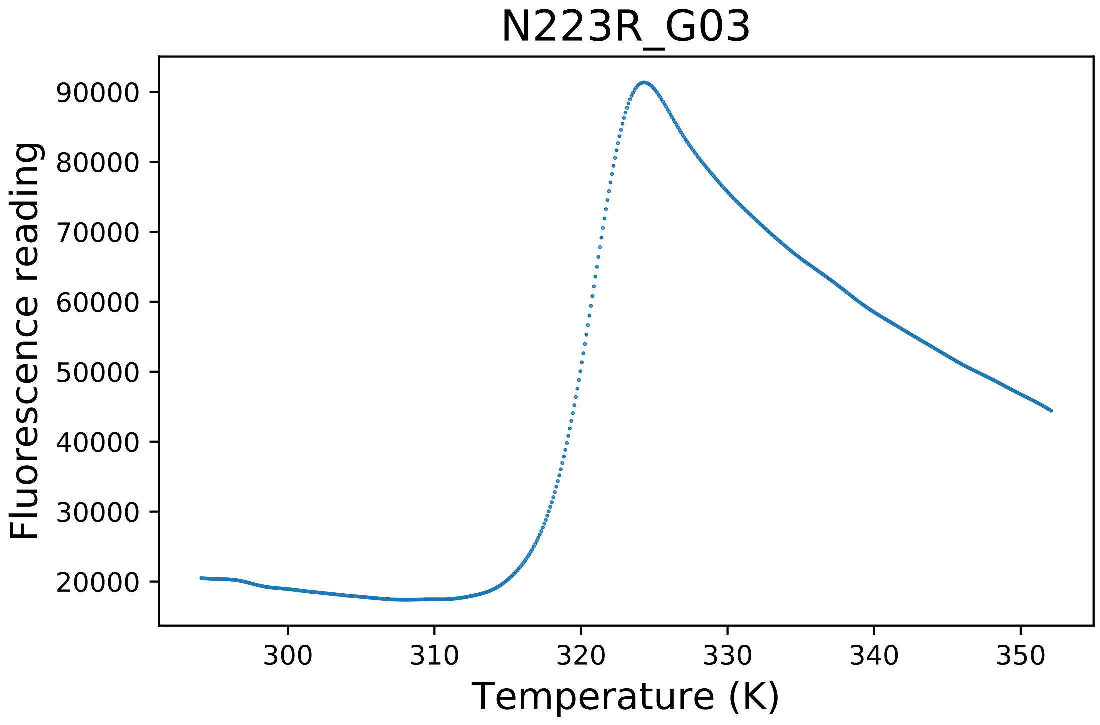

### N296A_B08_derivative.pdf

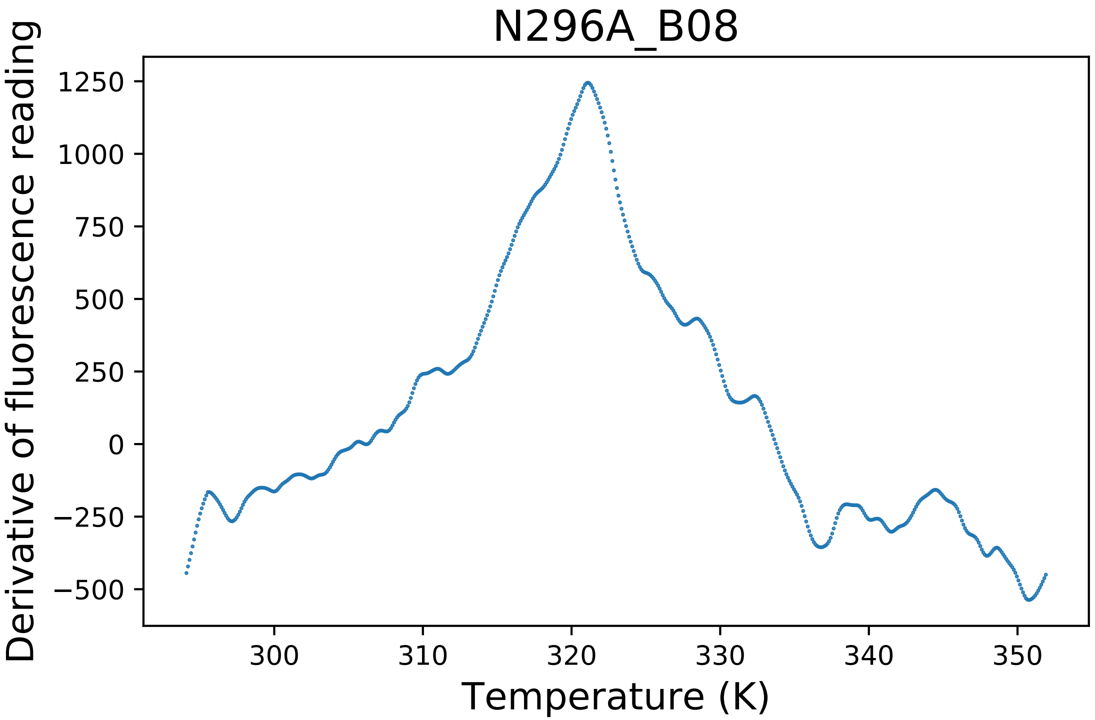

### N407A_A01_derivative.pdf

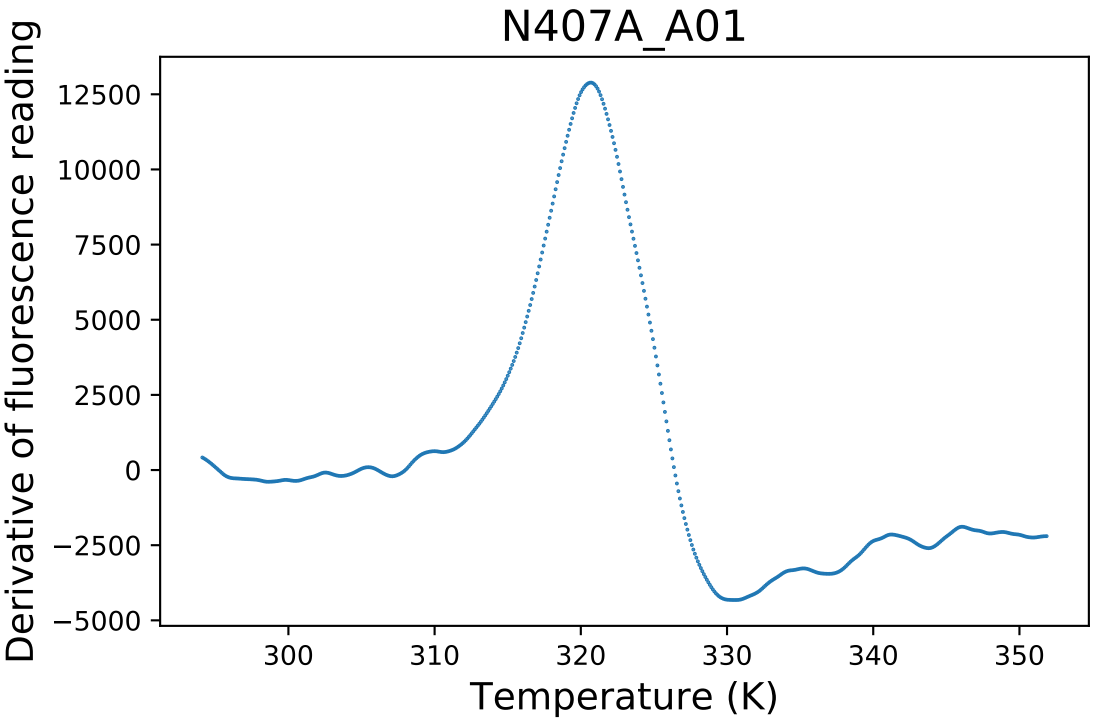

### N407A_A03_fluorescence.pdf

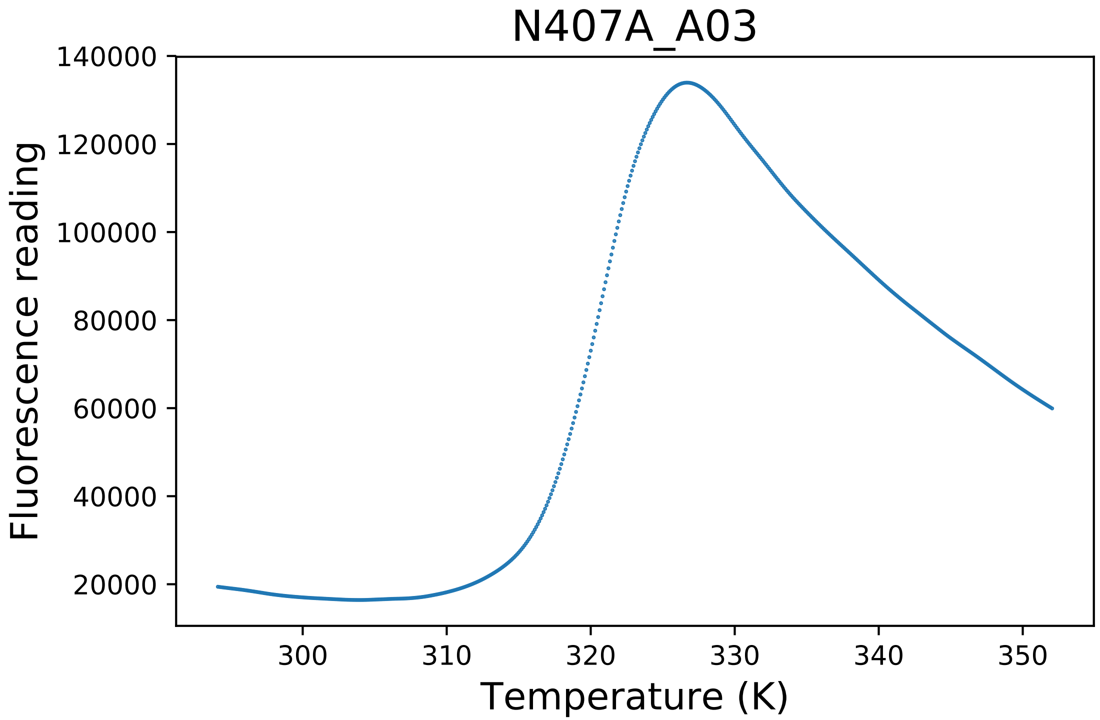

### N407C_C02_derivative.pdf

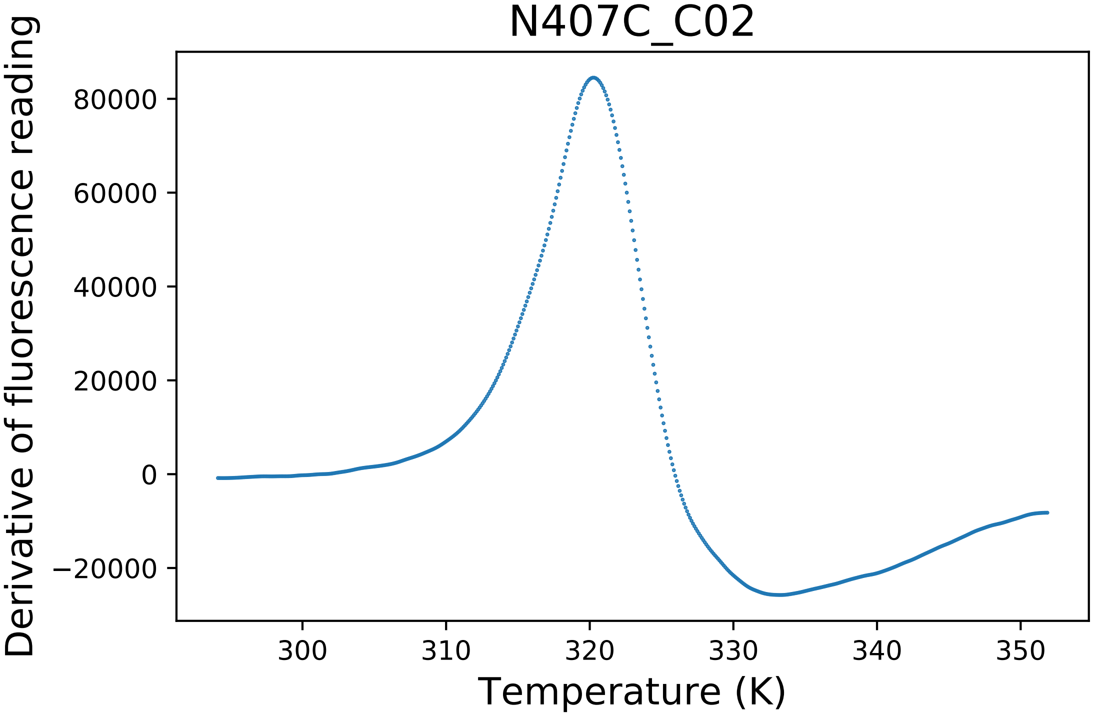

### N407C_C03_dg.pdf

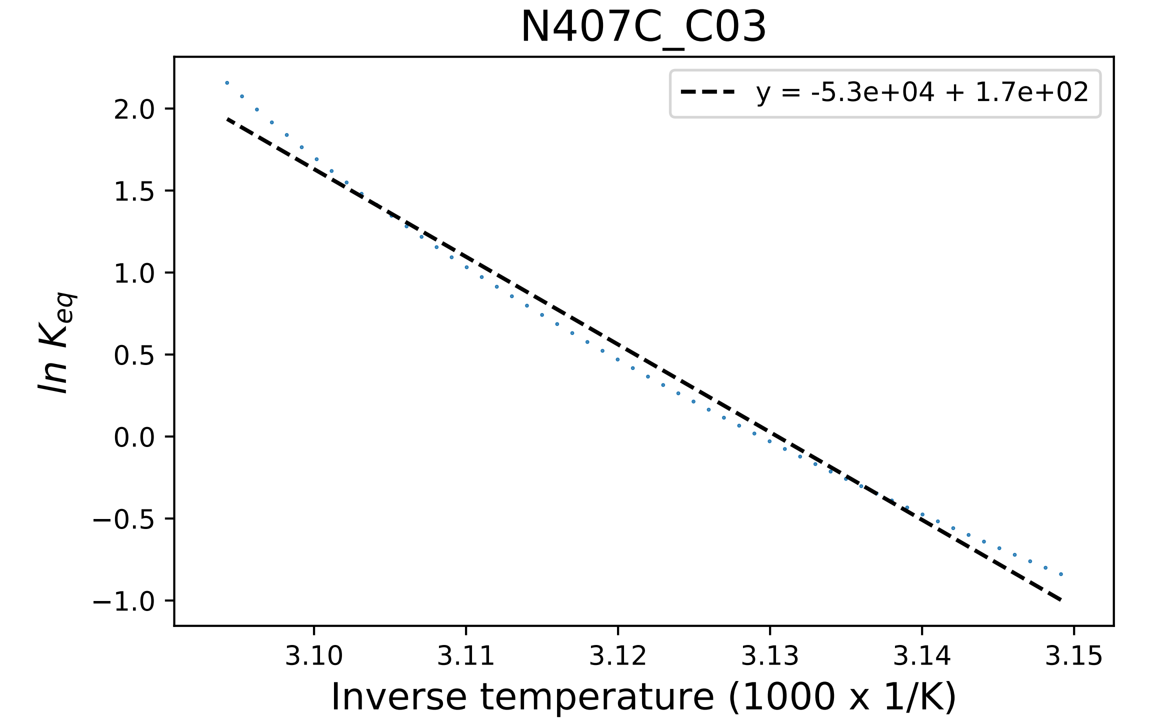

### N407C_C03_fluorescence.pdf

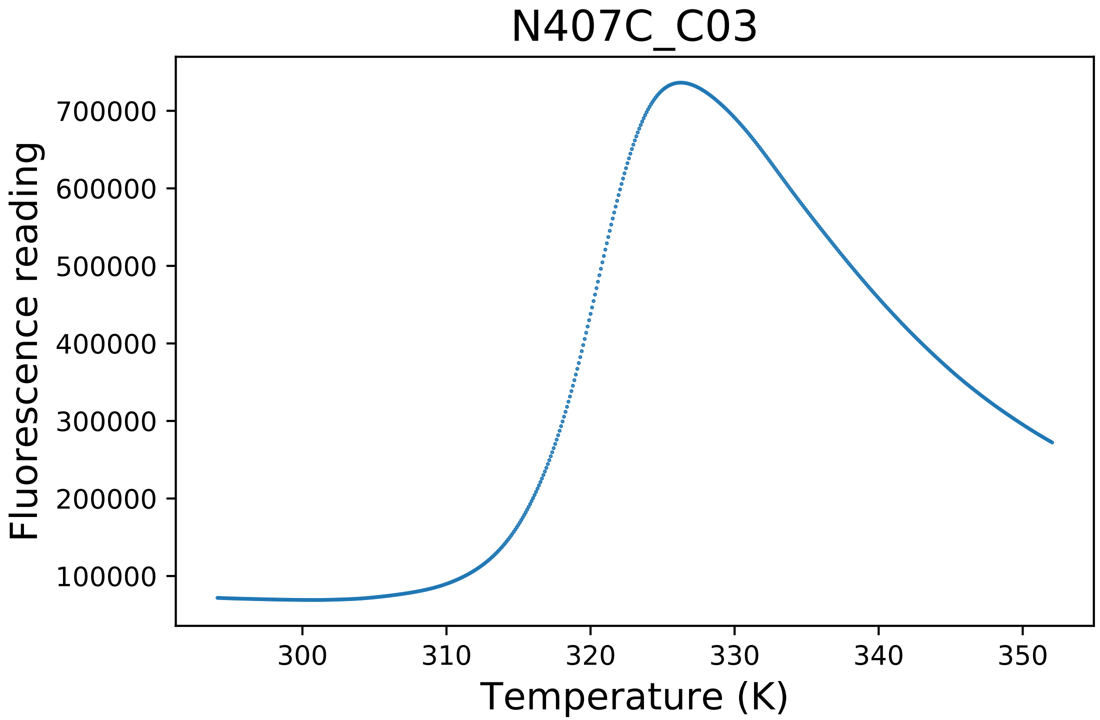

### Q22S_B06_derivative.pdf

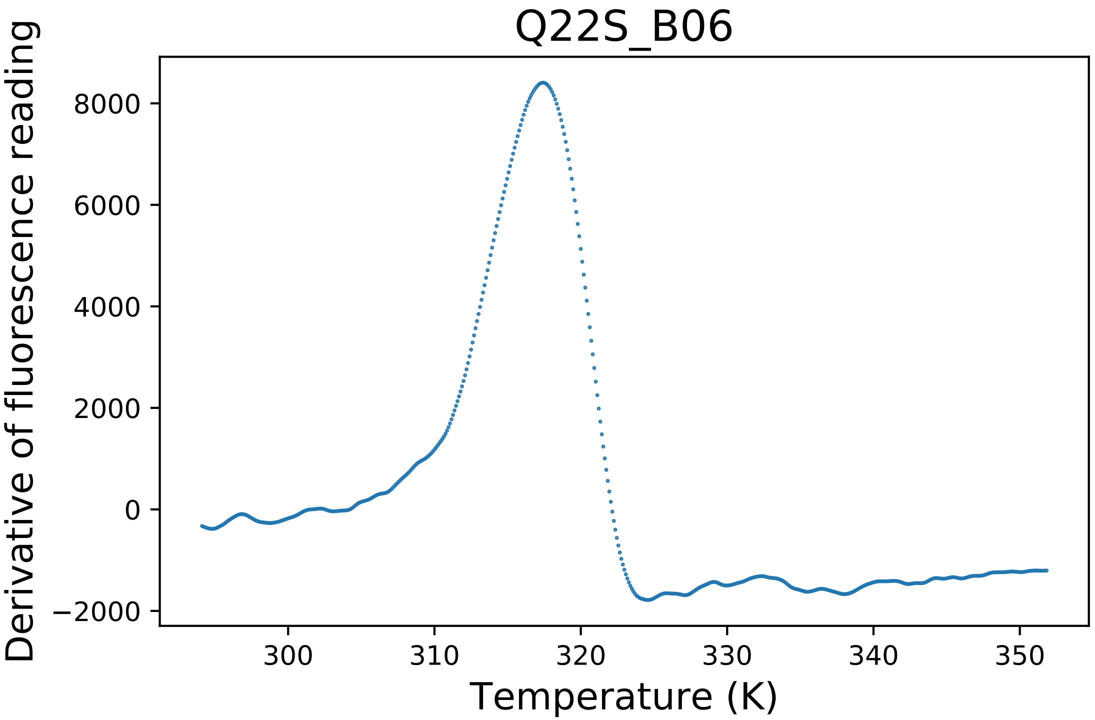

### S19A_A01_dg.pdf

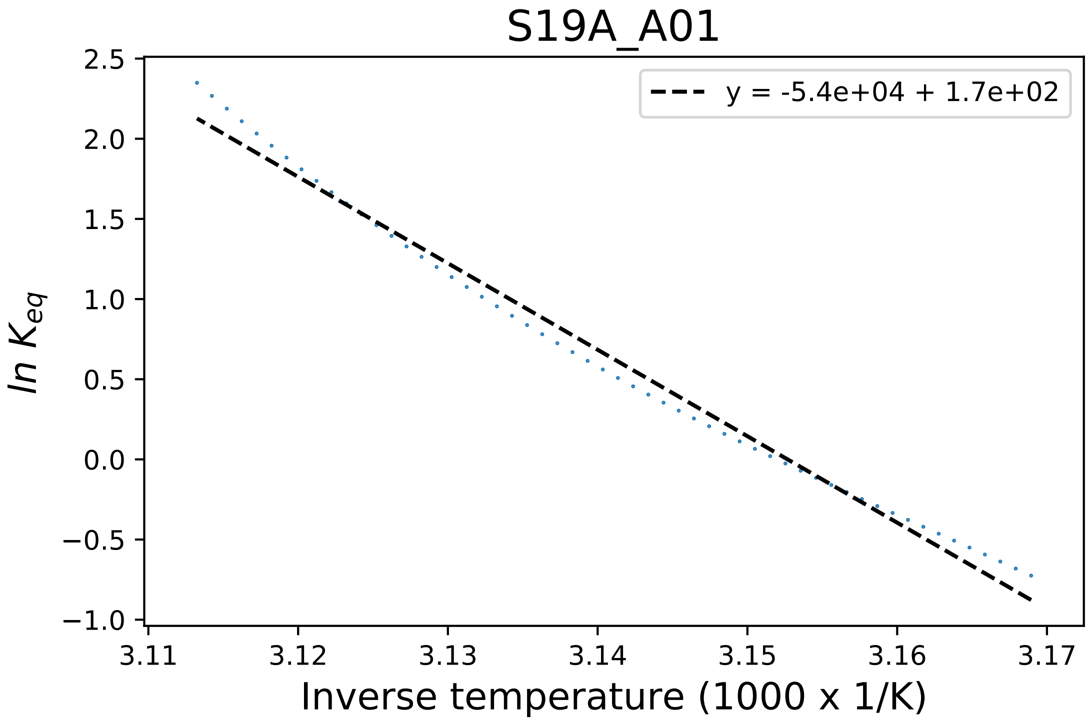

### S19A_A04_derivative.pdf

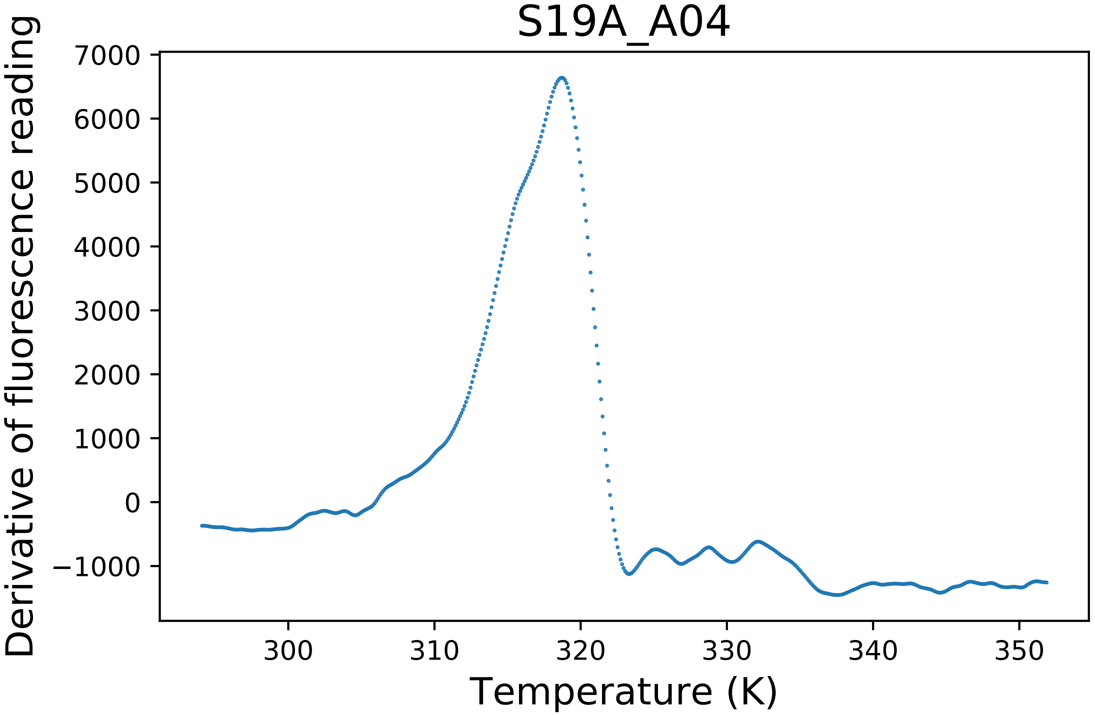

### T221A_F01_dg.pdf

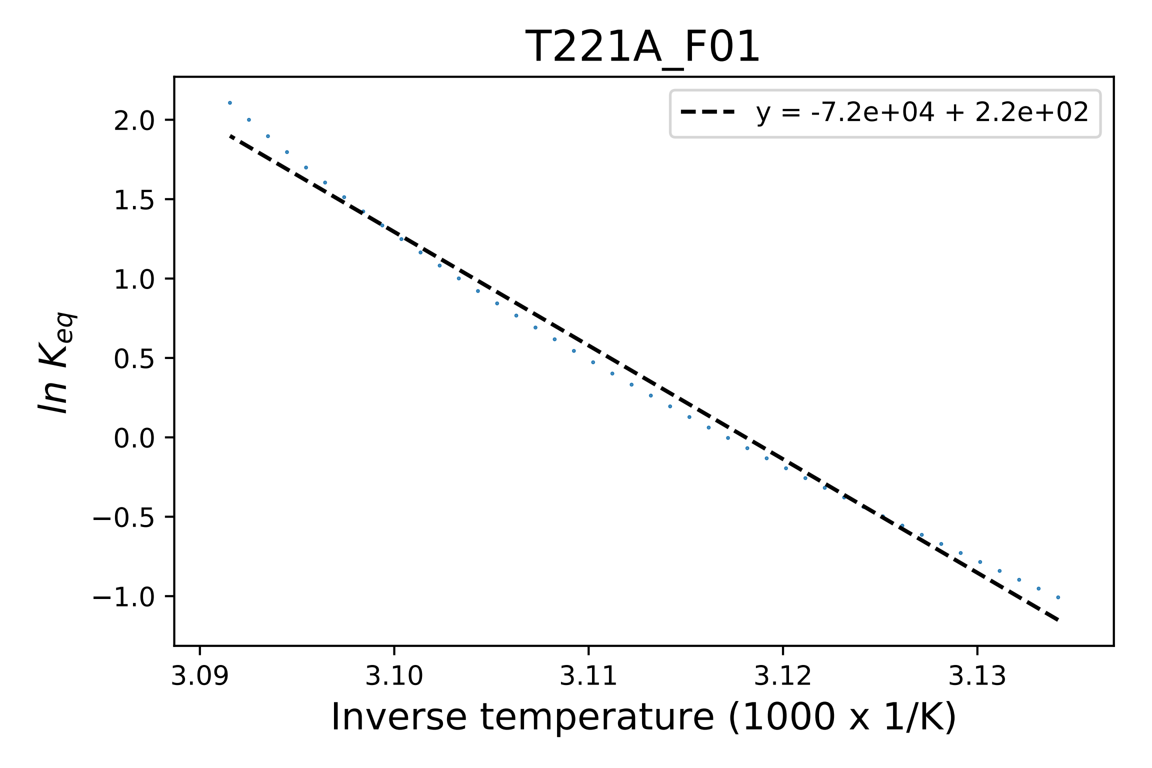

### W328A_C04_derivative.pdf

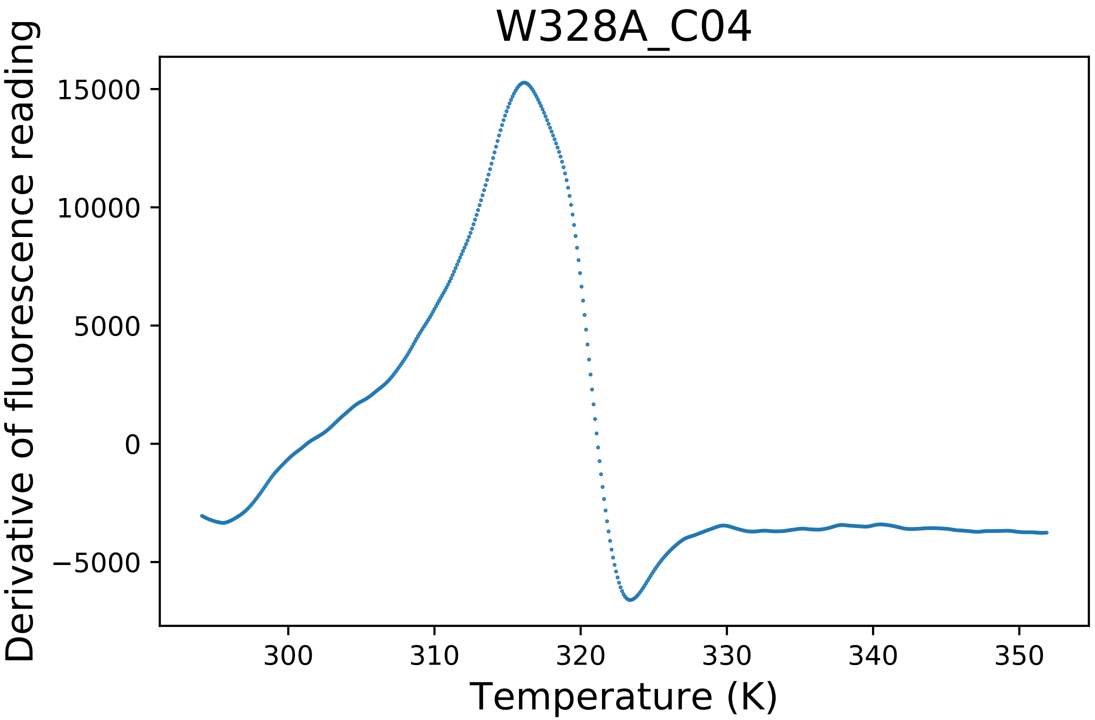

### W328C_A12_derivative.pdf

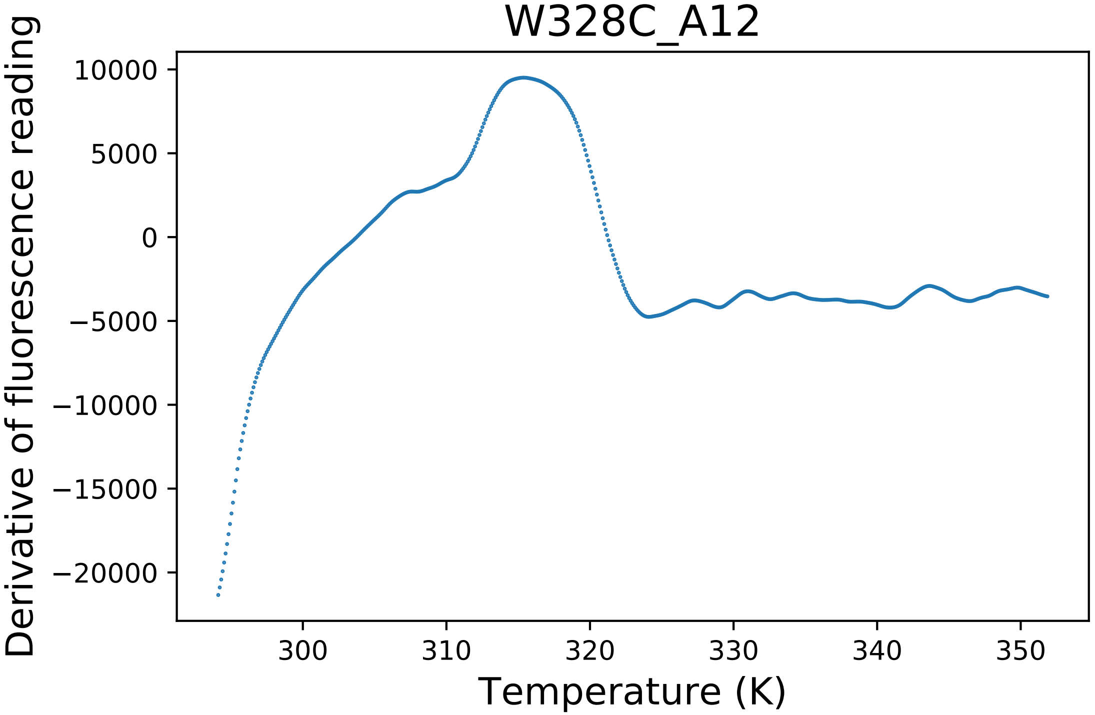

### W328C_A12_dg.pdf

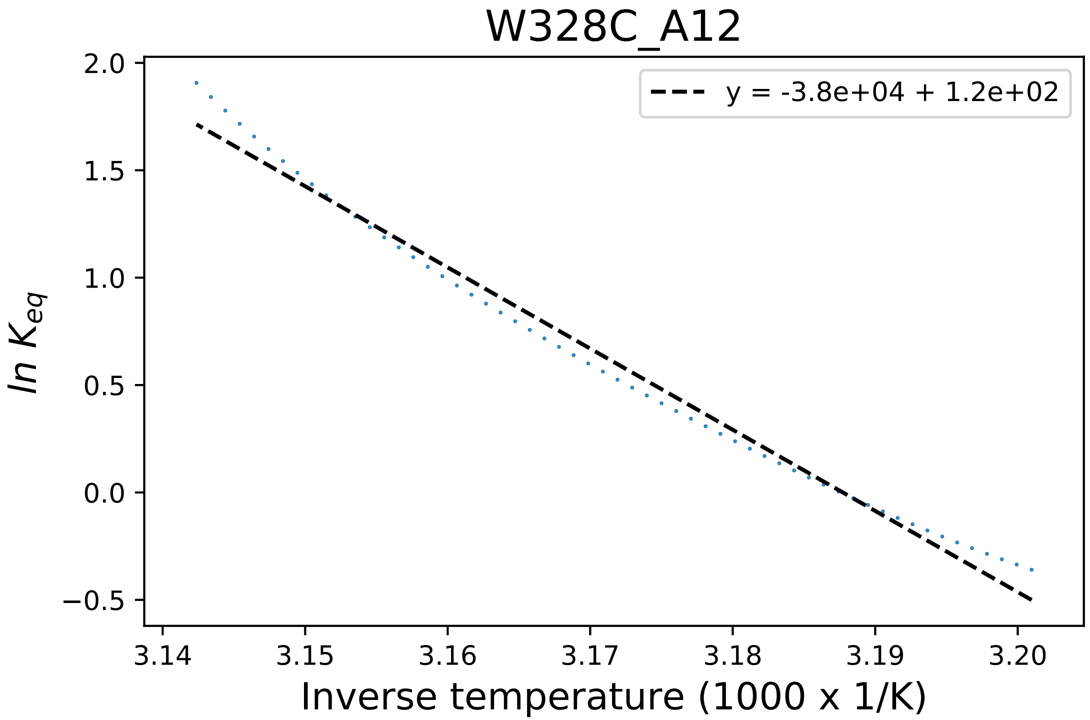

### W328H_A05_derivative.pdf

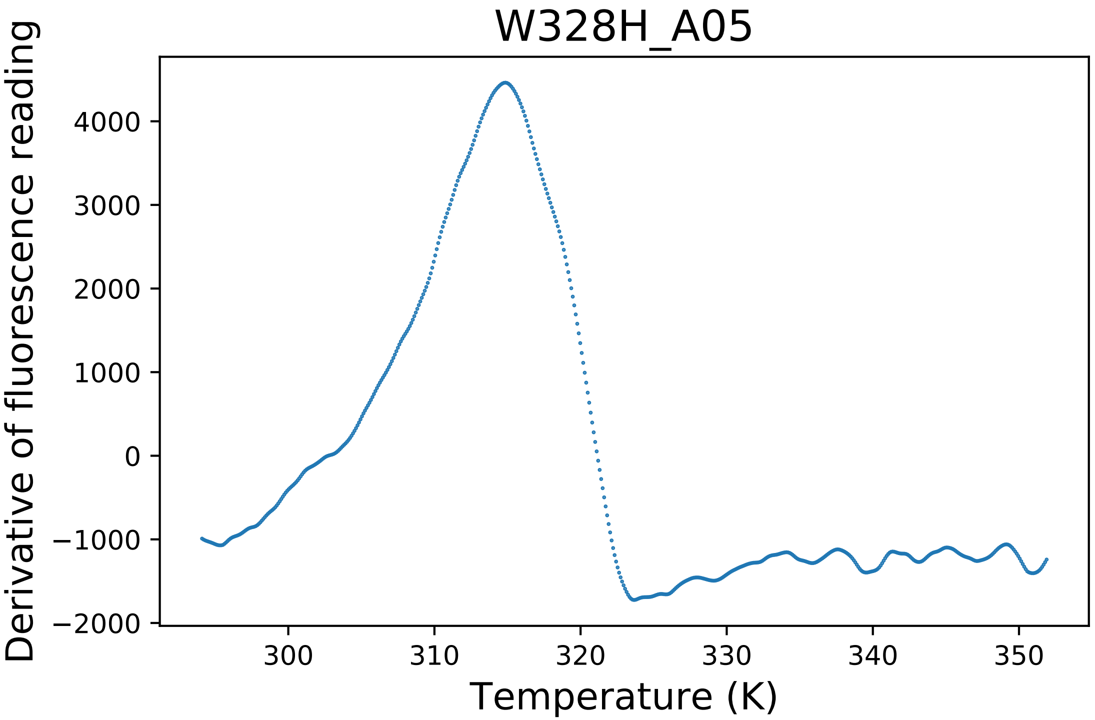

### W328R_D06_fluorescence.pdf

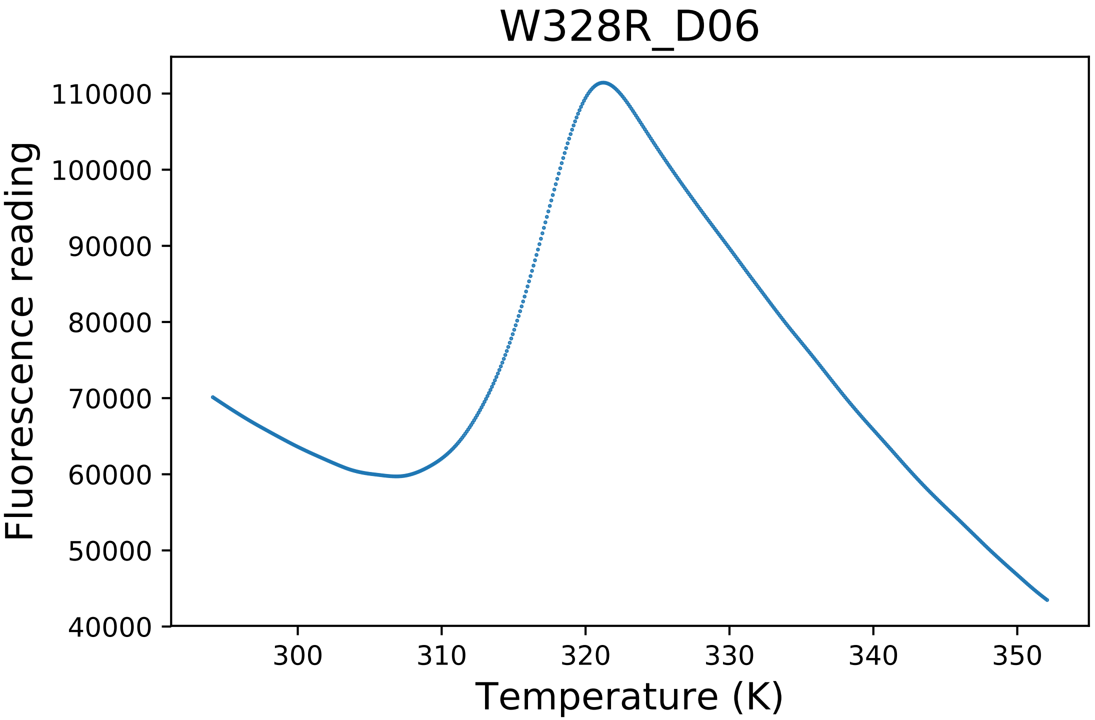

### W328R_D07_derivative.pdf

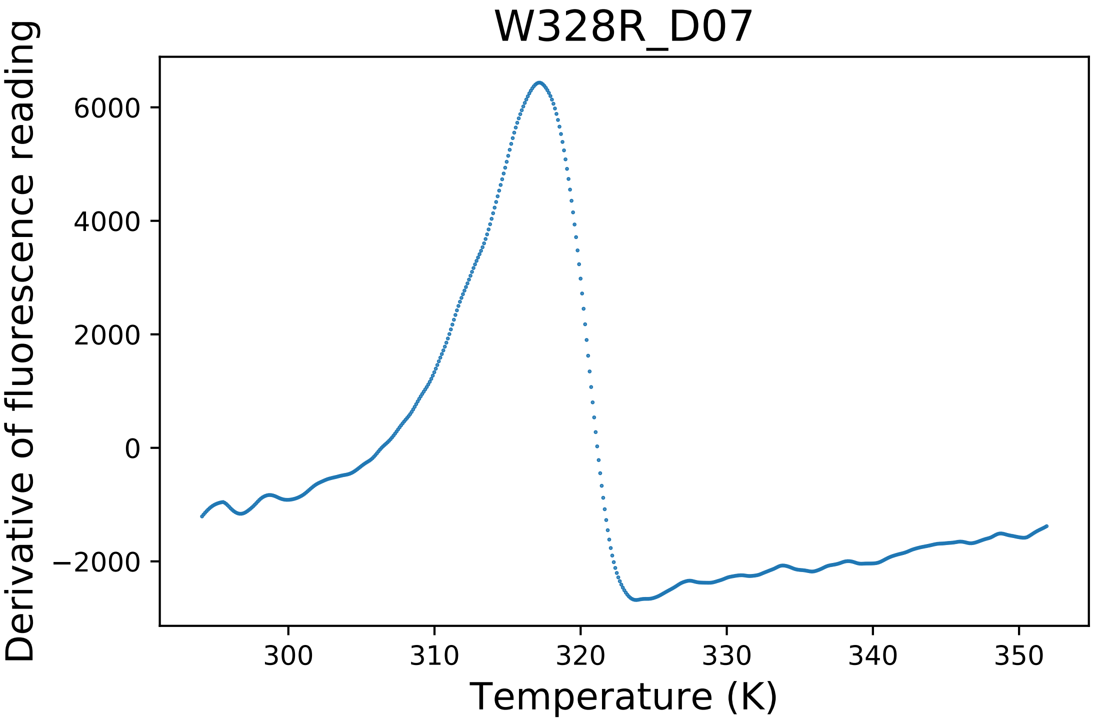

### W402R_A08_fluorescence.pdf

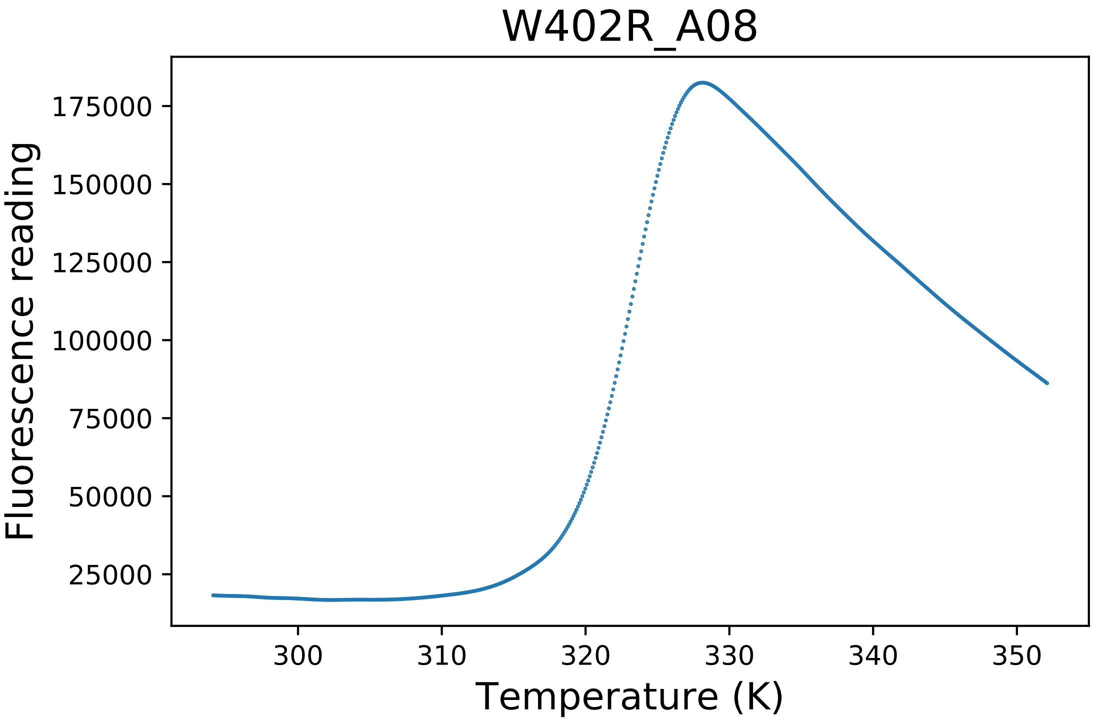

### W412Y_B08_fluorescence.pdf

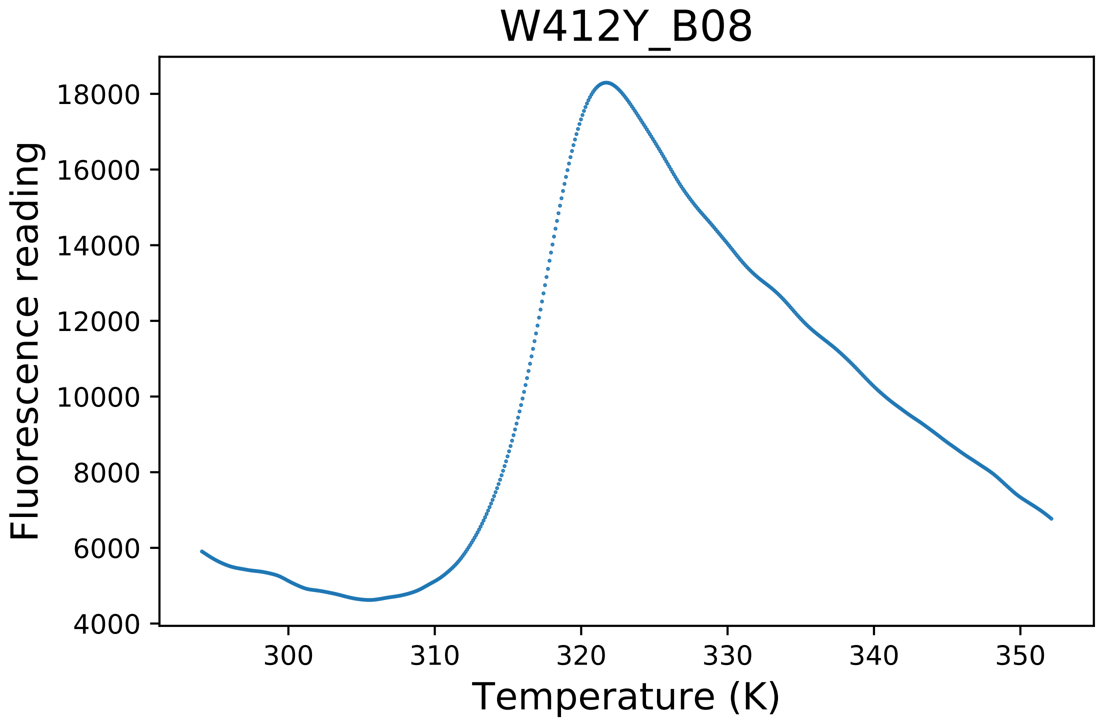

### WT6_D05_derivative.pdf

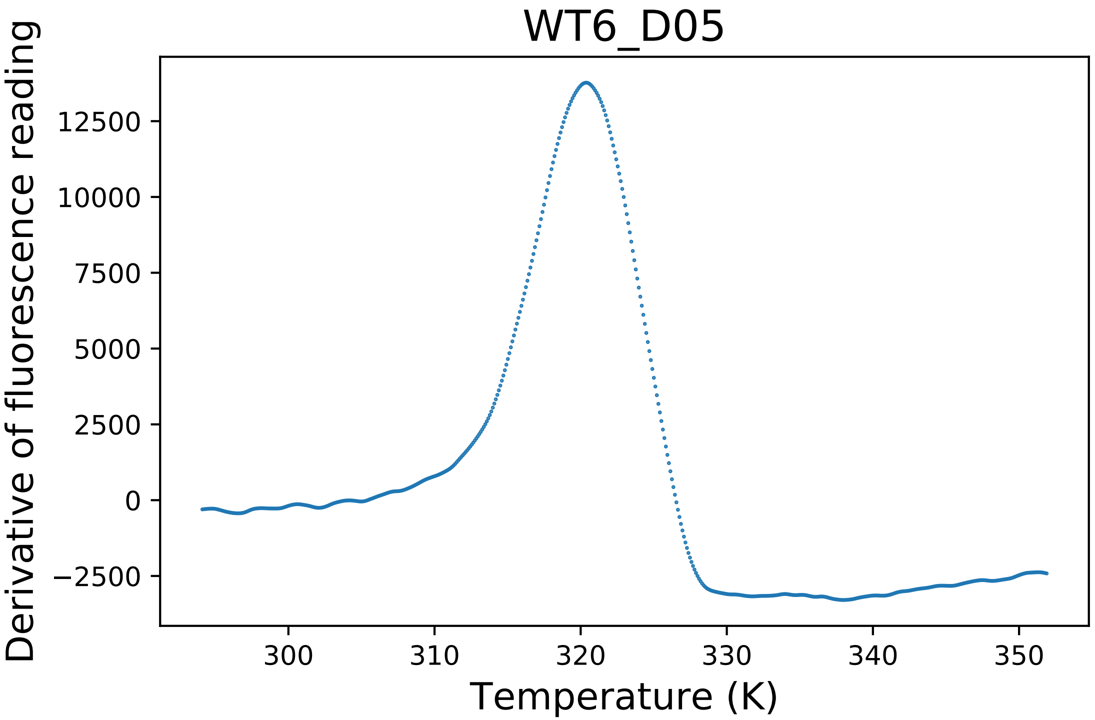

### Y21A_B02_dg.pdf

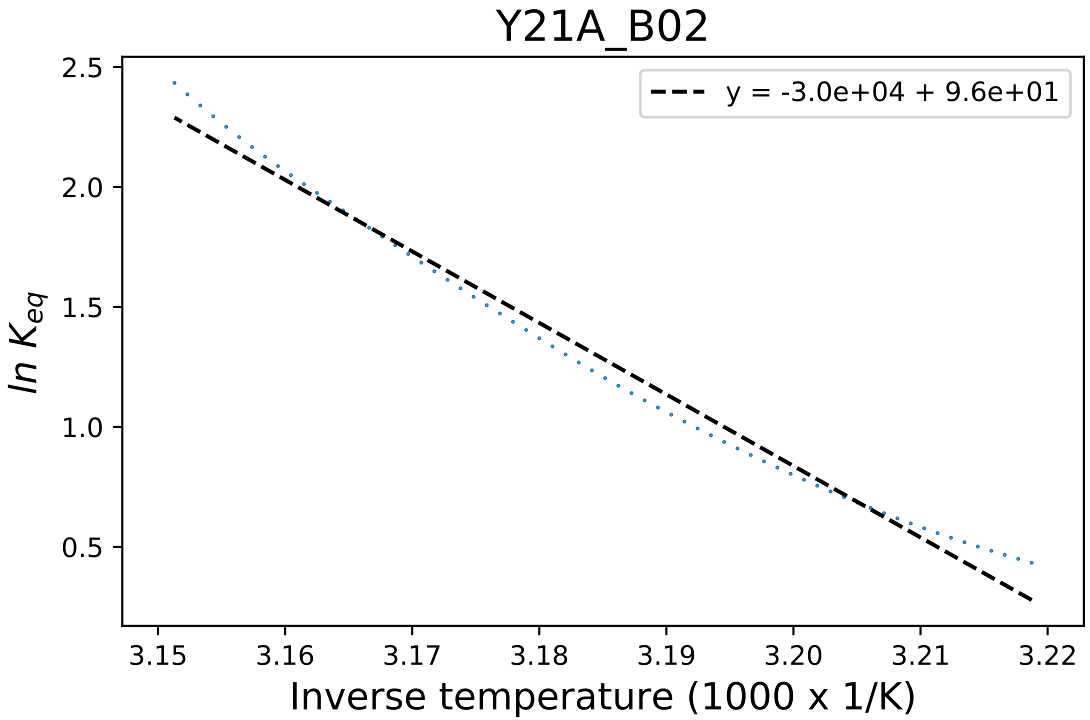

### Y297A_H04_fluorescence.pdf

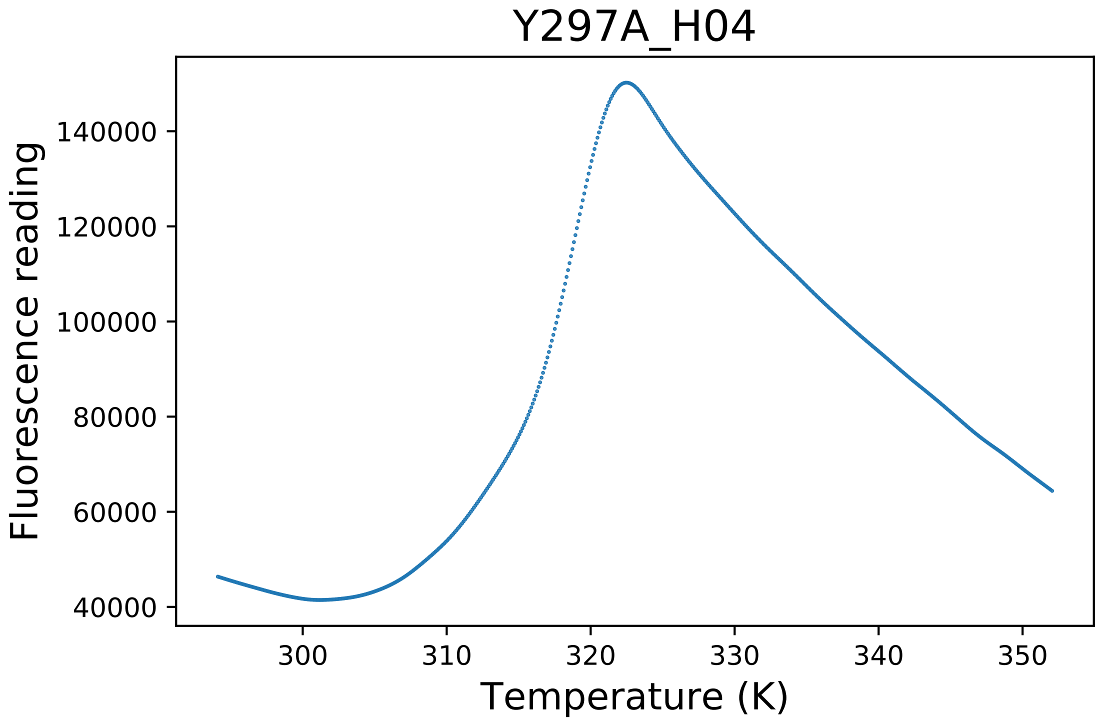
