## Supplementary figures and images for "Evaluating molecular modeling tools for thermal stability using an independently generated dataset"

### C170A_C05_fluorescence.pdf

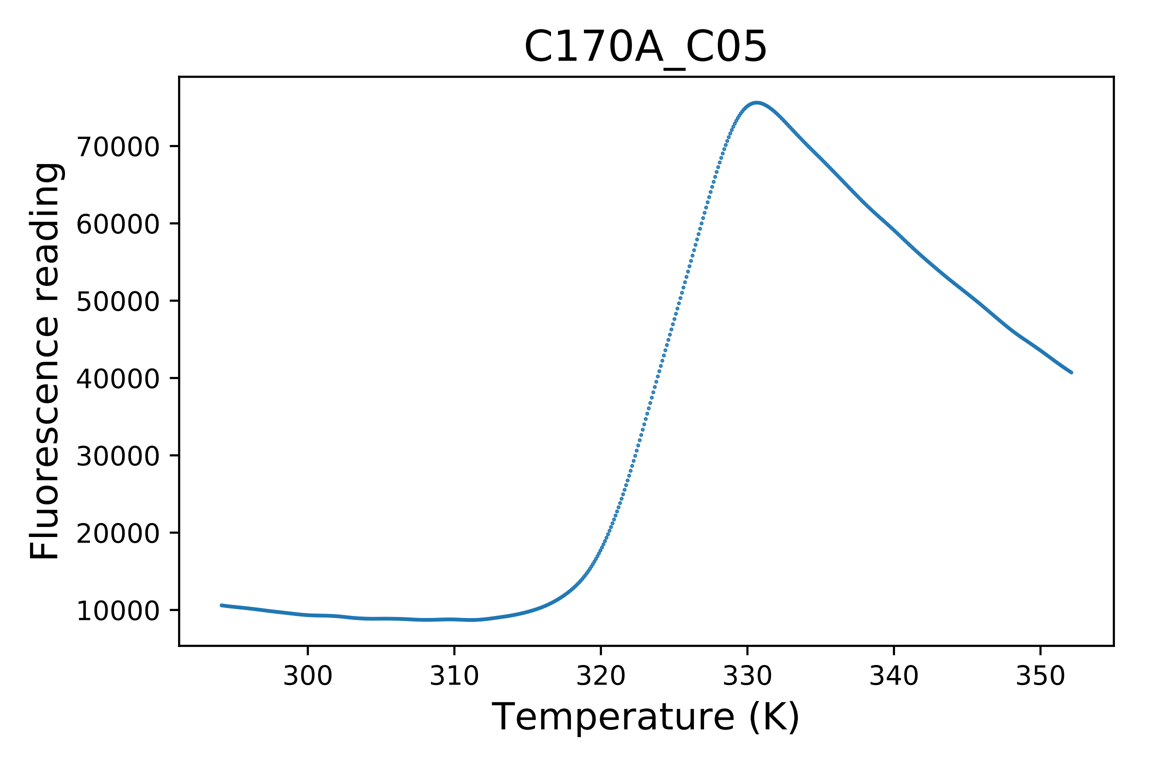

### C170Q_C10_derivative.pdf

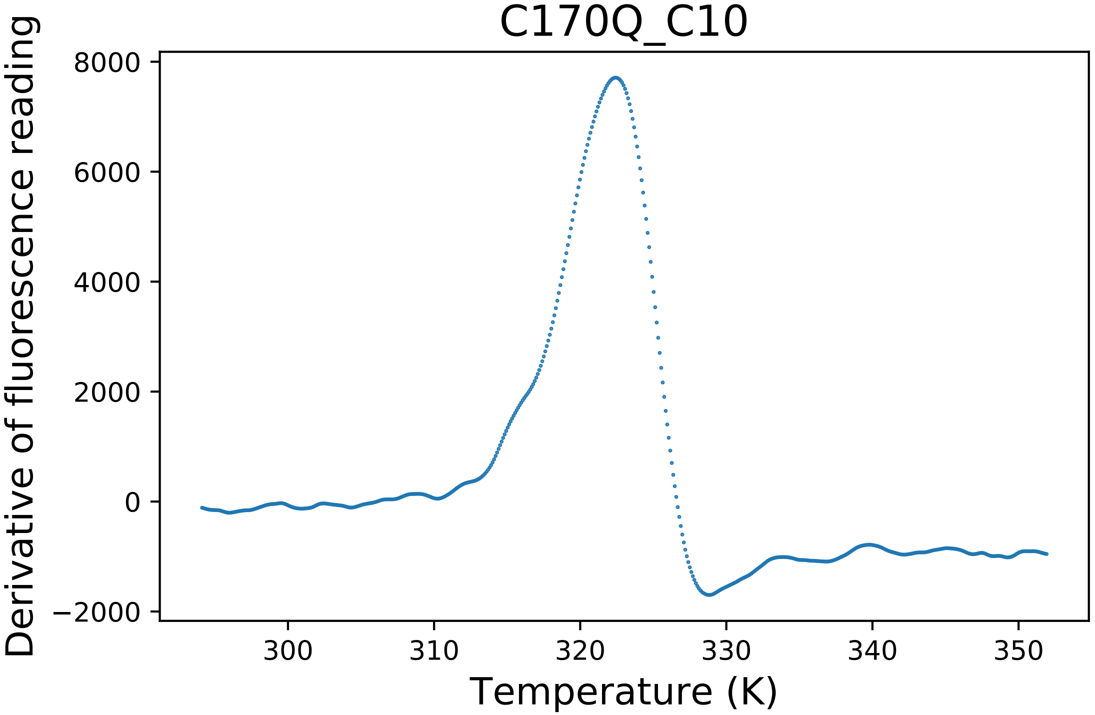

### C170Q_C11_derivative.pdf

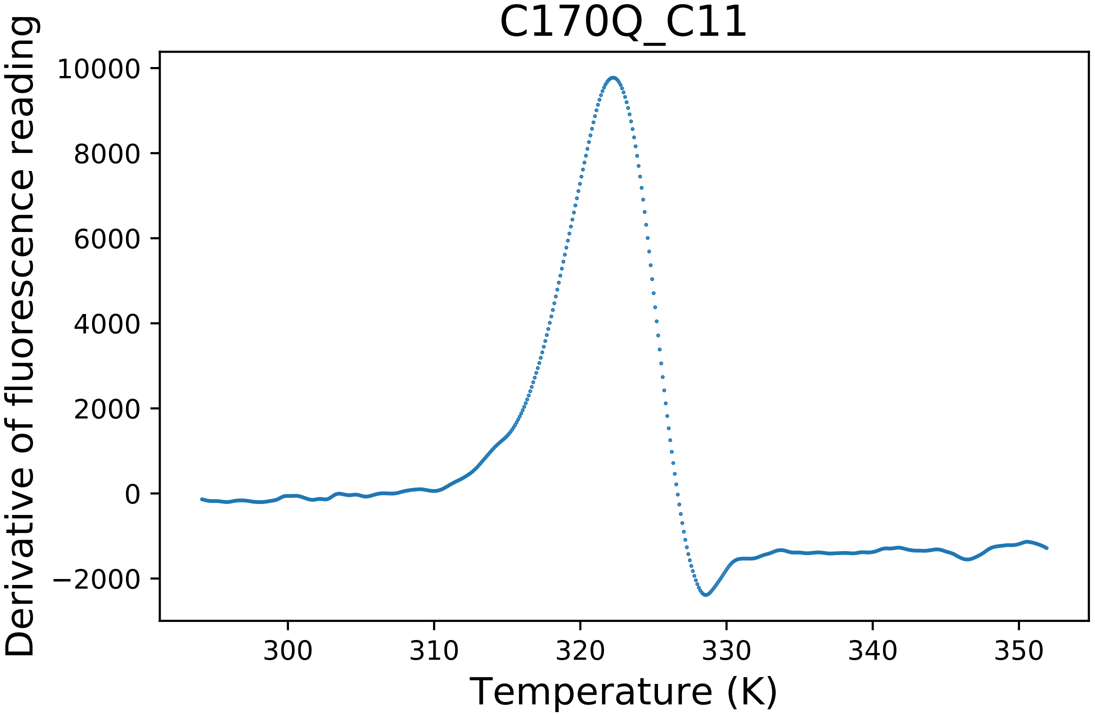

### E157D_A07_fluorescence.pdf

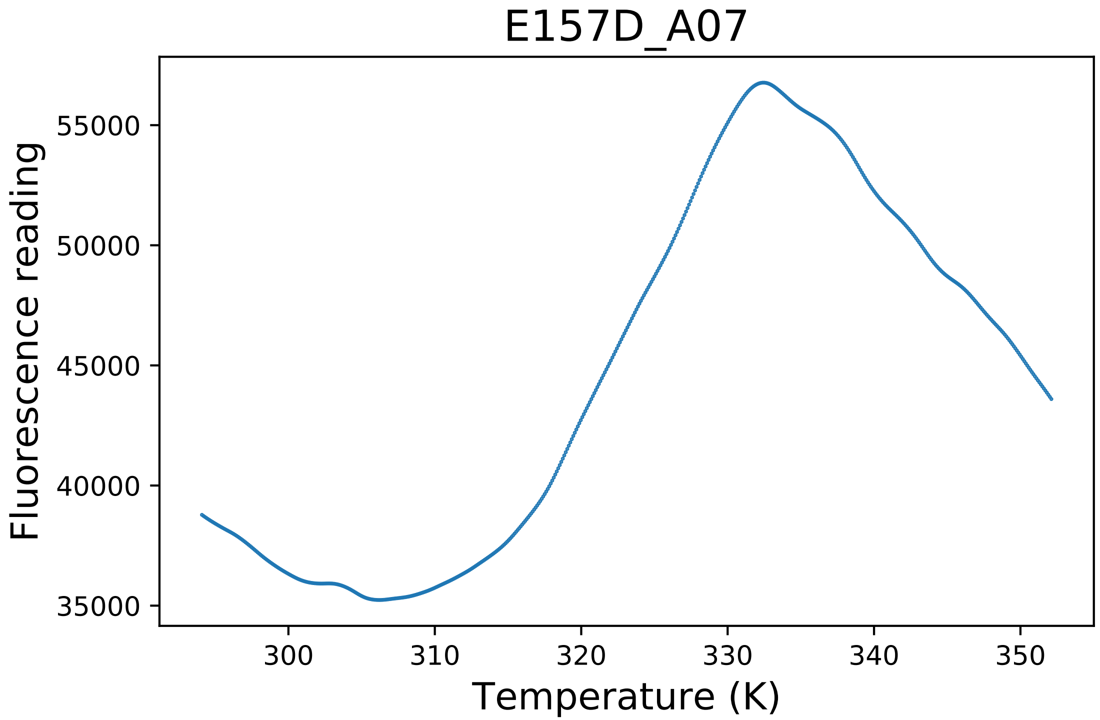

### E157D_A08_fluorescence.pdf

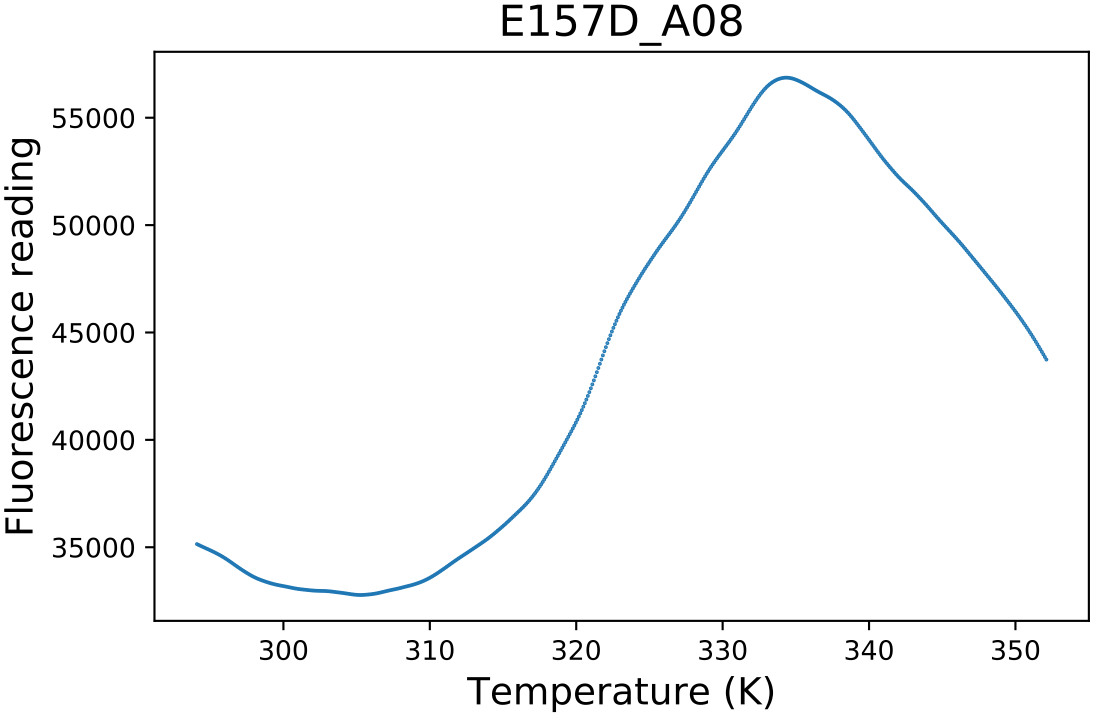

### E167A_A09_dg.pdf

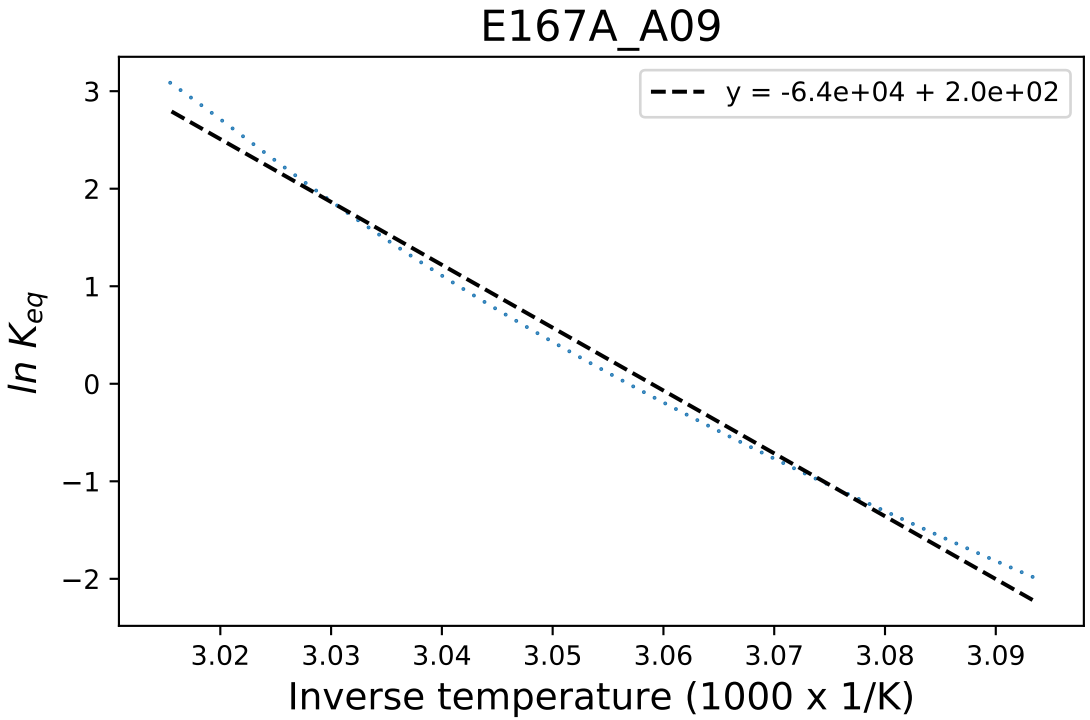

### E167A_A10_derivative.pdf

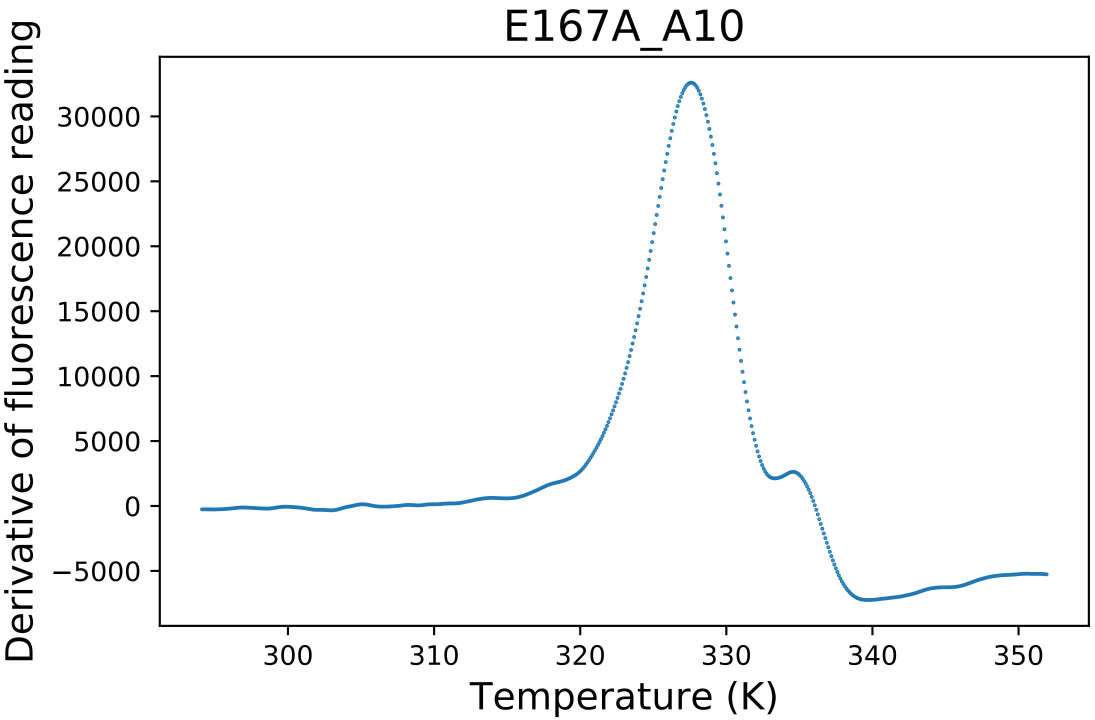

### E167A_A11_derivative.pdf

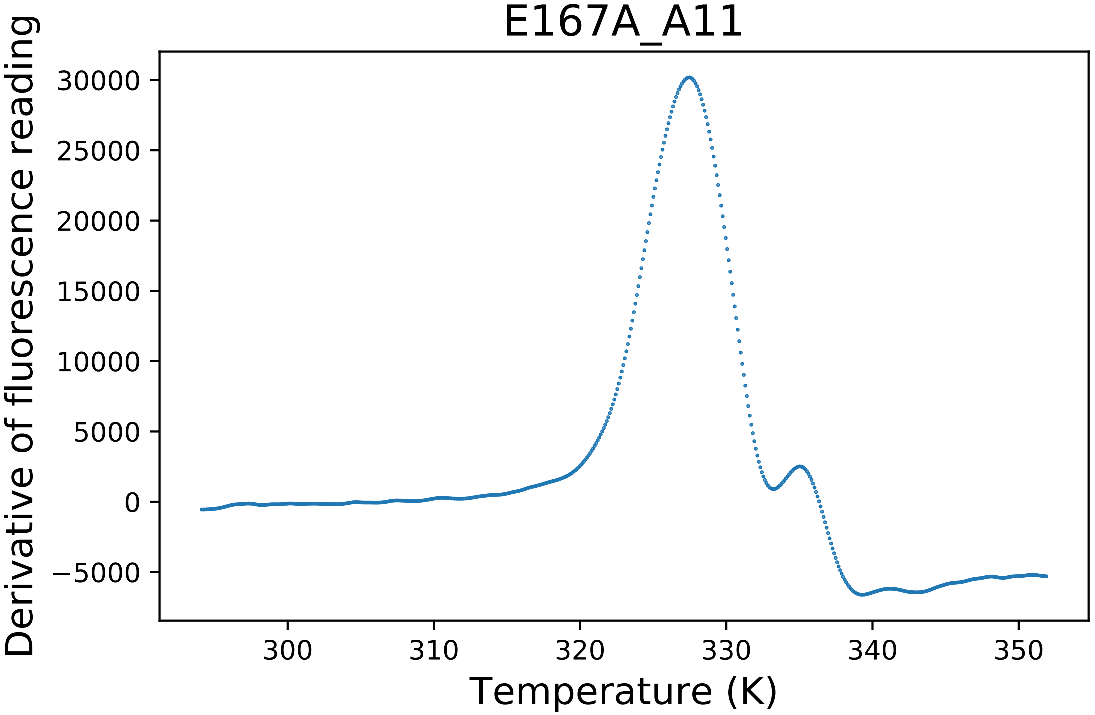

### E225A_A02_fluorescence.pdf

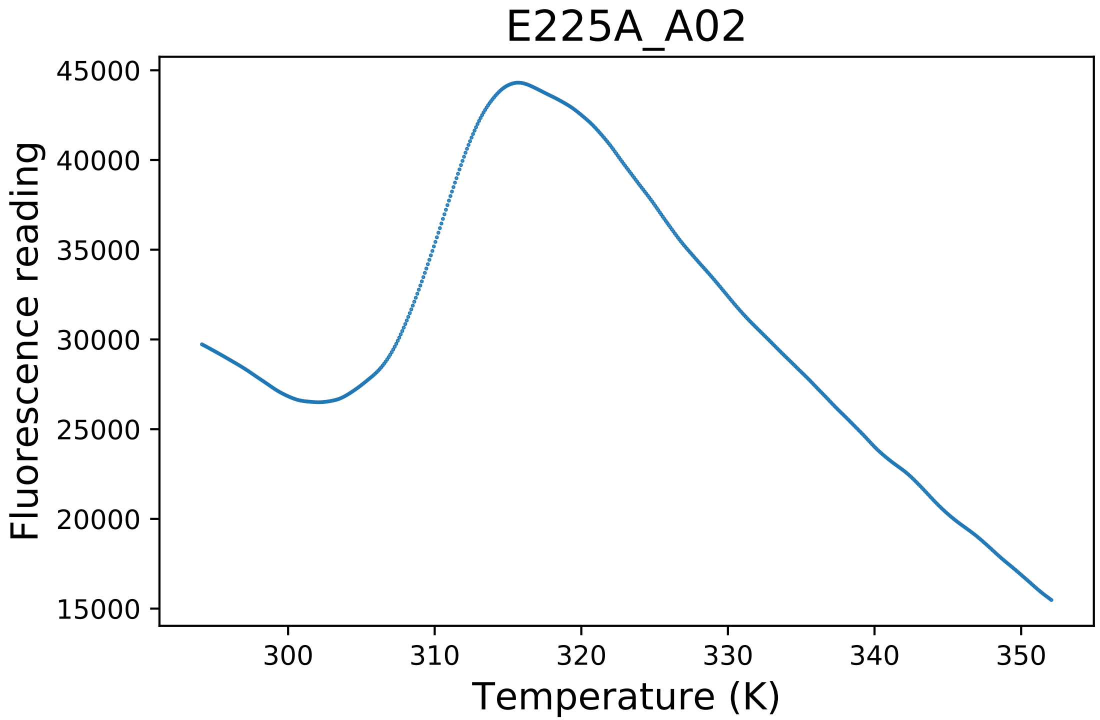

### E225H_C09_dg.pdf

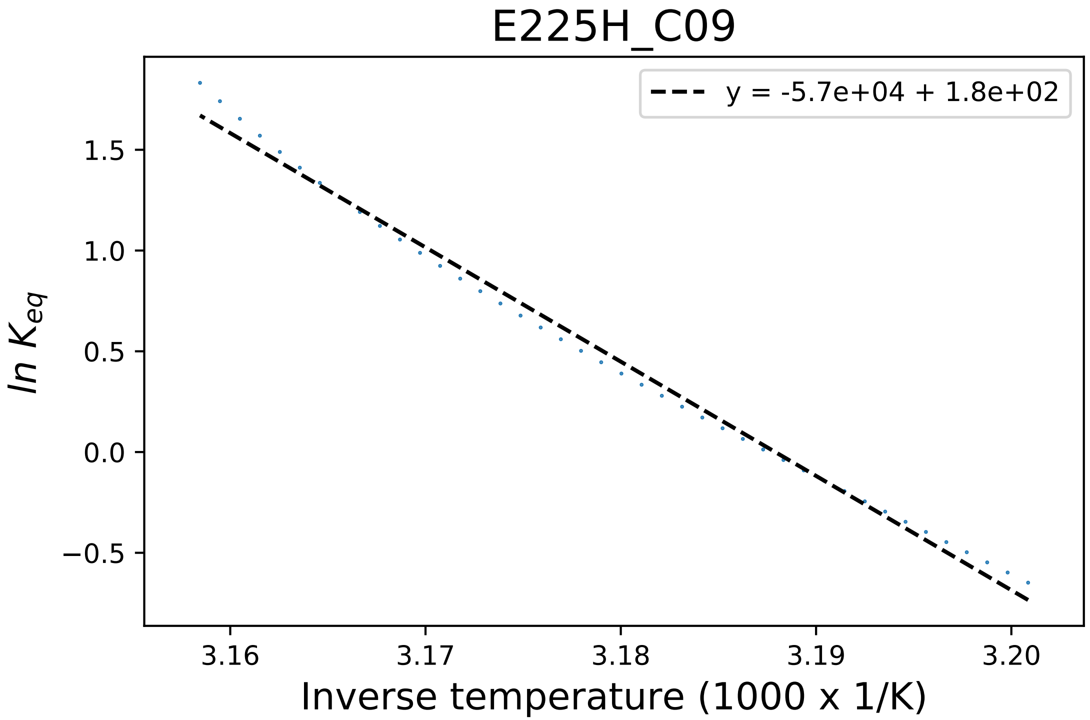

### E225H_C12_fluorescence.pdf

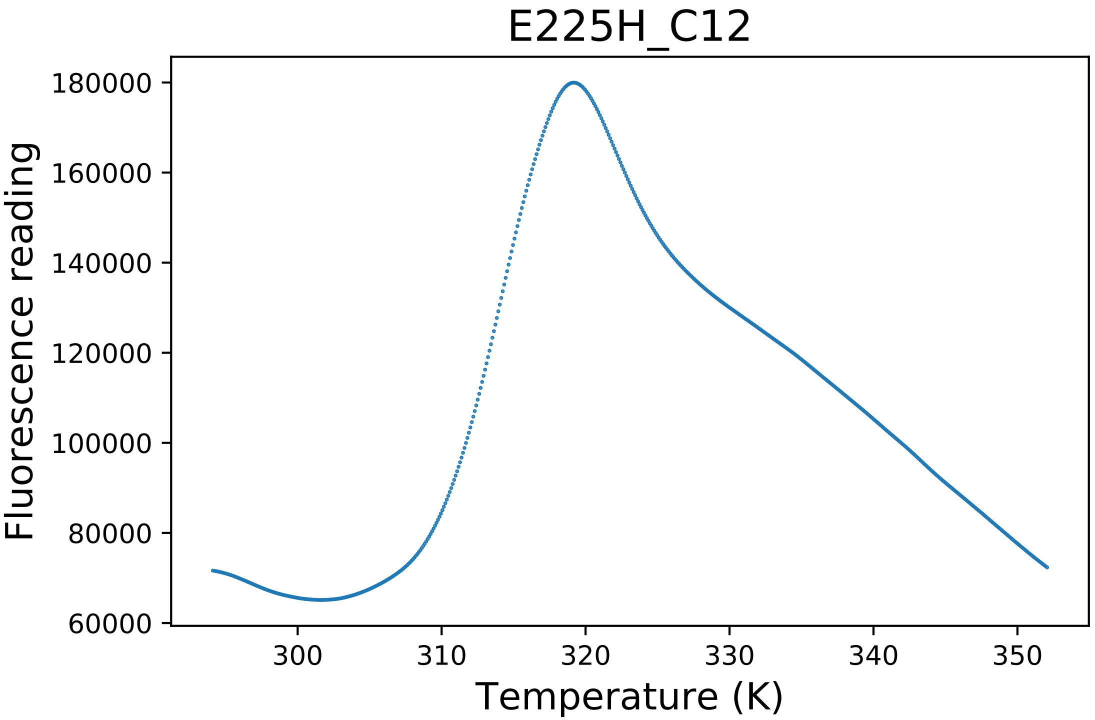

### E409A_C06_dg.pdf

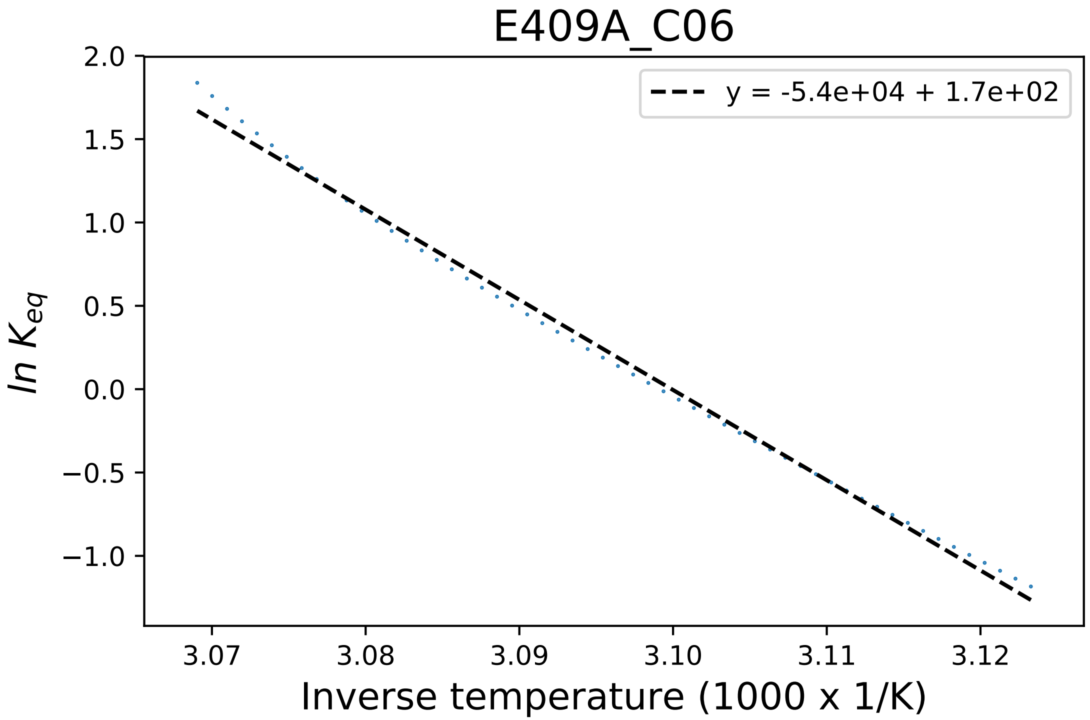

### E409A_C06_fluorescence.pdf

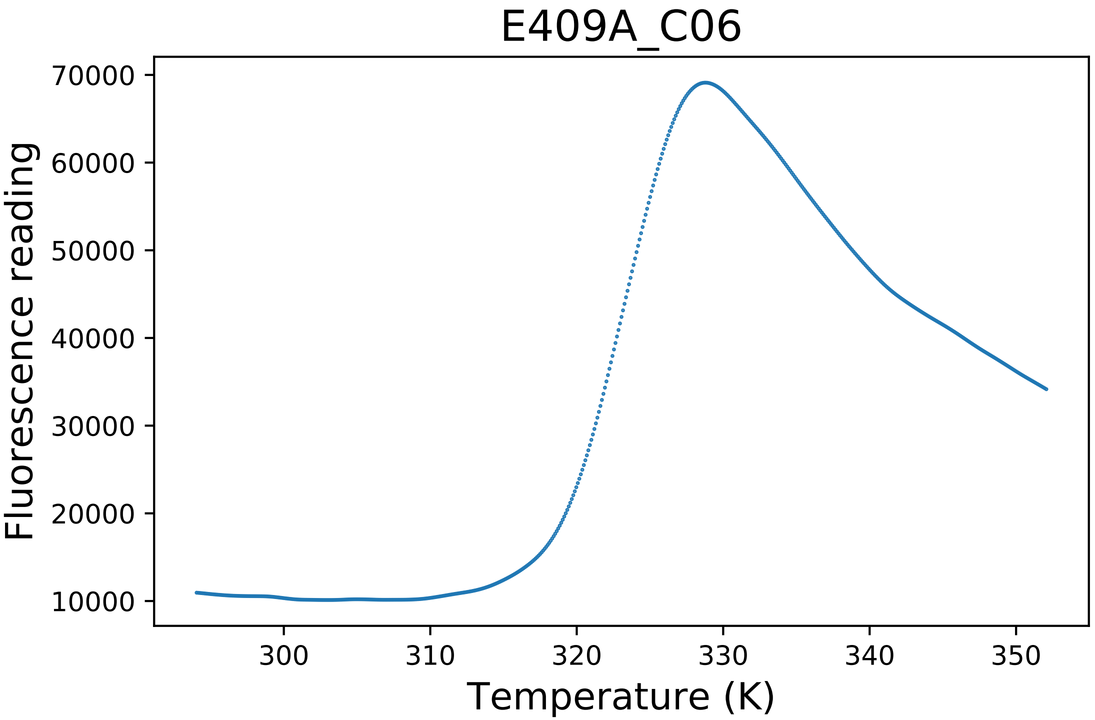

### E409D_F03_dg.pdf

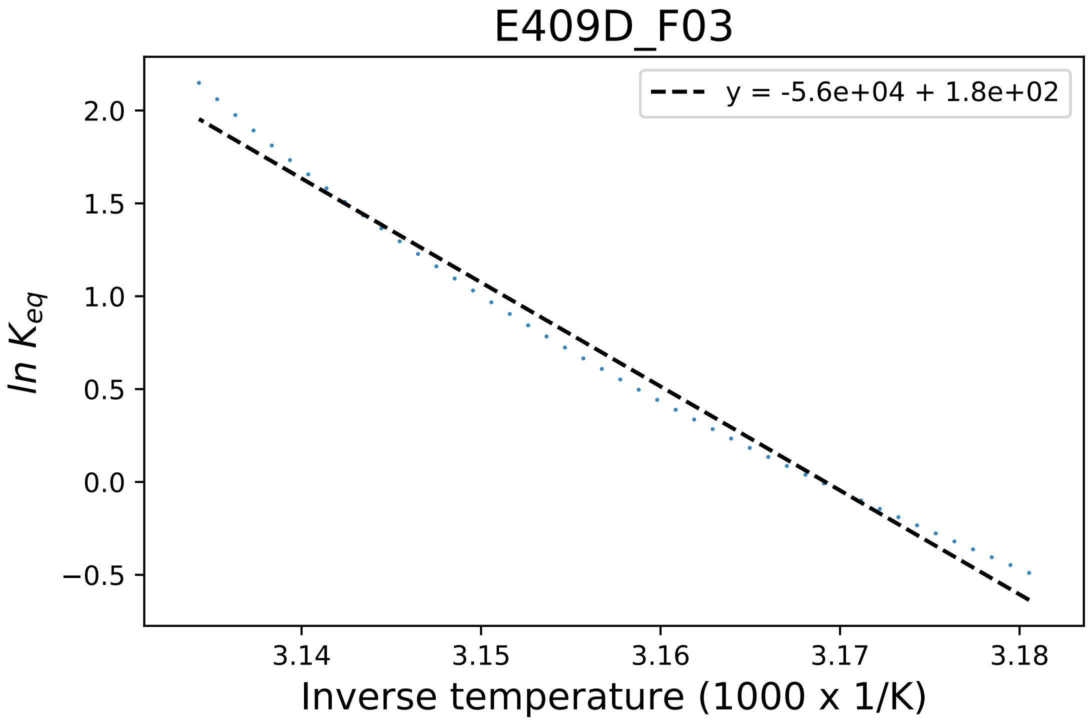

### E409D_F03_fluorescence.pdf

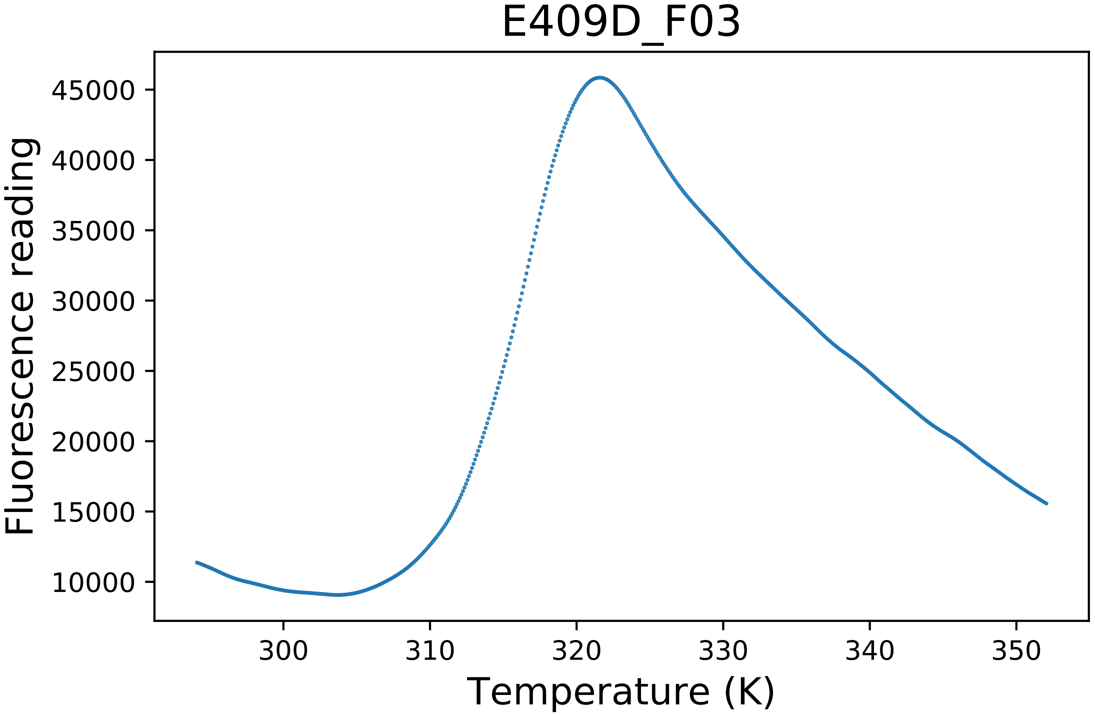

### E426S_E06_fluorescence.pdf

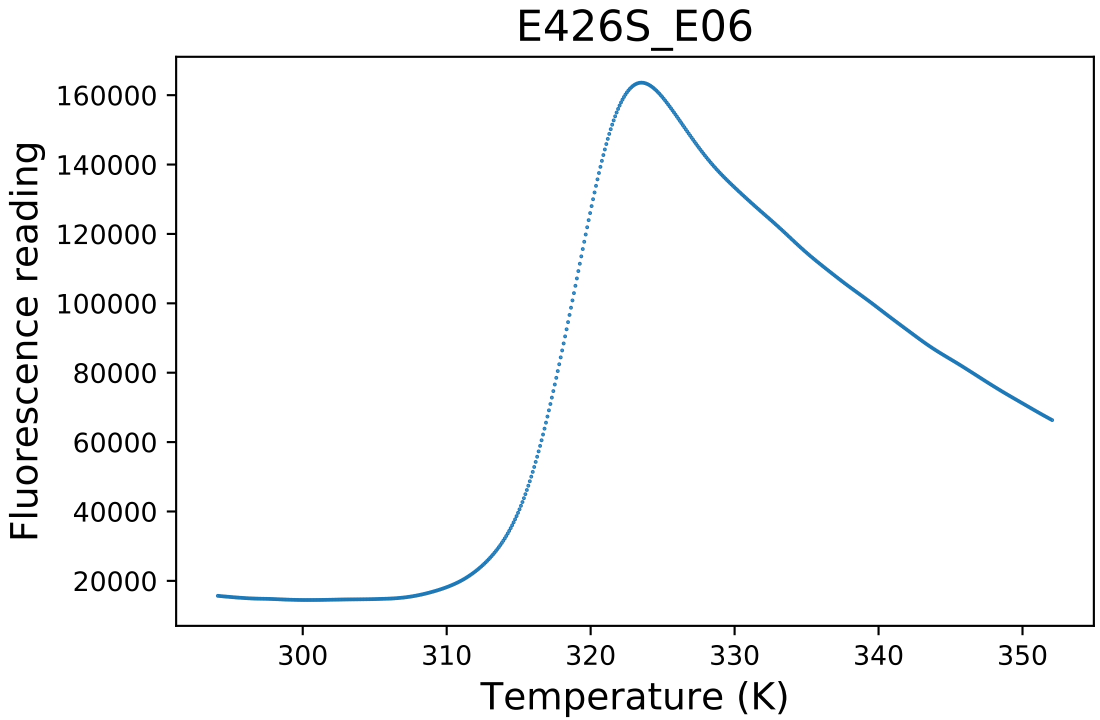

### F418A_H01_dg.pdf

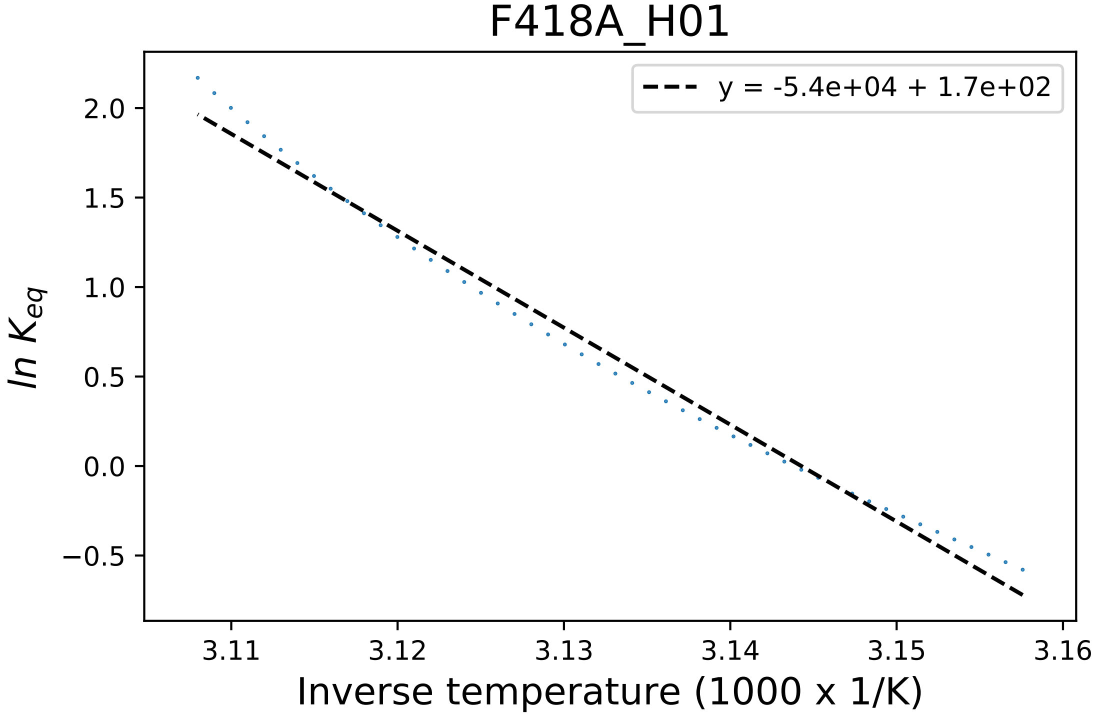

### F418A_H03_dg.pdf

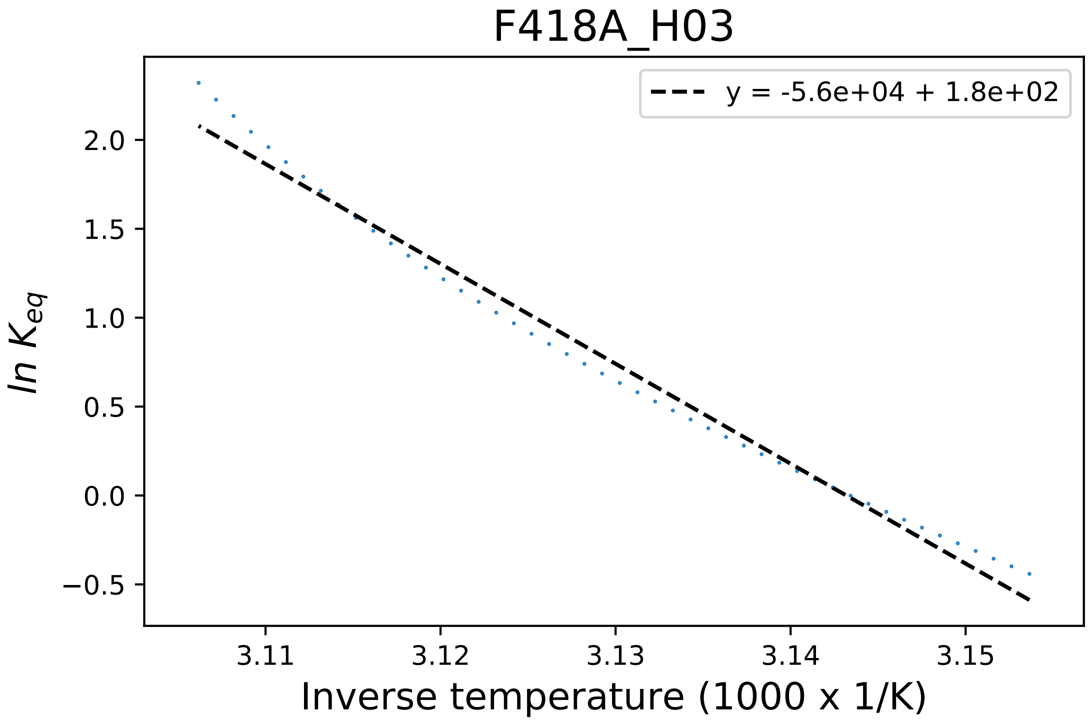

### I247E_D02_dg.pdf

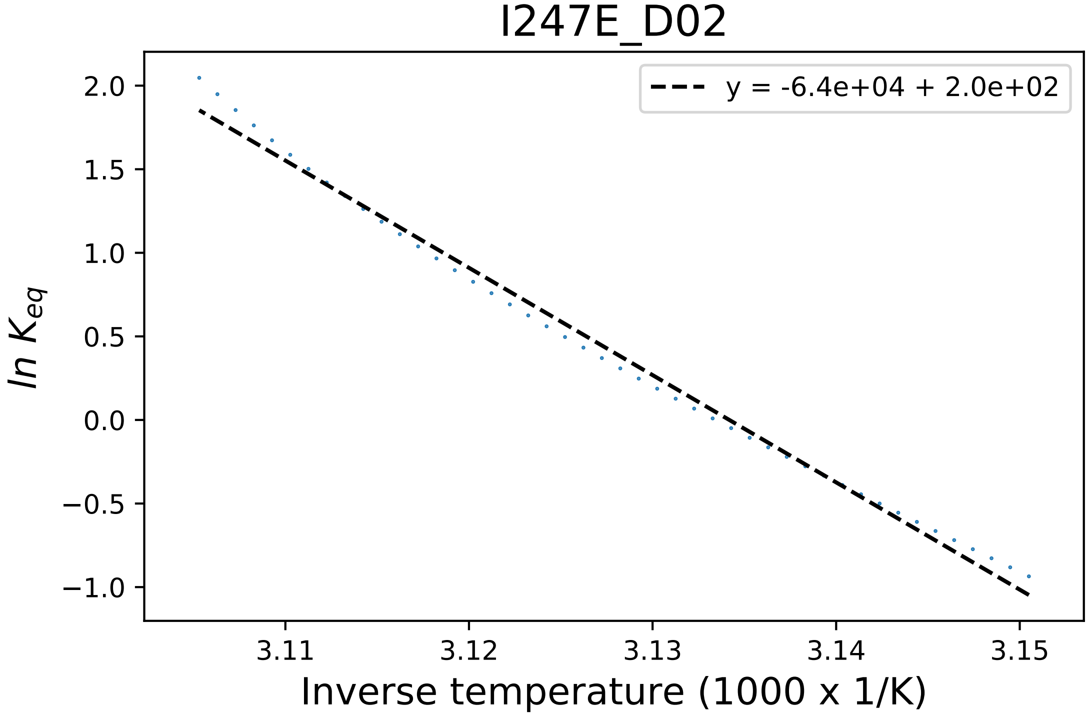

### I247E_D03_fluorescence.pdf

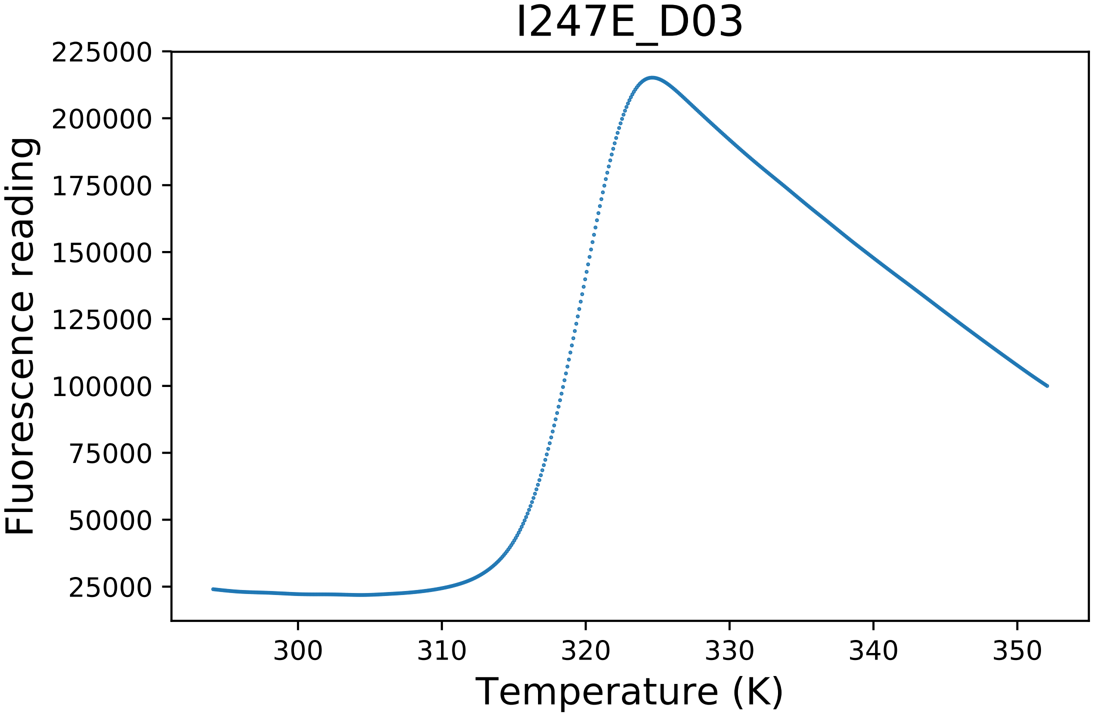

### I247N_B03_dg.pdf

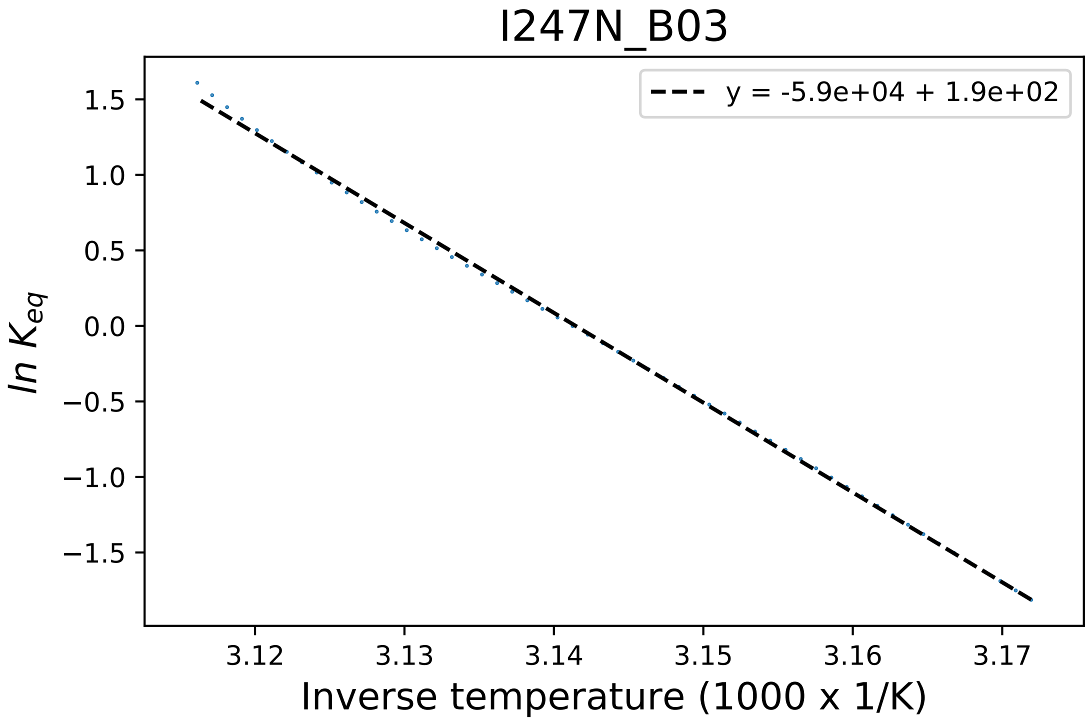

### I247N_B03_fluorescence.pdf

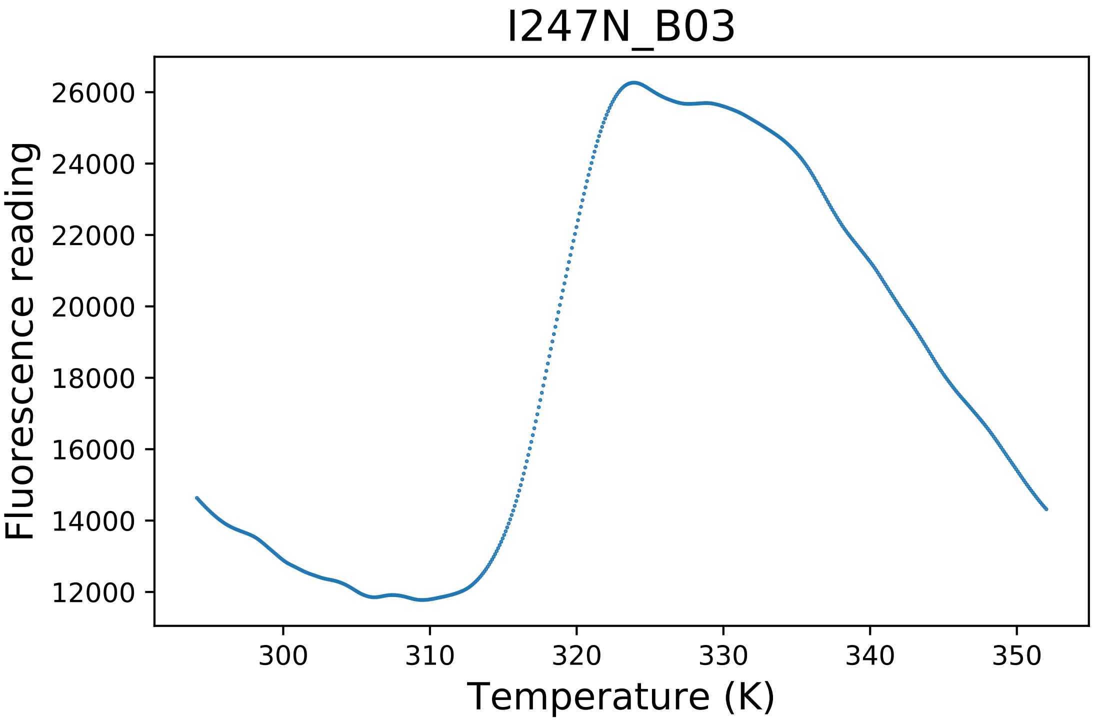

### L174A_E01_derivative.pdf

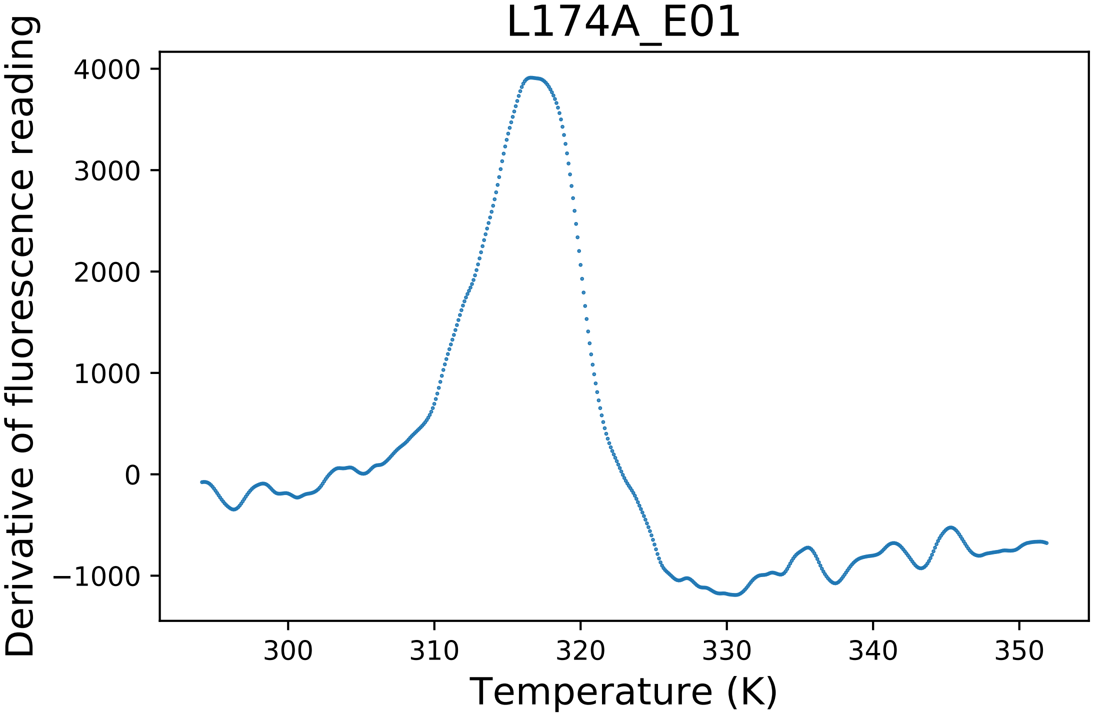

### L174R_G12_derivative.pdf

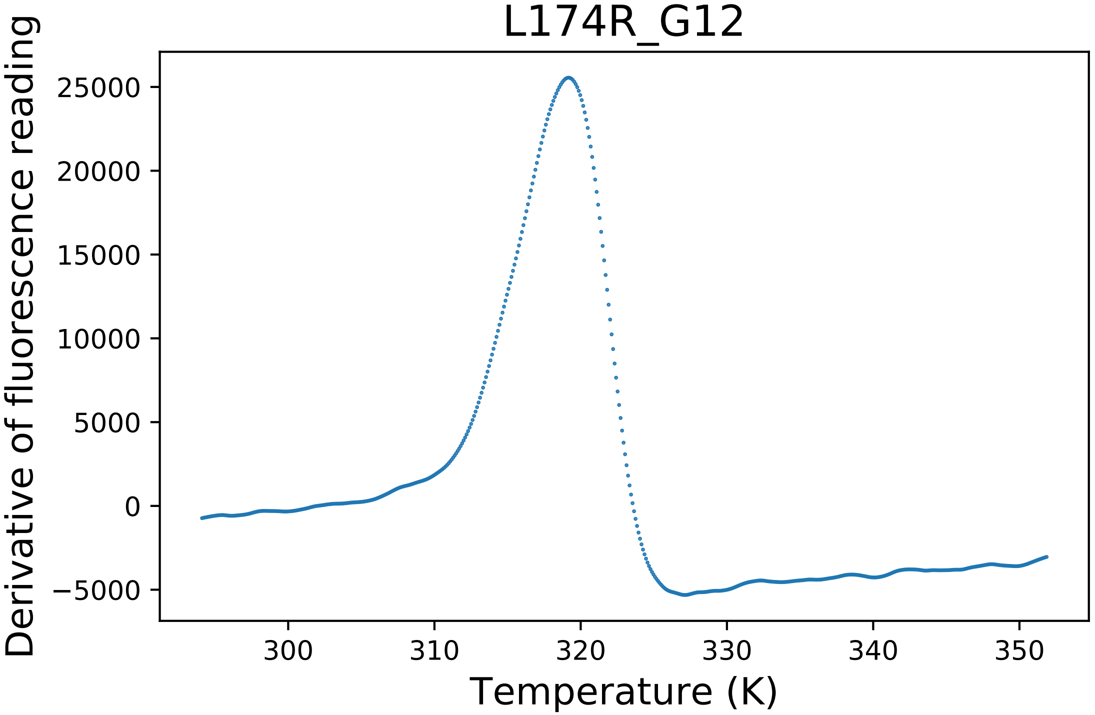

### L174R_G12_fluorescence.pdf

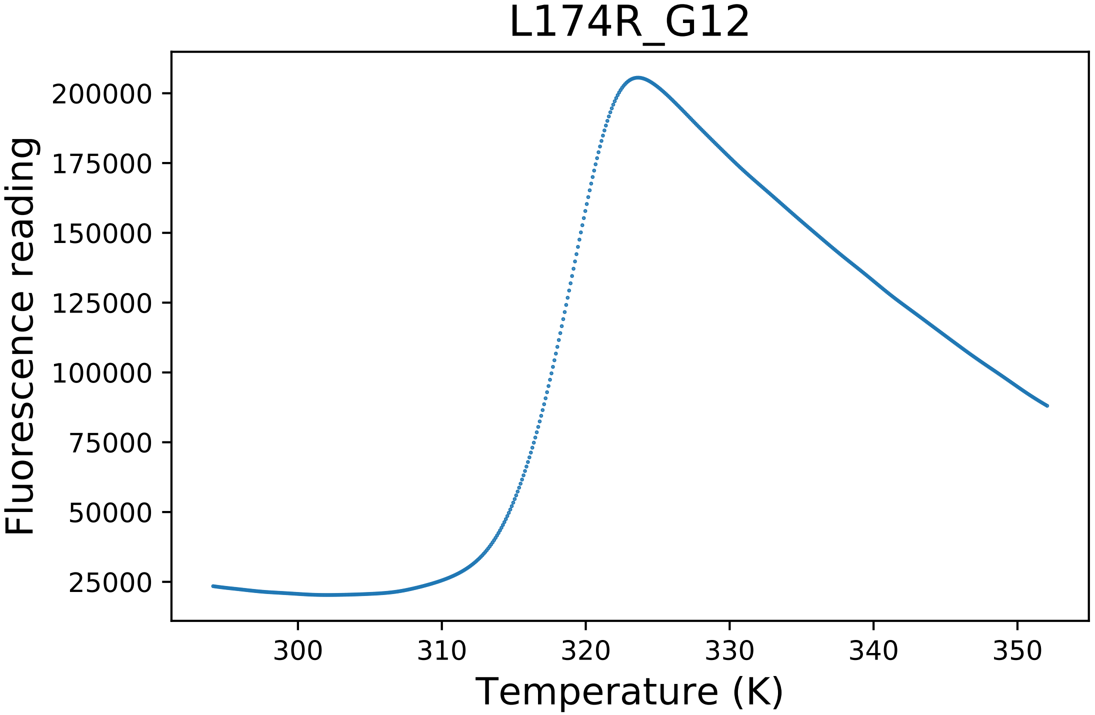

### M224A_E09_fluorescence.pdf

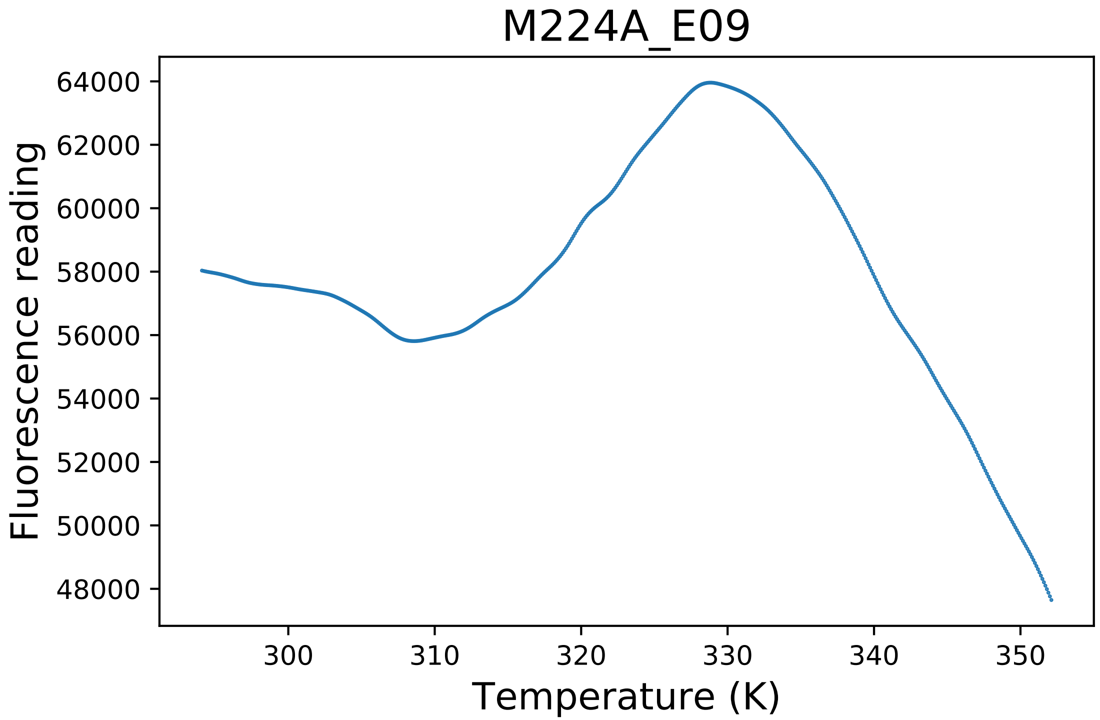

### M224A_E12_fluorescence.pdf

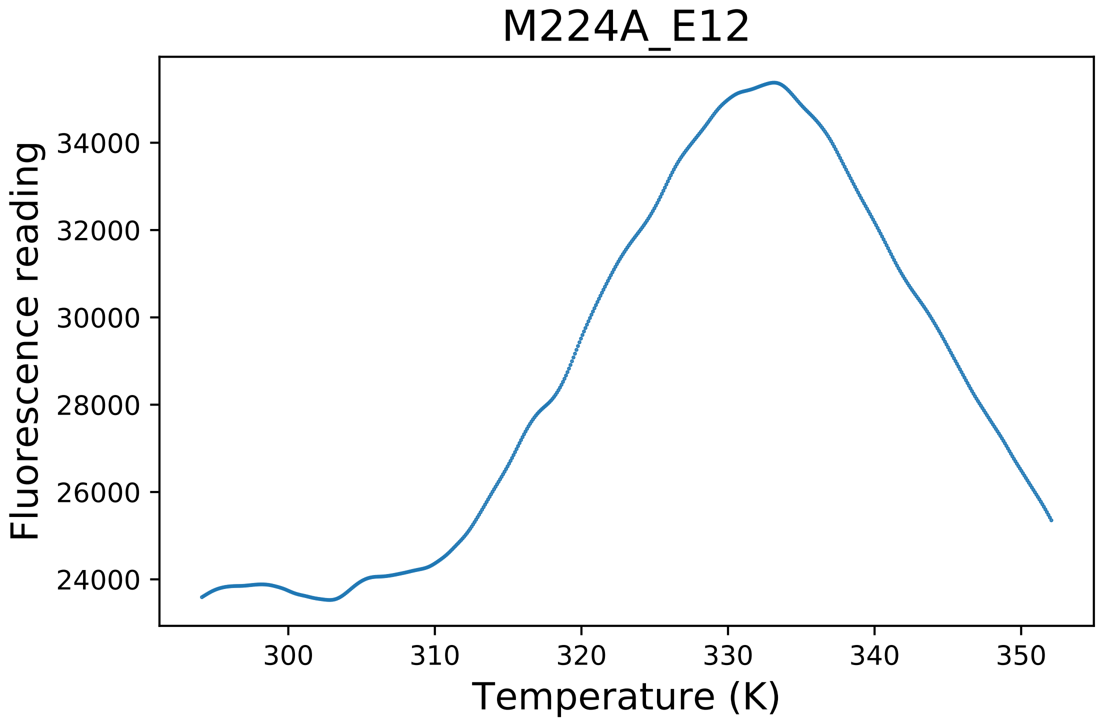

### M326A_D06_derivative.pdf

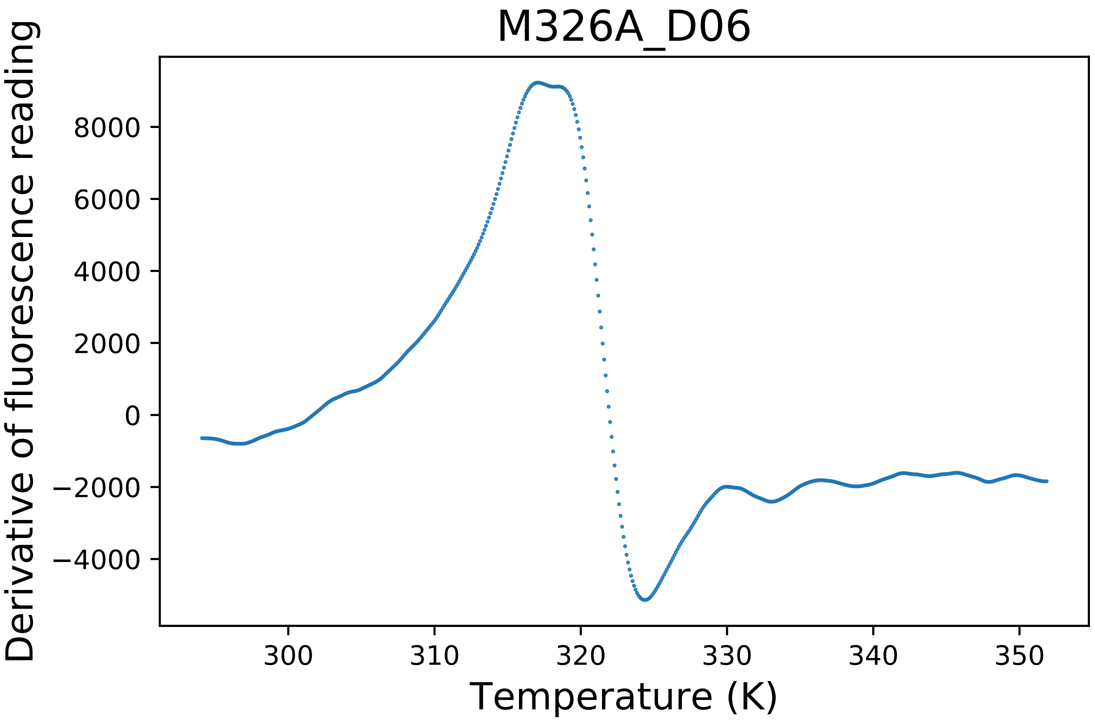

### M326A_D07_derivative.pdf

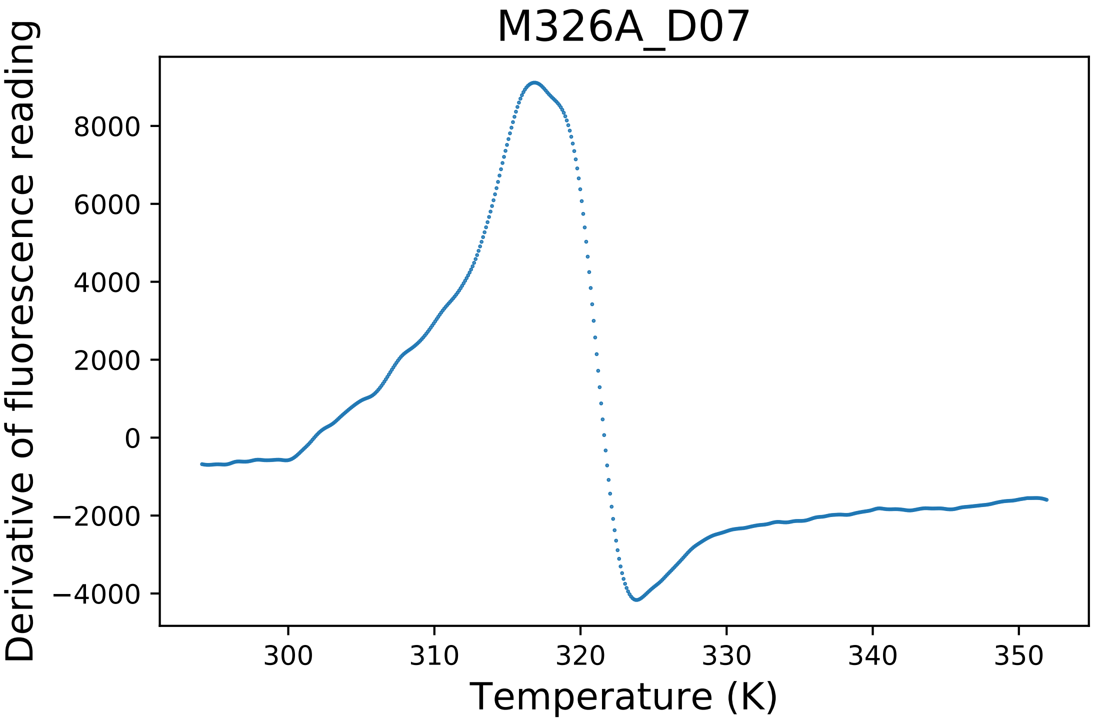

### M326A_D08_fluorescence.pdf

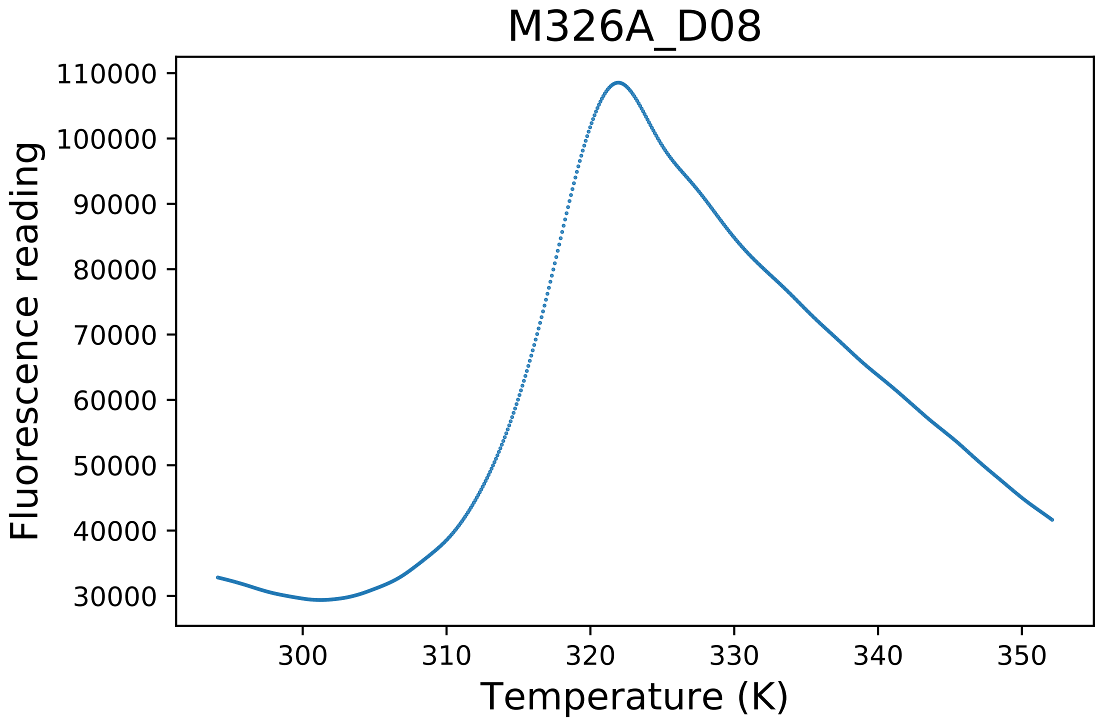

### N166C_F01_derivative.pdf

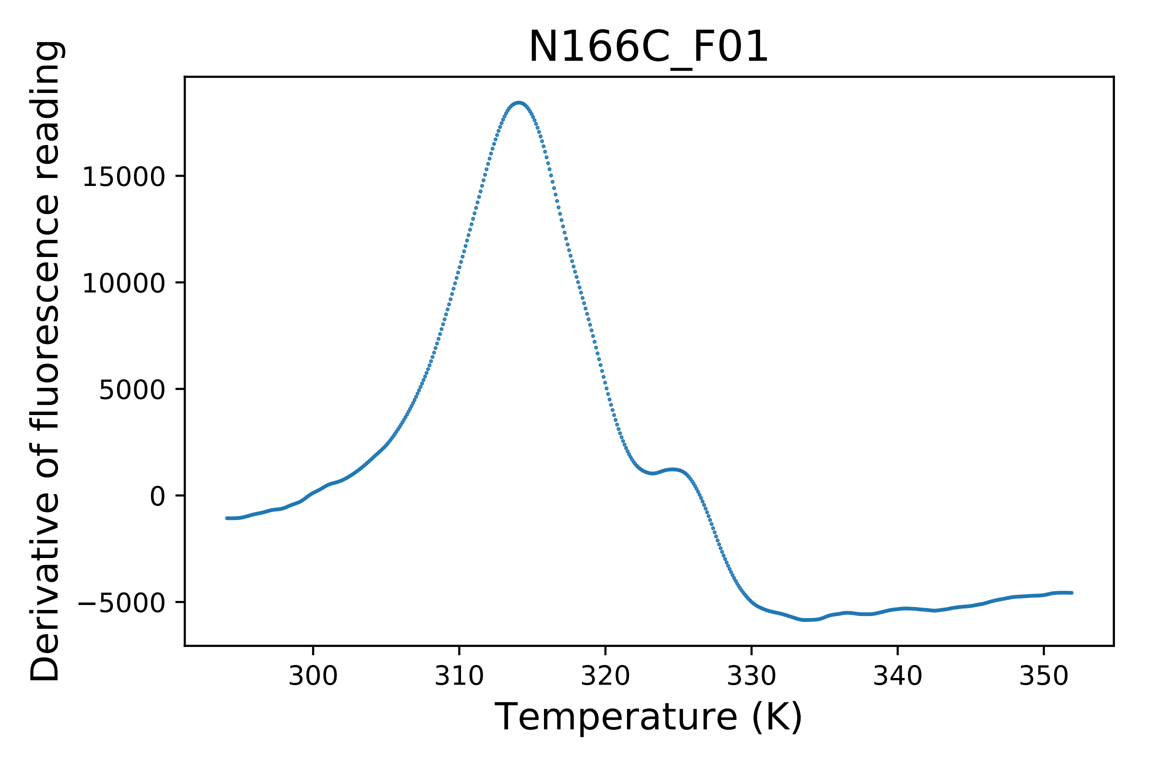

### N166C_F03_fluorescence.pdf

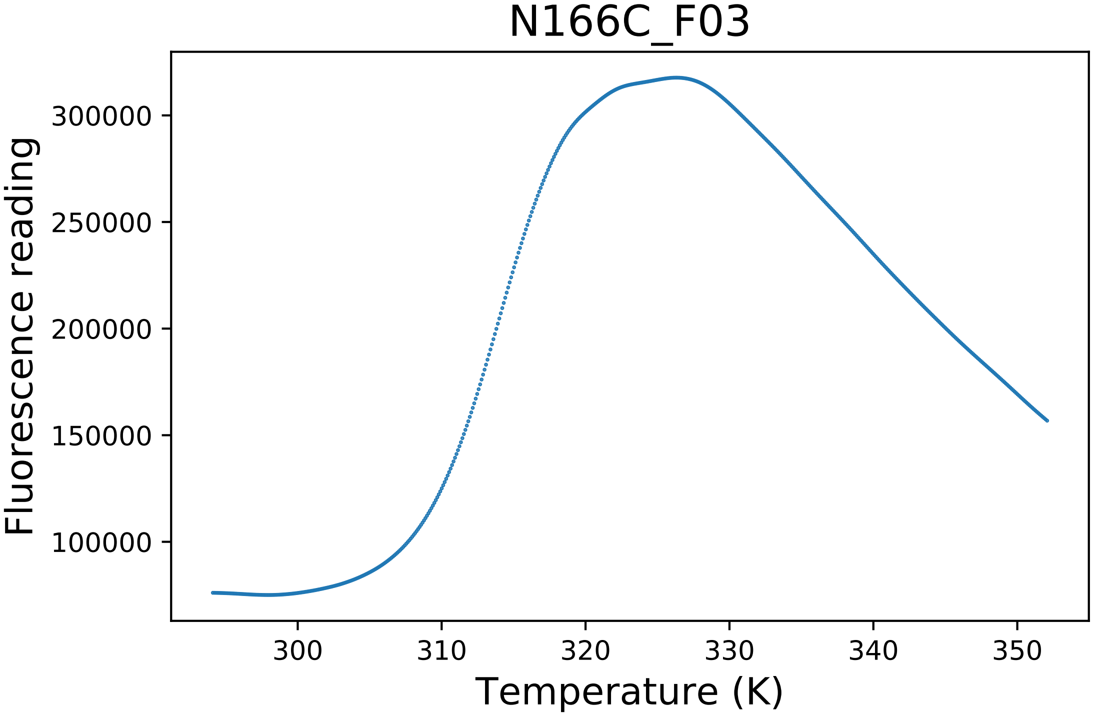

### N223A_F07_fluorescence.pdf

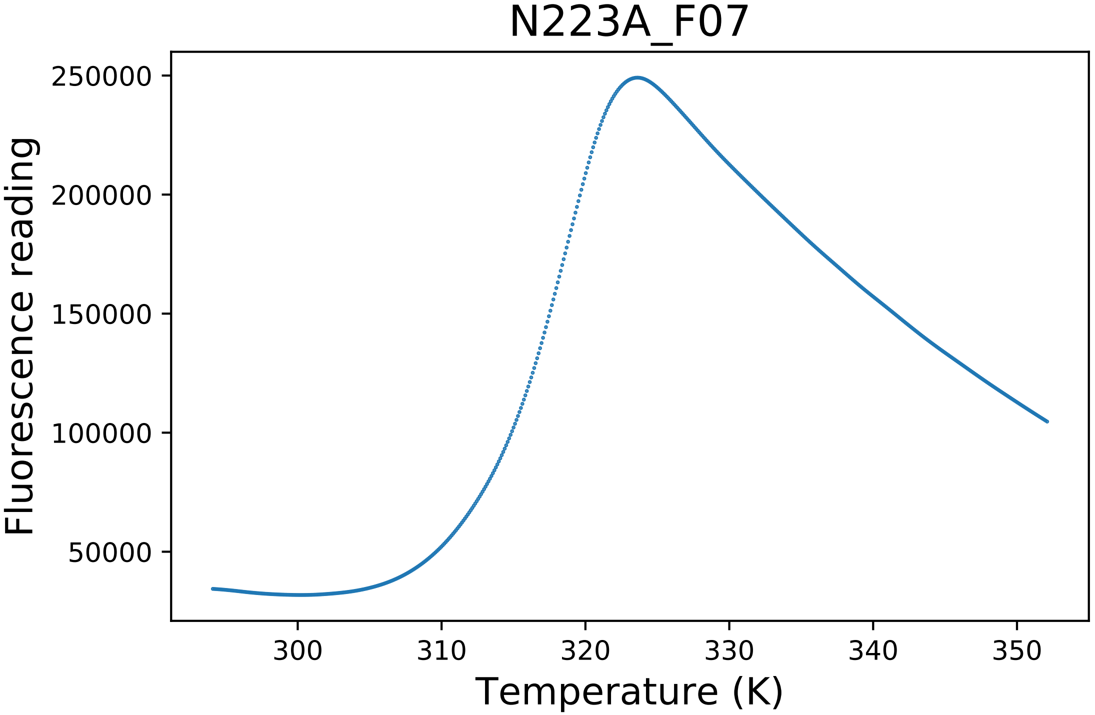

### N223A_F08_derivative.pdf

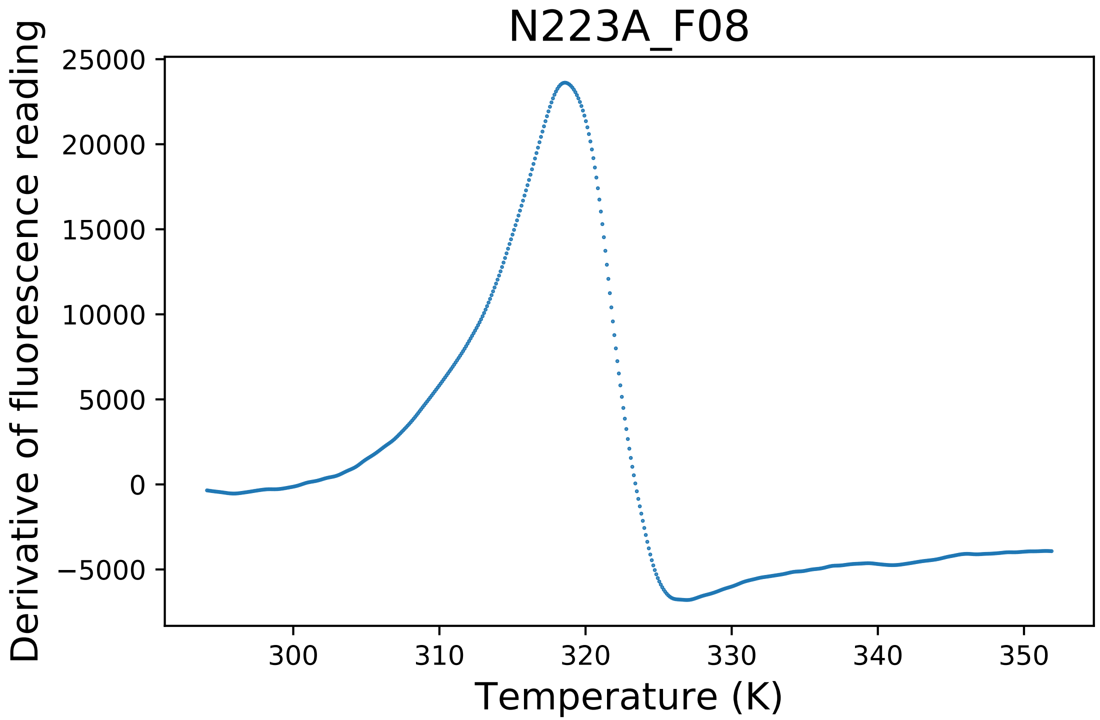

### N223A_F08_fluorescence.pdf

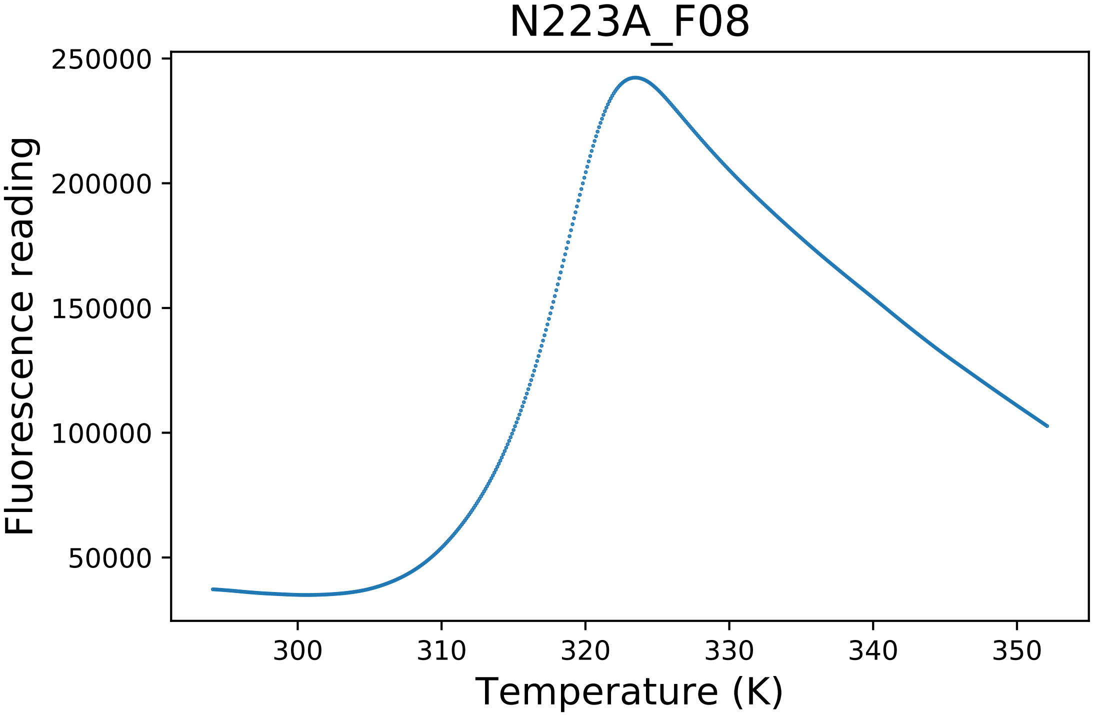

### N223G_C01_derivative.pdf

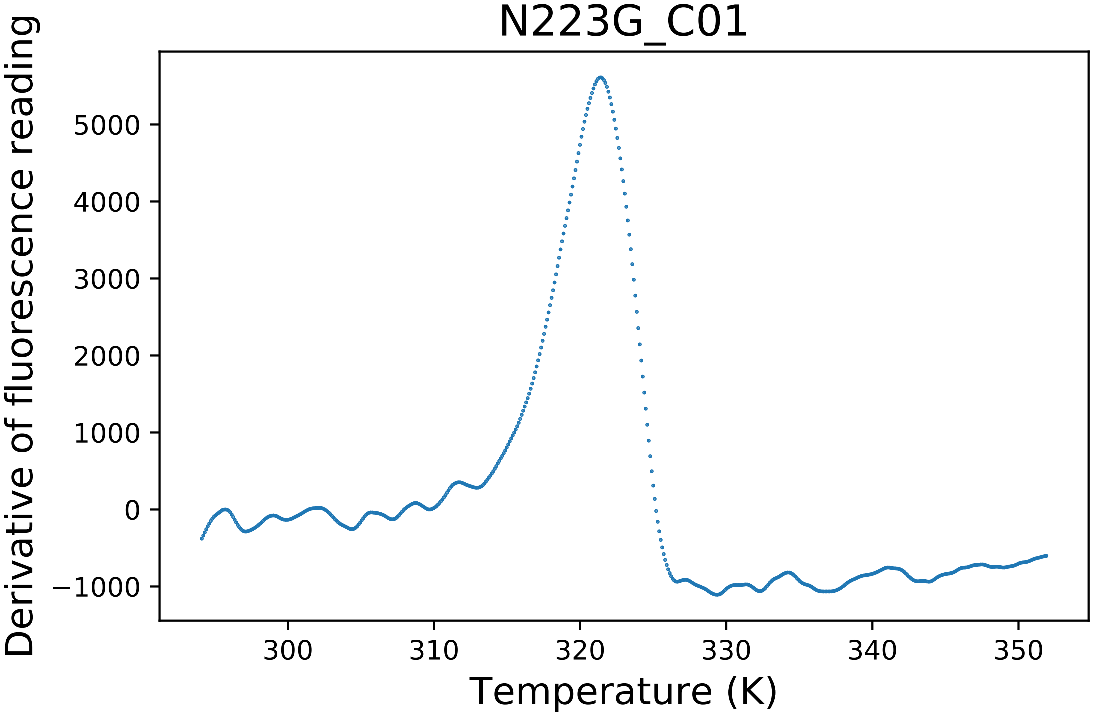

### N223G_C01_dg.pdf

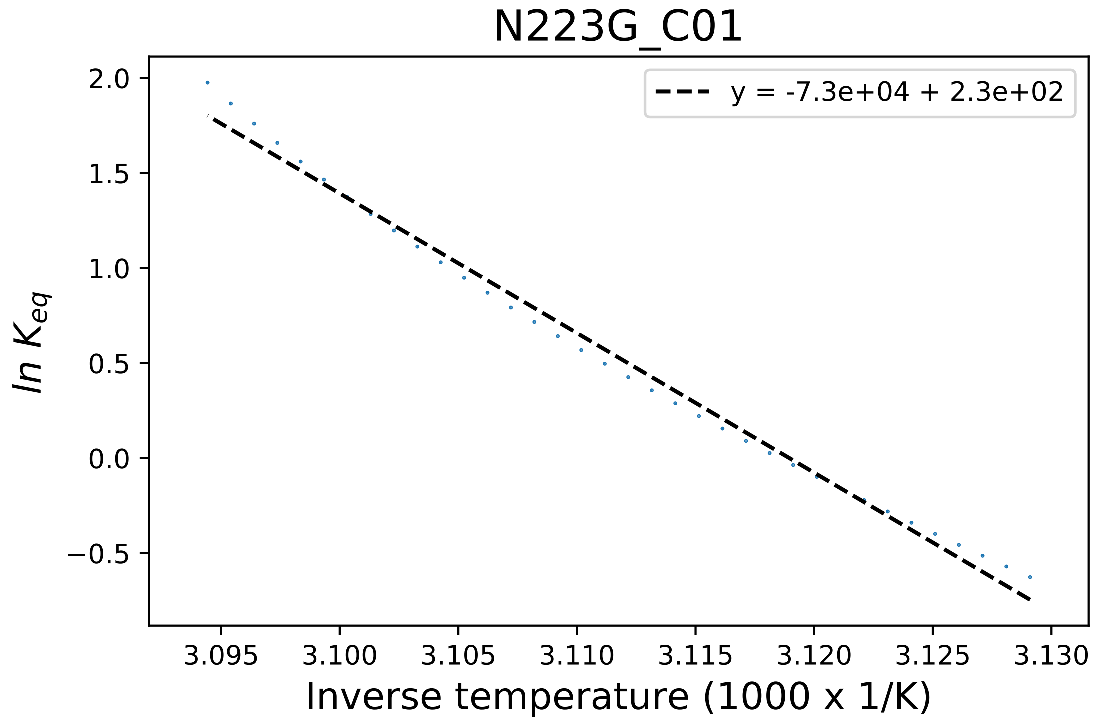

### N223R_G01_dg.pdf

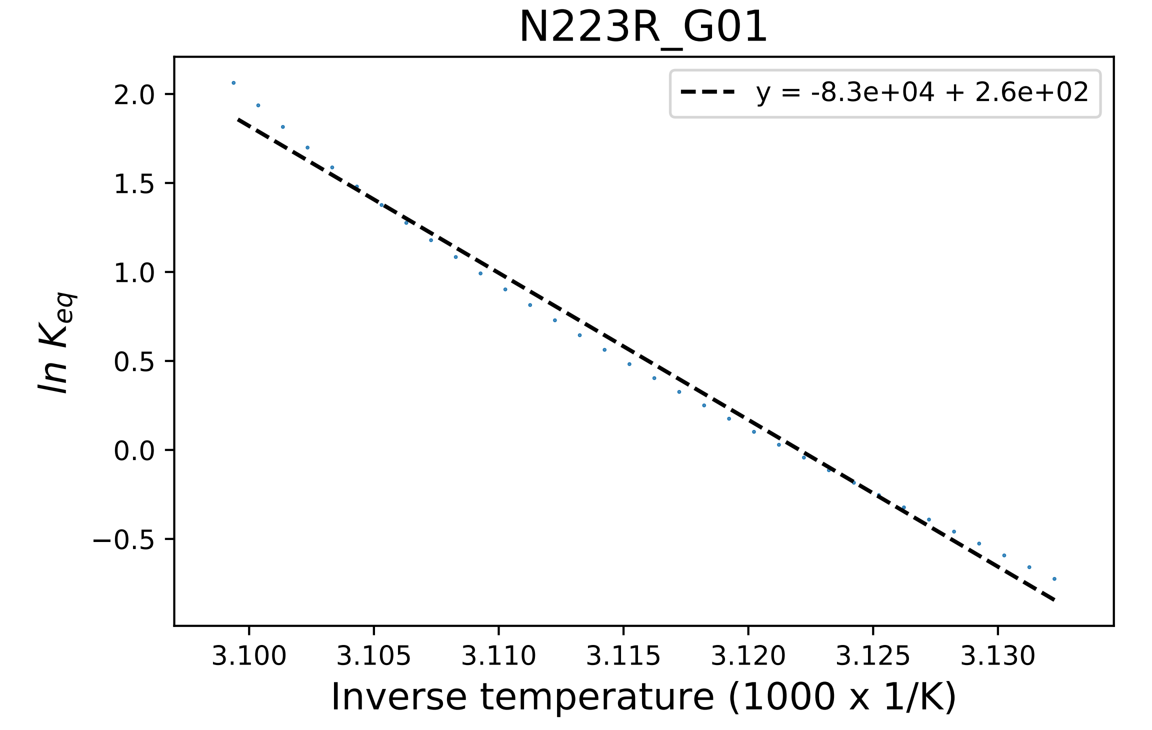

### N223Y_B01_dg.pdf

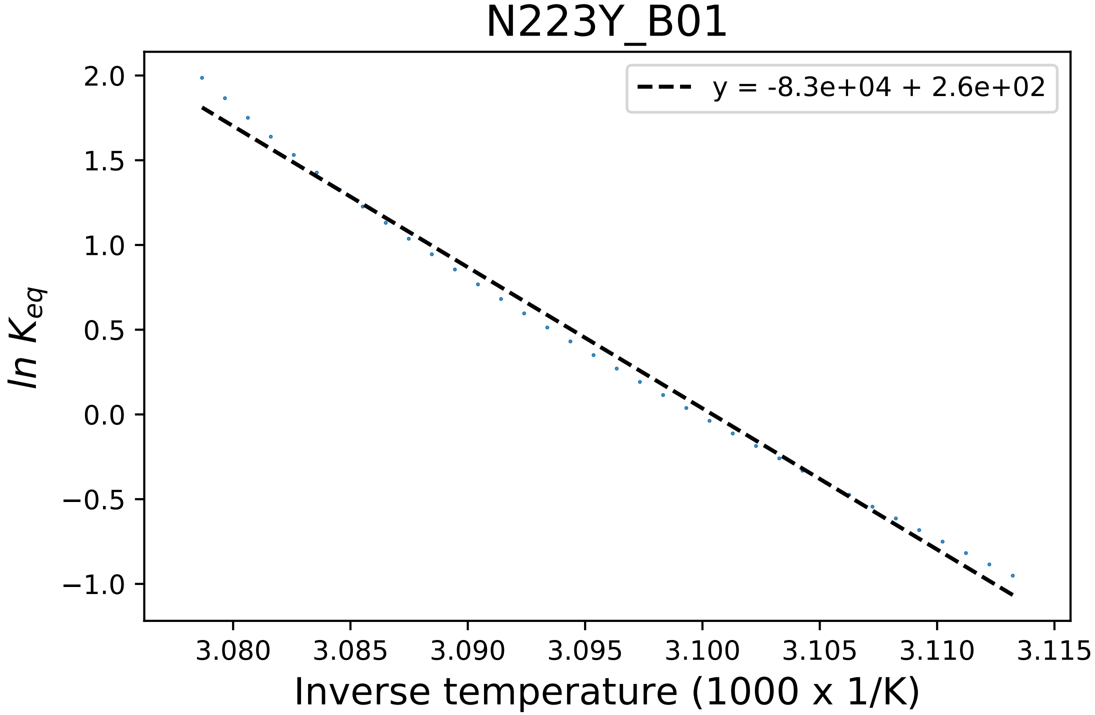

### N223Y_B03_dg.pdf

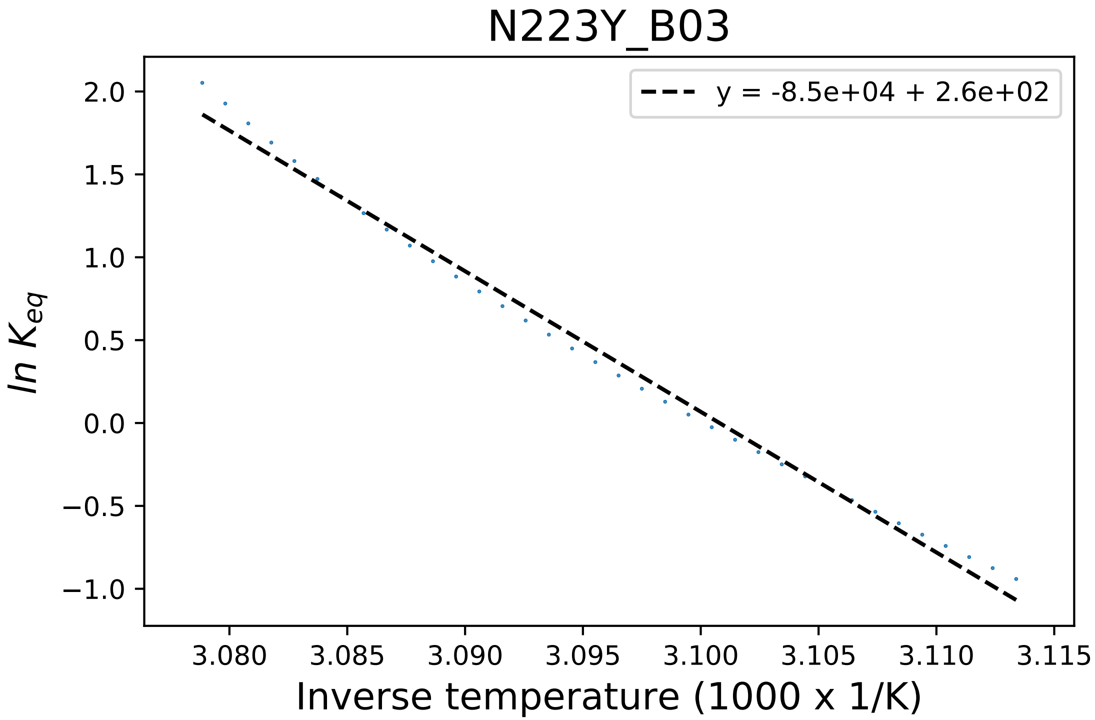

### N296A_B05_derivative.pdf

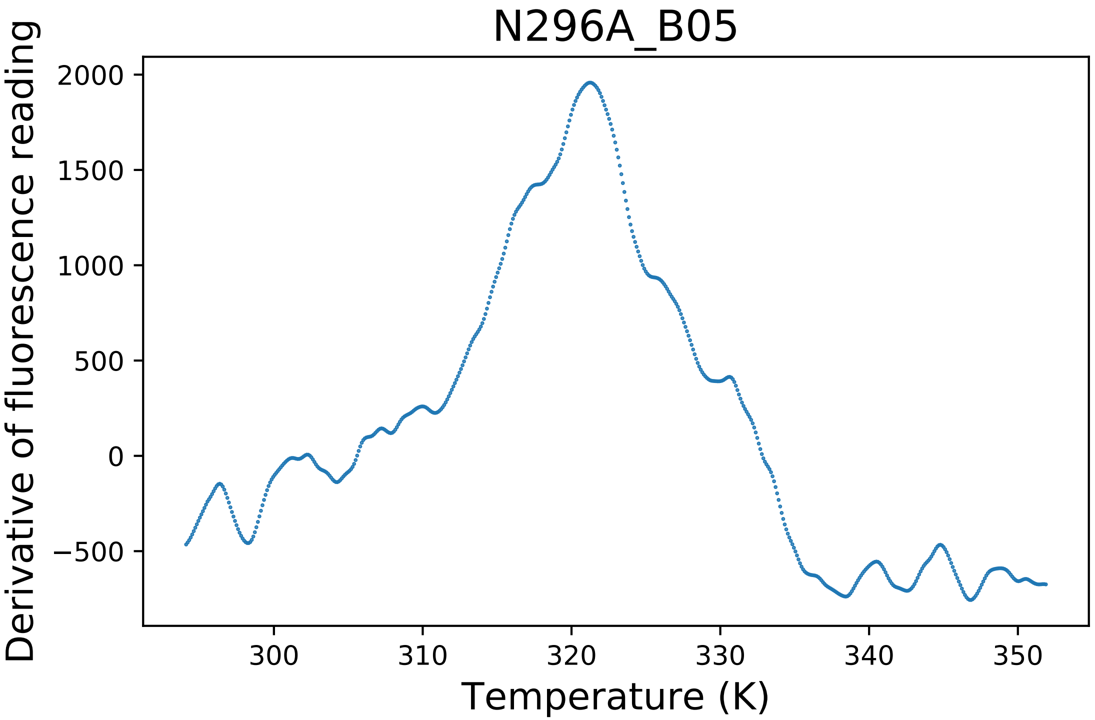

### N296A_B08_fluorescence.pdf

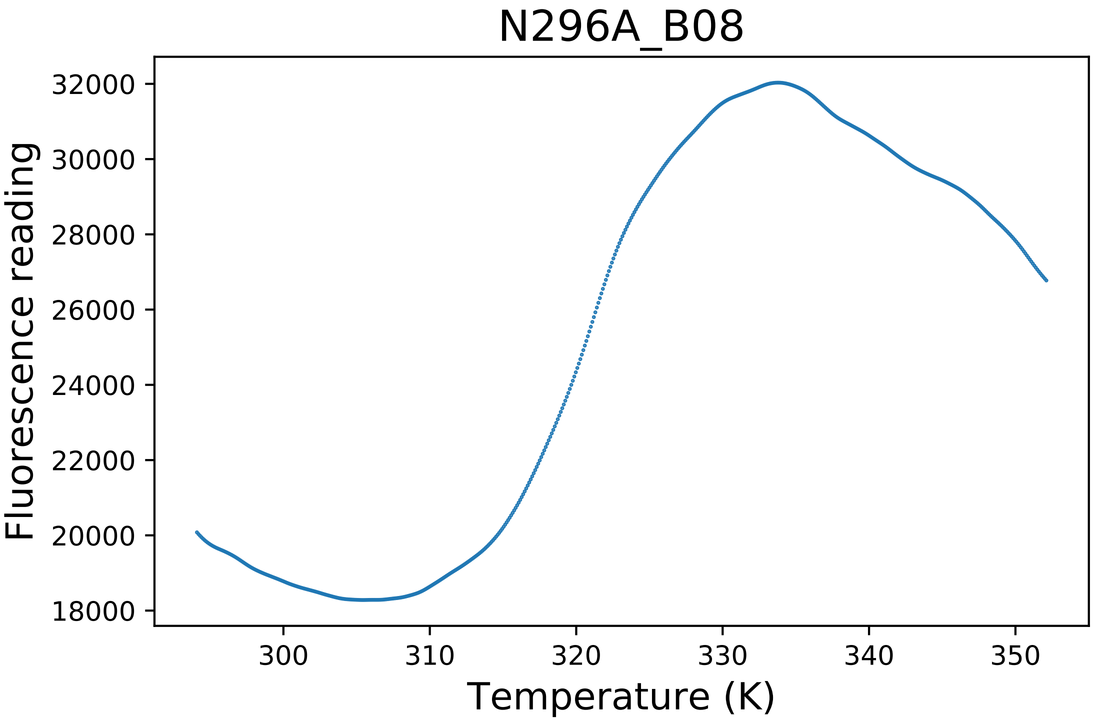

### N296C_B05_fluorescence.pdf

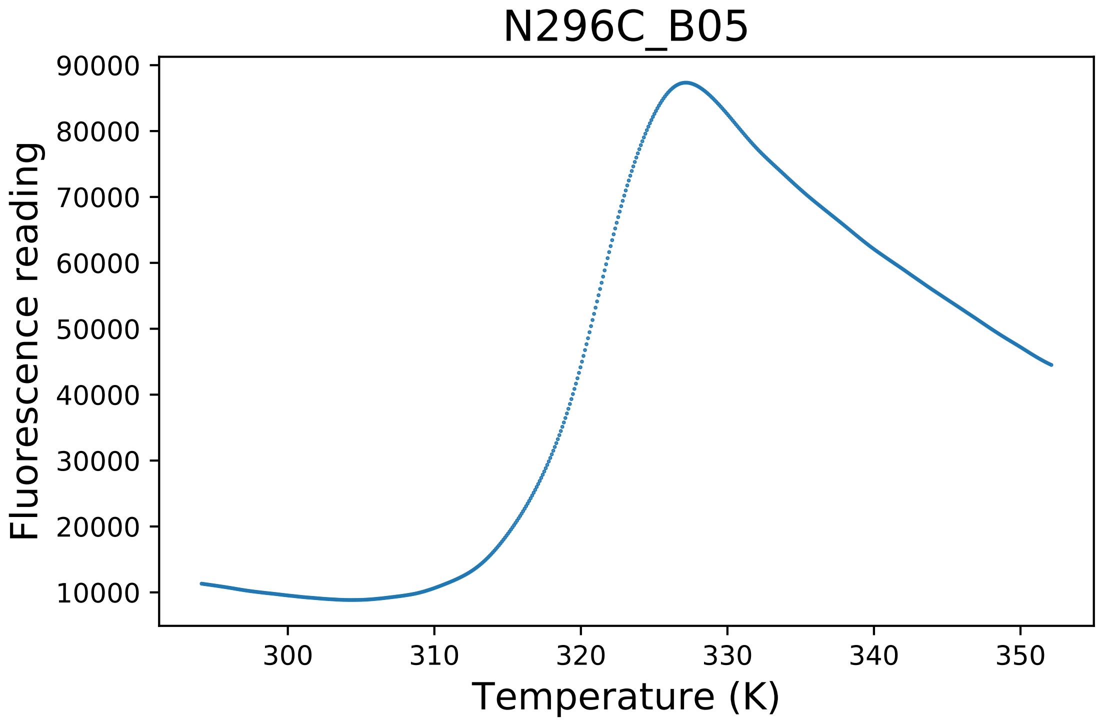

### N296C_B08_dg.pdf

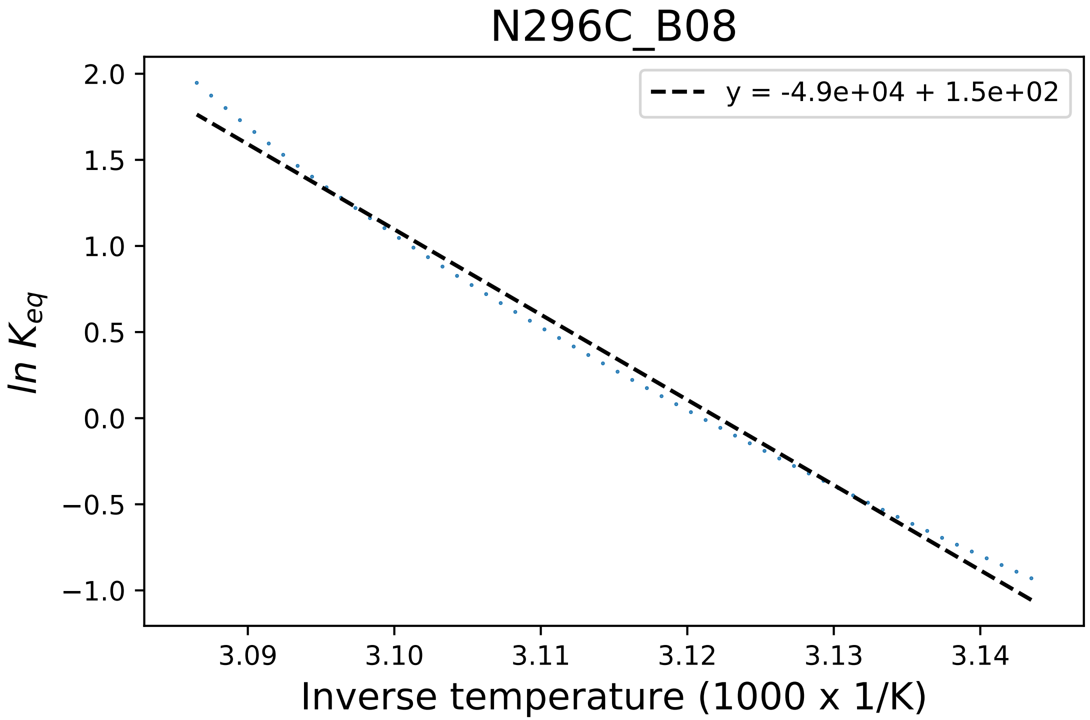

### N407C_C04_derivative.pdf

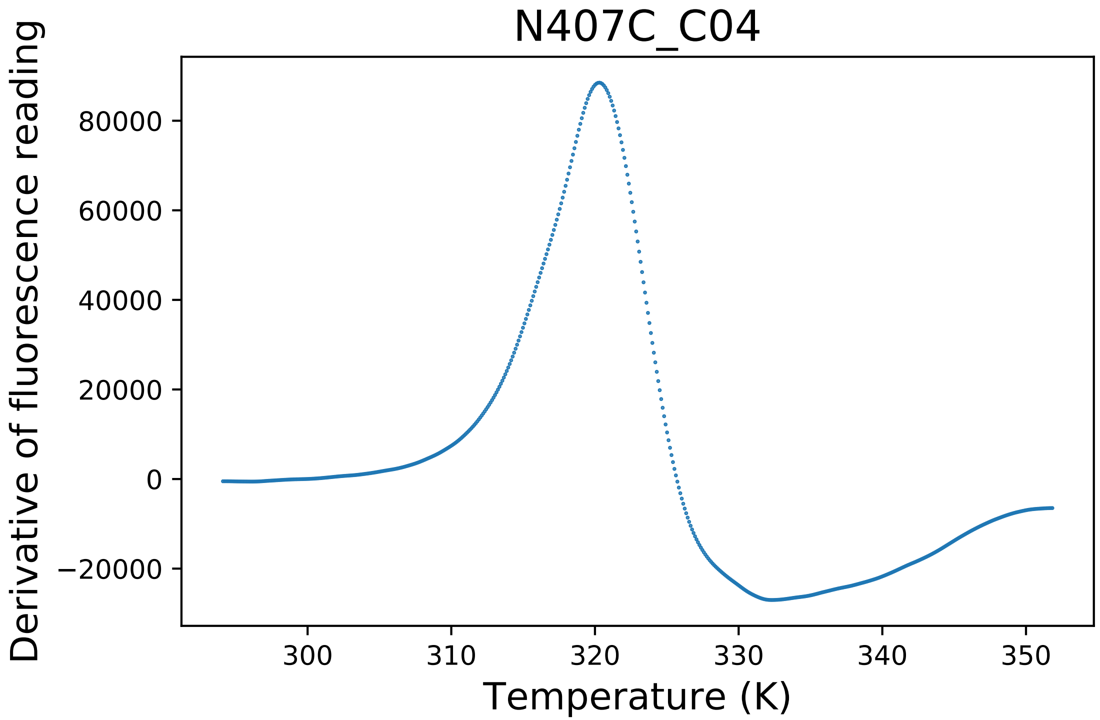

### Q22S_B06_dg.pdf

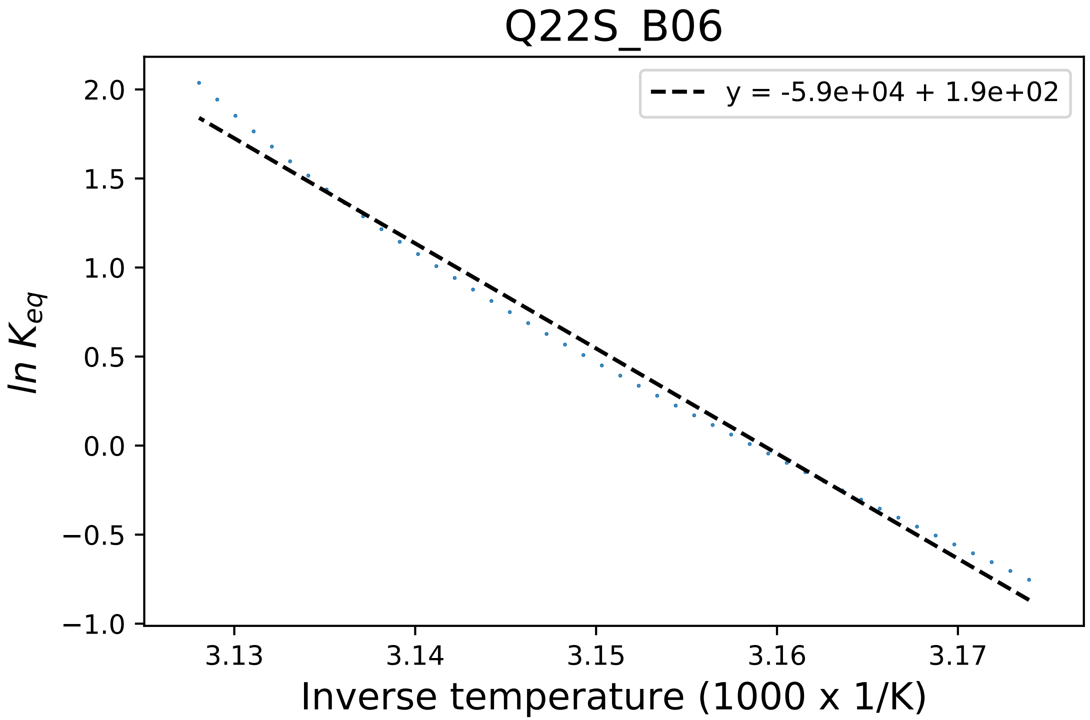

### Q22S_B06_fluorescence.pdf

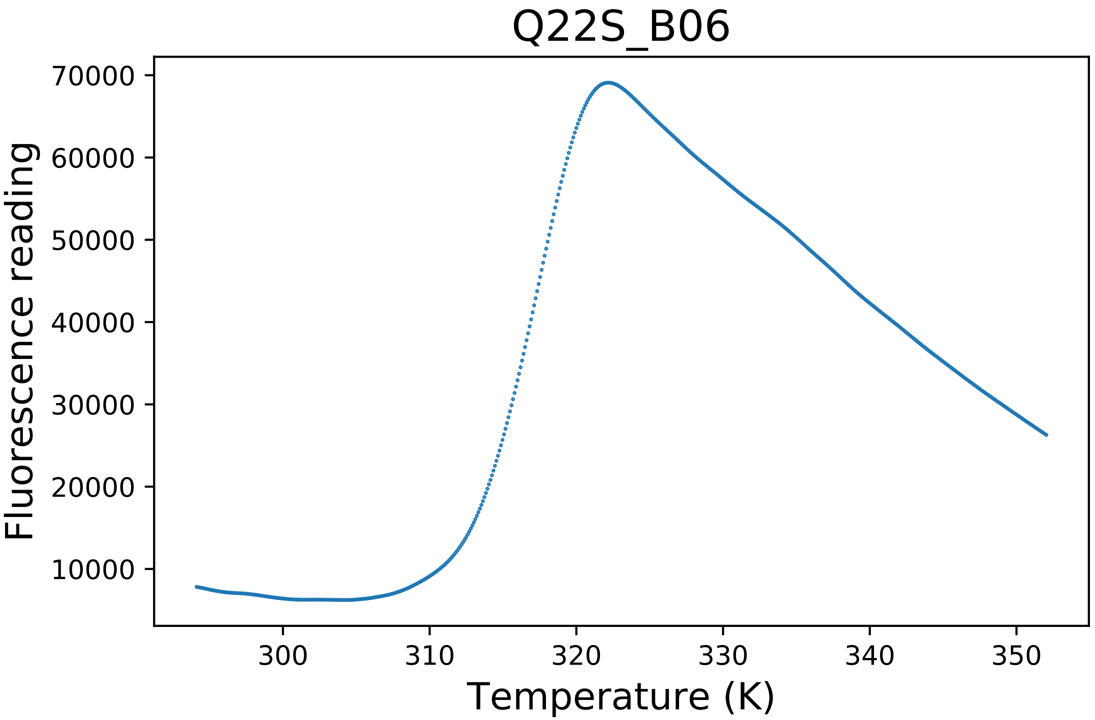

### R243A_A02_dg.pdf

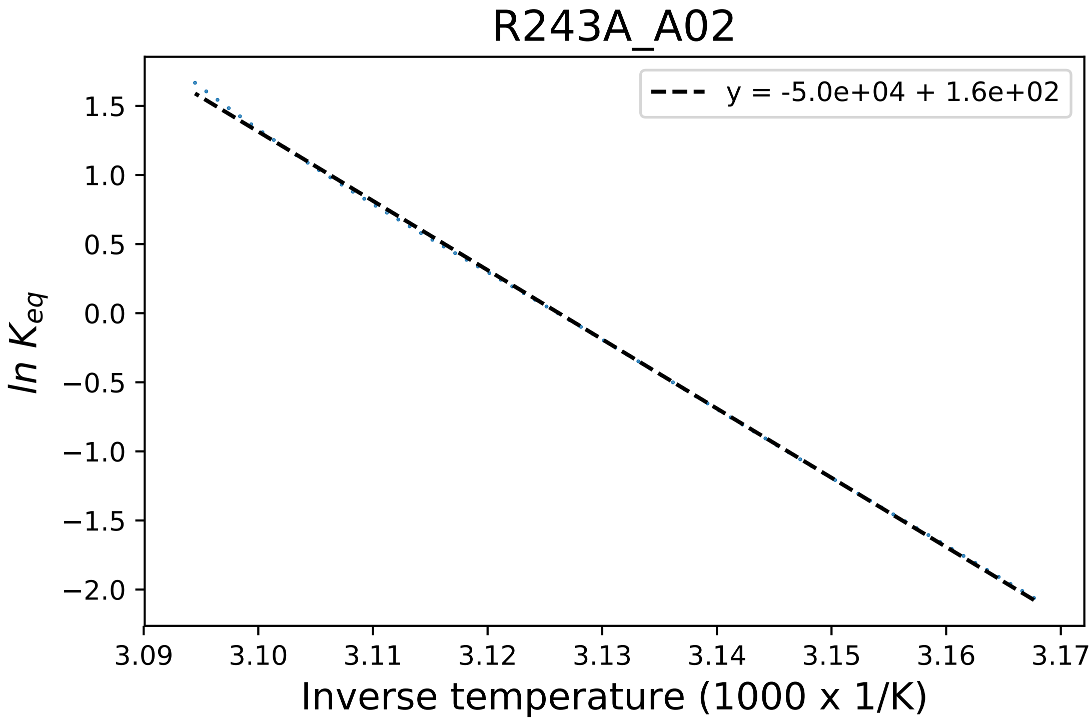

### R243D_E06_fluorescence.pdf

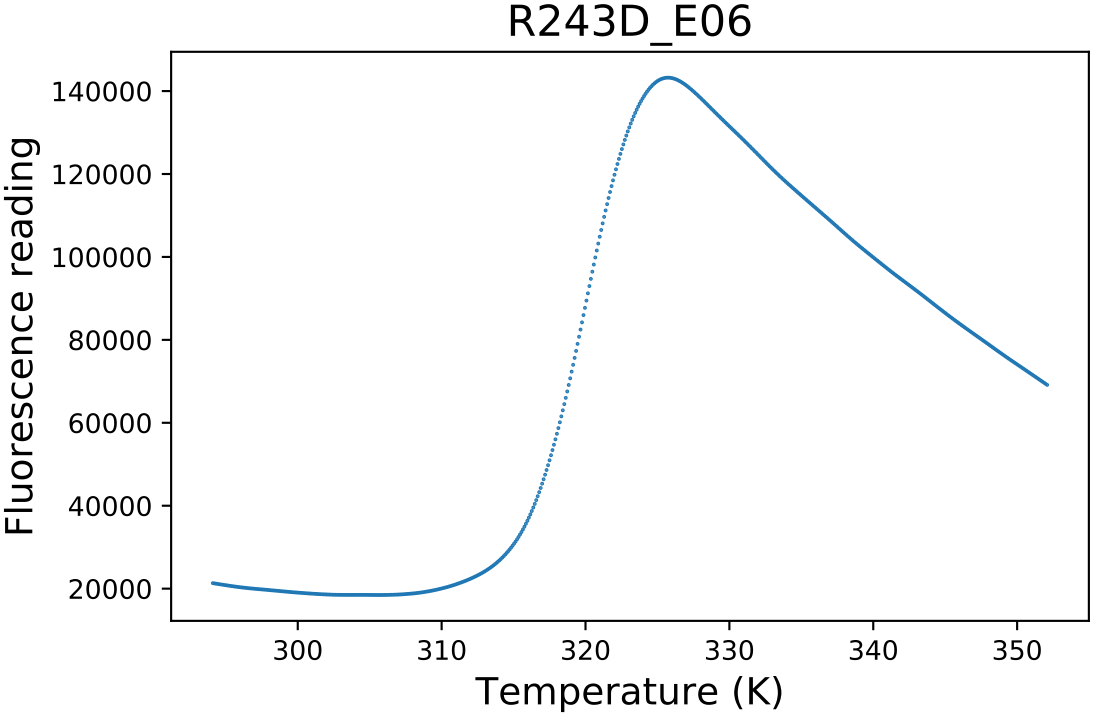

### R243D_E07_dg.pdf

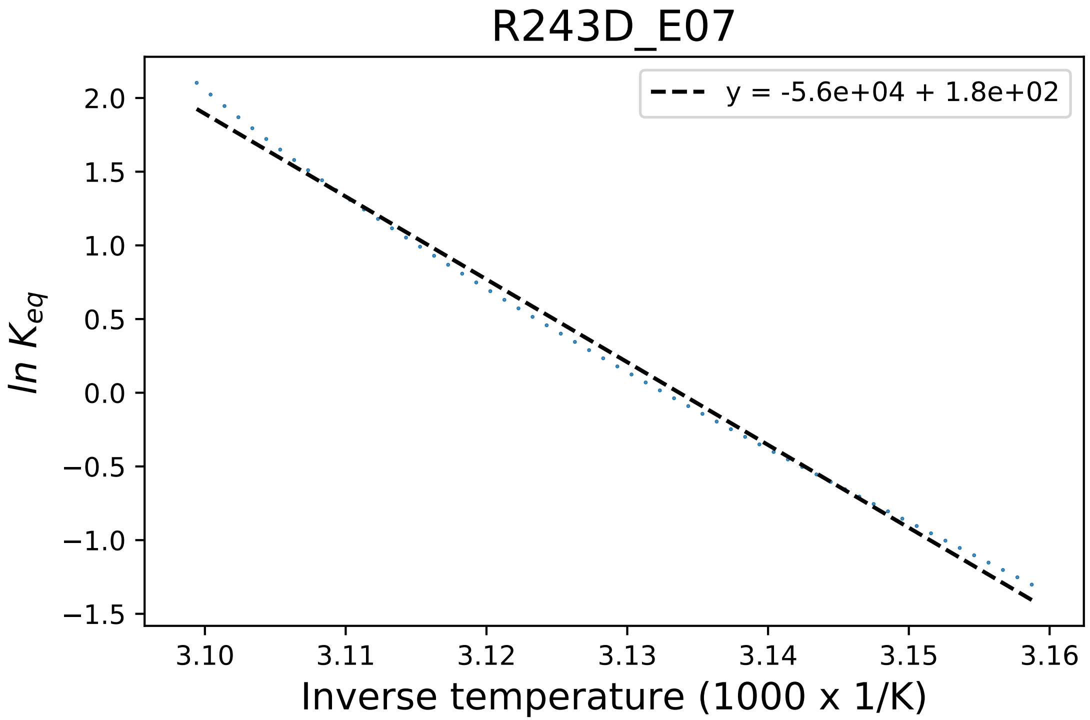

### R243K_C05_dg.pdf

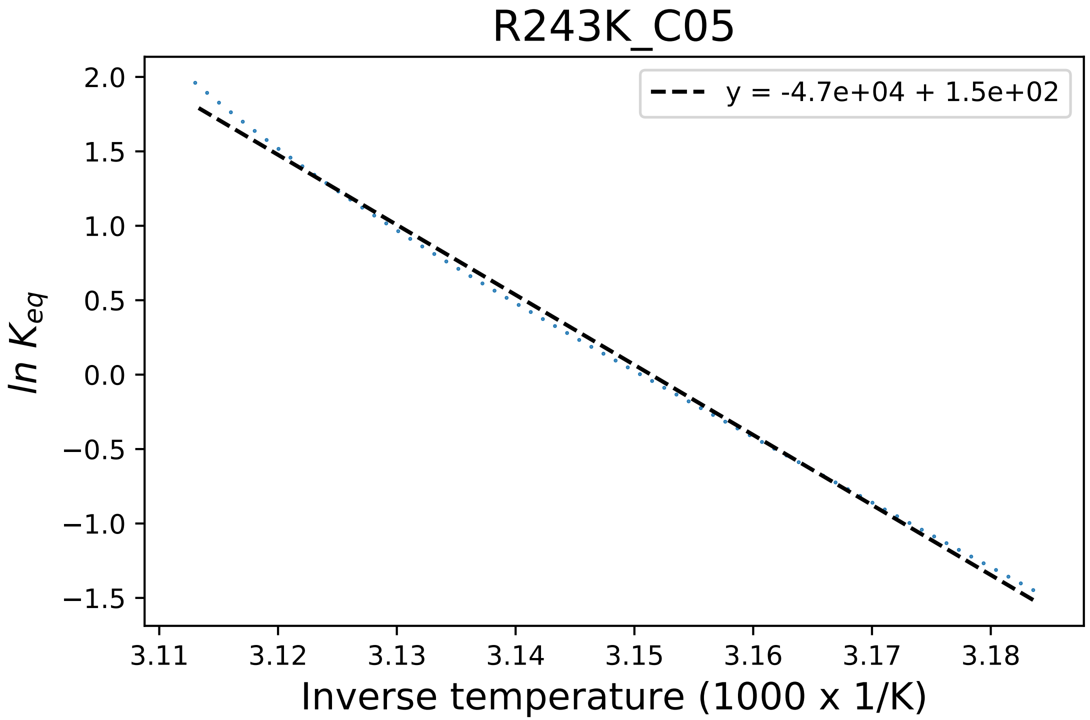

### R243K_C07_dg.pdf

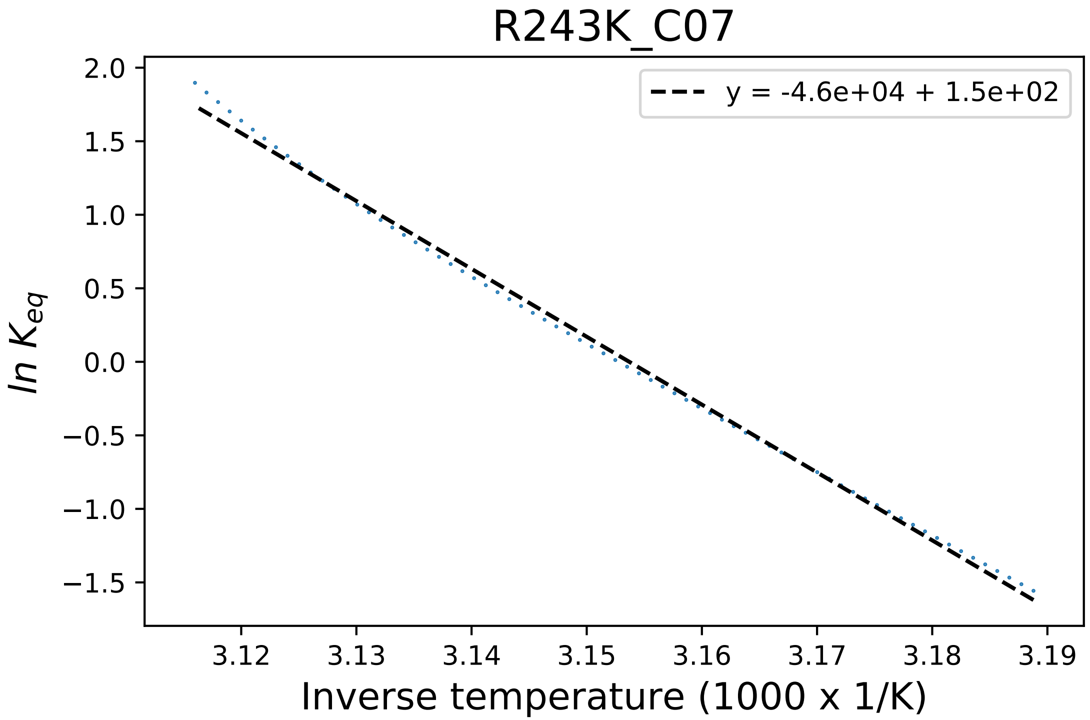

### R243K_C08_fluorescence.pdf

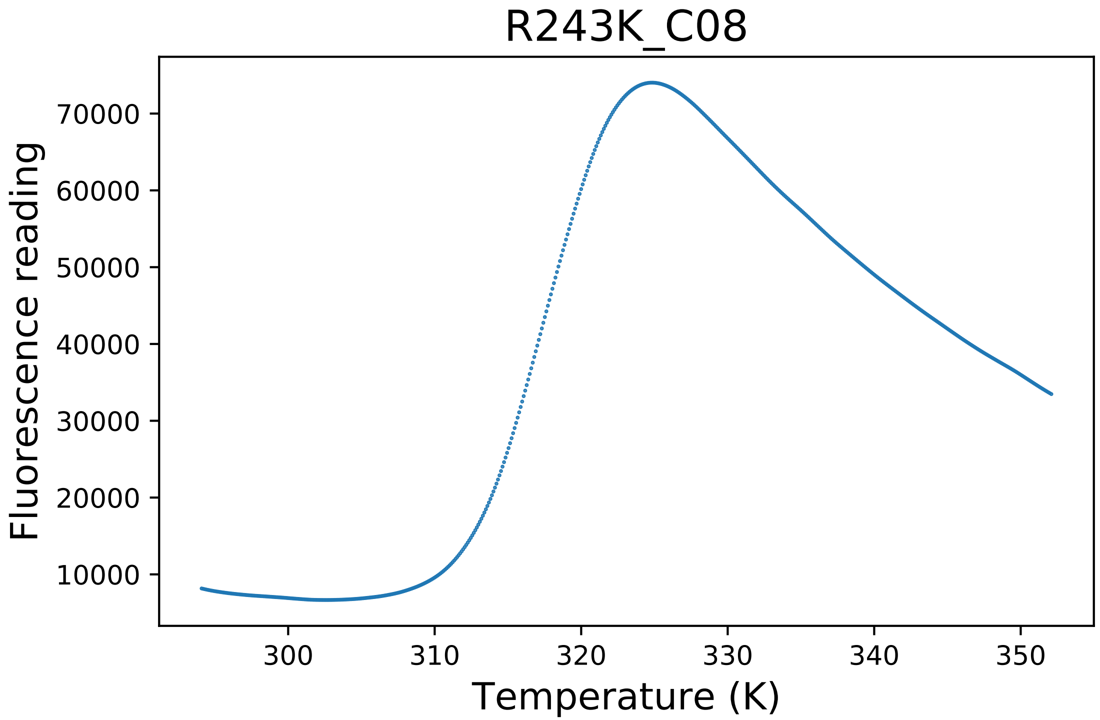

### S17A_G12_dg.pdf

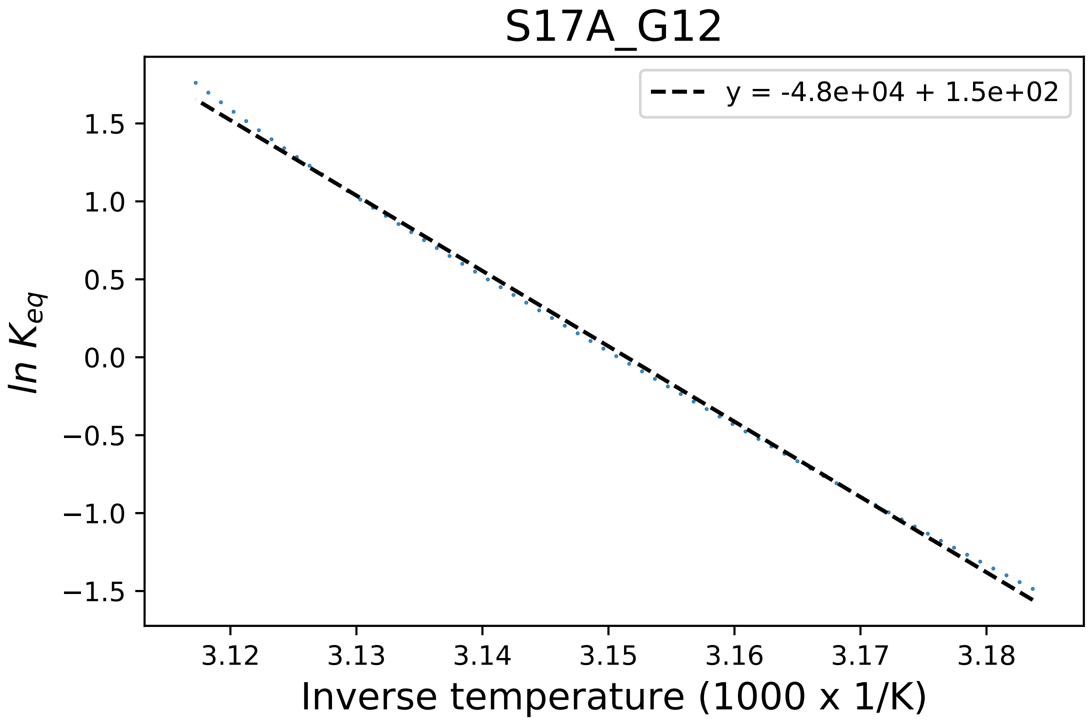

### S19A_A01_derivative.pdf

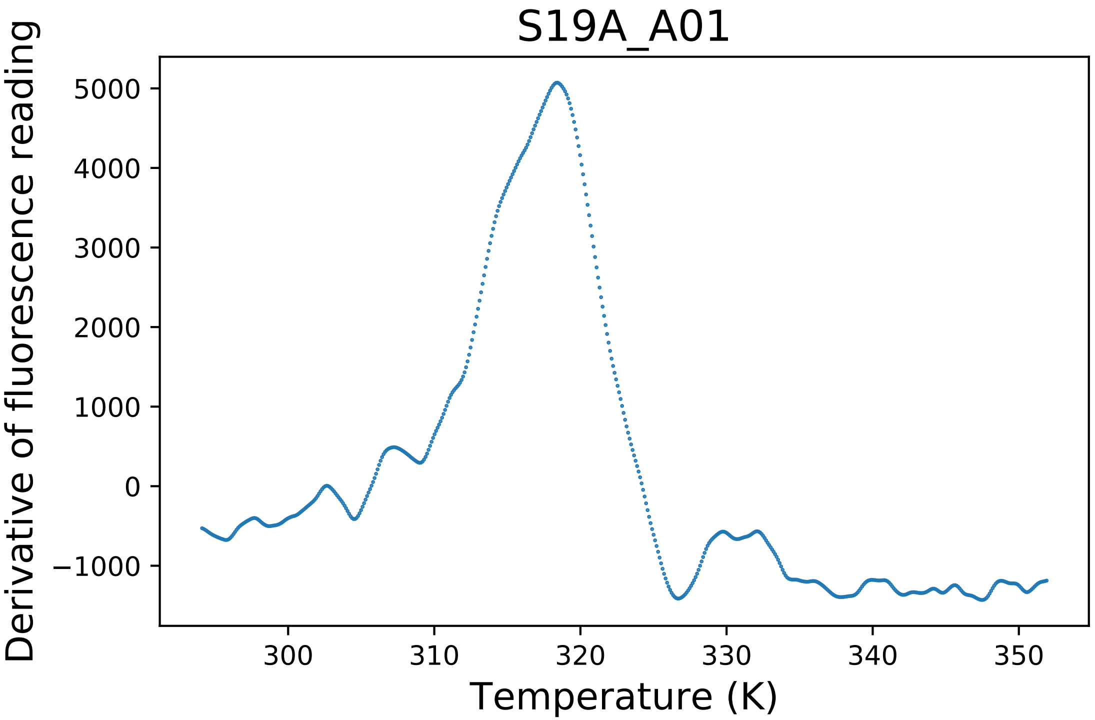

### S334A_B02_derivative.pdf

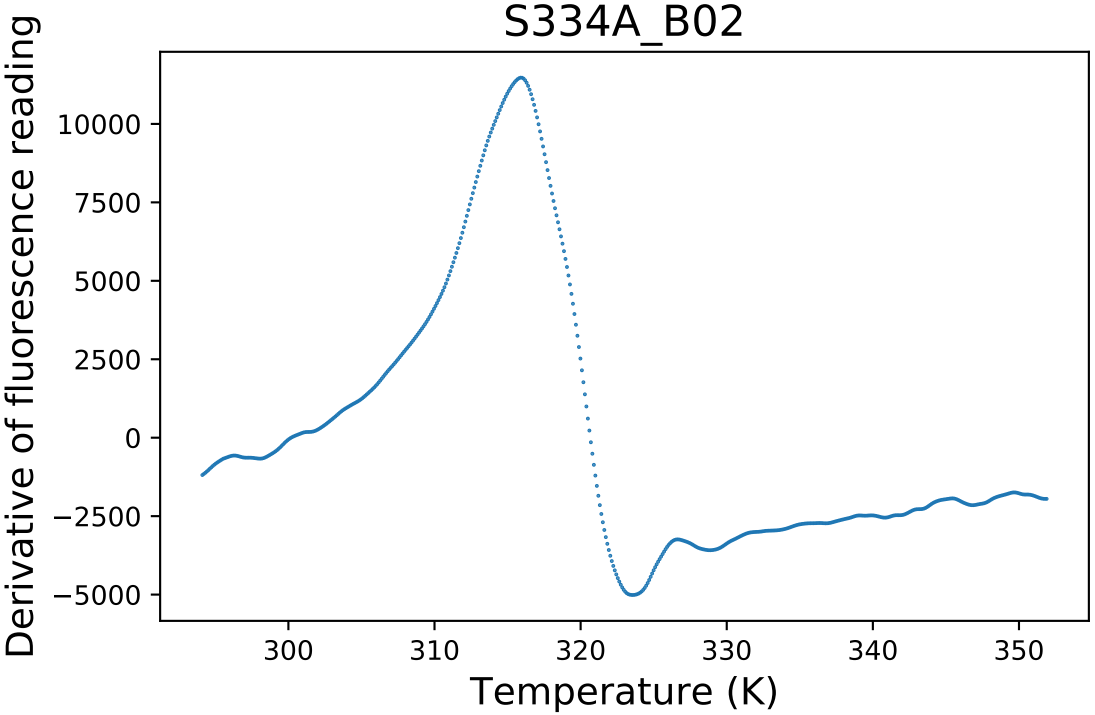

### S334A_B03_derivative.pdf

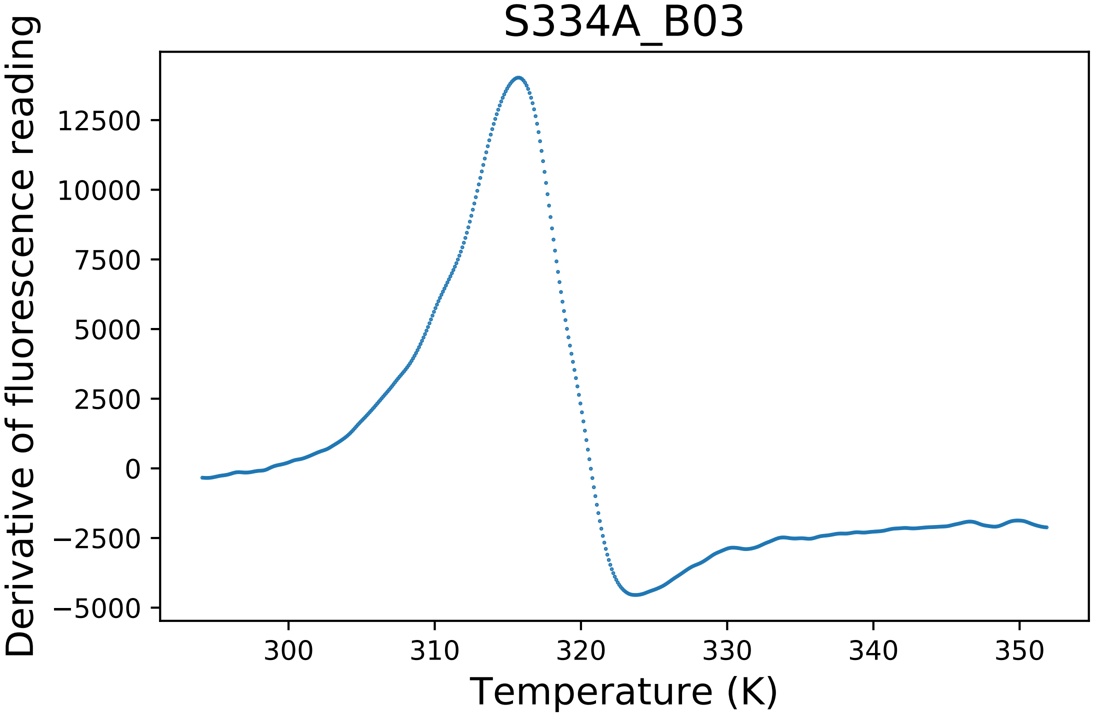

### S334A_B03_dg.pdf

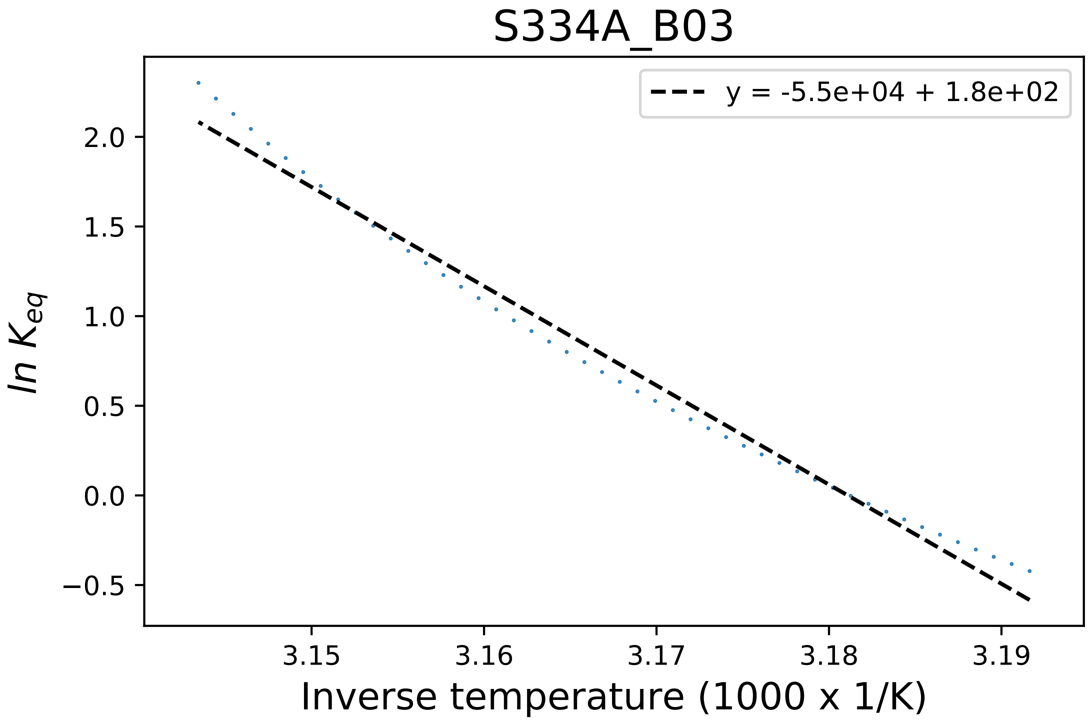

### T18A_E01_fluorescence.pdf

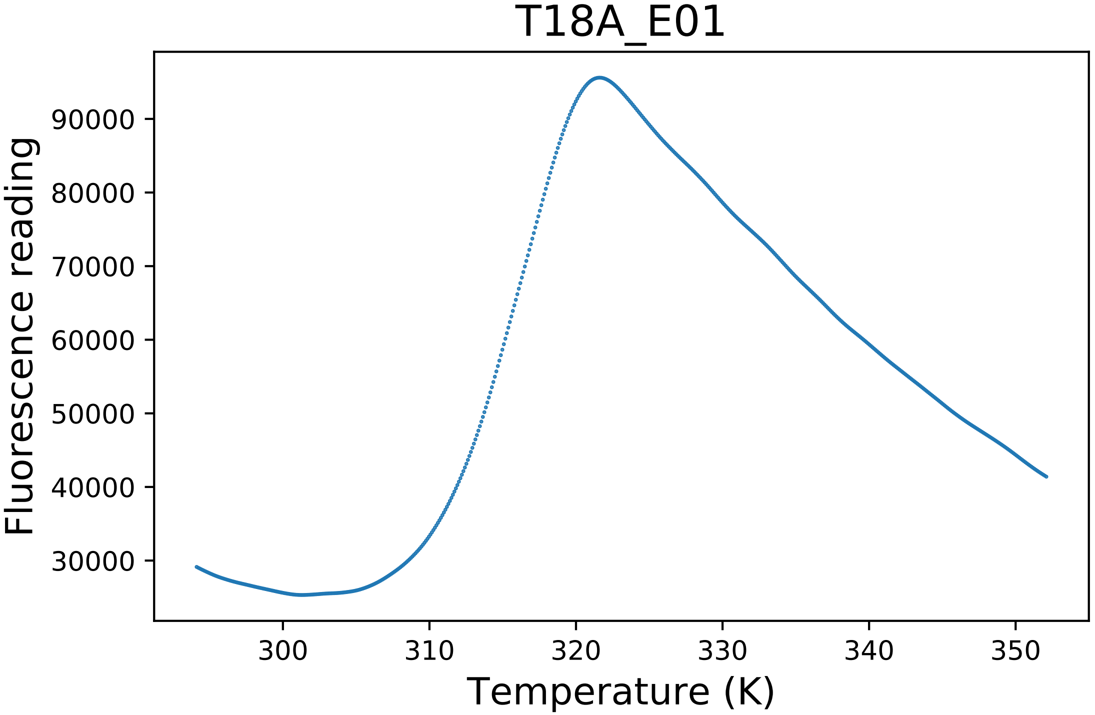

### T355A_B05_fluorescence.pdf

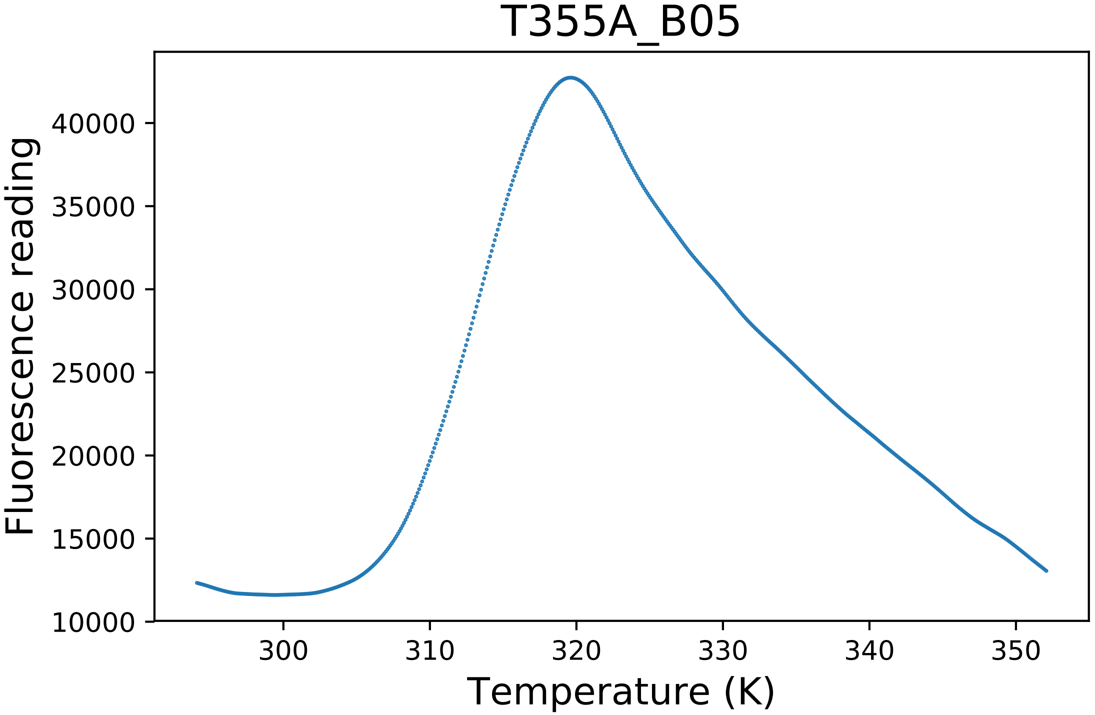

### W123F_G06_dg.pdf

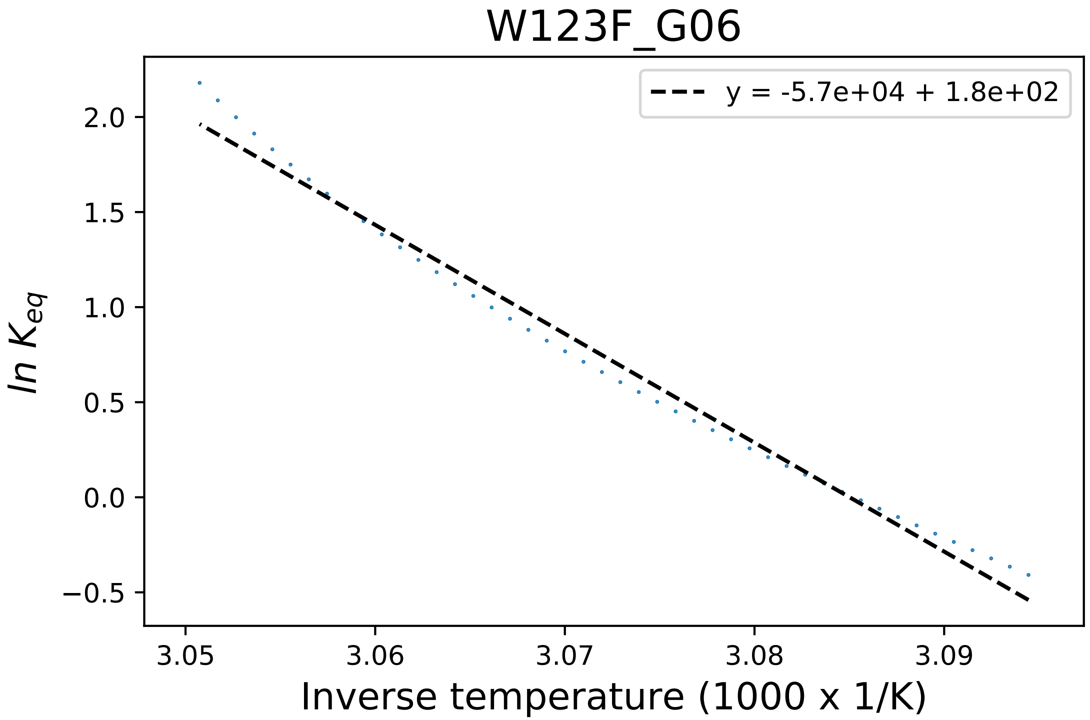

### W123F_G06_fluorescence.pdf

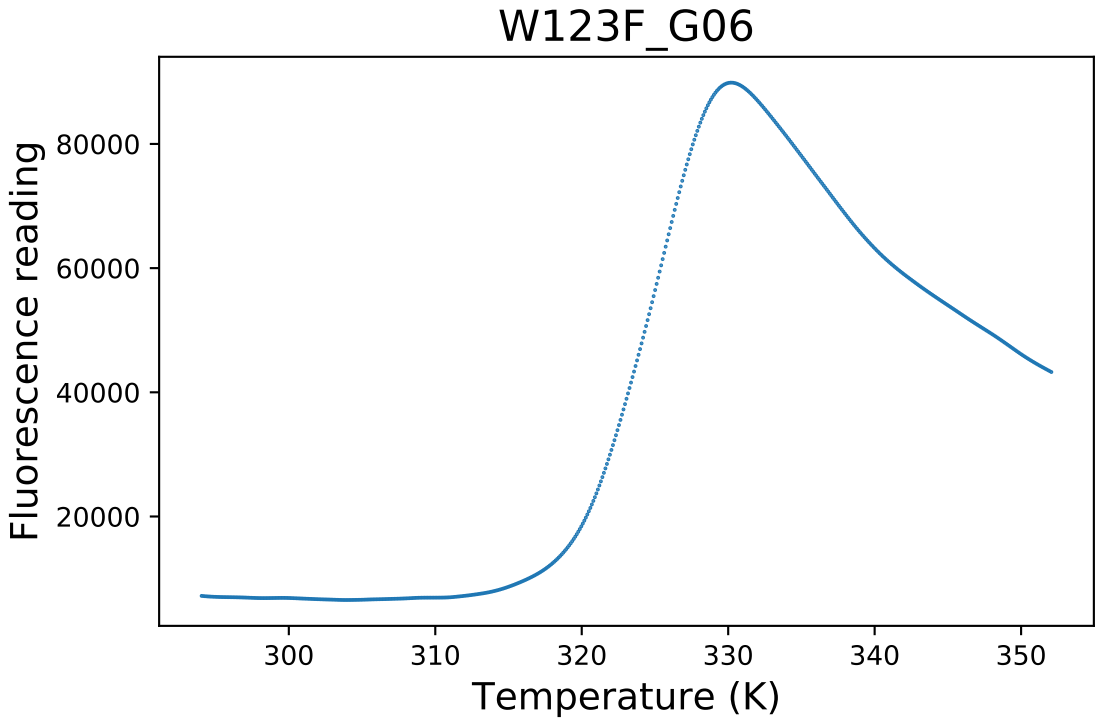

### W123H_F03_dg.pdf

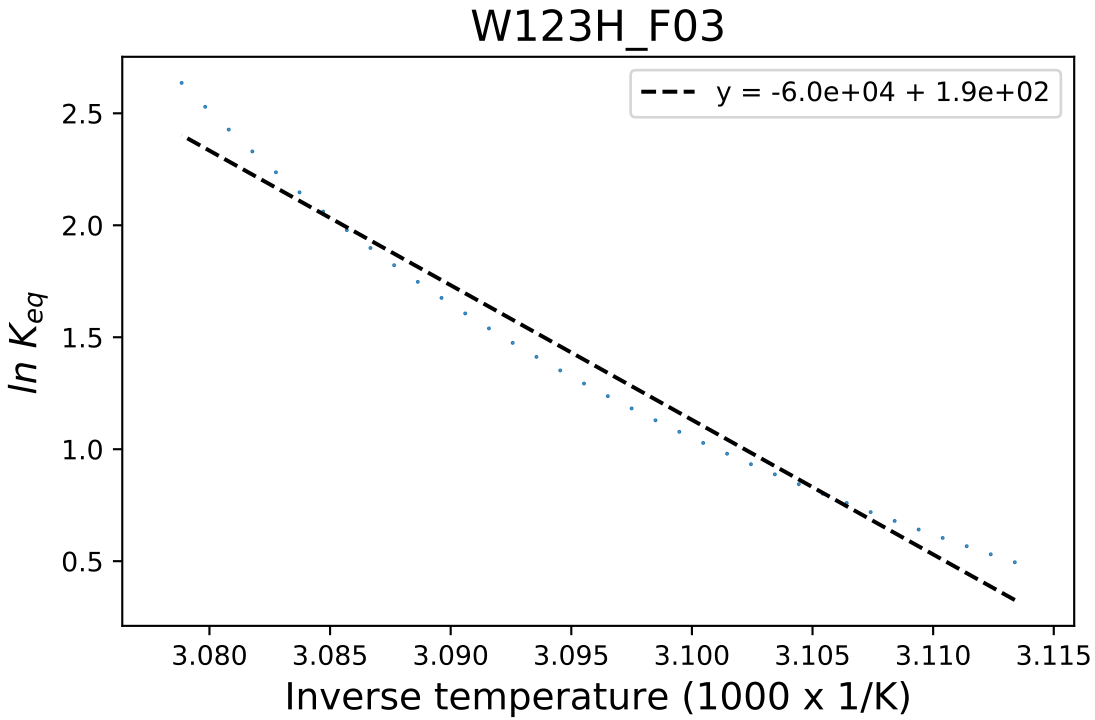

### W328A_C01_derivative.pdf

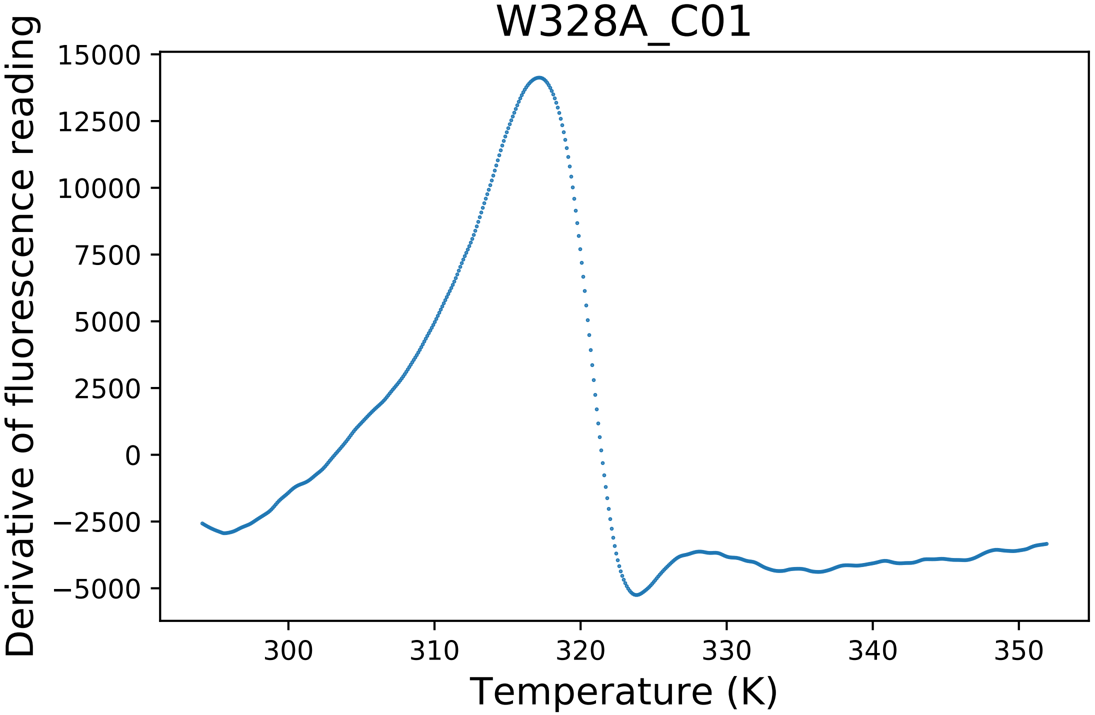

### W328A_C03_fluorescence.pdf

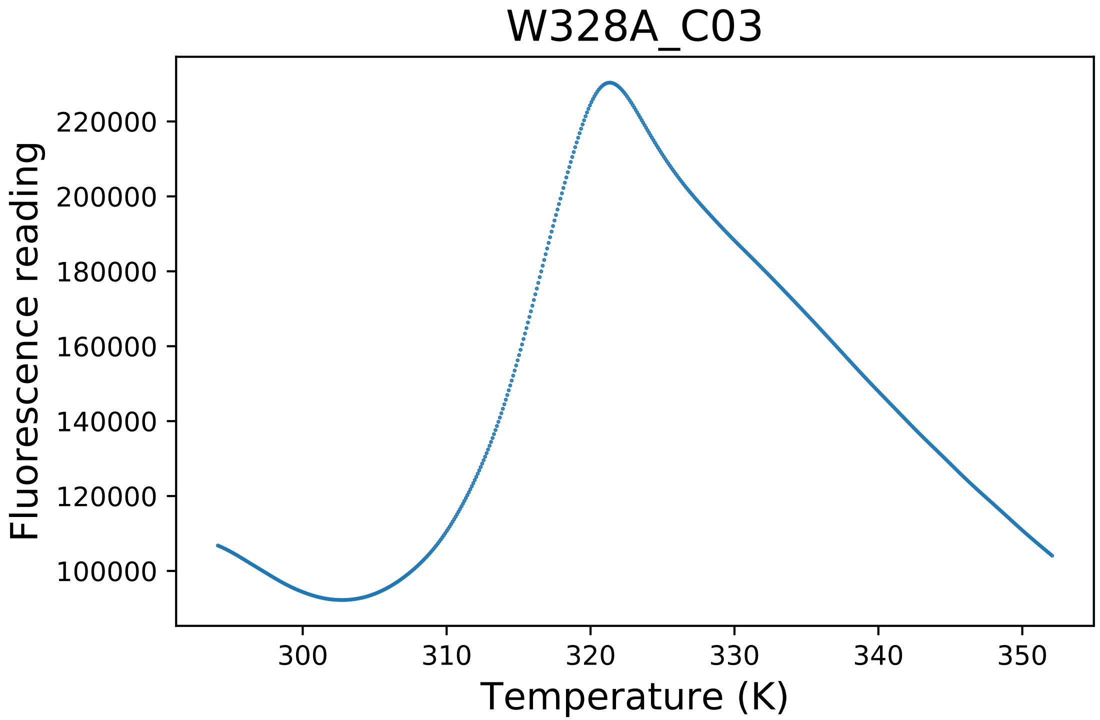

### W328C_A10_derivative.pdf

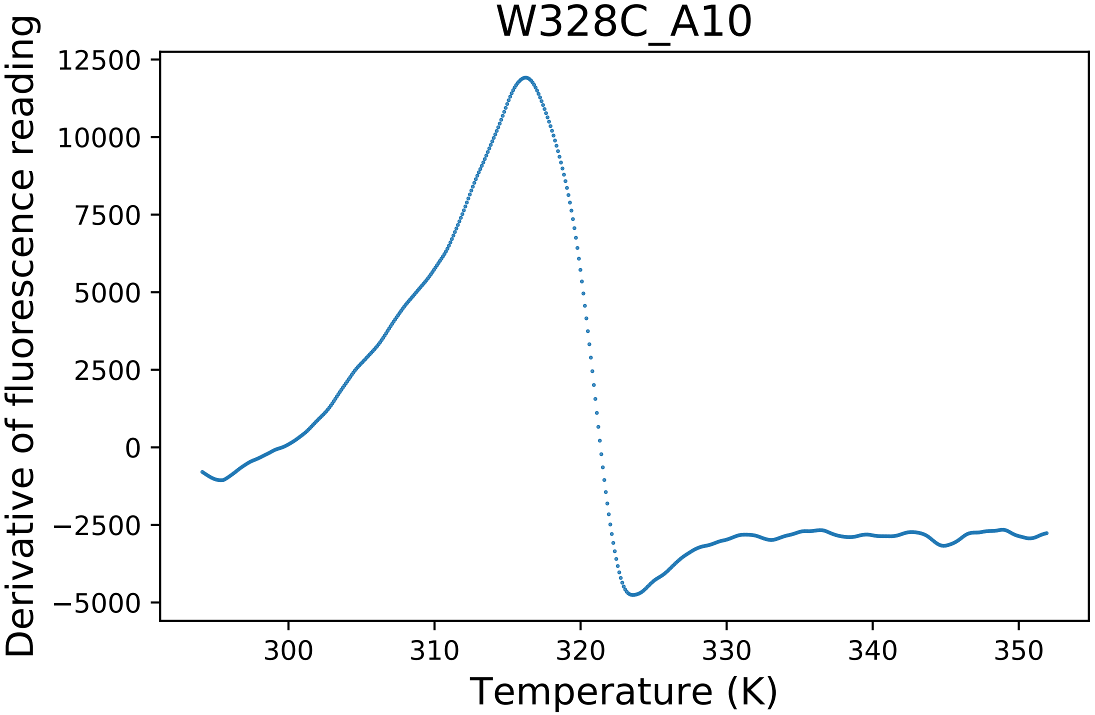

### W328C_A12_fluorescence.pdf

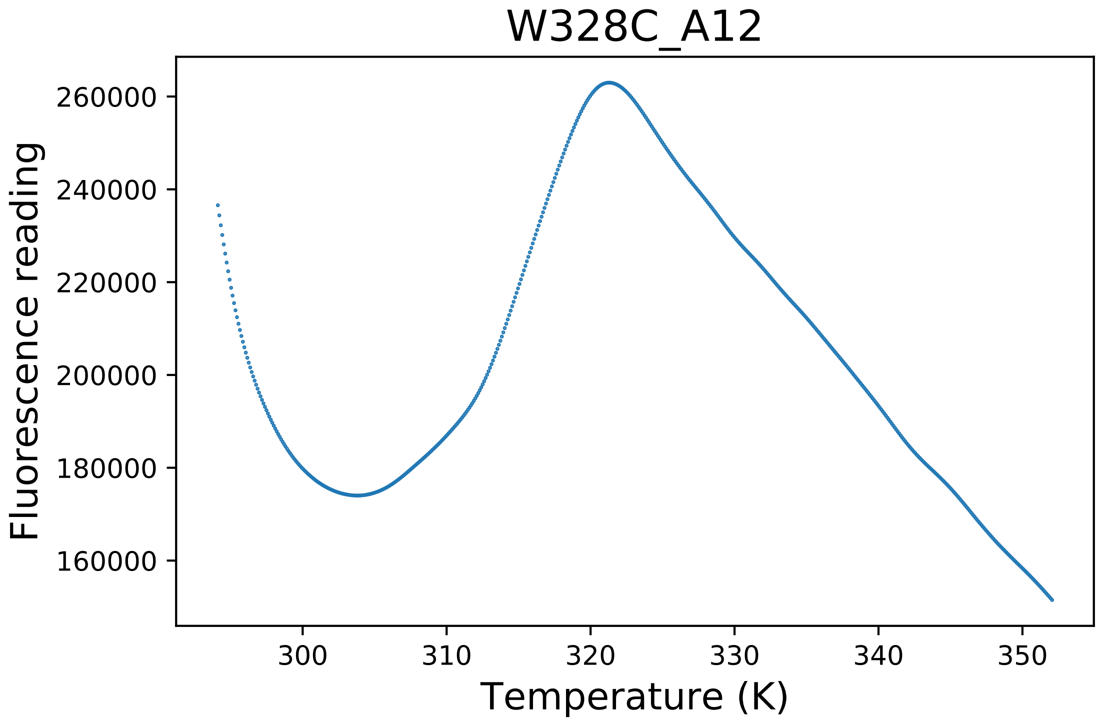

### W328H_A07_derivative.pdf

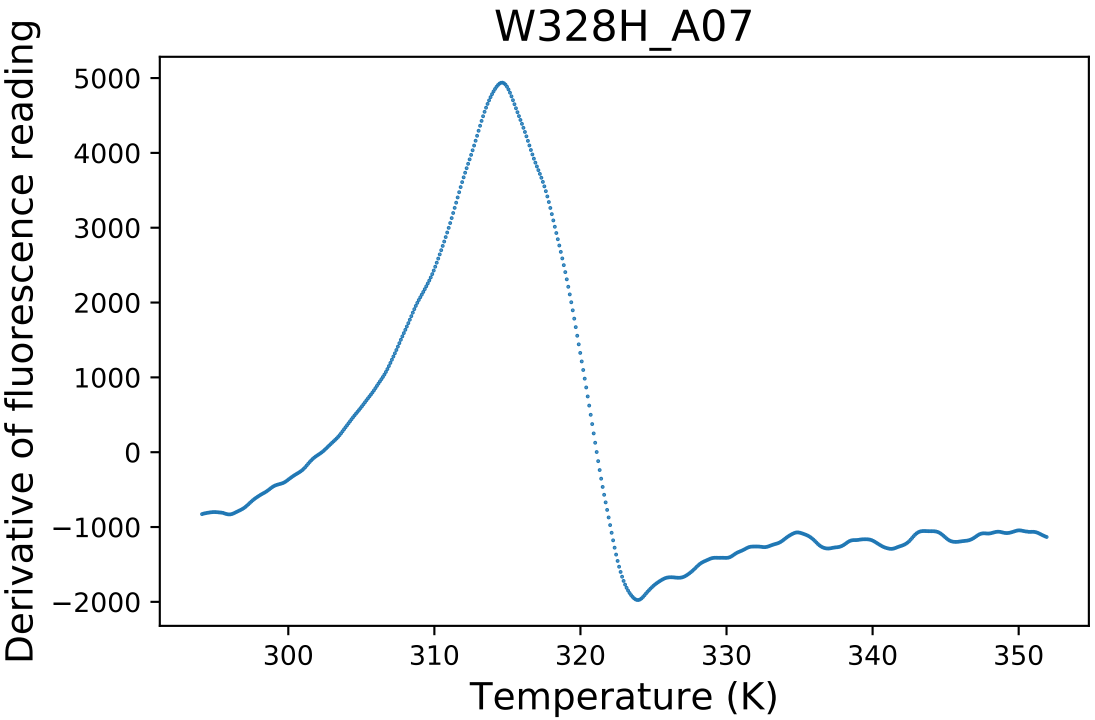

### W328H_A07_dg.pdf

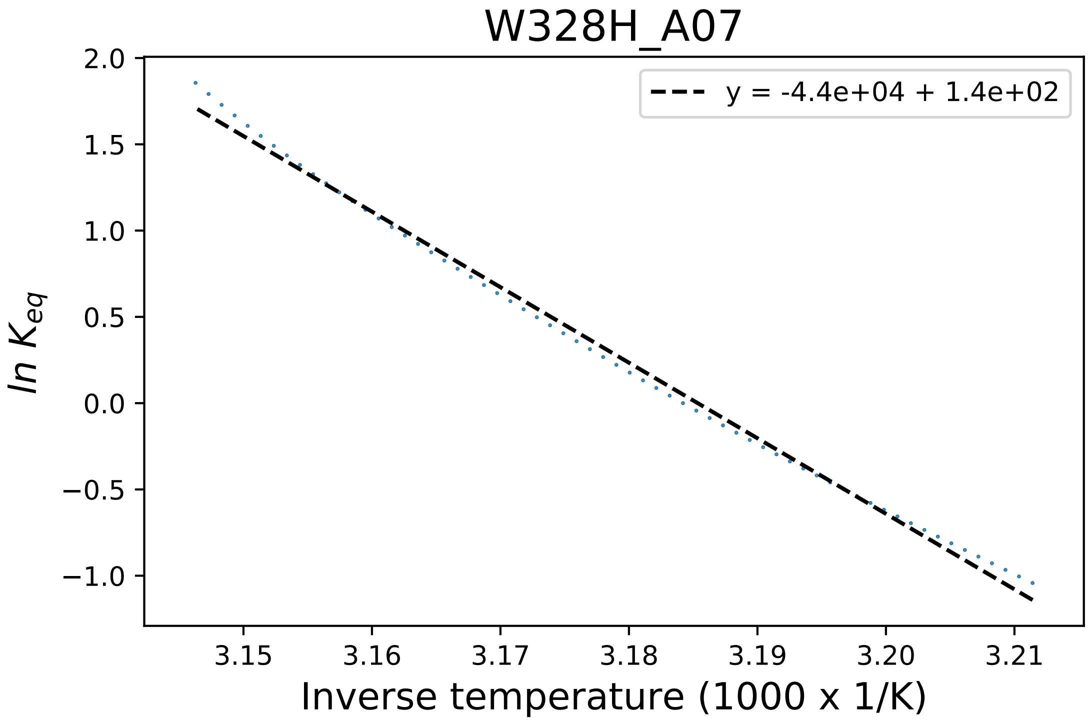

### W328H_A07_fluorescence.pdf

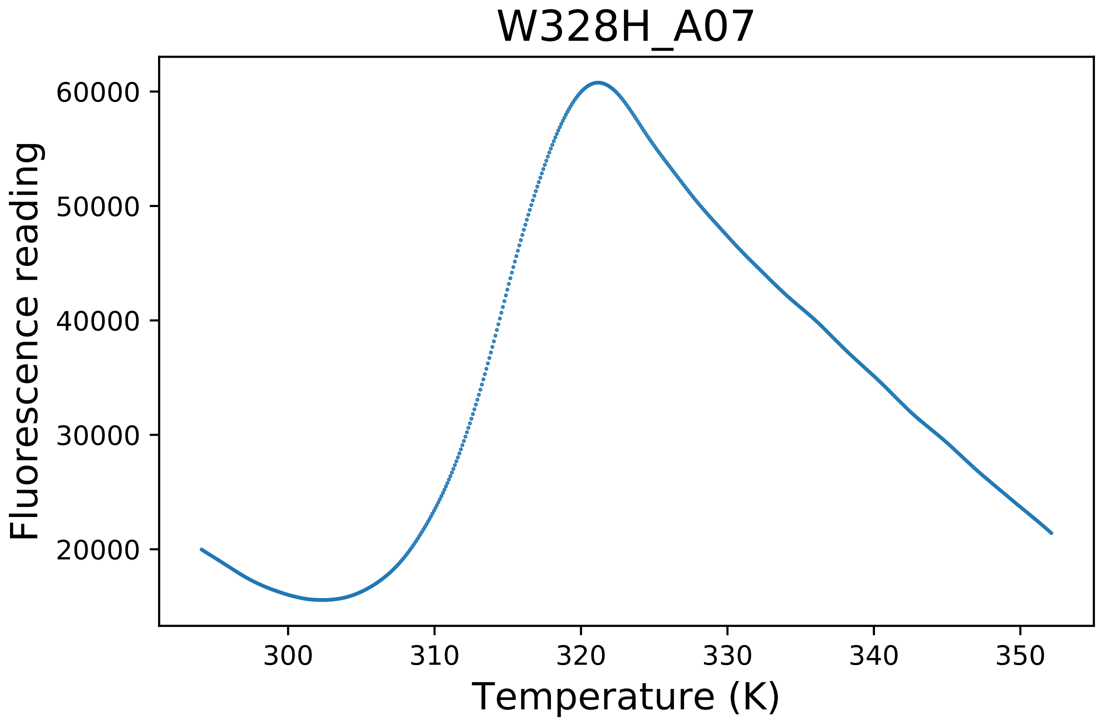

### W328H_A08_derivative.pdf

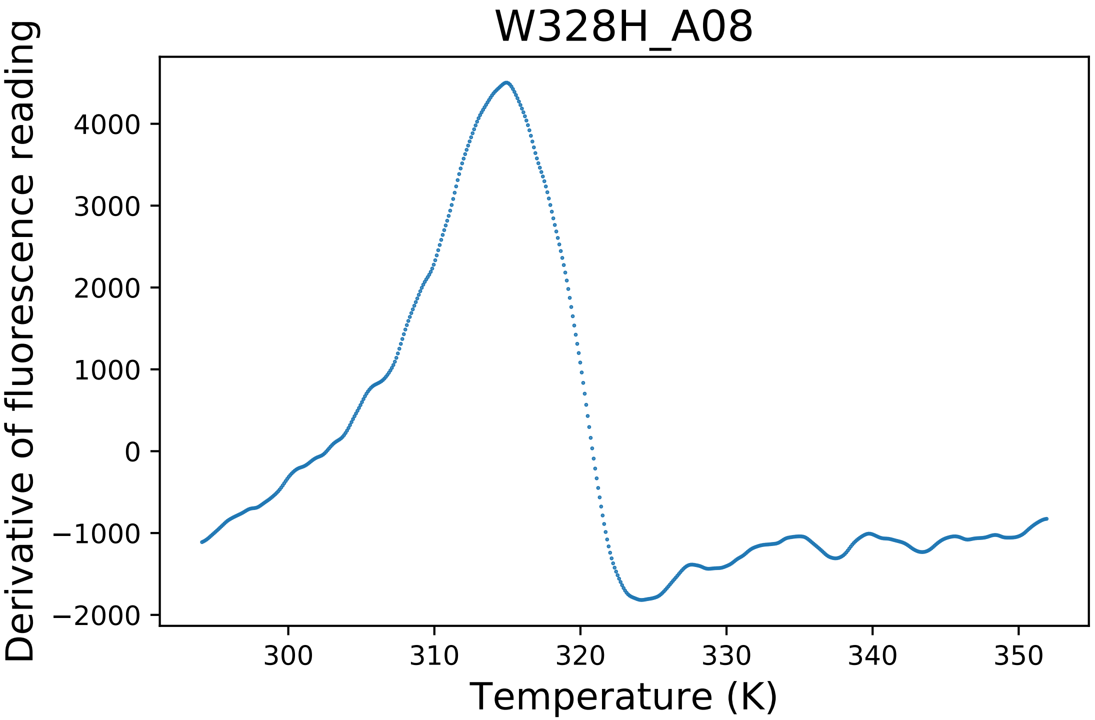

### W328R_D05_derivative.pdf

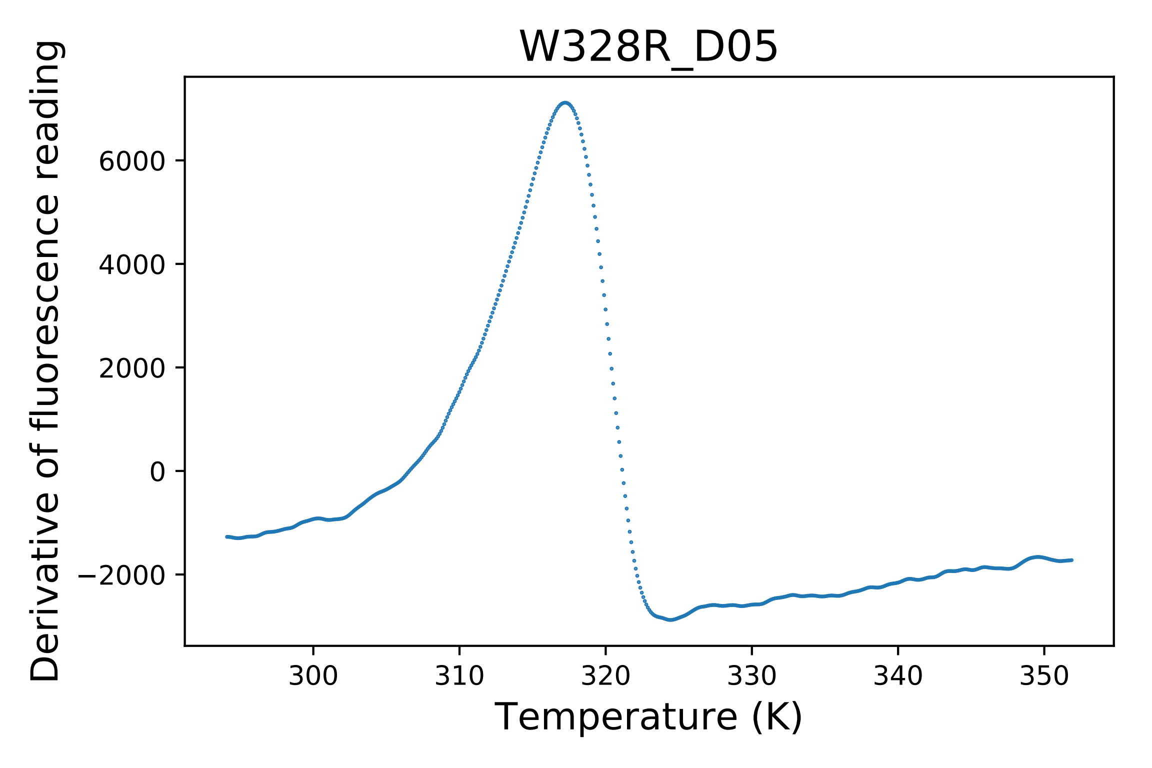

### W328R_D08_fluorescence.pdf

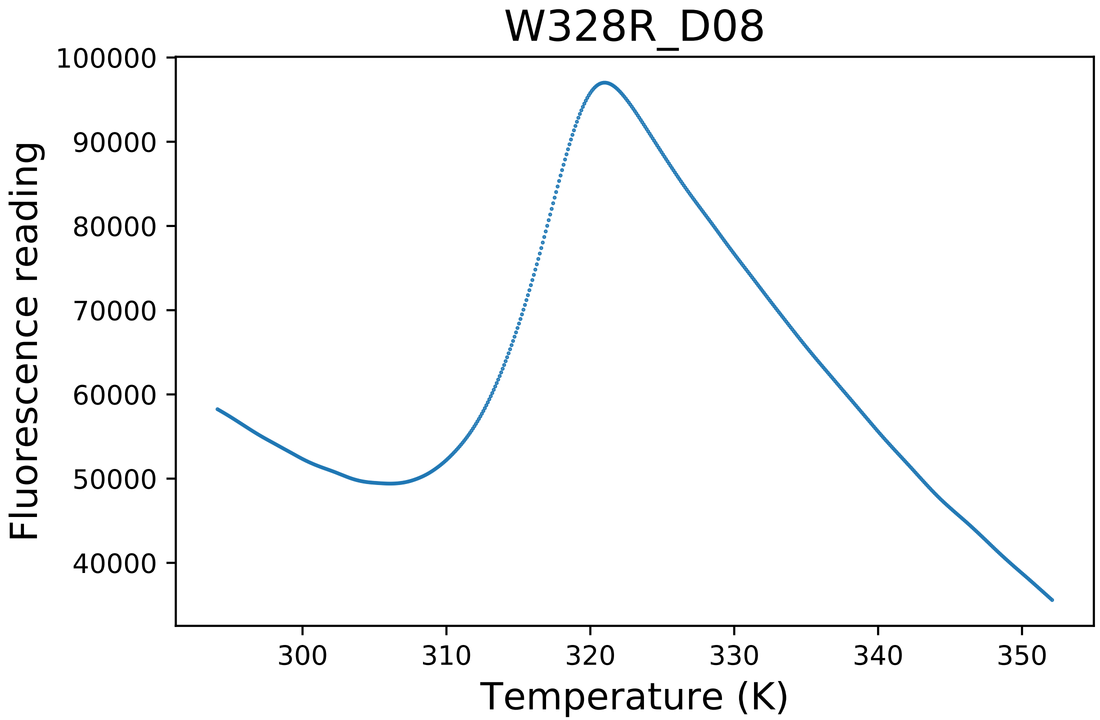

### W402C_E08_dg.pdf

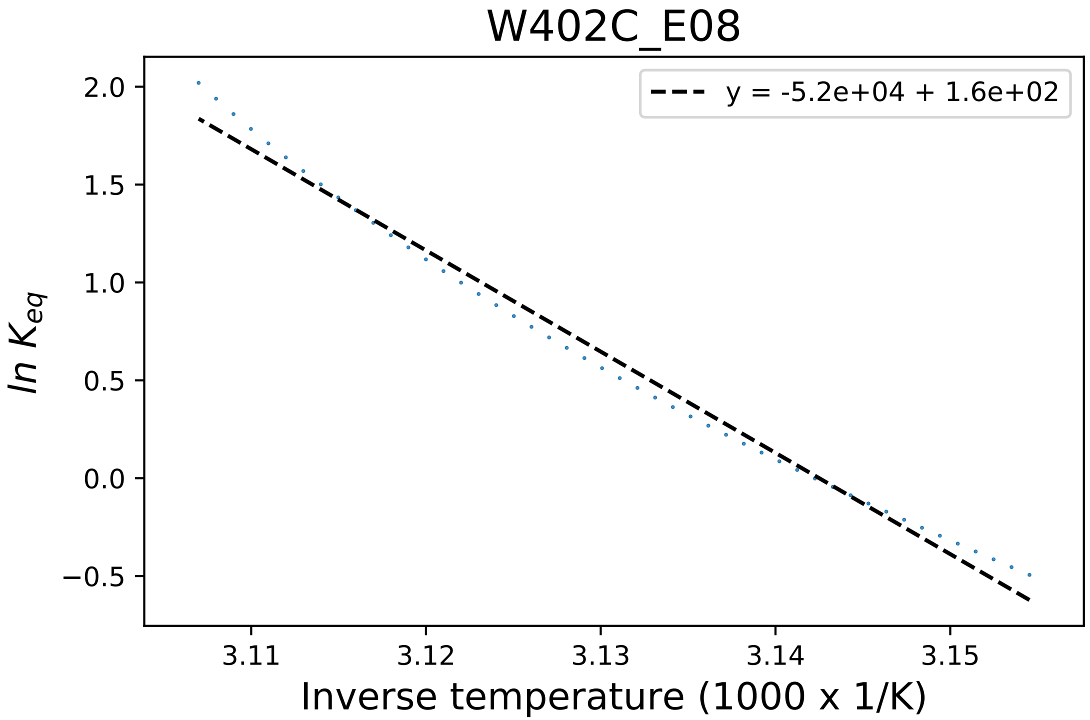

### W402R_A05_dg.pdf

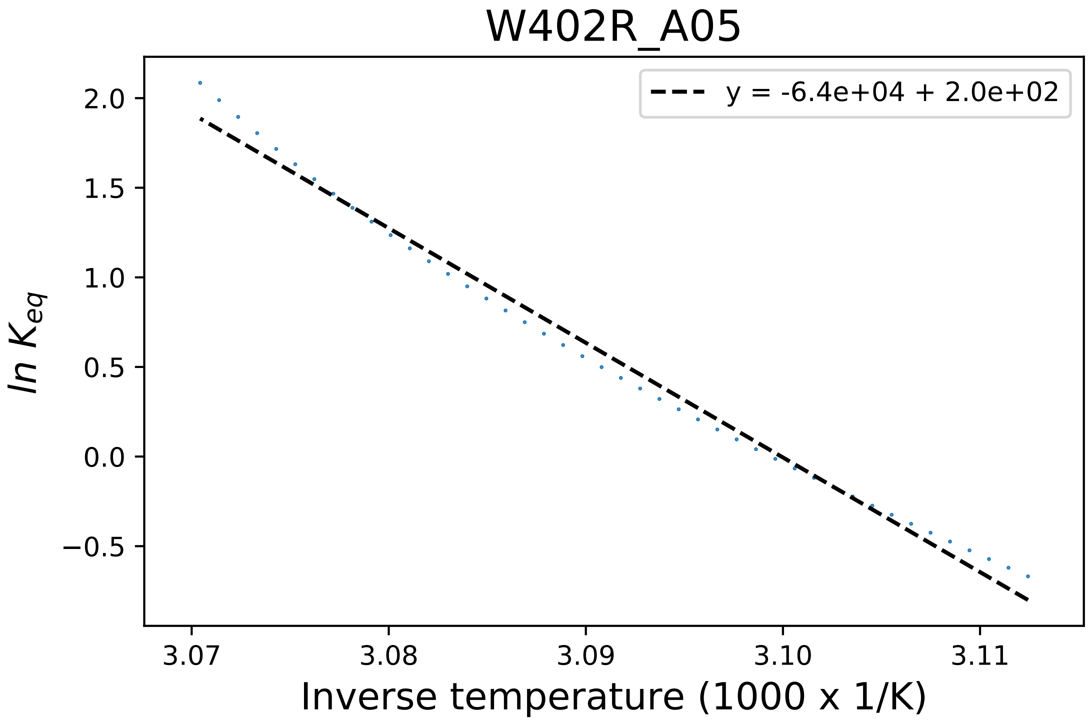

### W402R_A06_fluorescence.pdf

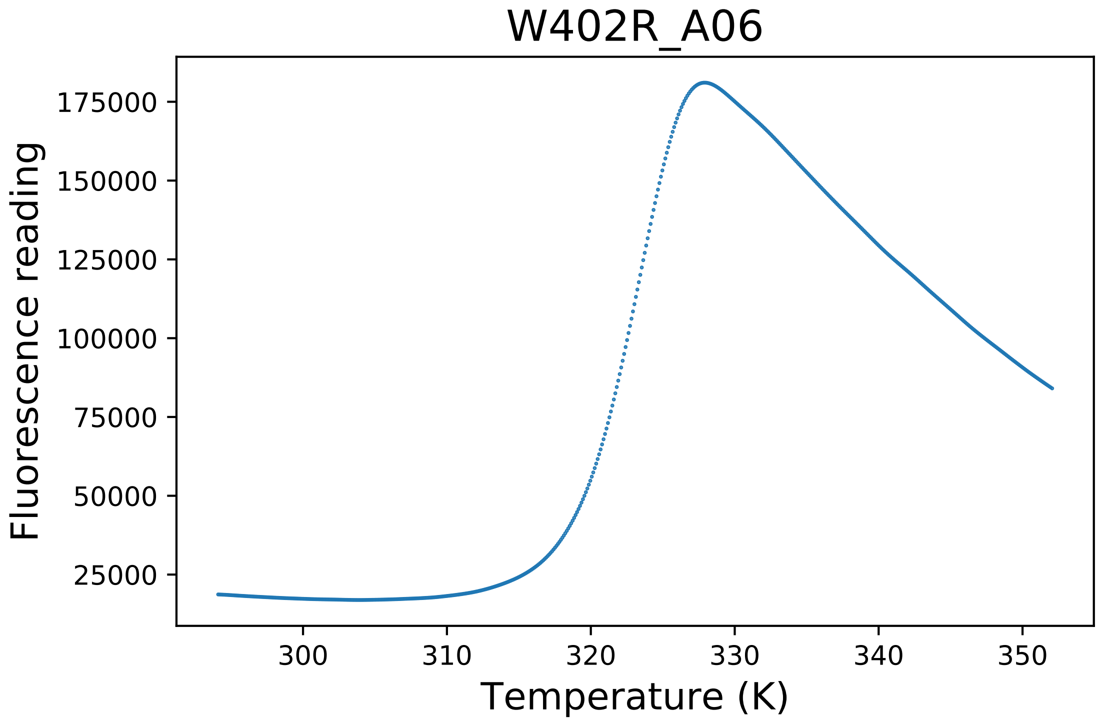

### W402R_A07_derivative.pdf

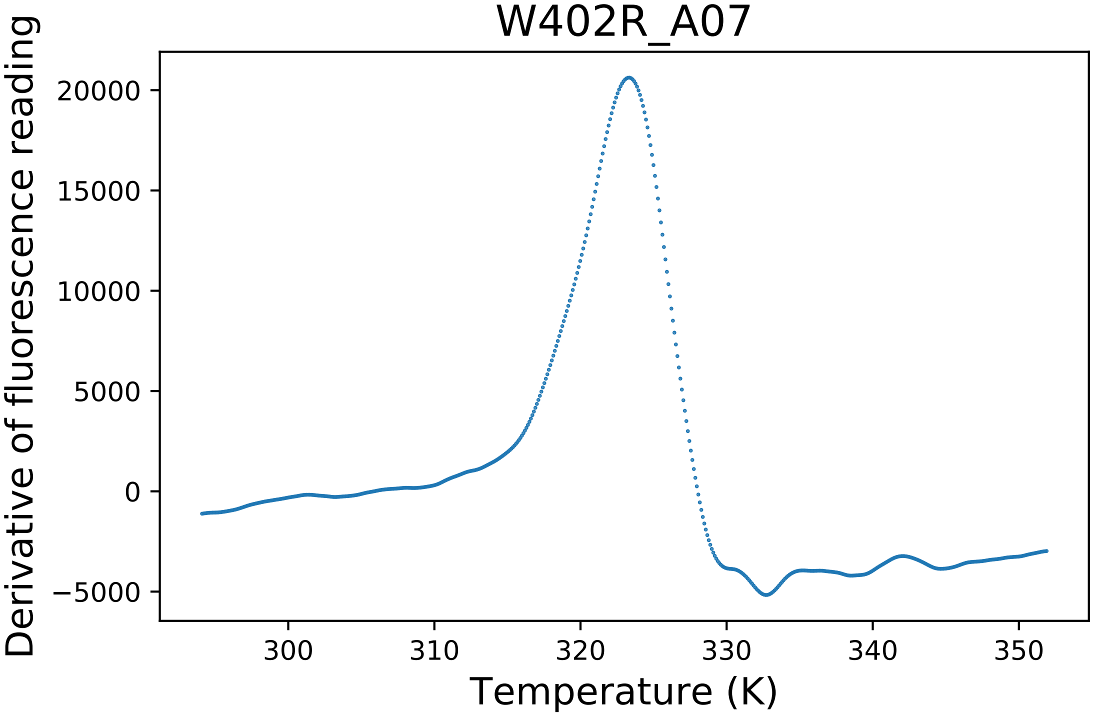

### W402R_A07_dg.pdf

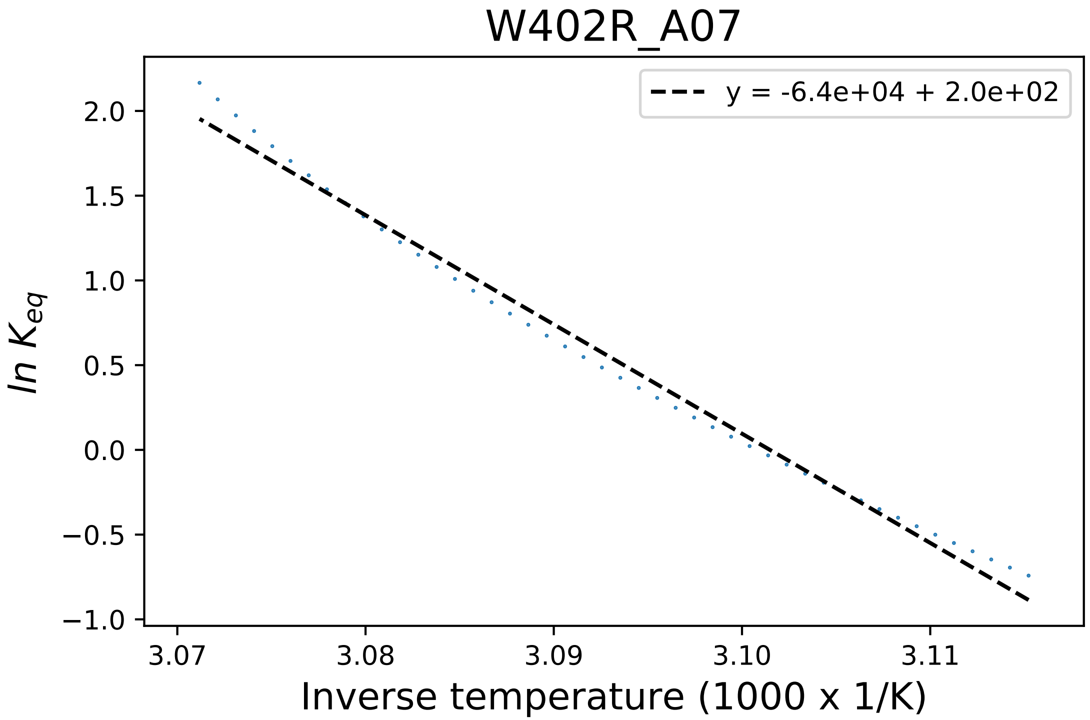

### W402R_A08_derivative.pdf

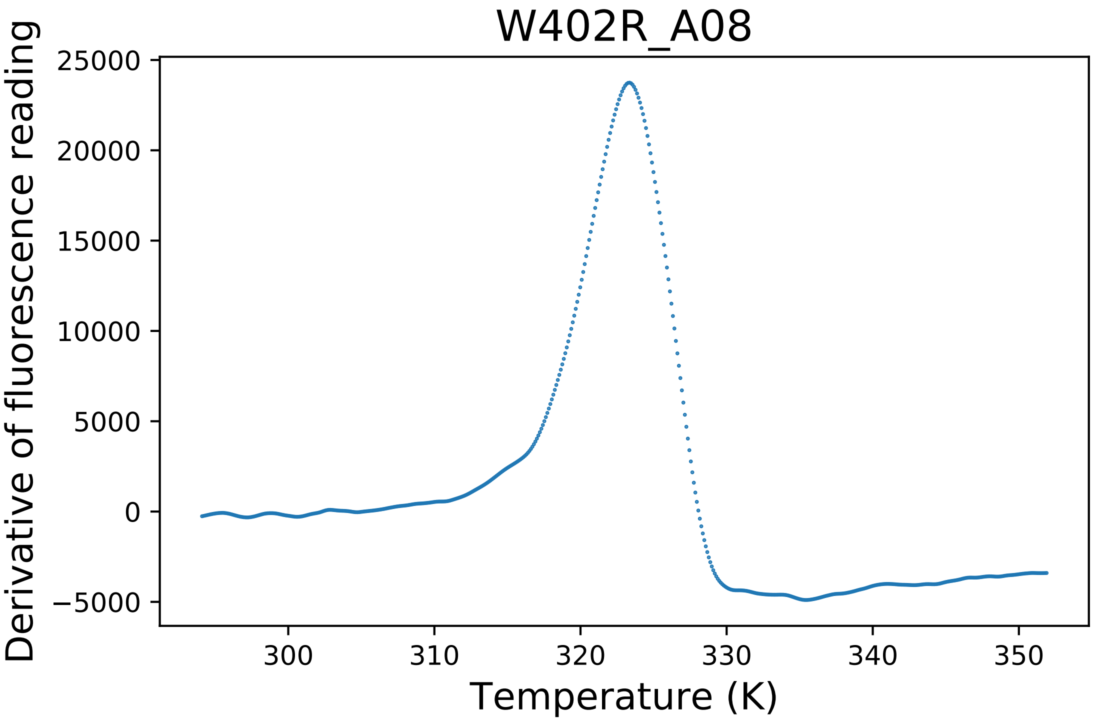

### W412Y_B06_fluorescence.pdf

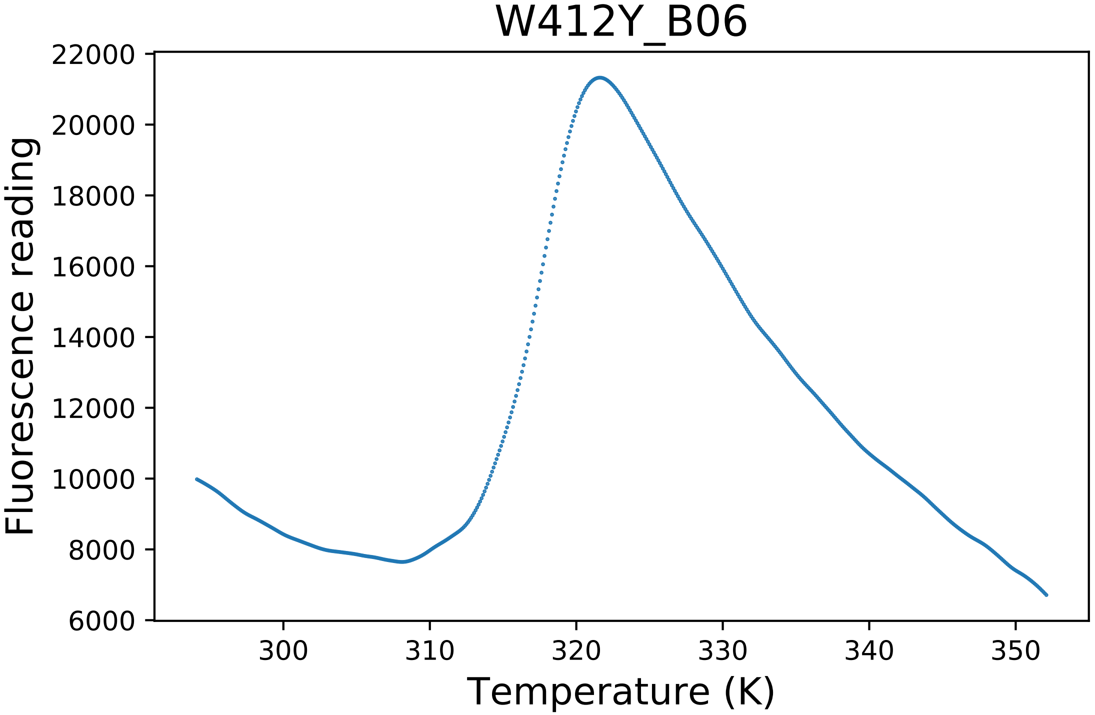

### WT1_A01_dg.pdf

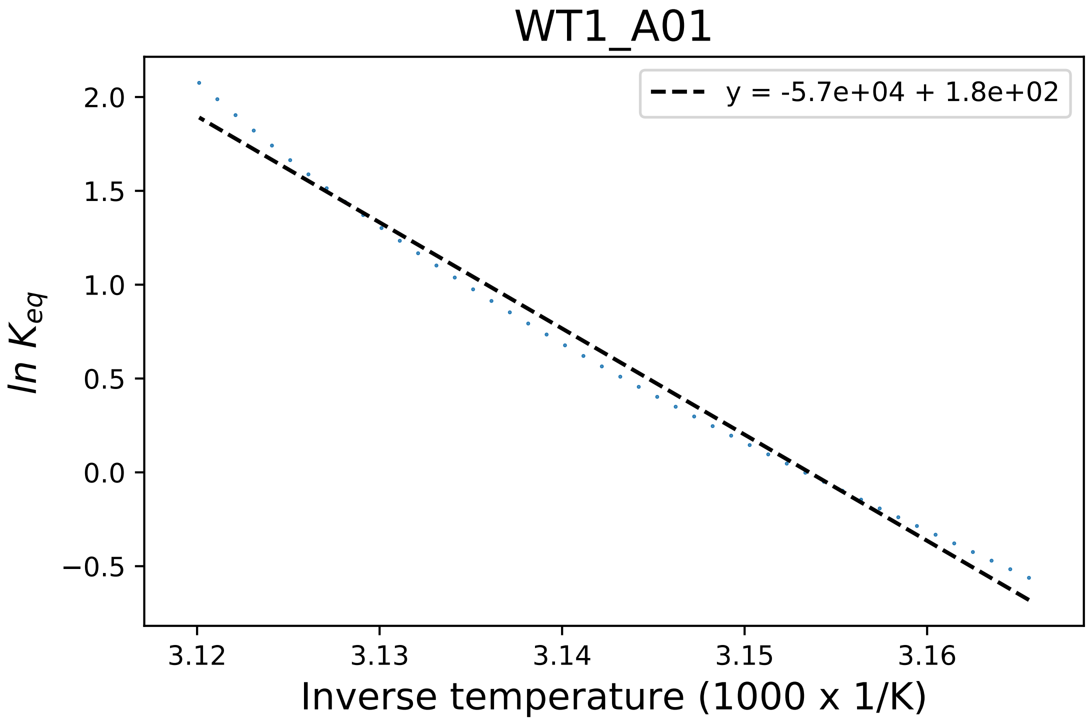

### WT1_A04_fluorescence.pdf

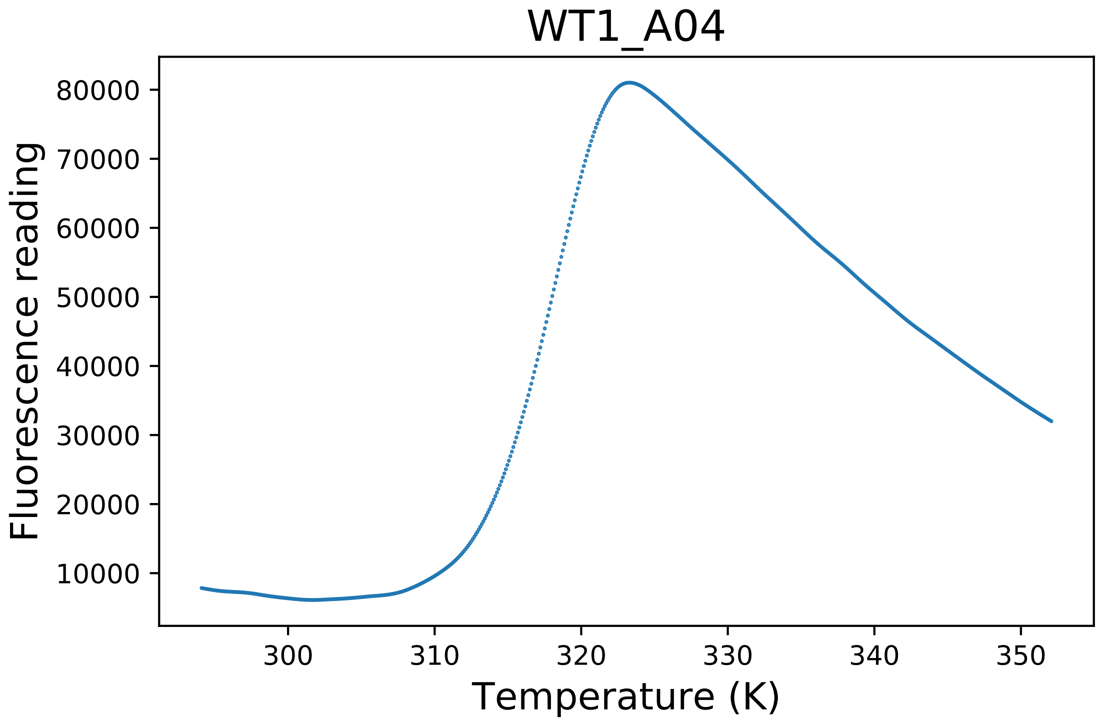

### WT2_H10_derivative.pdf

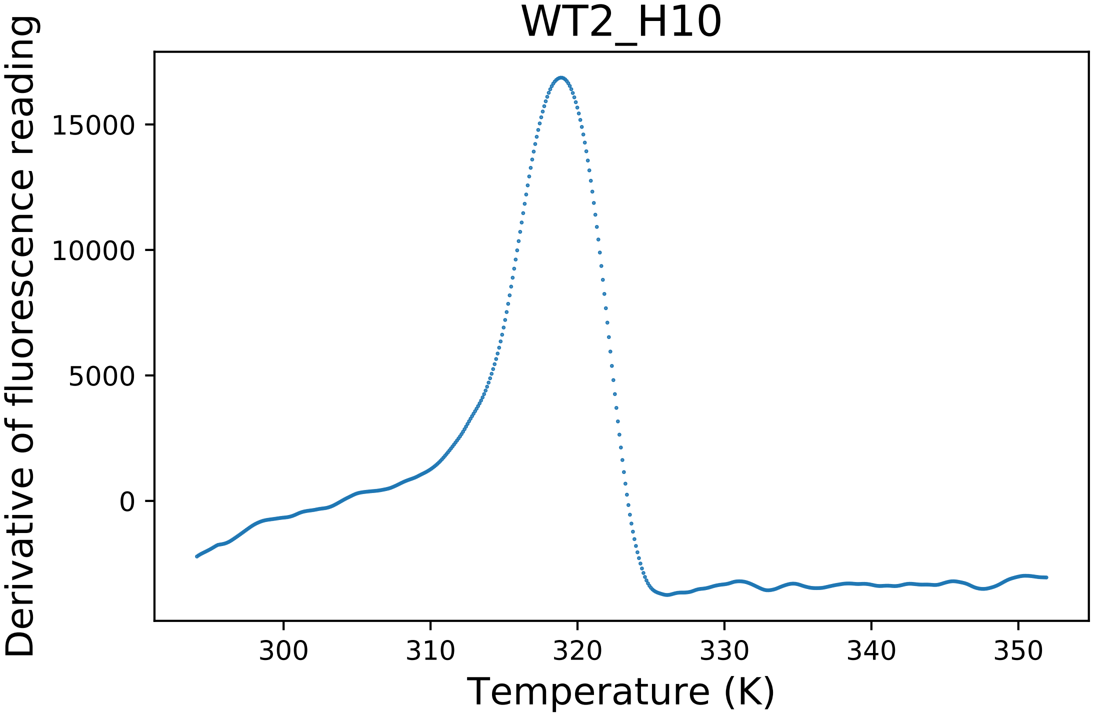

### WT2_H11_derivative.pdf

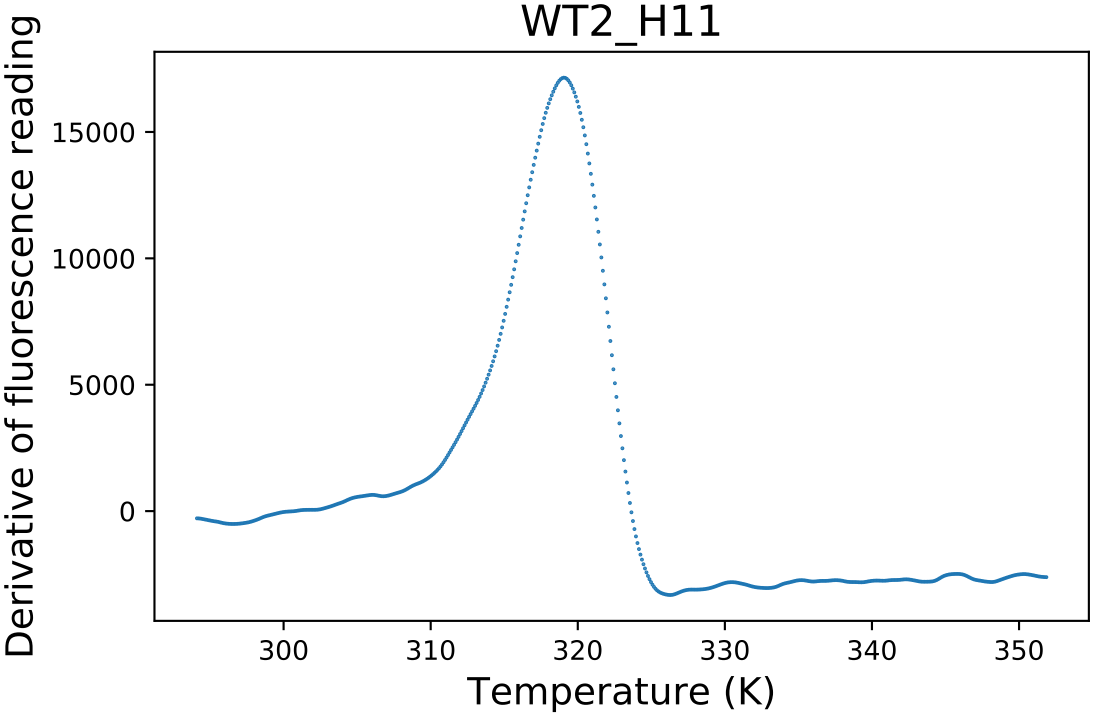

### WT2_H11_fluorescence.pdf

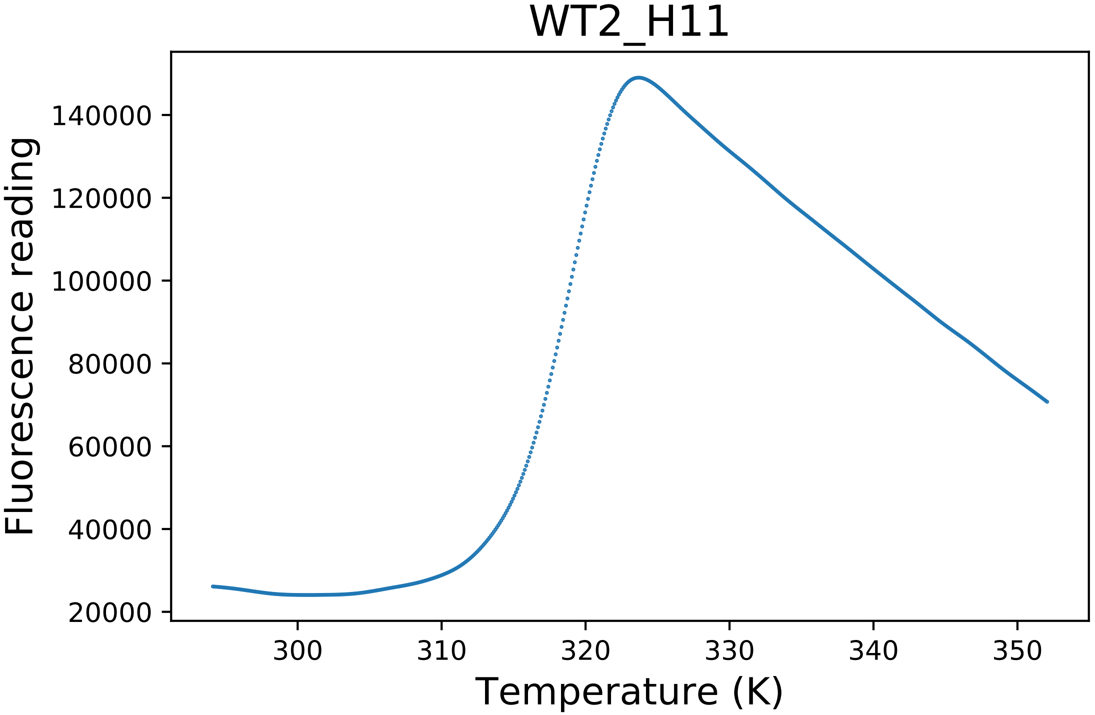

### WT3_A01_derivative.pdf

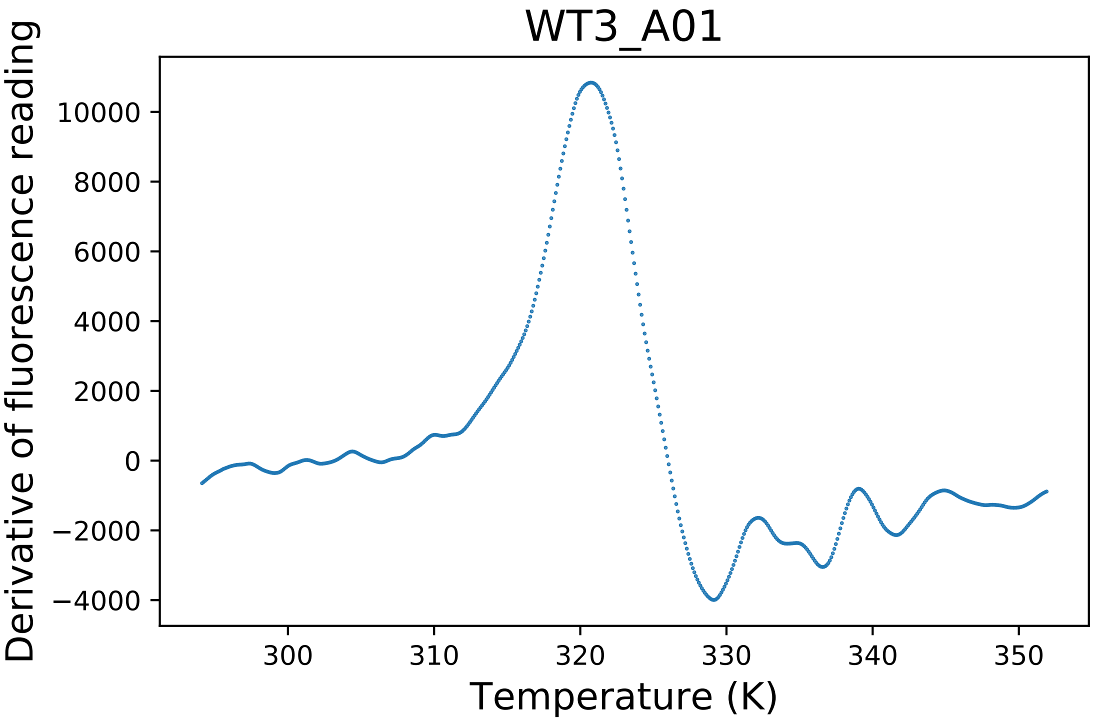

### WT3_A02_fluorescence.pdf

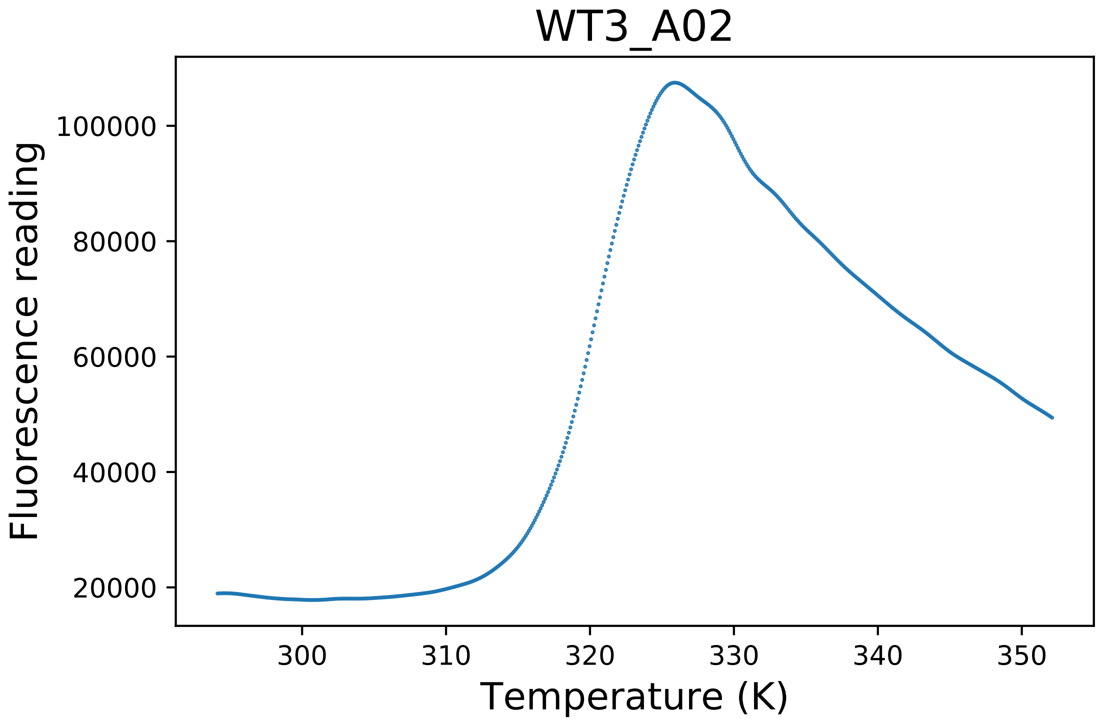

### WT4_A03_dg.pdf

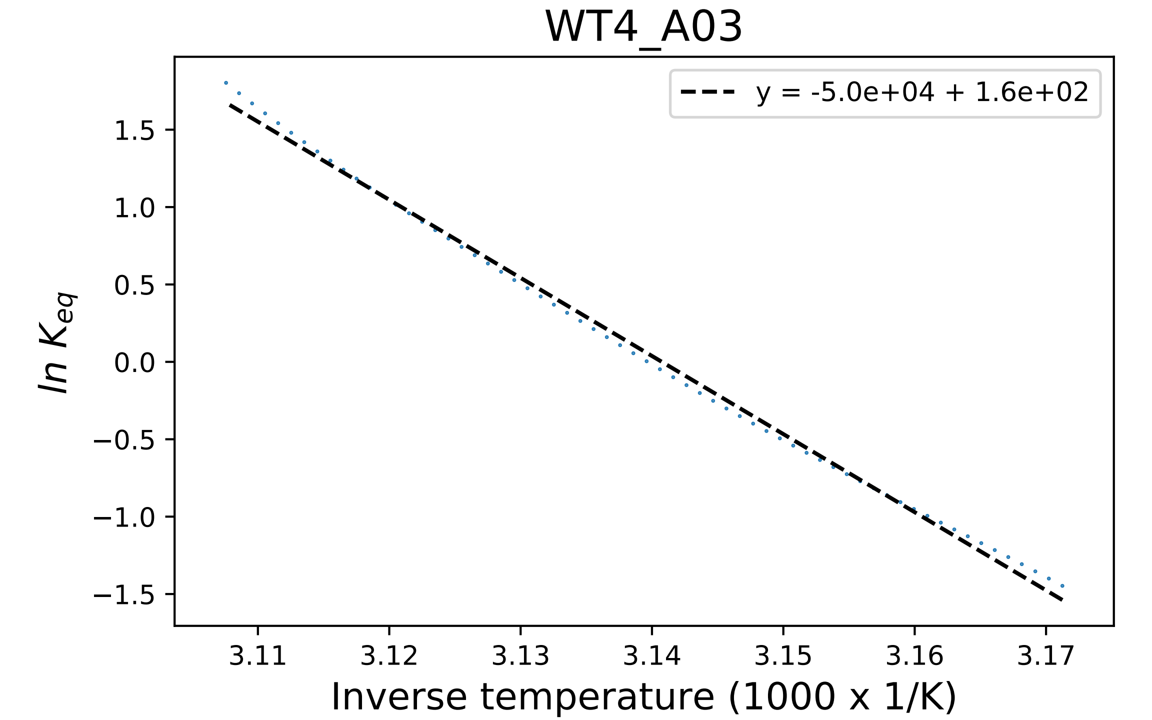

### WT5_F03_fluorescence.pdf

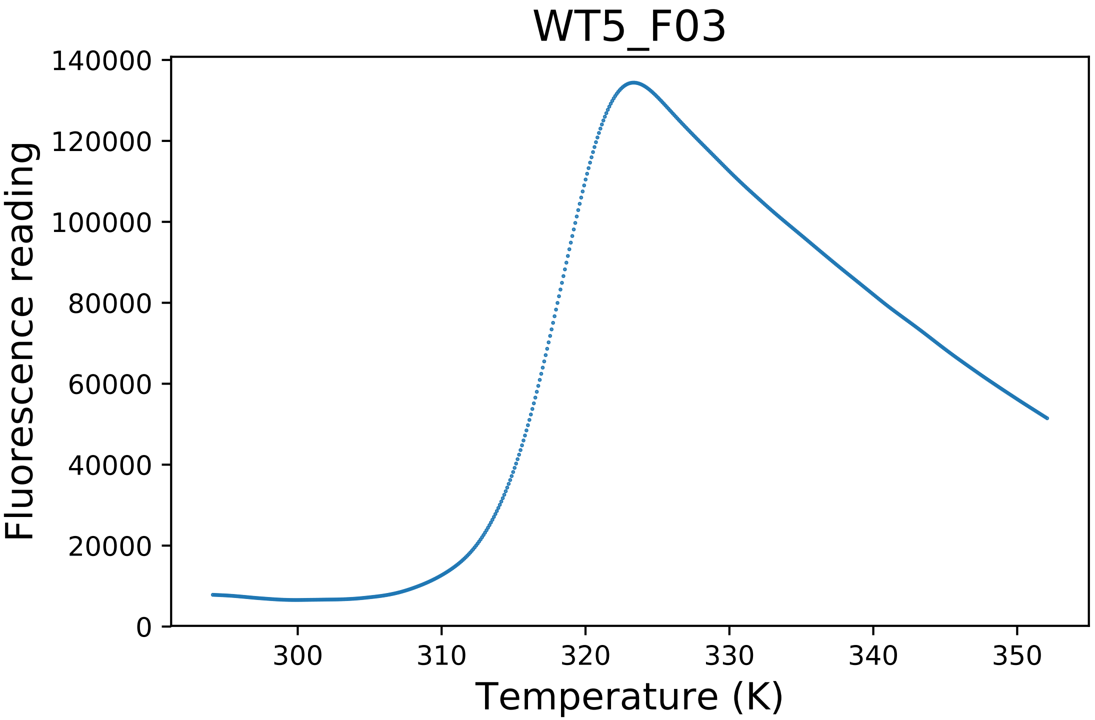

### WT5_F04_dg.pdf

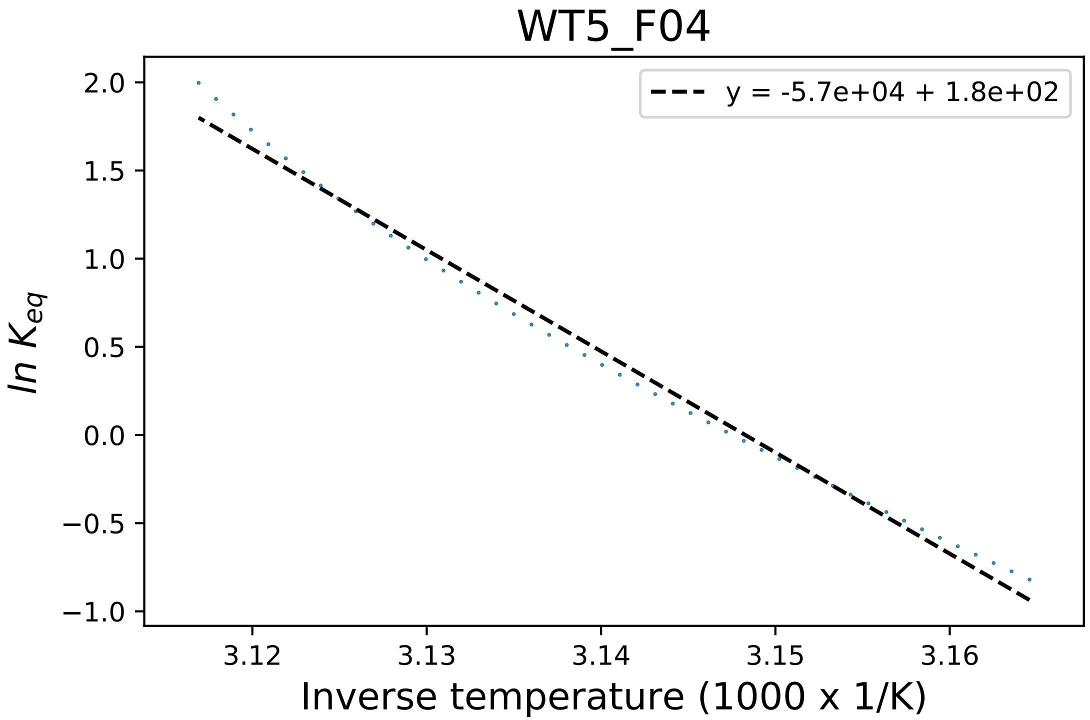

### WT6_D07_derivative.pdf

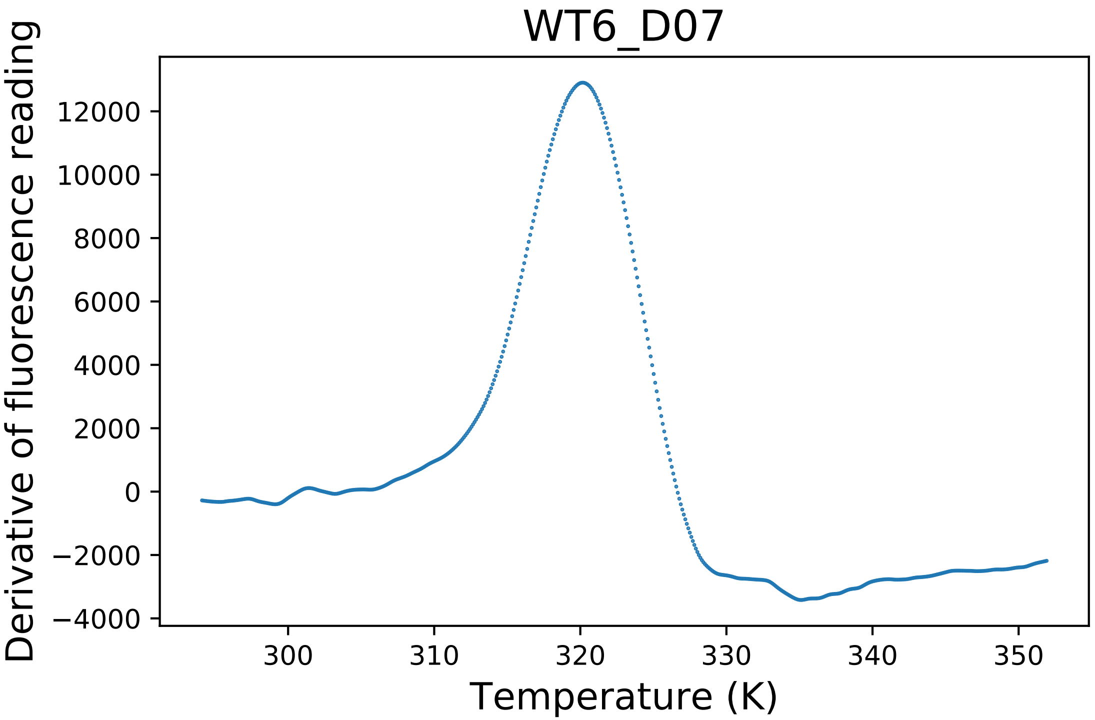

### WT6_D07_fluorescence.pdf

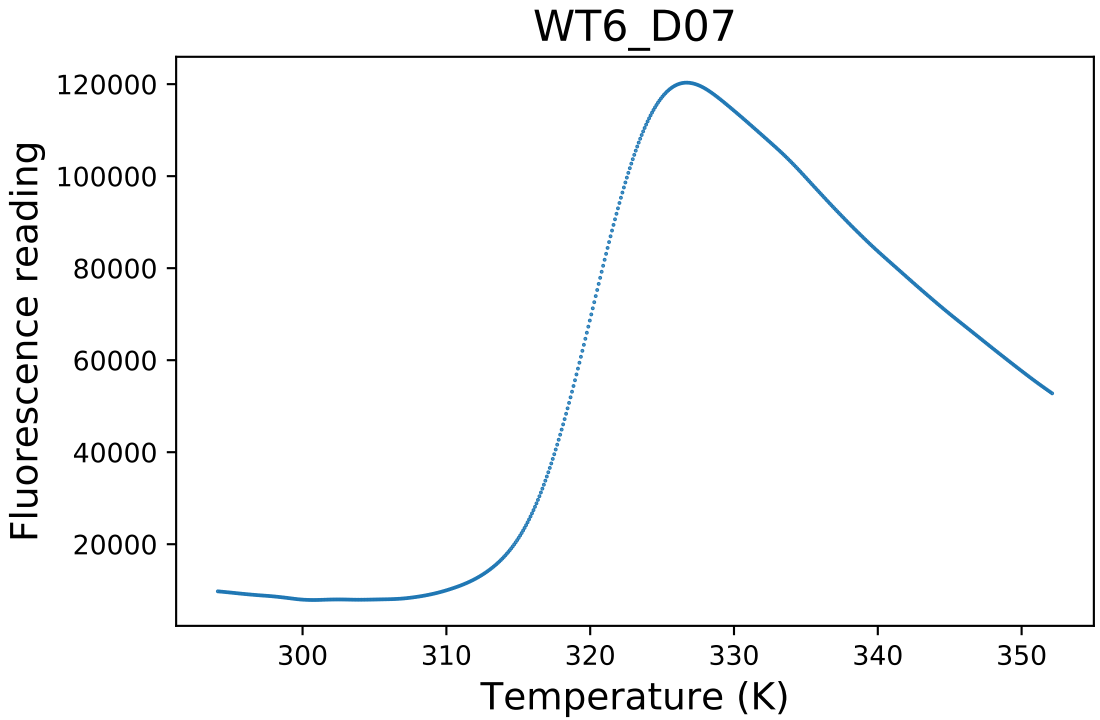

### WT6_D08_derivative.pdf

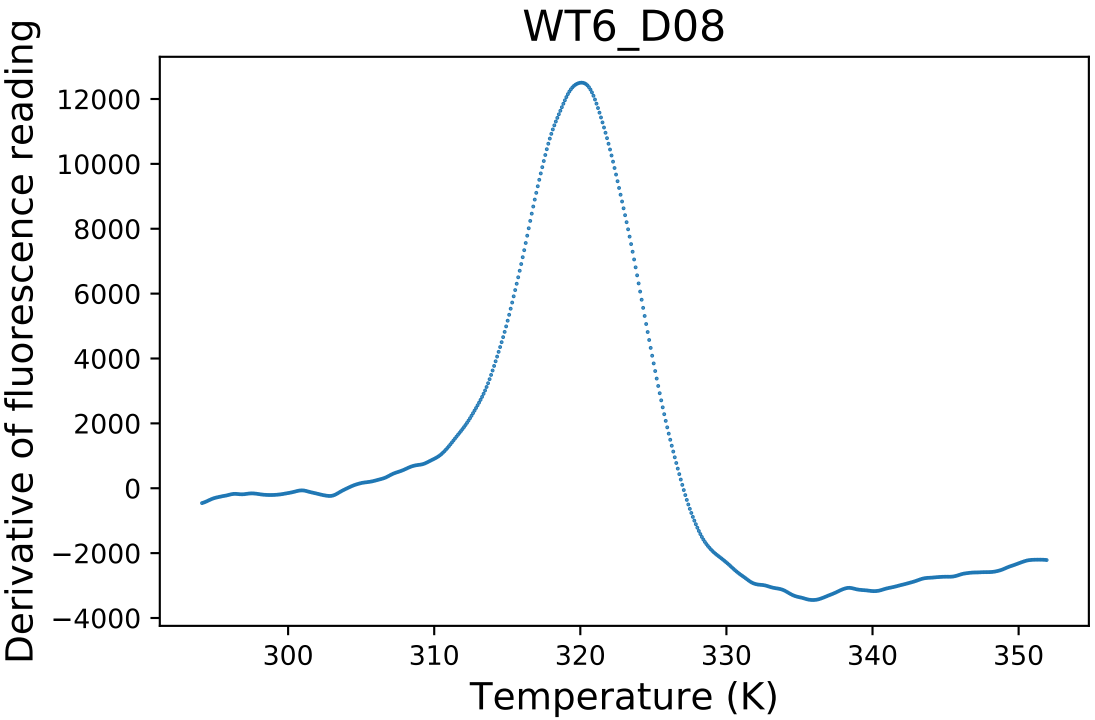

### Y21A_B02_derivative.pdf

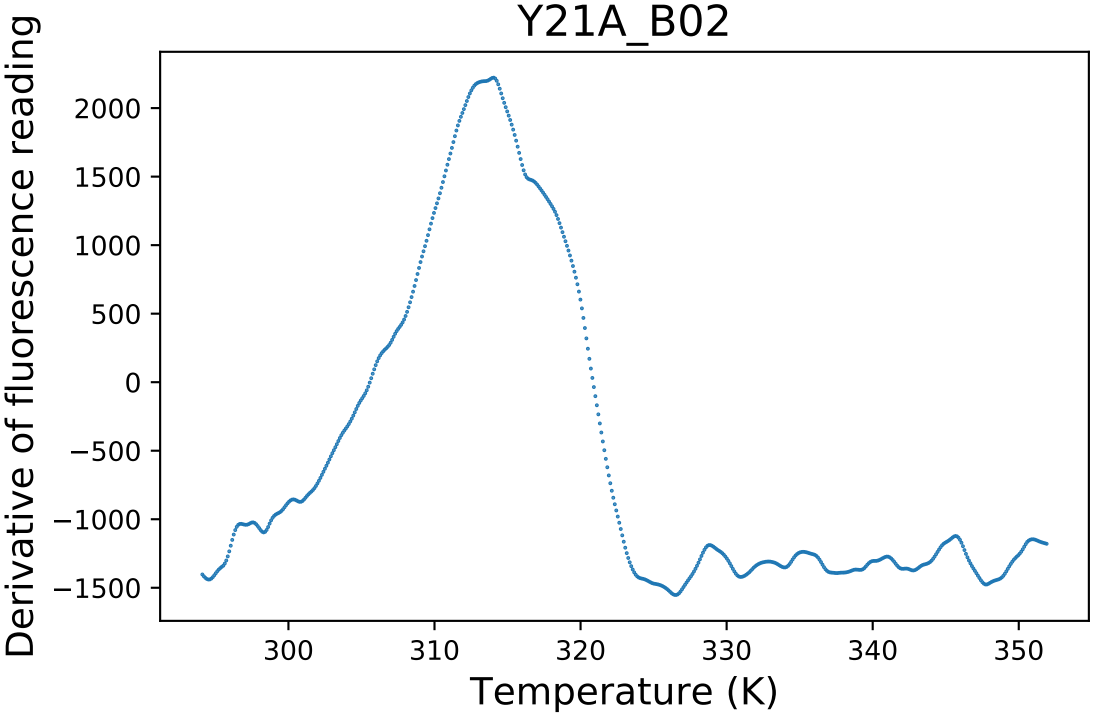

### Y21A_B02_fluorescence.pdf

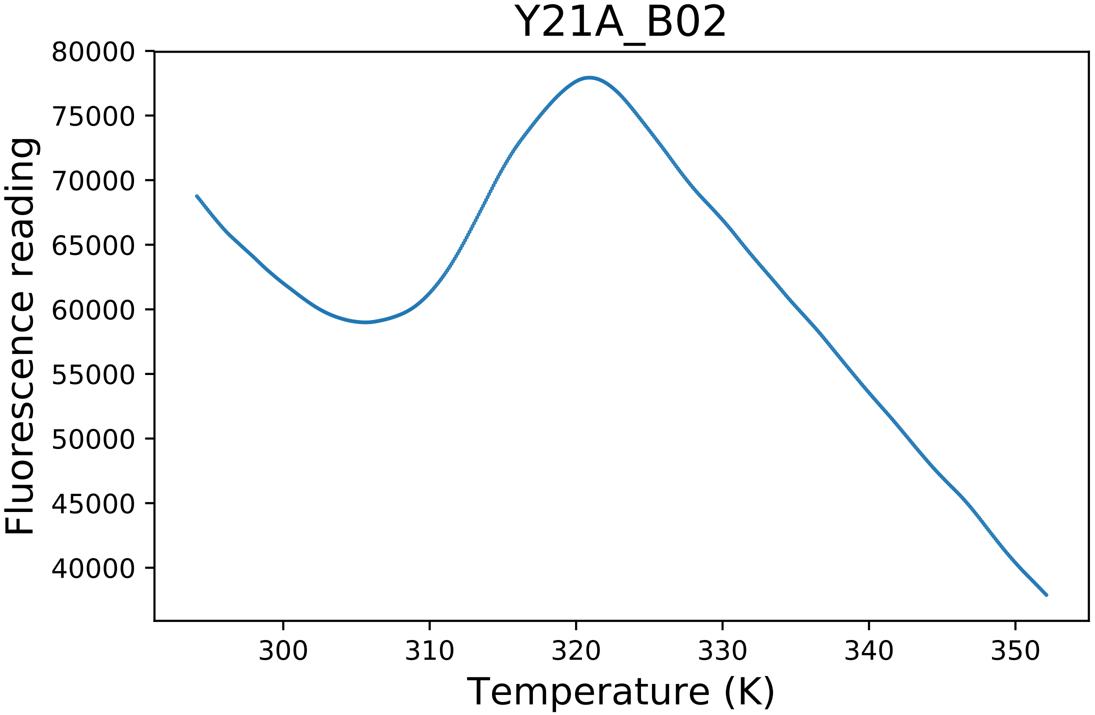

### Y21A_B03_derivative.pdf

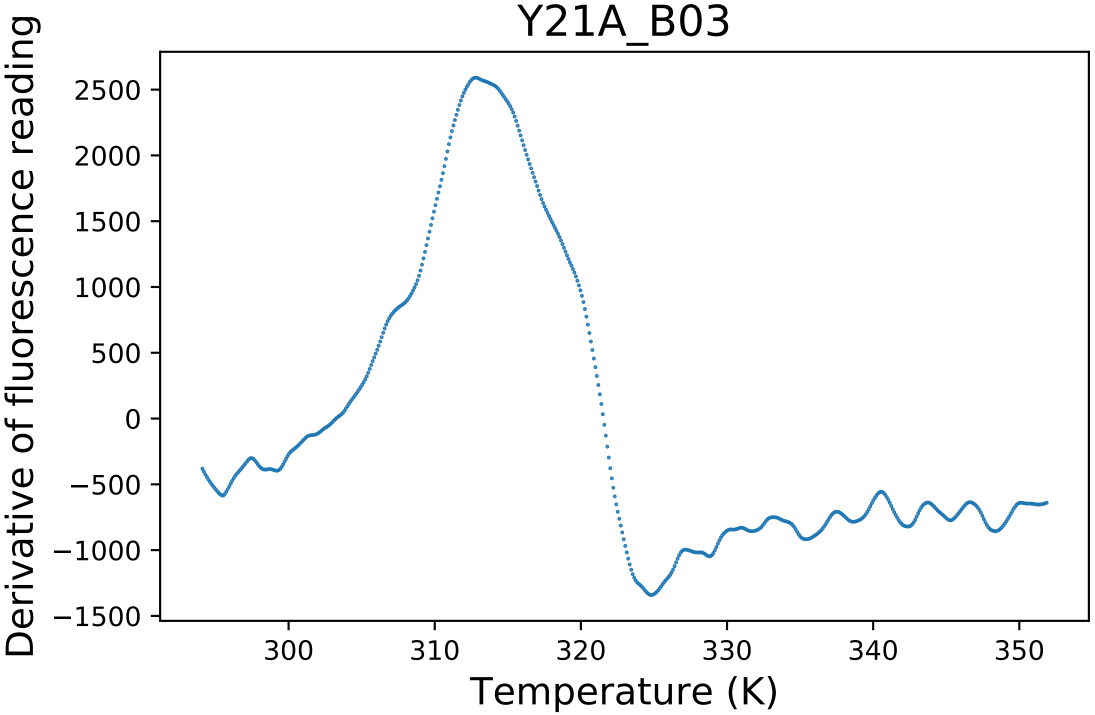

### Y297A_H01_fluorescence.pdf

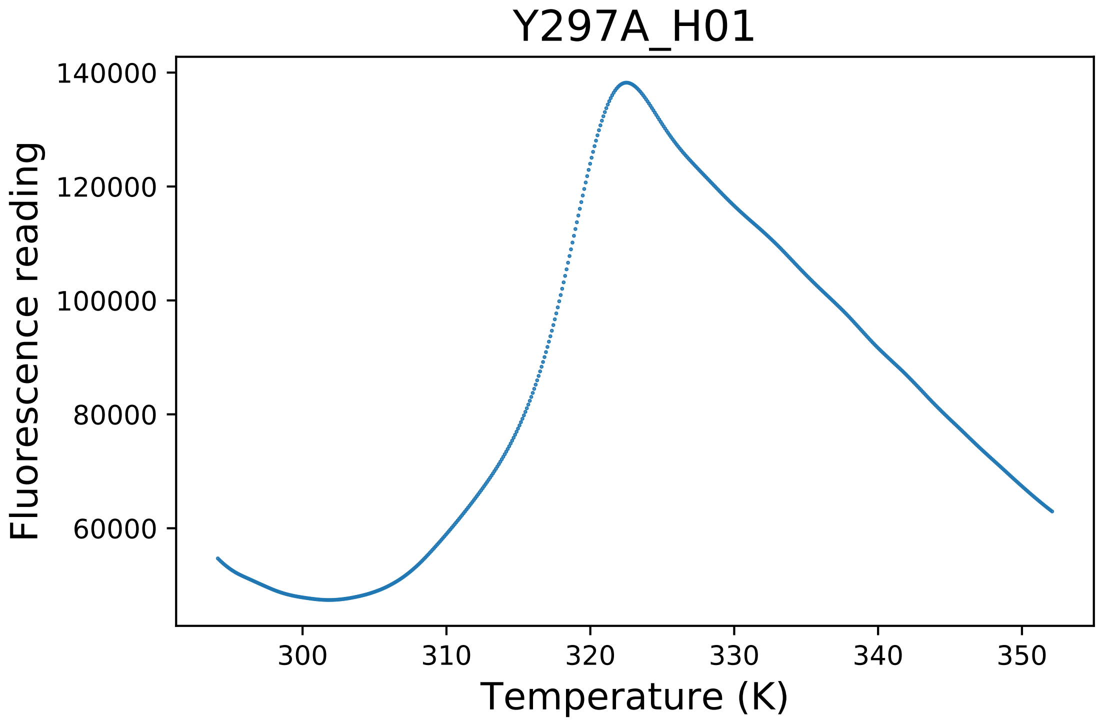

### Y297A_H02_dg.pdf

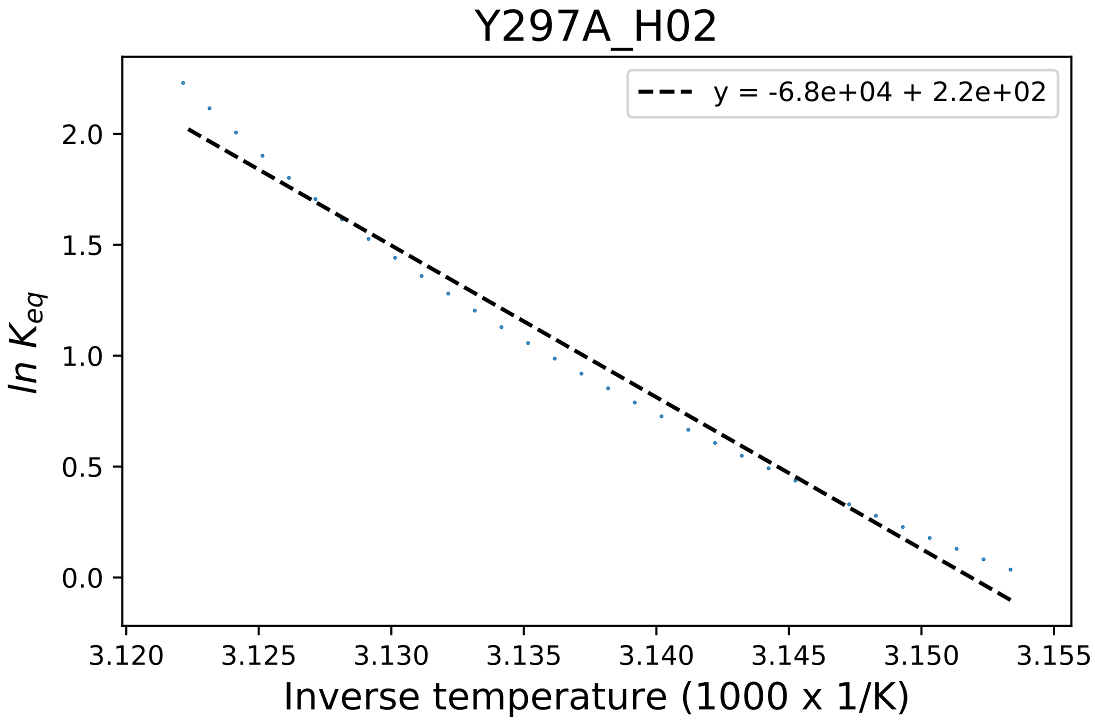

### Y297F_B02_derivative.pdf

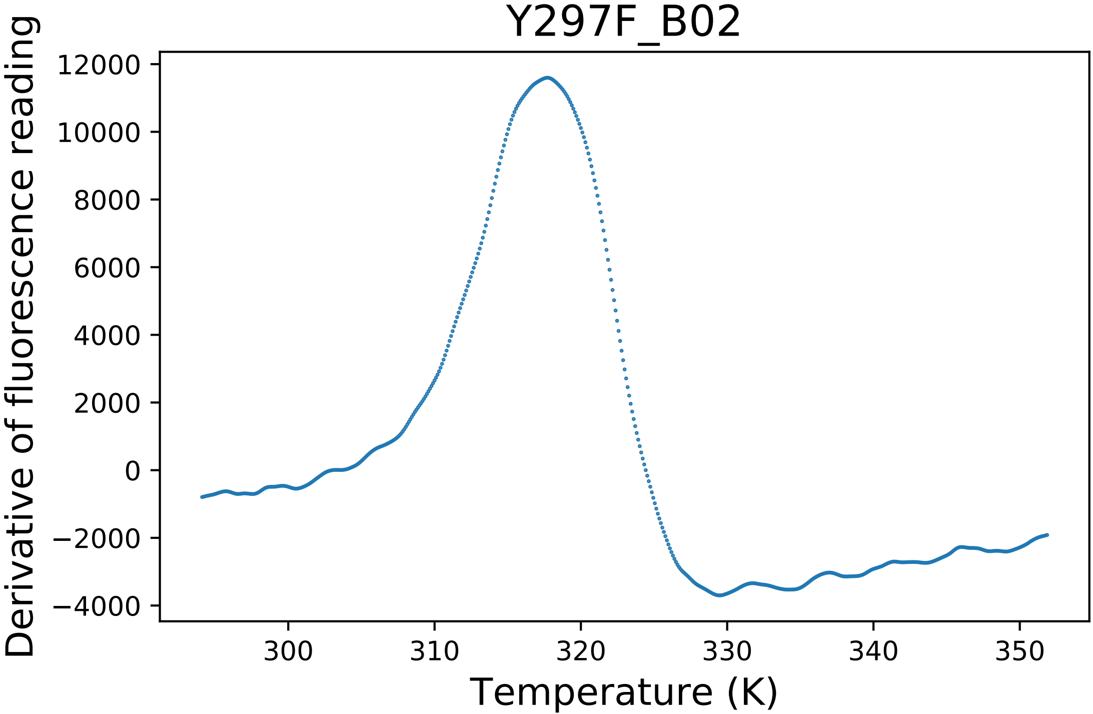

### Y297F_B03_derivative.pdf

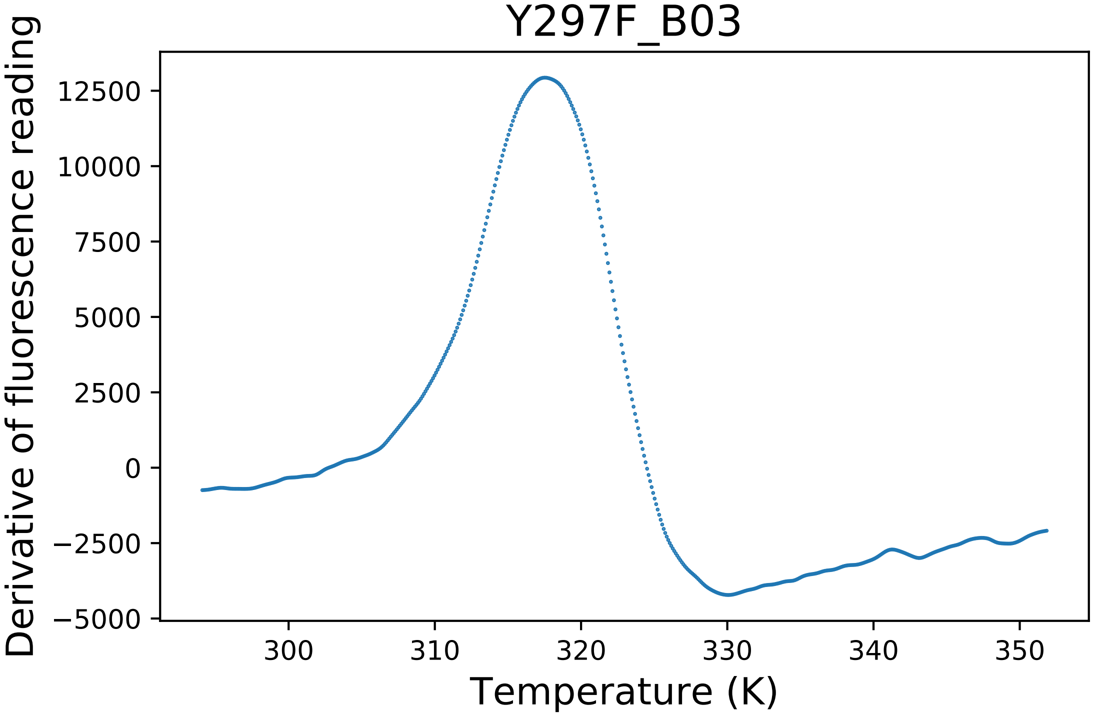
