## Supplementary material for "Evaluating molecular modeling tools for thermal stability using an independently generated dataset": SI_4: SI_script.ipynb

```
# %% [markdown]
# ddG calculation
This script calculates melting temperature, ddG and other thermodynamic quantities from input experimental data.

Unless specified, the temperatures in input files are assumed to be in Celcius. However, the output temperature is always in Kelvin while the energy unit is in kcal/mol.

The code is written in python3.

# %% [code]
import os
import numpy as np
import pandas as pd
import matplotlib.pyplot as plt
from scipy import stats
from scipy.optimize import curve_fit
from pylab import text

# scientific constant
R_gas = 1.987E-3 # kcal /(K mol)

# %% [markdown]
# Inputs and options
The user can specify the location at which is the experiment data is stored.

The user can also choose to skip figure output and speed up the analysis. Although it is possible to skip window-average smoothening, it is not advised due to the possible roughness of protein melting curve. Image output format is also supported, including but not limited to pdf, png and jpg.

If necessary, the temperature at which folding energy is calculated, is adjustable.

# %% [code]
src_dir     = 'raw_data'        # directory containing all experimental data
filename_master_csv = 'tm_t50.csv'   # file containing list of mutations and their respective csv and wells

flag_save_fig = True                 # output figures for all individual wells?
flag_smoothen = True                 # window-average smoothening on protein melting curve
N_kernel = 20                        # window-average size by temperature bin
image_suffix = 'pdf'                 # format of output figures

ddG_temperature = 298.               # the temperature at which the folding energy is calculated

# %% [markdown]
# Load input and output analyzed data
This script load data and analyze. Three csv files:
1. data_raw.csv       : combining all relevant well data into a single file
2. data_fitted.csv    : analyzed quantities categorized into individual wells
3. data_finalized.csv : analyzed quantities and their error categorized into individual mutants

# %% [code]
# Load and clean table csv
path_csv = os.path.join(src_dir, filename_master_csv)
df_table = pd.read_csv(path_csv)
del df_table['Unnamed: 3']

# store info
proteins            = df_table.Protein.values
wells               = df_table.wells.values
filenames_table_csv = df_table.csv.values

# organize wells label
wells_list = []
for i, entry in enumerate(wells):
    wells_list = []
    
    for position in range(int(entry[1:3]), int(entry[-2:])+1):
            wells_list.append('%s%s' % (entry[0], str(position).zfill(2)))

    wells[i] = wells_list

# %% [code]
# Construct data_raw.csv

df_raw = []
filenames_missing_csv = []
for i, filename_table_csv in enumerate(filenames_table_csv):
    # specific csv
    if filename_table_csv == 'S33-55':
        filename_table_csv = 'S31-54'
    
    filename_full = '%s_raw.csv' % filename_table_csv
    if filename_full in os.listdir(src_dir):        
        # load csv file
        path_csv_full = os.path.join(src_dir, filename_full)
        df = pd.read_csv(path_csv_full)

        # extract all wells for data
        df_list = []
        
        for well in wells[i]:
            df_list.append(df[df['Well Position'] == well])
        
        # include protein info in dataframe
        df_list = pd.concat(df_list)
        df_list['Protein'] = np.full(df_list.values.shape[0], proteins[i])
        
        # include label info in dataframe
        well_arr = df_list['Well Position'].values
        labels = []
        for j in range(well_arr.shape[0]):
            labels.append('%s_%s' %(proteins[i], well_arr[j]))
        df_list['Label'] = labels
        
        # append dataframes
        df_raw.append(df_list)
    
    else:
        # append missing csv filenames
        if filename_table_csv not in filenames_missing_csv:
            filenames_missing_csv.append(filename_full)
        
# convert multiple dataframes into single one
df_raw = pd.concat(df_raw, sort=True)

# show missing csv filenames
if len(filenames_missing_csv) != 0:
    print('The following data files are mising')
    print(filenames_missing_csv)
    
# save dataframe
filename_csv = 'data_raw.csv'
df_raw.to_csv(filename_csv, sep=',')

# %% [code]
# Construct data_fitted.csv

# The following mutant samples will not be analyzed for data collection due to low fluorescence reading
labels_remove = ['H376R_H09', 'H376R_H10', 'H376R_H11', 'H376R_H12',
                 'R243K_C09', 'N357A_H09', 'N357A_H10', 'N357A_H11',
                 'N357A_H12', 'M221A_E11']

# T, fluorescence, label
labels = list(set(df_raw.Label.values))
labels_iterable = [label for label in labels if label not in labels_remove]
label_num = len(labels_iterable)
labels_abnormal = []

# isolate temperature, fluorescence and their derivatives
temperatures, fluorescences, pf_full_range_list = [], [], []
pf_list, inverse_temperatures, ku_list, ln_ku_list = [], [], [], []
Tms, dGs = np.full(label_num, np.nan), np.full(label_num, np.nan)
dHs, dSs = np.full(label_num, np.nan), np.full(label_num, np.nan)

for i, label in enumerate(labels_iterable):
    # progress update
    print('\rProgress: %d/%d' % (i+1, len(labels_iterable)), end='', flush=True)
    
    # extract raw data
    df = df_raw[df_raw.Label == label]
    temperature = df.Temperature.values.astype(float) + 273.15
    fluorescence = df.Fluorescence.values.astype(float)

    # convolution smoothenning
    if flag_smoothen:
        kernel = np.ones(N_kernel,)/N_kernel
        temperature = np.convolve(temperature, kernel, mode='valid')
        fluorescence = np.convolve(fluorescence, kernel, mode='valid')
    
        # calculate Tm
        temperature_diff = np.subtract(temperature[1:-1], temperature[:-2])
        fluorescence_diff = np.subtract(fluorescence[1:-1], fluorescence[0:-2])
        fluorescence_derivative = np.divide(fluorescence_diff, temperature_diff)
        index_derivative_max = list(fluorescence_derivative).index(fluorescence_derivative.max())
        Tm = temperature[index_derivative_max]
        Tms[i] = Tm
    
    # plot fluorescence and derivative graphs
    if flag_save_fig:
        # dir for fig output
        dir_fig = 'figures'
        if not os.path.exists(dir_fig):
            os.mkdir(dir_fig)

        # fluorescence graph
        filename_fig = os.path.join(dir_fig, '%s_fluorescence.%s' % (label, image_suffix))
        fig, ax = plt.subplots(figsize=(6,4))
        ax.scatter(temperature, fluorescence, s=0.2)

        ax.set_title(label, fontsize = '16')
        ax.set_ylabel('Fluorescence reading', fontsize = '14')
        ax.set_xlabel('Temperature (K)', fontsize = '14')
        # ax.set_xlim(-30, 5)

        plt.tight_layout()
        plt.savefig(filename_fig, dpi=300)
        plt.close()
        
        # derivative of fluorescence graph
        filename_fig = os.path.join(dir_fig, '%s_derivative.%s' % (label, image_suffix))
        fig, ax = plt.subplots(figsize=(6,4))
        ax.scatter(temperature[:-2], fluorescence_derivative, s=0.2)

        ax.set_title(label, fontsize = '16')
        ax.set_ylabel('Derivative of fluorescence reading', fontsize = '14')
        ax.set_xlabel('Temperature (K)', fontsize = '14')
        # ax.set_xlim(-30, 5)

        plt.tight_layout()
        plt.savefig(filename_fig, dpi=300)
        plt.close()
    
    # crop window before and after Tm
    index_end = list(fluorescence).index(max(fluorescence))
    if index_end != 0:
        index_start = list(fluorescence).index(min(fluorescence[:index_end])) + 1
    else:
        index_start = 0

    index_middle = index_derivative_max # take center around Tm
    shrink_ratio = 0.5
    index_span   = int(min(index_middle-index_start, index_end-index_middle) * shrink_ratio)
    index_start  = index_middle - index_span
    index_end    = index_middle + index_span
    
    # calculate Tm and dG 
    if index_start >= index_end or index_start < 0 or index_end > len(fluorescence):
        labels_abnormal.append(label)
    else:
        # calculate dG
        pf_full_range = 1. - (fluorescence-fluorescence.min())/(fluorescence.max()-fluorescence.min())
        pf = pf_full_range[index_start:index_end]
        inverse_temperature = 1./temperature[index_start:index_end]
        ku = (1. - pf)/pf
        ln_ku = np.log(ku)

        # fit linear curve
        fit = np.polyfit(inverse_temperature, ln_ku, 1)
        fit_func = np.poly1d(fit)
        dH = -R_gas * fit[0]
        dS = R_gas * fit[1]
        dG = dH - ddG_temperature * dS
        dHs[i] = dH
        dSs[i] = dS
        dGs[i] = dG
        
        # save into lists
        temperatures.append(temperature)
        fluorescences.append(fluorescence)
        pf_full_range_list.append(pf_full_range)
        pf_list.append(pf)
        inverse_temperatures.append(inverse_temperature)
        ku_list.append(ku)
        ln_ku_list.append(ln_ku)
        
        # generate figures for pf and dG
        if flag_save_fig:            
            # linear dG graph
            filename_fig = os.path.join(dir_fig, '%s_dg.%s' % (label, image_suffix))
            fig, ax = plt.subplots(figsize=(6,4))
            ax.scatter(1000. * inverse_temperature, ln_ku, s=0.2)
            fit_label = 'y = %1.1e + %1.1e' % (fit[0], fit[1])
            ax.plot(1000 * inverse_temperature, fit_func(inverse_temperature), 
                    '--k', label=fit_label)

            ax.set_title(label, fontsize = '16')
            ax.set_ylabel(r'$ln\ K_{eq}$', fontsize = '14')
            ax.set_xlabel('Inverse temperature (1000 x 1/K)', fontsize = '14')
            # ax.set_xlim(-30, 5)

            plt.legend()
            plt.tight_layout()
            plt.savefig(filename_fig, dpi=300)
            plt.close()
            
# show labels of missing data
if len(labels_abnormal) != 0:
    print('The following label(s) has abnormal data.')
    print(labels_abnormal)
    
# construct dataframe
df_fitted = pd.DataFrame(labels_iterable, columns=['label'])

proteins_df = []
for label in labels_iterable:
    protein = label.split('_')[0]
    if 'WT' in protein:
        protein = protein[:-1]
    proteins_df.append(protein)

df_fitted['protein'] = proteins_df
df_fitted['Tm'] = Tms
df_fitted['dH'] = dHs
df_fitted['dS'] = dSs
df_fitted['dG'] = dGs

df_fitted.sort_values('label')

# save dataframe
filename_csv = 'data_fitted.csv'
df_fitted.to_csv(filename_csv, sep=',')

# %% [code]
# Construct data_finalized.csv

# final dataframe combining wells of the same protein into a single entry
dTm_means = []
dTm_stds  = []
Tm_means  = []
Tm_stds   = []
ddH_means = []
ddH_stds  = []
ddS_means = []
ddS_stds  = []
ddG_means = []
ddG_stds  = []

for protein in ['WT'] + list(set(proteins_df)):   
    # extract data from dataframe
    df = df_fitted[df_fitted.protein == protein]
    Tm = df.Tm.values
    dH = df.dH.values
    dS = df.dS.values
    dG = df.dG.values
    
    # store wildtype Tm, dG and dTm, ddG
    if protein == 'WT':
        Tm_wt_mean = np.mean(Tm)
        Tm_wt_std  = np.std(Tm)
        dTm_mean   = 0.
        dTm_std    = Tm_wt_std
        
        dH_wt_mean = np.mean(dH)
        dH_wt_std  = np.std(dH)
        ddH_mean   = 0.
        ddH_std    = dH_wt_std
        
        dS_wt_mean = np.mean(dS)
        dS_wt_std  = np.std(dS)
        ddS_mean   = 0.
        ddS_std    = dS_wt_std
        
        dG_wt_mean = np.mean(dG)
        dG_wt_std  = np.std(dG)
        ddG_mean   = 0.
        ddG_std    = dG_wt_std
    
    # store mutant dTm, ddG
    else:
        dTm_mean = np.mean(Tm) - Tm_wt_mean
        dTm_std  = ((np.std(Tm))**2 + (Tm_wt_std)**2)**0.5
        
        ddH_mean = np.mean(dH) - dH_wt_mean
        ddH_std  = (np.std(dH)**2 + dH_wt_std**2)**0.5
        
        ddS_mean = np.mean(dS) - dS_wt_mean
        ddS_std  = (np.std(dS)**2 + dS_wt_std**2)**0.5
        
        ddG_mean = np.mean(dG) - dG_wt_mean
        ddG_std  = (np.std(dG)**2 + dG_wt_std**2)**0.5
        
    # save into list
    Tm_means.append(np.mean(Tm))
    Tm_stds.append(np.std(Tm))
    
    dTm_means.append(dTm_mean)
    dTm_stds.append(dTm_std)
    
    ddH_means.append(ddH_mean)
    ddH_stds.append(ddH_std)
    
    ddS_means.append(ddS_mean)
    ddS_stds.append(ddS_std)
    
    ddG_means.append(ddG_mean)
    ddG_stds.append(ddG_std)

# create dataframe
df_final = pd.DataFrame(list(set(proteins_df)), columns=['label'])
df_final['mean Tm']  = Tm_means[1:]
df_final['mean dTm'] = dTm_means[1:]
df_final['mean ddG'] = ddG_means[1:]
df_final['mean ddH'] = ddH_means[1:]
df_final['mean ddS'] = ddS_means[1:]
df_final['std Tm']   = Tm_stds[1:]
df_final['std dTm']  = dTm_stds[1:]
df_final['std ddG']  = ddG_stds[1:]
df_final['std ddH']  = ddH_stds[1:]
df_final['std ddS']  = ddS_stds[1:]
df_final.sort_values('label')

# save dataframe to file
filename_csv = 'data_finalized.csv'
df_final.to_csv(filename_csv, sep=',')

# %% [markdown]
# Analysis visualization
1. Visualize the distribution of wildtype dG measurement (wt_dG_distribution)
2. Correlation between wildtype Tm and dG (correlation_wt_Tm_dG)
3. Corrleation between all mutants' dTm and ddG (correlation_dTm_ddG)

# %% [code]
# Plot the distribution of wildtype dG
dGs_wt = df_fitted[df_fitted.protein == 'WT']['dG'].values
bins = np.arange(dGs_wt.min(), dGs_wt.max(), 0.2)

# plot 
fig, ax = plt.subplots(figsize=(6,4))
plt.hist(dGs_wt, bins)

ax.set_title(r'Wildtype $\Delta$G distribution', fontsize = '16')
ax.set_xlabel(r'$\Delta$G (kcal/mol)', fontsize = '14')
ax.set_ylabel(r'Frequency', fontsize = '14')

plt.tight_layout()
filename_fig = 'wt_dG_distribution.%s' % image_suffix
plt.savefig(filename_fig, dpi=800)

# %% [code]
# Statistical assessment of correlation between wildtype Tm and dG
df_wt = df_fitted[df_fitted.protein == 'WT']
Tm_wt = df_wt.Tm
dG_wt = df_wt.dG

pcc_wt = stats.pearsonr(Tm_wt, dG_wt)
pd.DataFrame([pcc_wt], ['Tm and dG'], columns=['PCC', 'p value'])

# %% [code]
# Plot correlation between wildtype Tm and dG
fig, ax = plt.subplots(figsize=(6,4))
ax.scatter(Tm_wt, dG_wt, s=2)

text(0.3, 0.9, 'Pearson\'s Correlation = %1.2f' % pcc_wt[0],
     fontsize=12, ha='center', va='center', transform=ax.transAxes)

ax.set_title(r'Correlation between T${}_{M}$ and $\Delta$G', fontsize = '16')
ax.set_xlabel(r'T${}_{M}$ (K)', fontsize = '14')
ax.set_ylabel('$\Delta$G (kcal/mol)', fontsize = '14')

plt.tight_layout()
filename_fig = 'correlation_wt_Tm_dG.%s' % image_suffix
plt.savefig(filename_fig, dpi=800)

# %% [code]
# Statistical assessment of correlation between dTm and ddG
pcc_dtm_ddg = stats.pearsonr(df_final['mean dTm'].values,
                             df_final['mean ddG'].values)
pd.DataFrame([pcc_dtm_ddg], ['dTm and ddG'], columns=['PCC', 'p value'])

# %% [code]
# Plot correlation between dTm and ddG
fig, ax = plt.subplots(figsize=(6,4))
ax.scatter(dTm_means, ddG_means, s=1.5)

text(0.3, 0.9, 'Pearson\'s Correlation = 0.76',
     fontsize=12, ha='center', va='center', transform=ax.transAxes)

ax.set_title(r'Correlation between $\Delta$T${}_{M}$ and $\Delta\Delta$G', fontsize = '16')
ax.set_xlabel(r'$\Delta$T${}_{M}$ (K)', fontsize = '14')
ax.set_ylabel('$\Delta\Delta$G (kcal/mol)', fontsize = '14')

plt.tight_layout()
filename_fig = 'correlation_dTm_ddG.%s' % image_suffix
plt.savefig(filename_fig, dpi=300)

# %% [code]
```
